# TrailMix: Mixture Proportion and Placement Estimation of Ancient Hominin mtDNA Samples using a Pangenomic Reference

**DOI:** 10.64898/2026.11.21.768339

**Authors:** Jolan Boucher, Nicola A. Vogel, Viviane Slon, Peter Wad Sackett, Elena I. Zavala, Gabriel Renaud

## Abstract

Hominin mitochondrial DNA extracted from sediments or ancient environments is sometimes a mixture of multiple contributing sources. Researchers must subsequently computationally identify from which mitochondrial lineages the DNA sequences comes (e.g. Denisovan vs. modern human superhaplogroup) and ideally, quantify their abundance. Bioinformatics approaches described in the literature to achieve this involve in-house scripts or heuristics applied to tools designed for other datasets. However, such approaches have neither been fully tested nor do they provide credible intervals on model parameters. To address this, we introduce TrailMix, a C++ program for estimating contributing mixture proportions as well as individual source placements on the hominin mitochondrial tree for ancient environmental DNA. Using a pangenome reference graph to mitigate reference-related bias, TrailMix works well to accurately deconvolve mixtures of multiple mitochondrial lineages and place each contributor within its respective phylogenetic context. Crucially, it allows users to place minor contributing sources on the tree even on ancestral nodes without needing to reach consensus in the presence of multiple sources. TrailMix is part of the vgan suite of tools and is available at https://github.com/grenaud/vgan.

## 1 Introduction

DNA left behind by ancient individuals is often an invaluable source of information about, among other things, their demographic trends, migrational histories, burial practices, or even cultural and gender dynamics [21, 39, 2, 36]. Mitochondrial DNA is especially useful in this regard owing to the lack of recombination, providing a stable phylogenetic tree of maternal lineage in humans [41]. Ancient DNA (aDNA) has been recovered from skeletal or dental remains for a few decades now. However, in recent years, the potential of ancient DNA deposited in sediment or environmental samples to retrace human population history has emerged, leveraging the ability to recover aDNA even in the absence of visible skeletal or dental remains (e.g. [40, 49, 44, 48, 14, 13]). However, such samples are also difficult to analyze, because they not only share all the hallmarks of ancient DNA (i.e. fragmentation, contamination, deamination, and low coverage), which pose formidable difficulties in their own right, but furthermore a given sample may well contain DNA from a mixture of distinct contributing sources. Furthermore, we must distinguish between two related but different problems: i) the problem of detecting and quantifying present-day human contamination in aDNA by the researchers and handlers of ancient hominin samples who will invariably be of said species and ii) the problem of finding ancient modern human DNA mixed with DNA from archaic hominins like Denisovans or Neanderthals or even mixtures of ancient modern humans from different lineages.

For the first problem where the sample contains a mixture of present-day and ancient DNA, a number of programs are currently available to estimate the contamination rate on the mitochondria. One such program is schmutzi, which employs a Bayesian model to estimate the deamination rate and proportion of endogenous material in a two-source (endogenous plus contaminant) mitochondrial aDNA mixture, while jointly calling a consensus on both the endogenous and the exogenous material, iteratively [35]. Another widely used program is contamMix [11], which though widely used in the aDNA community [35, 11, 28, 43, 10, 30, 16] has never been properly published in its own right as a method. contamMix uses Gibbs sampling techniques to estimate the contamination rate.

The above programs are useful, but they may not be applicable to the second problem, which is the deconvolution of ancient hominin DNA. For example, both schmutzi and contamMix are applicable only in the case of a two-source mixture where one source is negligibly damaged (the contaminant) and the other non-negligibly damaged (the endogenous). The latter program further stipulates that the endogenous proportion of the sample be greater than 50%, and moreover it requires a consensus as an input — which it assumes to exactly reflect the endogenous mitotype — and it cannot handle ambiguous nucleobase encodings which are ubiquitous for consensus FASTA files called from ancient samples. Due to the assumptions made by the programs, they are not readily applicable to the second problem i.e., deconvolution of hominin DNA in sediments.

For handling mixtures of modern samples, MixEMT deconvolves mtDNA mixtures using a maximum likelihood estimation approach after alignment to the linear rCRS reference genome, reporting both mixture proportions and haplogroup assignments for each contributor. However,mixEMT does not allow the user to specify as a hyperparameter the number of contributing sources in the mixture if they so desire [46, 1]. EMMA2 can deconvolve mtDNA mixtures using ambiguous nucleotide information, but only works off of a called consensus (and therefore it cannot estimate the contamination rate) [7]. HaploCheck can detect human contamination in an mtDNA sample, and reports the contamination rate if two sources are detected, but it does not provide a haplogroup call for the contaminant. The tool works by looking at the allele frequency of polymorphic sites in the sample with respect to the reference, and taking into account phylogenetic distances of these polymorphisms to try to differentiate between genuine contamination and potential heteroplasmy or private mutation [47]. None of these aforementioned tools is capable of inferring that an ancestral state is present in the sample or deal with highly degraded DNA.

To tackle the second problem, for hominin mtDNA recovered from sediments, fragments have been assigned to modern humans, Neanderthals or Denisovans by counting those that carry lineage-specific states at diagnostic positions [48], and kallisto [3], a pseudoalignment-based tool originally designed to quantify transcripts from RNA-seq data, has been used to identify the reference mitogenomes to which the hominin fragments of a sediment sample are most similar [44, 48]. kallisto has since also been applied to the poorly preserved and heavily contaminated mtDNA of the Teshik-Tash 1 Neandertal [42]. These approaches used for sediments summarize the hominin fragments by predefined groups or by the reference sequences to which they are most similar, which requires the groups of interest to be defined in advance, and they do not jointly estimate the proportions and phylogenetic placements of the contributors together with their uncertainty. Another algorithm called soibean [45] determines the phylogenetics placement and relative abundance for clades of eukaryotic DNA, but its ability to handle highly similar clades like hominins is undocumented, as its seeding algorithm is dependent on nodes unique to certain species. Also, pathphynder [25] has been used for phylogenetic placement of sedaDNA, but cannot separate between different components.

However, the primary challenges in developing a method for joint mixture proportion and phylogenetic placement estimation, particularly one which addresses the gaps described above, are twofold. For one, the parameter space that must be explored is vast, primarily due to the combinatorial explosion involved in a mixture of haplotypes. The second challenge is that robust quantification of uncertainty is absolutely crucial, especially given the inherently sparse and noisy nature of aDNA data. Both of these issues point towards an approach based on Markov Chain Monte Carlo (MCMC) sampling methods. MCMC sampling methods are perhaps the most widely used approaches for inferring phylogenies, as the space of possible trees that could relate some given sequence data is enormous. The task of phylogenetic placement, i.e. placing data at its most probable source location on a fixed phylogenetic reference tree, is less common in general, and no extant program of which we are aware uses MCMC methods to do this. However other Bayesian methods are used in this context, e.g. pplacer [26]. In addition, such a method should handle the high similarity within hominin mitogenomes as these are not species separated by several million years.

Up to now, accurate mixture proportion and phylogenetic placement estimation from arbitrary hominin mtDNA mixtures in the absence of an endogenous reference has been an unsolved problem, especially for samples where little data aligns to the mitogenome. To address this we introduce TrailMix, a C++ subcommand of the vgan suite of tools for pangenomes. TrailMix infers approximate phylogenetic placement and source contribution proportions via MCMC sampling methods from next-generation sequencing (NGS) data for hominin single-source or mixture samples (see graphical abstract 1). By using an aDNA-aware aligner that takes deamination events into account during the mapping process, as well as by using a pangenomic reference to mitigate reference-related biases during inference, the program is capable of drawing accurate inferences even for very low-coverage samples. Our method is aimed at profiling sediments with very low coverage and can identify that a given sample is a mix of Neanderthals, Denisovans and ancient modern humans and estimate their relative proportion at a coverage of even 1X. For ancient modern humans, we can distinguish between macro-haplogroups (e.g. L2 versus M) but not provide specific haplogroups at very low coverage. Although present-day human contamination estimation remains an important application, in this work we focus on mixtures in which every contributor is ancient, where several hominin individuals or even several hominin groups may have contributed DNA to the same deposit; all sources are therefore modeled as carrying ancient DNA damage. The special case of a present-day human contaminant is not considered. TrailMix accepts as input either FASTQ reads (single-end or paired-end, possibly interleaved), GAM files, or consensus FASTA files if the input is single-source. Additionally the user may provide estimated nucleotide misincorporation matrices for the input sample to be used by the inference model for damage, as discussed in Section 2.2.

## 2 Methods

### 2.1 Reference Pangenome Graph Construction and Phylogeny Inference

We used the tool IQ-TREE (version 1.6.12, HKY+F model) for the tree topology estimation and the marginal ancestral state reconstruction [29], after aligning the 64 mitogenomes with MAFFT [18]. The graph comprises 126 sequences (64 derived, 62 ancestral) and has 5,425 nodes and 6,664 edges. This includes 7 archaic hominins (4 Neanderthals and 3 Denisovans). Our updated graph was constructed with PGGB [12]. The corresponding reference tree has 64 tips and 125 branches. We also include scripts and instructions for users to build their own bespoke databases. The exact composition of the database is found in Supplementary Section S2.

### 2.2 Inference

The inference model for TrailMix is adapted from that of soibean [45]. We generally follow the notation introduced therein.

We begin by defining our model, *M* = (***β***, **Θ**). Here **Θ** are mixture proportions (in the data, not in the sample) expressed as a point on the probability simplex of dimension *k* − 1, where the number of contributing sources *k* is fixed as a hyperparameter. These are proportions of fragment counts (Θ*_i_* is the probability that a given fragment originates from source *i*), not of nucleobase counts, and that is a distinction with a difference since the fragment length distribution will tend to be shorter for more ancient sources. The vector ***β*** denotes (continuous) phylogenetic placements on our fixed reference tree as fractions along the branch from zero to one, with zero being ancestral and one derived. All models are *a priori* equiprobable, meaning that Bayes’ Rule tells us

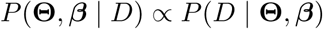

That is to say the likelihood function and posterior are equal up to a normalizing constant. Here *D* is the set of aligned reads after PCR duplicate removal has been performed (duplicates should be removed by the user before running TrailMix). Briefly, we seek to sample from *P* (*D* | **Θ***, **β***) to estimate the distributions over **Θ** and ***β***. Since duplicates have been removed it is reasonable to model each fragment as independent:

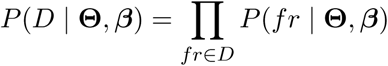

A given fragment *fr* originates from exactly one of the *k* possible sources. Let *P* (Θ*_i_, β_i_*) denote the probability that a fragment is from source *i* and is sampled from the mitogenomic sequence corresponding to phylogenetic placement *β_i_*. Rather than assign individual fragments to individual sources, we compute the likelihood of a fragment by marginalizing over each source from which it may have arisen:

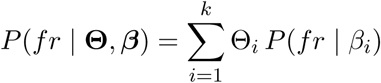

To calculate the likelihood of a branch placement on the tree, we make the further simplifying assumption of treating each base *b* of fragment *fr* as independent, allowing us to multiply the probabilities of each nucleotide observation for each fragment:

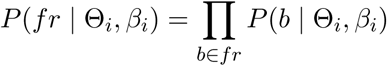

We note that this assumption of independence among bases is not entirely accurate from a biological perspective as there are in fact strong correlations between certain base observations, especially in mtDNA where linkage disequilibria persist due to lack of recombination. However, these dependencies would be very difficult to model and would also prohibit parallelization of the program, exacerbating issues with runtime. We acknowledge that future work to improve modeling of base dependencies, e.g. dinucleotide frequencies, may improve the model.

An aligned base is either correctly aligned or incorrectly aligned. These are disjoint possibilities. Let us denote the event that the base is correctly aligned *M*. Then by the law of total probability we have

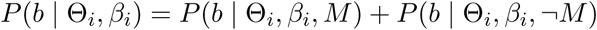

We consider the event ¬*M* to include not only the case where the read is mitochondrial in origin but is mismapped, but also where it is a nuclear mitochondrial DNA segment (or NuMT) of autosomal origin:

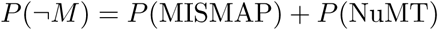

The probability of mismapping is readily computed from the mapping quality score for the read in question, which is reported on the PHRED scale:

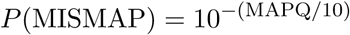

The probability of the second possibility, i.e., that the aligned read is a NuMT from the autosomes, can be calculated based on the GenMap mappability score for the base in question as mapped against a reference comprising all chromosomes. These are identical to what was done for the HaploCart model [31, 37]. In the current implementation, however, the probability of mismapping enters the likelihood only through the bases that are not supported by the tree node under consideration (see below), and the mappability term is not used: the likelihood of a supported base is computed from the mutation, damage and sequencing error model described below. Adding these terms to every base of a fragment would, for fragments with a low mapping quality, drive the fragment likelihood to its lower bound on every branch, so that such fragments would no longer carry any information.

Branch placements are modeled as fractions (between zero and one) along a branch between the derived node *N_D_* and ancestral node *N_A_*. Even though the putative placement is some fraction between these two nodes (,and may well be on a branch representing a huge amount of evolutionary time) our reference will only contain the sequences at the termini. Therefore we will be evaluating the likelihood at the two nodes terminal to the branch, and then compute a weighted average. Let us suppose we have fixed one such tree node, call it *N_t_*, which will be either ancestral to or derived from the putative placement. Every node in the tree represents a full mitogenome sequence embedded in the reference graph, which translates to the concatenation of a large number of pangenome nodes. Therefore an arbitrary base *b* will either be supported by *N_t_* (i.e. the haplotype sequence represented by *N_t_* in our reference graph traverses the node on which *b* is located) or else *b* is unsupported by *N_t_*. In the latter case the first base of each aligned segment on such a node, and every fourth base thereafter, incurs a penalty commensurate with the likelihood of a sequencing error for a base with base quality 50 on the PHRED scale, while the remaining bases are assigned the probability of a correct base call multiplied by the probability that the read is correctly mapped. Since most nodes not shared by all haplotypes are single-base variant nodes, in practice almost every unsupported base at a variant site incurs the penalty. The stringency of this penalty term was empirically derived through experimentation, and constitutes a minor departure from the model introduced by HaploCart.

In case *b* is supported by the tree node *N_t_* we have several potential explanations for how the observed base in the read can have arisen, given that it has aligned to the base in the graph. There could have been a private mutation from the reference to the sample. Alternatively, seeing as this is an aDNA sample there may have been a deamination event from the reference base to the read base. Also, there may have been a sequencing error from the sequencing machine, leading us to observe the incorrect base. To take all of this into account the data generation mechanism has to be viewed as a Markov process, with hidden states representing the outcome of these different events at each stage in the steps leading up to the aligned base observation (Figure 2).

**Figure 1:**
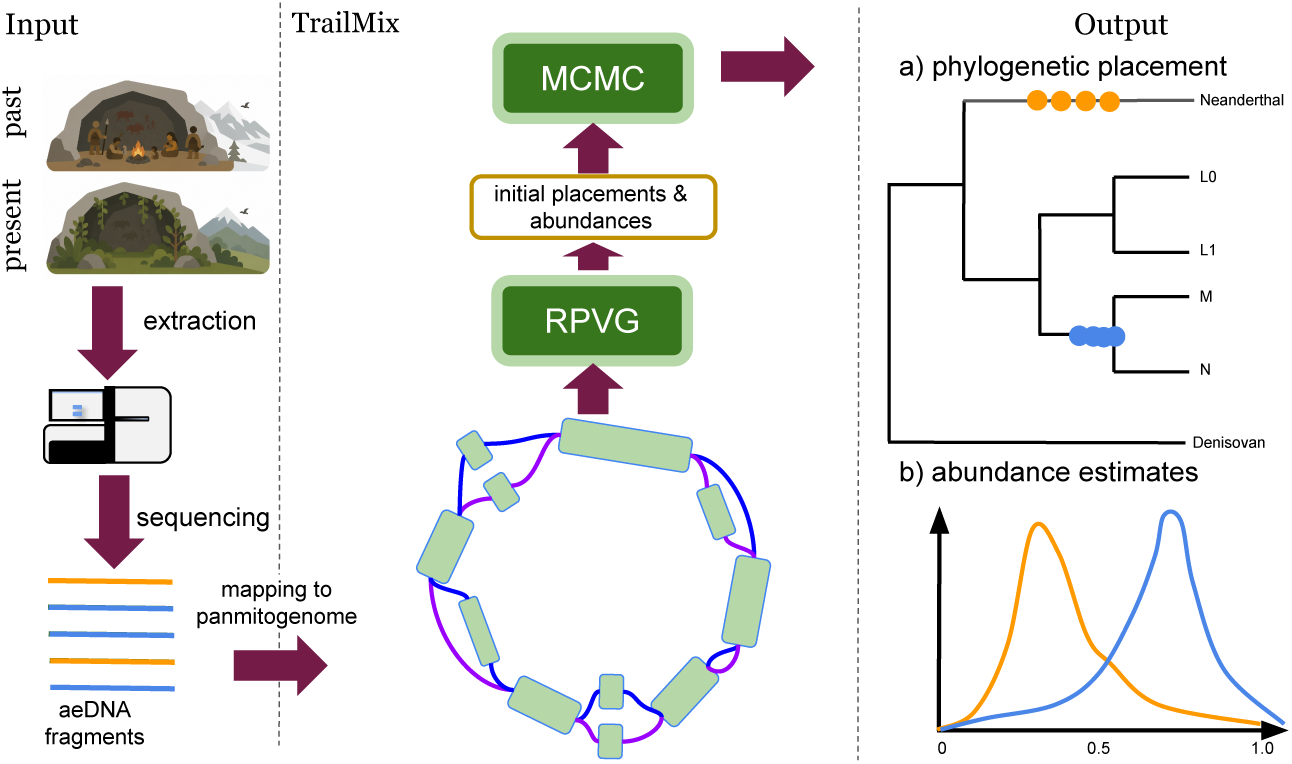
When ancient environmental DNA (aeDNA) is extracted from a site with potential hominin presence, it may contain a very small fraction of hominin DNA. In TrailMix, the sequenced aeDNA fragments are aligned to a panmitogenome for hominins containing both archaic hominins (Neanderthals and Denisovans), modern humans, as well as reconstructed ancestral states. RPVG provides an initial placement and abundance estimates. These initial values are refined on the phylogenetic tree via MCMC.

**Figure 2:**
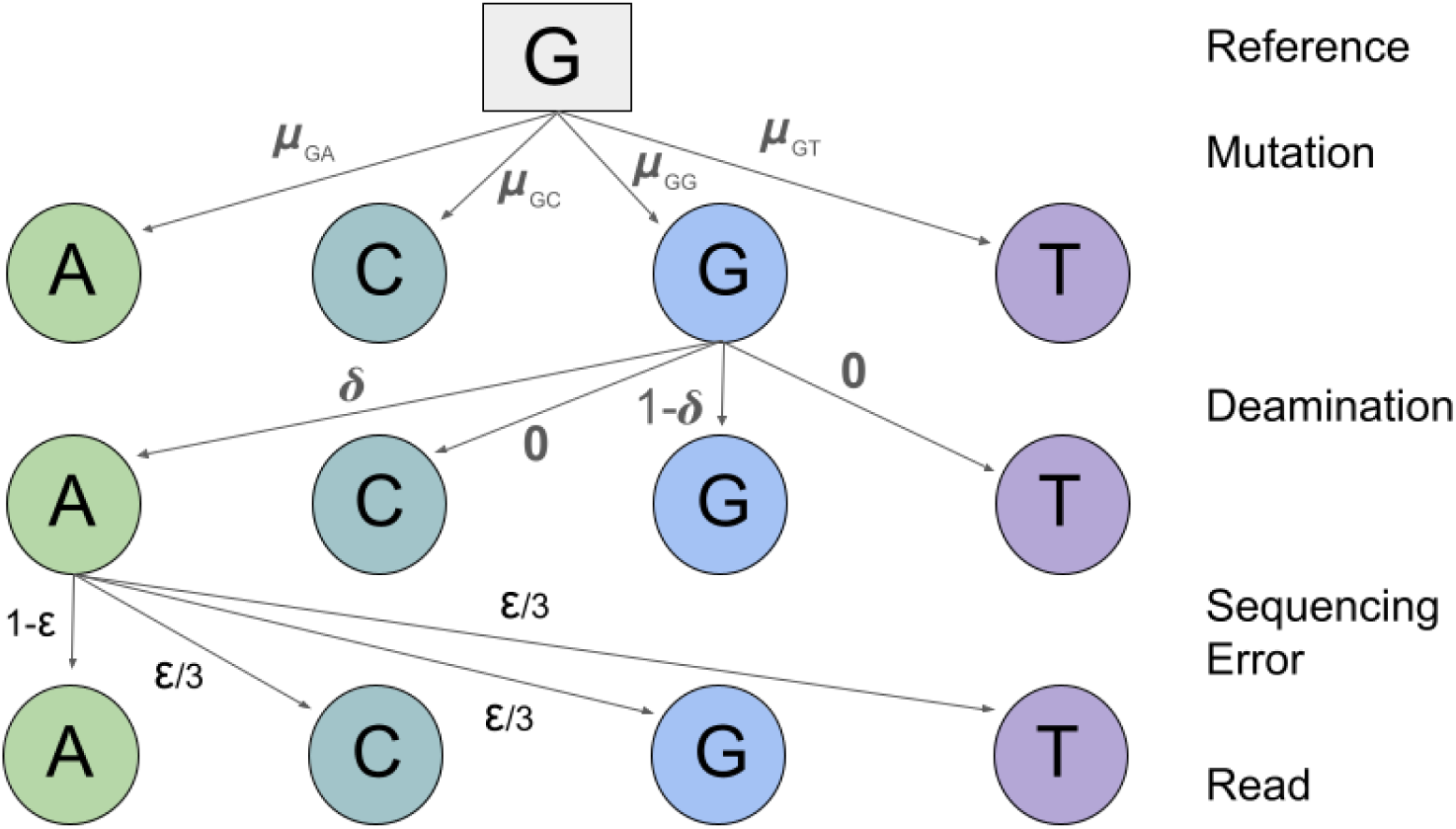
The observation of a correctly aligned nucleobase is modeled as the final state of a hierarchical Markov process with multiple hidden states corresponding to steps in the data-generating process from canonical reference base to observed base. The probability of observing the aligned base (the final state) is obtained by marginalizing over the joint probabilities for each path in the process.

The first event that we must consider is that a private mutation from the reference to the sample has occurred. We compute the probability of this event via the HKY substitution model (the model under which our tree was constructed) given the base in the reference, the base in the read, the length of the branch on which the putative placement is situated, and the fraction along this branch at which the source is placed for the given Metropolis-Hastings sample [17]. Note that, of course, different putative mutations will have different likelihoods under this model, e.g., transitions will be far more probable than transversions, as one would expect for mtDNA.

The second event in our Markov process is the potential for a deamination event from the reference base to the read base. This is only possible if the reference base is a C or G, and the read base is T or A, respectively. More precisely, if we let *b_d_* denote the observed base in the fragment (post-damage) and let *b_s_* denote the base in the original genome, then the deamination rates are given by

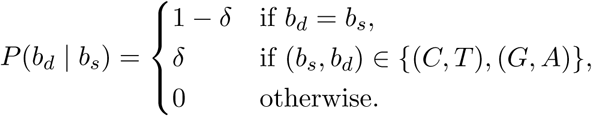

where *δ* depends on the base position within the fragment. *δ* will depend on the read base and reference base but also the position of the base on the input fragment, taken from a model implemented in ROHan [34]. The rate matrices dictating *δ* are provided by the user at runtime. As such, TrailMix is compatible with any type of DNA library preparation (single-stranded or double-stranded, either with or without enzymatic treatment to remove uracil residues).

The final hidden state in our Markov process is the potential for a sequencing error, denoted *ɛ*. *ɛ* is quickly computed based on the base quality score provided by the sequencing machine on the PHRED scale [8]. In particular if a sequencing error has occurred we disregard any potential biases in the sequencing machine and consider it equally likely to see any of the three alternate bases being called, so we have:

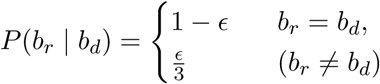

where *b_r_* is the base in the read called by the sequencer. Following HaploCart we assign probability 0.02 to observed gaps due to indels, as this is roughly the background frequency of indels for a given base in the human mitogenome [37]. Now although the putative source placement is in general somewhere along the branch, we only store the mitogenome sequences of the terminal tree nodes. Therefore we need to compute separately the probability of observing the read base given the reference base for two different reference bases, namely the ancestral base *b_A_* and the derived base *b_D_*. Now the putative placement *β_i_* represents the fraction along the branch (from ancestral to derived) from which we are sampling. Thus if the branch has length *l_b_*, the distance of the putative placement to ancestral node *N_A_* is *l_b_β_i_* and the distance of the putative placement to derived node *N_D_* is (*l_b_* − *l_b_β_i_*).

So the question then becomes, for a given branch on the reference tree how can we calculate the likelihood for arbitrary *β_i_* ∈ [0, 1]? The answer is that we compute the likelihood of the fragment (the product over its bases above) at each of the two nodes and calculate a weighted average of these terms, where the weighting factor is the proportion along the branch of the putative placement:

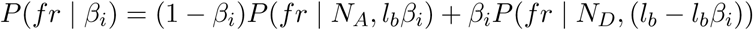

Each of the two fragment log-likelihoods is bounded to [log 10*^−^*^10^, 0] before the weighted average is taken.

#### 2.2.1 Metropolis-Hastings Sampling

Given the size of the parameter space, it is clear that exhaustively computing the posterior of every possible model is out of the question, and sampling methods are therefore indicated. TrailMix uses Metropolis-Hastings sampling, a particular version of Markov Chain Monte Carlo (MCMC) estimation, to generate samples from the posterior distribution in an efficient way [27]. We again follow soibean in using the topology of our reference tree to guide our sampling choices when it comes to the phylogenetic placements *β*.

More specifically, at every MCMC iteration we select one source uniformly at random and move its placement along the reference tree by a Gaussian step (in units of branches), whose standard deviation *λ* is sampled from a mixture distribution (empirically derived):

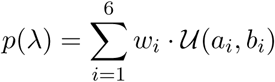

where:

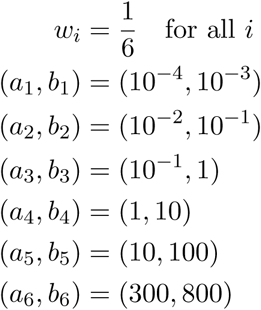

Given this sampled step we then move the state of the phylogenetic placement *β* the appropriate amount along the tree, uniformly randomly selecting a branch wherever there is a choice. For more details about movement along the tree please see [45], Supplementary Material Section 2.3. To sample the proposed proportion vector Θ*_i_*_+1_ we draw each component independently from *N* (Θ*_i_,* 0.1) truncated to [0, 1] and then normalize so the components sum to unity. This also follows Vogel *et. al.* [45]. The placement and proportion proposals are accepted or rejected jointly. Since this proposal is symmetric, no Hastings correction is required.

The crucial benefit of our MCMC approach to inference is that it naturally lends itself to clear and unambiguous confidence estimates within a phylogenetic context. Samples which are easily placed by the program will exhibit very narrow intervals of search along the tree, while difficult samples (due to low-coverage, unknown private variation, etc.) will lead to a wide search across the tree, reflecting the wideness of the underlying posterior distribution over models. The user can therefore discern the confidence in the program output and make their interpretations accordingly. Moreover, the program comes with detailed diagnostic reports on the MCMC chains for the user to examine; these diagnostic reports are discussed in [45], Supplementary Section 2.3.1. Programmatically, navigating the reference tree topology is accomplished via functionality in the SPIMAP library for C++ [32].

#### 2.2.2 Initialization

The parameter space of the TrailMix model is vast. Each contributing source to the input sample is modeled as originating from one of the 125 branches of our reference tree, at any position along that branch, and on top of that, we also estimate the contributing mixture proportions as well as the degree of mutational distance from the reference mitogenome in our graph. For this reason, it increases the odds that we will converge if we initialize each MCMC chain at a reasonable point in the likelihood landscape. If we start our search in a completely incorrect region of the tree, we might dwell in suboptimal areas without ever reaching the maximum likelihood abundance and phylogenetic placements. For this reason, TrailMix uses an initialization strategy which first runs the program RPVG on the sample in question [38]. Briefly, RPVG is a pangenome-based approach for estimating transcript abundance for RNA-seq which uses an expectation-maximization algorithm in a similar to kallisto. More details about the RPVG initialization procedure are provided in Supplementary Section S3. All chains of a run start from the same initial state.

#### 2.2.3 Whitelisting Branches

To reduce the memory demands and the running time of TrailMix, the model by default only explores branches within a given phylogenetic depth from one of the nodes identified under RPVG initialization. In particular, the user can provide a maximum phylogenetic depth parameter (--depth, 10 edges by default; --depth -1 lifts the restriction), and we only allow MCMC chains to explore branches off of nodes which are within that fixed depth away from a haplotype flagged in the Θ quantification (haplotype-transcripts) run of RPVG, i.e. a haplotype to which RPVG assigns more than one transcript per million. All results in this manuscript were obtained with a default depth of 10, which left between 32 and all 125 branches of the tree available to the chains, depending on the sample.

### 2.3 Visualization

To help the user interpret TrailMix results, we include with the software a utility script for ease of visualization.

#### 2.3.1 Visualizing Phylogenetic Placement

We provide for the user’s convenience a Python script (*plot trailmix trees.py* ) to visualize the posterior sampling across the tree in a way which should be useful and interpretable.

The script takes as input the output prefix of a TrailMix run, from which it reads the MCMC files of every chain, the run parameters and the reference tree file in Newick format [9]. For each source, every branch visited by the chain is drawn with a ‘swelling’ showing the posterior density of the position along the branch, and a dot at the median position whose area is the posterior probability of that branch; the tip of the best-supported branch is shown in bold, or its clade is shaded when that branch is internal. In the figures of the main text, every named lineage (Denisovans, Neanderthals, haplogroups L0–L6, macrohaplogroups M and N) that forms a single clade of the tree and in which no post-burn-in iteration placed a source is collapsed into a single bar, labeled with the lineage and its number of tips; the supplementary figures show the full tree. Branches are matched between the tree used by TrailMix and a tree re-rooted for display by the set of tips they separate, so the display rooting can be chosen freely; in the figures of this manuscript the tree is rooted on the branch leading to the Denisovans. Each run may need to be customized to an extent to best visualize the results, so we leave options open to the user. For example, the user can draw the tree with all branch lengths set to unity or shorten very long branches, which can be helpful when long branches would otherwise obscure more salient portions of the tree. We also allow the user to provide a list of clades to be sure to retain for the plot, so that the user has control over what to emphasize.

#### 2.3.2 Visualizing Mixture Proportions Θ

The same script generates a density plot for each source proportion Θ*_i_*, with the median and the 95% credible interval of each source indicated below the densities and in the legend. It also draws the trace of the proportions along the chain and compares the posterior mean proportions across chains. Chains are not pooled: the chain shown is the one with the highest minimum effective sample size over the sources and over the placement (position along the branch and location on the tree) and the mixture proportions.

We caution the user that, since sources are exchangeable, their labels may differ between chains and may occasionally ‘flip’ during the trajectory of a chain (e.g. in a two-source mixture, for source 1 to become source 2, and source 2 to become source 1, and therefore for each estimated Θ*_i_* to become 1 − Θ*_i_*). In our two-source simulations we observed such a flip in about 6% of the chains. The script therefore numbers the sources by decreasing mean proportion, and the chain-selection criterion above rarely selects a chain in which a flip occurred.

### 2.4 Experimental Design and Sample Generation

We conducted three experiments for our tool to ensure its robustness and the soundness of the model specification. Across all experiments we strove to represent the full spectrum of human mitodiversity as far as possible, as one of the primary benefits conferred by the pangenome reference graph is the mitigation of reference-related biases. Each simulation experiment featured a set number of contributing sources. Simulated samples for evaluation of our pipeline were generated using Snakemake [19], and further details about the generation of samples is provided in Supplementary Section S1.

Our first experiment aims to show that TrailMix can correctly place single-source hominin samples on the mitochondrial phylogeny regardless of provenance, as well as to demonstrate that accuracy in placement increases as a function of coverage depth, and to get a sense of the depth of coverage typically required for high-confidence placements. We do this for five simulated samples: one Neanderthal, one Denisovan, and three modern humans belonging to haplogroup L2 and to the macrohaplogroups M and N (see Supplementary Section S1). None of these five mitogenomes is part of our reference graph, so each must be placed relative to its closest relatives in the reference. For each sample, we run the program at varying levels of coverage to demonstrate that the predicted placement of the sample on the reference tree will become more and more certain (i.e. the posterior will be more narrowly concentrated around the correct location on the tree) as coverage increases.

In the two-source experiment we turn our attention to samples in which both sources exhibit the chemical damage patterns characteristic of ancient DNA, as for instance might be found in sedaDNA. Each of the five samples above was mixed with a second, more weakly damaged, modern human mitogenome at 21 mixture proportions between 1% and 99% and at mean coverage depths between 0.5X and 32X. For these samples we also compare the mixture proportions estimated by TrailMix to those of its RPVG initialization and of kallisto [3], which has previously been used to quantify hominin mtDNA in sediments [44].

Finally, we apply TrailMix to empirical sediment samples from two publications [48, 44] whose hominin mtDNA was characterized with independent methods, which allows us to compare our inferences with those of the original authors.

## 3 Results

Here we show results of our program on various input samples to demonstrate its robustness. We begin with simulated samples, from which it is possible to compare to a well-established ground truth. The downside to simulations is that they can only approximate what is encountered in reality, which is why we will then showcase the results of TrailMix run on empirical samples.

### 3.1 Simulated Data

Our simulated samples are based on five published mitogenomes which are not part of our reference graph: the Neanderthal Chagyrskaya 8 (ref: MK388903.1 from [23]), the Denisovan Denisova 21 (ref: MT576653.1 from [4]), and three modern humans, one of haplogroup L2 (ref: AF381981.1 from [22]), one of haplogroup D4a1d in macrohaplogroup M (ref: FJ951517.1 from [6]) and one of haplogroup T2a1 in macrohaplogroup N (ref: KY671094.1 from [24]). Their closest relatives in our reference are the Les Cottés Neanderthal, Denisova 20, L2a1d, D4a3a and T1a1a1, respectively, so that TrailMix must place each of them on a branch of the tree rather than on an exact reference sequence. Reads were simulated using gargammel 1.1.4 [33] with the fragment size distribution of the ATP2 sample from [15] and its measured ancient damage rate (see Supp. Fig. S1). The rates of damage were empirically derived using bam2prof from [20] obtained by aligning the simulated fragments to the rCRS (ref. NC 012920.1) using SHRiMP version 2.2.2 [5], and the samples were downsampled to the desired mean coverage depths. For the two-source samples, each of the five mitogenomes was combined with a sixth simulated ancient mitogenome, described in Section 3.1.2. Further details about the generation of samples are provided in Supplementary Section S1.

#### 3.1.1 Single-source Samples

Here we illustrate TrailMix results on single-source simulated samples, in order to demonstrate the robustness of our method in this case. We will then generalize to simulated mixture samples in Section 3.1.2. Figure 3 shows TrailMix results on two single-source samples, the Neanderthal Chagyrskaya 8 and the modern human D4a1d. The damage rates estimated as described above along with parameters --iter 100000 -k 1 --chains 5 were used in TrailMix.

**Figure 3:**
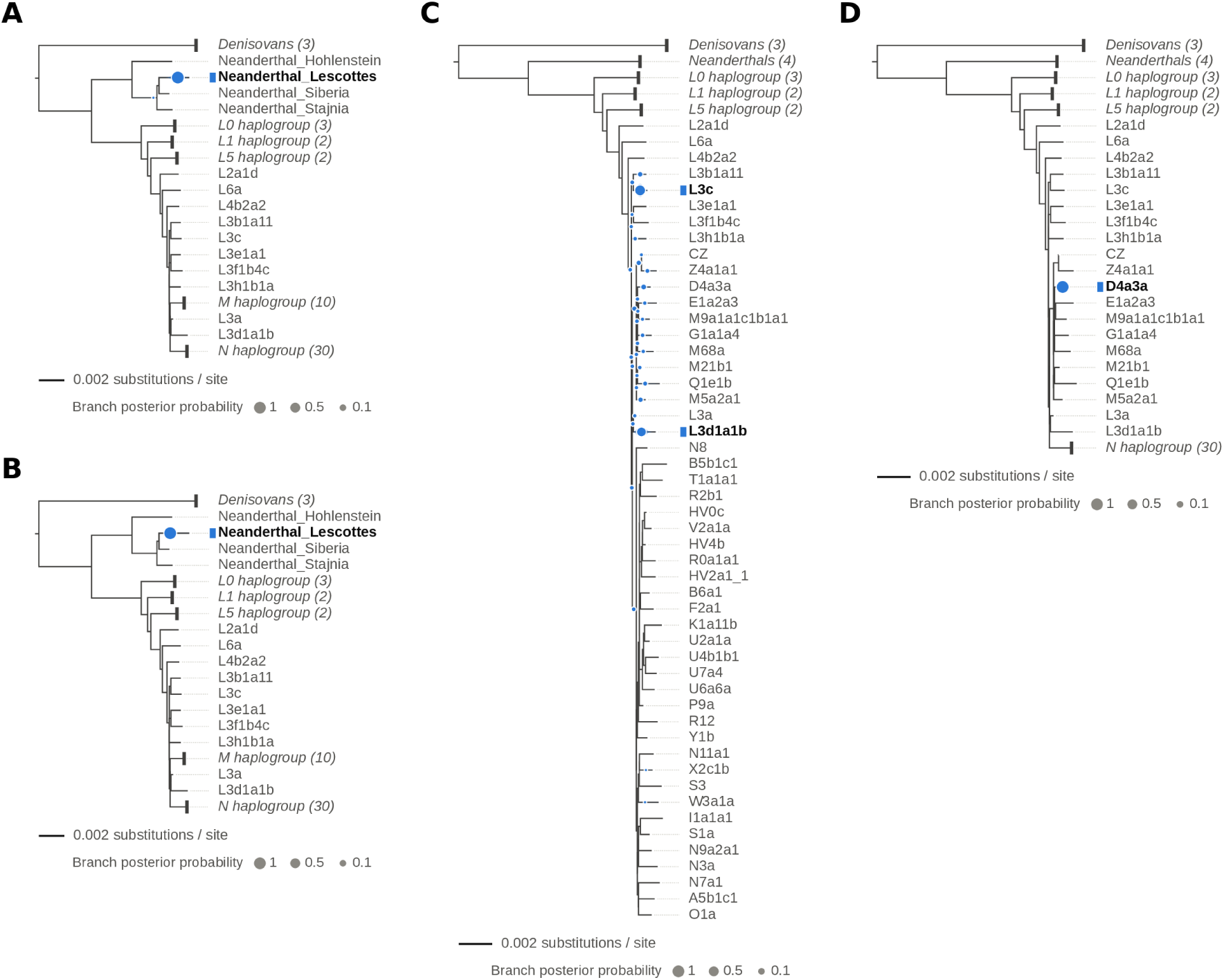
Examples of single-source simulated samples: the Neanderthal Chagyrskaya 8 [23] (A, B) and the modern human D4a1d [6] (C, D), neither of which is part of our reference. Chagyrskaya 8 should fall close to the Les Cottés Neanderthal and D4a1d close to D4a3a in our tree. These plots represent where the MCMC has dwelled for the best of five chains: the area of a dot is the posterior probability of the branch, the swelling along the branch the posterior over the position on that branch, and the best-supported tip is in bold. We have downsampled the original file to test the robustness. Chagyrskaya 8 is placed on the Les Cottés branch already at 0.5X, and its position along the branch is narrowed down at 4X. At 0.1X, the algorithm is unsure about the placement of D4a1d, which spreads over haplogroup L3 and macrohaplogroup M; at 1X, the placement has converged to D4a3a. A) Chagyrskaya 8 at 0.5X, B) Chagyrskaya 8 at 4X, C) D4a1d at 0.1X, D) D4a1d at 1X. Clades in which no source was placed are collapsed into a bar labeled with the lineage and its number of tips; the full trees are shown in Supplementary Figures S2 and S6.

We present results at varying levels of mean coverage depth obtained via downsampling (Chagyrskaya 8: Subpanel (A), 0.5X; Subpanel (B), 4X; D4a1d: Subpanel (C), 0.1X; Subpanel (D), 1X). Already at 0.5X mean coverage depth the posterior over *β* for Chagyrskaya 8 is concentrated on the branch leading to the Les Cottés Neanderthal, with every chain placing more than 99% of its samples on this branch. As coverage increases, the posterior over the position along this branch narrows and moves towards the ancestor that Les Cottés shares with the Denisova 17 Neanderthal (labeled Neanderthal Siberia in our figures), which differs from Chagyrskaya 8 at fewer substitutions (23 versus 34) than the Les Cottés sequence itself. At 0.1X (17 reads) the D4a1d sample, on the other hand, cannot be placed: the posterior over *β* is spread over many branches of haplogroup L3 and, to a lesser extent, of macrohaplogroup M, indicating a high degree of uncertainty. At 1X, and in fact already at 0.5X, the posterior has collapsed onto the branch leading to D4a3a, its closest relative in our reference, as expected.

We have repeated this experiment for all five mitogenomes and all coverage depths between 0.05X and 4X, each in two independent replicates; please find Supplementary Figures S2–S11 in the Supplementary Material. As for Chagyrskaya 8, the Denisovan, L2 and T2a1 samples are placed on the branch of their closest relative in the reference (Denisova 20, L2a1d and T1a1a1, respectively) from 0.1X onwards, and increasing coverage depth narrows the posterior over the position along that branch. At 0.05X (fewer than a dozen reads), Chagyrskaya 8 is still placed on the Les Cottés branch and Denisova 21 on the stem of the Denisovan clade, whereas the placements of the three modern humans are spread over many branches. The two replicates agreed on the placement in every case.

#### 3.1.2 Two-source Samples

Having established that our method is quite robust on simulated single-source samples, we now turn our attention to simulated two-source mixtures. We illustrate the robustness of our tool by showcasing simulated mixtures of the 5 samples described in the preceding subsection spiked with one N macrohaplogroup (ref: HM229344.1; haplogroup U5a2a) at varying mixture proportions. Haplogroup U5 is not represented in our reference; the closest relatives of this mitogenome are the reconstructed ancestral sequences and tips of the U clade (e.g. U2a1a, U6a6a). We used the same fragment size distribution as described above, but to add greater realism, we changed the damage pattern as multiple empirical sources may have different damage rates. We scaled the damage of the previously used rates by 0.75 (see Supp. Fig. S1). The *in silico* mixing was performed at rates of 1%, 5%, 10% … 95%, 99% of the reads of the first source, the remainder being U5a2a, and each of these 105 mixtures was downsampled to mean coverage depths of 0.5X, 1X, 2X, 4X, 8X, 16X and 32X. TrailMix was run as described in the above section except with the “-k 2” flag invoked.

##### Simulated Denisovan + U5a2a Mixture at Varying Levels of Coverage, Fixed Mixture Proportions

Figure 4 illustrates TrailMix predicted placements and mixture proportions in a simulated sedimentary sample comprising a Denisova 21 Denisovan at 30% rate and the remaining 70% from a modern human of haplogroup U5a2a, at varying levels of mean coverage depth. Already at our lowest level of mean coverage depth (0.5X; Subpanels (A) and (B)) the Denisovan source is placed on the branch leading to Denisova 20, and the credible intervals over Θ, while wide, span the ground truth values (Denisovan source: 0.34, 95% CI [0.24, 0.45]). The modern human source, on the other hand, is placed within macrohaplogroup R on the branch leading to haplogroup B6a1 rather than in the U clade. As coverage increases to 1X (Subpanels (C) and (D)) the posterior distributions narrow (0.33 [0.25, 0.41]), and at 2X (Subpanels (E) and (F)) and 8X (Subpanels (G) and (H)) the modern human source moves to the U clade, while the Θ estimates continue to narrow around the true underlying ground truth Θ values for this sample (0.35 [0.29, 0.41] at 2X and 0.31 [0.28, 0.34] at 8X). Please find the remaining coverage depths of this mixture and the simulations on U5a2a mixed with the Neanderthal, L2, M and N macrohaplogroup samples at all mixture proportions and coverage depths, in Supp. Figs. S12–S116.

**Figure 4:**
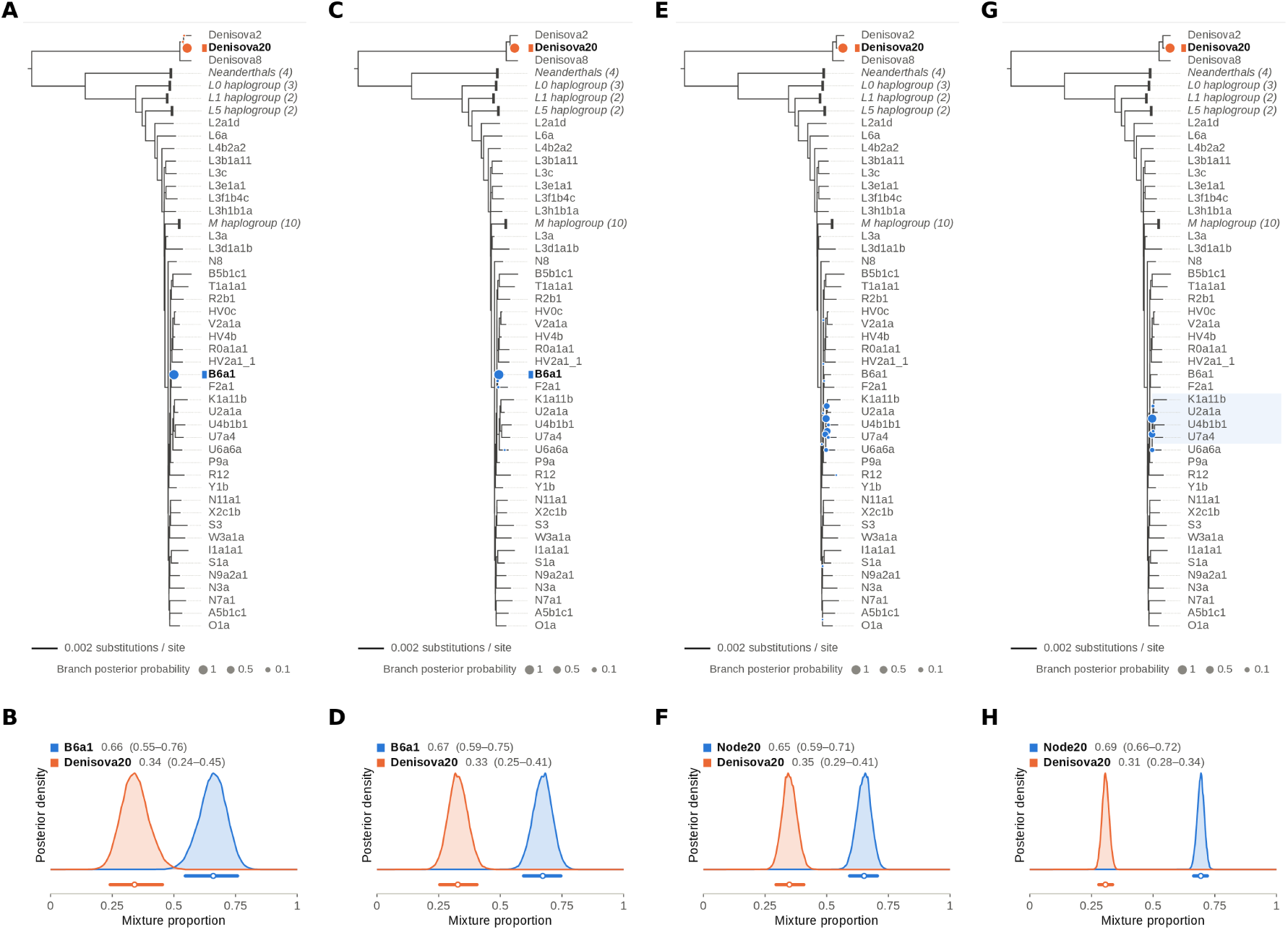
A mix of 30% Denisova 21 from [4], which should be close to Denisova 20, and 70% modern human with U5a2a haplogroup mitochondria, which should fall within the U clade (U5 is not part of our reference). Both sources carry ancient DNA damage. The upper panels represent the phylogenetic placement and the lower panels the inferred proportions (median and 95% credible interval) for the best of five chains. The Denisovan source is placed on the Denisova 20 branch at every coverage depth; the modern human source is placed on B6a1 (macrohaplogroup R) at 0.5X and 1X and within the U clade from 2X. At 0.5X: A)-B), 1X: C)-D), 2X: E)-F), 8X: G)-H). Clades in which no source was placed are collapsed as in Figure 3; the full trees are shown in Supplementary Figure S39.

##### Varying Mixture Proportions

In order to ensure that our inferred mixture proportions are robust for a wide variety of input samples, we stress-tested our program by looking at our Θ estimates for the Neanderthal Chagyrskaya 8 mixed with U5a2a in varying mixture proportions, namely 1%, 5%, 10%, 15%, & , 90%, 95% and 99% of Chagyrskaya 8, the remainder being U5a2a (21 mixtures). We see (Figure 5) that at 1X mean coverage depth (Subpanel (A)) the posterior distributions over Θ are quite wide for each input sample, with the estimates for the most imbalanced input samples (e.g. ⟨0.01, 0.99⟩) among the farthest from the ground truth. In general, samples in which the mixture proportions are not too imbalanced (i.e. the minor mixture proportion is above 10%) are nonetheless fairly centered around the ground truth Θ values, and the root-mean-square error (RMSE) of the minor proportion over the 21 mixtures is 0.056. As mean coverage depth increases to 4X (Subpanel B) we can see the same general pattern still holds, but (1) the posteriors over Θ are narrower, indicating greater confidence; and (2) the estimates are closer to the ground truth (RMSE 0.032). Increasing coverage depth even further, to 8X (Subpanel (C)) and 16X (Subpanel (D)) we see the posteriors over Θ increasingly narrow around the ground truth values (RMSE 0.027 at both coverage depths). On average, the minor proportion is overestimated by 0.02–0.04, and the highly imbalanced samples remain the most difficult: a 1% minor source is estimated at 0.10, 0.06, 0.05 and 0.04 at 1X, 4X, 8X and 16X, respectively. Since the credible intervals narrow faster than this bias decreases, the 90% intervals contain the ground truth in 14 of the 21 mixtures at 1X but only 9 of 21 at 16X, and they should be interpreted with this in mind. The corresponding results for Denisova 21 are of similar accuracy, except for the mixture containing only 1% of Denisova 21, whose proportion is overestimated at most coverage depths (e.g. 0.41 at 2X), while the proportions of the three modern human sources, which are mixed with another modern human, are estimated less accurately (RMSE at 16X between 0.08 and 0.18; Figs. S117–S126).

**Figure 5:**
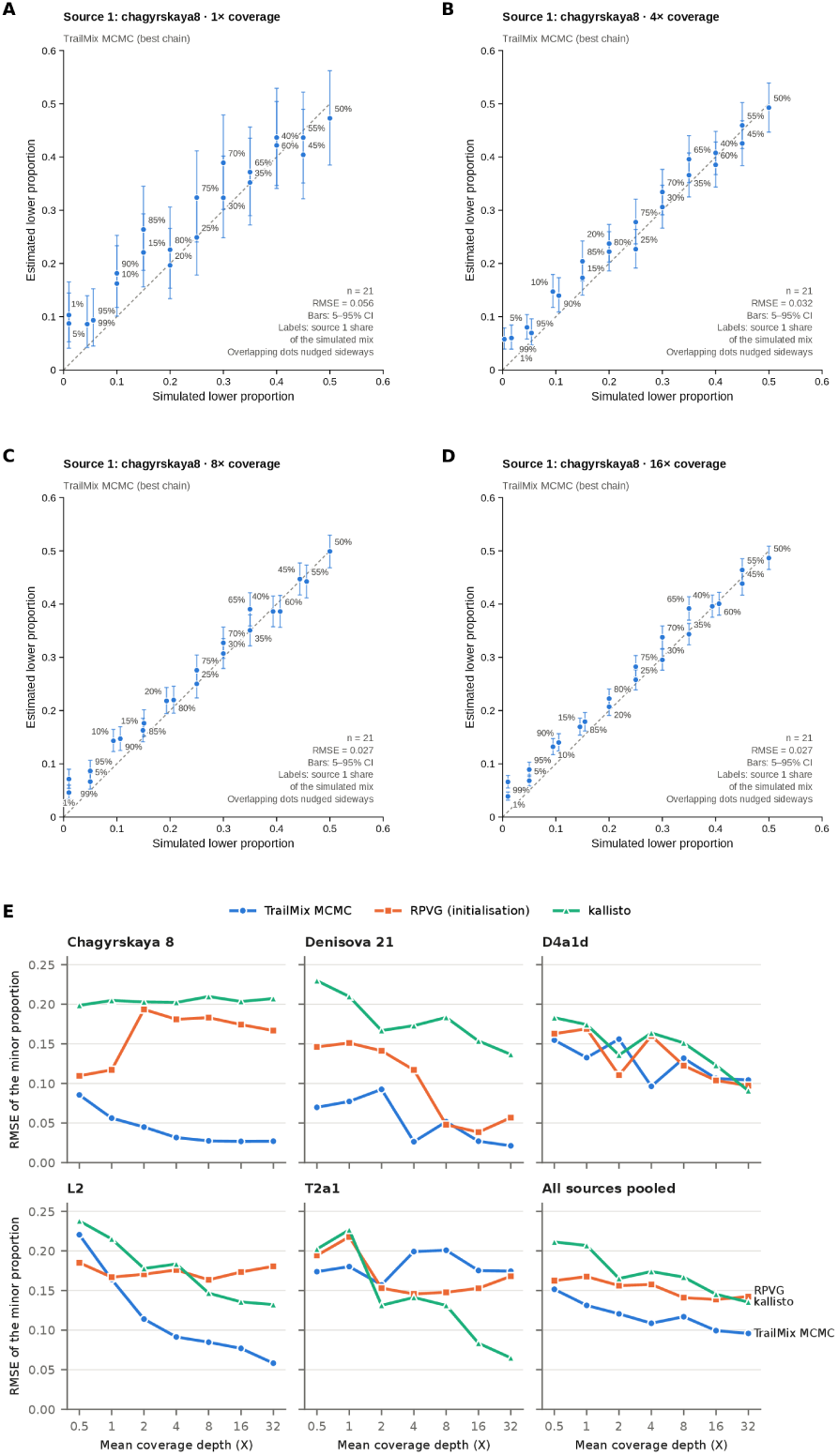
Accuracy of the inferred proportions. The plots represent the estimated versus the simulated minor mixing proportion for a Chagyrskaya8 Neanderthal mixed with modern human with haplogroup U5a2a at 21 rates between 1 to 99%. Each point is the posterior mean of the best chain, with bars showing the 5–95% credible interval, and is labeled with the share of Chagyrskaya 8 in the mixture; the dashed line is the identity, and the RMSE over the 21 mixtures is given in each panel. A) At 1X, B) At 4X, C) At 8X, D) At 16X. E) Root-mean-square error (RMSE) of the estimated minor mixture proportion over the 21 simulated mixtures of each of the five sources with U5a2a, as a function of the mean coverage depth, for TrailMix (best chain), its RPVG initialization and kallisto (two most abundant reference sequences); the last panel pools the five sources (105 mixtures per coverage depth). The individual estimates of the three methods are shown in Supplementary Figures S117–S126.

##### Placement Accuracy Across All Mixtures

Looking at all 735 mixtures (Supplementary Table S3), we see that the placement of the two sources is reliable once both coverage and the share of the minor source are sufficient: when the minor source contributes at least 25% of the reads, both sources are placed in the correct clade in 93–100% of the samples at 4X or more, compared with 27% at 2X and almost never at 1X or below, where the modern human sources are typically placed elsewhere in macrohaplogroups N/R or M. The archaic sources are recovered even as a minor component of 1–5% (Chagyrskaya 8 in 12 of 14 such samples, Denisova 21 in 8 of 14), whereas modern human sources contributing only 1–5% of the reads were placed correctly in only 1 of 112 samples. Conversely, a modern human source of 15% or less next to a Neanderthal majority is detected and quantified but frequently placed on deep African L branches rather than in the U clade. Finally, in 27% of the mixtures of two modern humans, one of the two components was placed on an archaic branch with a small proportion (median 0.08), mostly when one of the two sources contributed 20% or less of the reads; at 4X or more with a minor source of at least 25%, this happened in only 4 of the 132 such mixtures. Small archaic components inferred from low-coverage or imbalanced samples should therefore be interpreted with caution.

##### Comparison to Existing Heuristic Methods

In Vernot *et al.* [44], the authors pioneered the use of kallisto [3], a software tool which was originally intended for use on modern transcriptomics data, for the analysis of ancient hominin sediments instead. We replicated this approach using the same mitochondrial genomes as the ones in the TrailMix database, taking the two reference sequences with the highest estimated read counts as the two sources, and we compared it to TrailMix and to its RPVG initialization on all 735 simulated mixtures (Figure 5E and Supp. Figs. S117–S126). Pooled across all samples, TrailMix is the most accurate of the three methods at every coverage depth: the RMSE of the minor proportion it estimates decreases from 0.15 at 0.5X to 0.10 at 32X, while that of the RPVG initialization remains between 0.14 and 0.17 at all coverage depths, and that of kallisto decreases from 0.21 at 0.5X to 0.14 at 32X. The exception is T2a1, for which kallisto is more accurate than TrailMix from 2X onwards (and D4a1d at 2X and 32X). While the results of this heuristic thus seem to converge towards the ground truth at high coverage, it struggles with phylogenetic uncertainty, as the reads of one source are spread over several closely related reference and ancestral sequences, and it does not quantify the uncertainty of its estimates. Summing the kallisto estimates over all archaic reference sequences instead gives accurate archaic fractions for the Chagyrskaya 8 and Denisova 21 mixtures (RMSE between 0.03 and 0.05 at every coverage depth, better than TrailMix below 4X; Supplementary Section S7), but this requires the groups of interest to be defined in advance and provides no phylogenetic placement of the contributors.

### 3.2 Empirical Data

Finally, and crucially, we demonstrate that our method is robust on empirical samples as well by comparing our results to those obtained via independent methods from two publications. The empirical results shown below were obtained with *k* = 2 and otherwise the same parameters as the simulations, using damage rates estimated from each library; single-source (*k* = 1) runs of all libraries are shown in Supp. Fig. S127. In these figures, the stems of the Neanderthal and Denisovan clades are labeled Node47 and Node45, respectively, and the Neanderthal from Denisova Cave Denisova 17 is labeled Neanderthal Siberia.

#### 3.2.1 Zavala *et al*

We begin by showcasing the results of our tool run on sedimentary aDNA samples described in Zavala *et al.*, 2021 [48]. Figure 6 displays TrailMix estimated phylogenetic placements and mixture proportions for three such samples. Subpanels (A) and (B) result from input sample M76, which has previously been inferred to contain Neanderthal mtDNA with a substantial degree of modern human contamination; in particular, Zavala *et al.* found that the Neanderthal fragments of this sample support the branch shared by the Hohlenstein-Stadel and the other Neanderthal mtDNAs but neither of the branches defining these lineages, and concluded that it carries a previously unknown Neanderthal mtDNA lineage [48]. TrailMix, on the other hand, consistently (in all five chains) deconvolves this sample into a deep lineage on the stem of the Denisovan clade (median 0.61, 95% CI [0.55, 0.67]) and a modern human of macrohaplogroup M on the branch leading to D4a3a (0.39 [0.33, 0.46]). This library also has the shortest fragments of those analyzed here (mean length 49 bp, Supplementary Table S4), and additional data would be required to resolve this discrepancy. Subpanels (C) and (D) are TrailMix outputs from sample E251, which was previously inferred to contain a mixture of human and Neanderthal mtDNA. TrailMix concords with this assessment, with a modern human contributor placed within macrohaplogroup N on the branch leading to N8 (0.67 [0.57, 0.77]) and a Neanderthal contributor on the branch leading to Les Cottés (0.33 [0.23, 0.43]). Finally, Subpanels (E) and (F) are the result of input sample E256, which the authors inferred to be a mixture of Neanderthal and Denisovan. TrailMix corroborates both contributors: the Neanderthal (0.75 [0.68, 0.81]) is placed on the branch leading to the Denisova 17 Neanderthal, which was also found in Denisova Cave, and the Denisovan (0.25 [0.19, 0.32]) on the stem of the Denisovan clade, i.e. on a lineage that diverged before those of the Denisovans in our reference. All five chains agree on this combination of sources. The three remaining libraries from Zavala *et al.* that we analyzed (E221, E227 and E249; Supp. Figs. S128 and S129) are deconvolved into a Denisovan and a modern human contributor (E221, which was previously inferred to contain Neanderthal and Denisovan mtDNA), a Denisovan and a Neanderthal contributor (E227, as previously inferred), and a Neanderthal and a modern human contributor (E249, as inferred previously).

**Figure 6:**
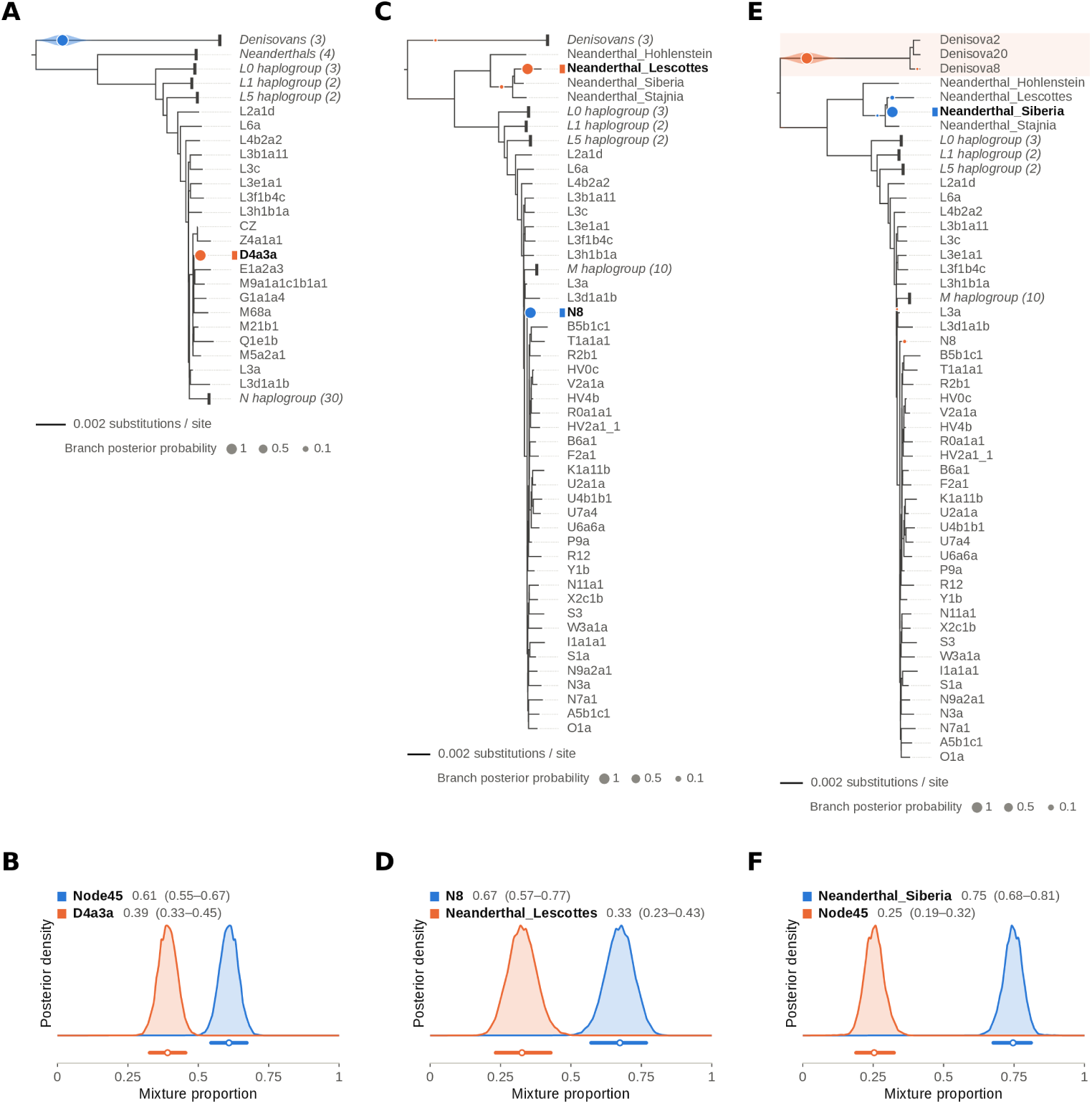
Various empirical samples from Zavala *et al.*[48]. A) and B): M76 has previously been inferred to contain Neanderthal mtDNA with heavy modern human contamination. TrailMix instead places one source on the stem of the Denisovan clade (Node45), which is shaded, and a modern human source on the branch leading to D4a3a (macrohaplogroup M), in all five chains (see text). C) and D): E251, inferred to be a mixture of modern humans and Neanderthal mtDNA. We confirm this finding, with a modern human placed on the branch leading to N8 and a Neanderthal on the branch leading to Les Cottés. E) and F) E256 inferred to be a mixture of Neanderthal and Denisovan mtDNA; we confirm a Neanderthal placed on the branch leading to the Denisova 17 Neanderthal (labeled Neanderthal Siberia) and a second source placed on the stem of the Denisovan clade (Node45), which is shaded. Clades in which no source was placed are collapsed as in Figure 3; the full trees are shown in Supplementary Figures S128 and S129.

#### 3.2.2 Vernot *et al*

We next display the results of TrailMix run on samples from Vernot *et al.* [44]. The Vernot *et al.* sediment samples (Figure 7) provide a complementary test spanning both a well-characterised Neanderthal site and the more divergent Neanderthal population history documented at the Galería de las Estatuas. For Chagyrskaya Cave, whose sediment mtDNA has been shown to agree with the skeletal record [44], sample Ch-3007 was reported as Neanderthal, and TrailMix recovers a predominantly Neanderthal composition (Neanderthal source on the branch leading to Les Cottés, median 0.933, 95% CI [0.921, 0.943]), together with a minor modern human component (0.067, 95% CI [0.057, 0.079]) placed within macrohaplogroup M (Figure 7A–B). The two Estatuas samples are more interesting, since Vernot *et al.* document a population turnover at this site around 100,000 years ago accompanied by a loss of mtDNA diversity, implying the presence of early, deeply divergent Neanderthal lineages [44]. Sample GE-I-B09, reported as Neanderthal, is deconvolved by TrailMix into a dominant Neanderthal contributor (median 0.906, 95% CI [0.894, 0.917]) whose placement sits with the Les Cottés/Denisova 17/Stajnia group of Neanderthals, on the branch leading to Stajnia, together with a minor (∼9%) modern human component placed within macrohaplogroup N (Figure 7C–D). Sample GE-I-A4, inferred to be a highly divergent Neanderthal, yields the most striking result: TrailMix places the dominant contributor (∼91%) on the branch leading to the Hohlenstein-Stadel Neanderthal rather than with the Les Cottés/Denisova 17/Stajnia group, an unusual and deeply divergent placement that is congruent with the early, diversity-rich Neanderthal population inferred for this site, while the remaining ∼9% is a modern human component placed on the branch leading to L5b1a (haplogroup L5) (Figure 7E–F). All three samples therefore contain 7–9% of modern human mtDNA, which may stem from contamination; since in our simulations a minor modern human source next to a Neanderthal majority was well quantified but often misplaced, we do not interpret the haplogroups of these minor components further.

**Figure 7:**
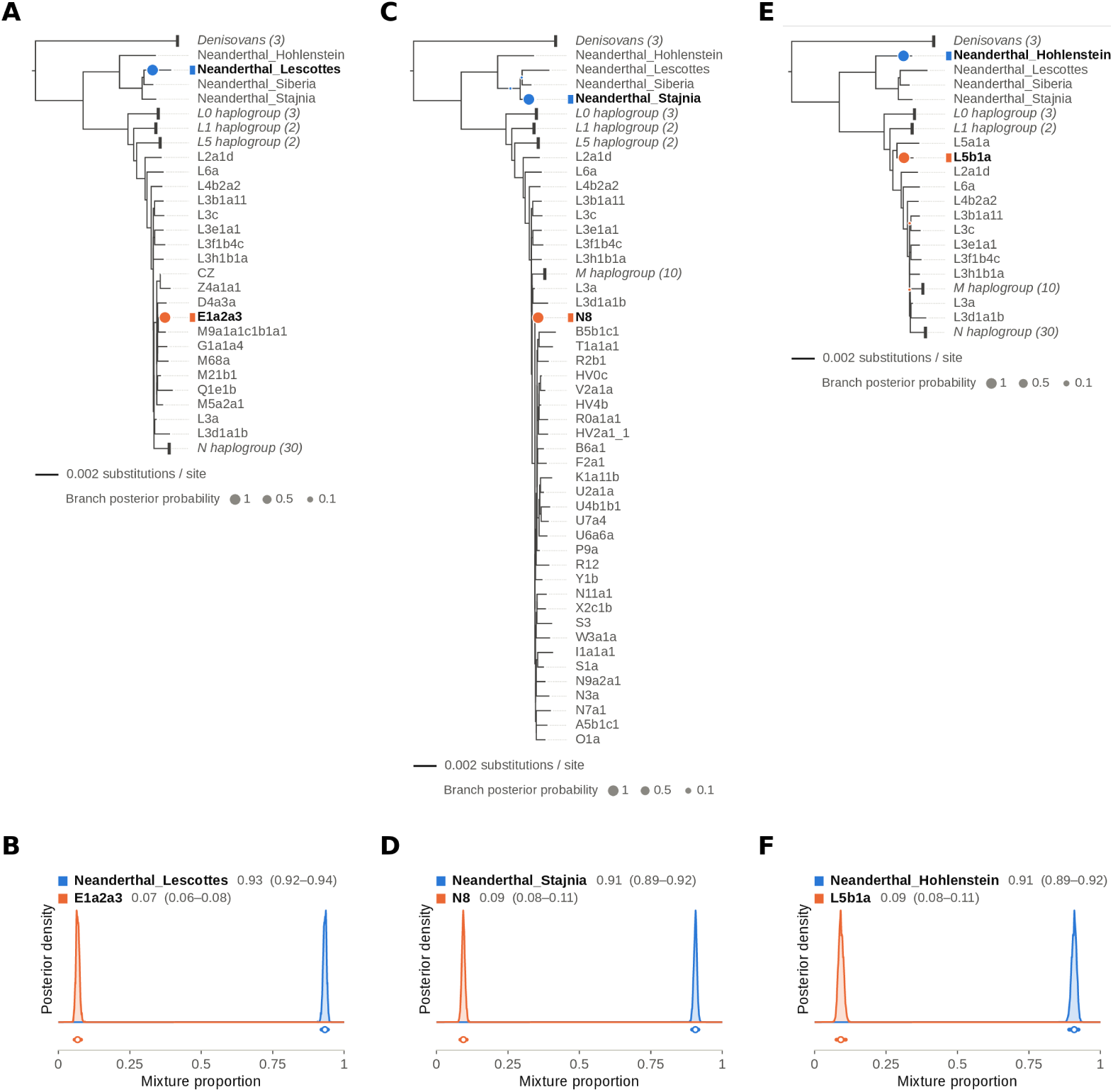
Various empirical samples from Vernot *et al.*[44]. A) and B): Ch-3007 from Chagyrskaya cave predicted to be Neanderthal, which we confirm to be 93% Neanderthal (placed on the branch leading to Les Cottés), with 7% of a modern human placed within macrohaplogroup M. C) and D): GE-I-B09 from the Estatuas cave, predicted to be Neanderthal. We confirm this with 91% of a Neanderthal from the Les Cottés, Denisova 17 and Stajnia group (placed on the branch leading to Stajnia) and 9% of a modern human placed within macrohaplogroup N. E) and F) GE-I-A4 from Estatuas cave, which was predicted to be a very divergent Neanderthal. We find 91% of a Neanderthal placed on the branch leading to Hohlenstein-Stadel (not Les Cottés/Denisova 17/Stajnia), which is rare, and 9% of a modern human placed on the branch leading to L5b1a (haplogroup L5). Clades in which no source was placed are collapsed as in Figure 3; the full trees are shown in Supplementary Figure S130.

Taken together, the empirical results from Zavala *et al.* and Vernot *et al.* show that on authentic, previously analyzed mixtures, TrailMix largely concords with the authors’ findings obtained via independent methods. Unlike competing tools, TrailMix is not restricted to a database of modern haplogroup sequences and is therefore capable of representing contributors as ancestral to the lineages of the reference (e.g. the Denisovan contributors of samples E256 and E227 from Zavala *et al.*, placed on the stem of the Denisovan clade), which is more biologically accurate and helps place such contributors in their proper archaeological context.

## 4 Discussion

We have shown that TrailMix successfully generalizes the Bayesian inference model introduced by the HaploCart program to the more difficult cases in which the sample may constitute a mixture of multiple distinct contributors (modern human or archaic), in which each contributing mitogenome in the sample may represent a population not found among extant humans (and therefore not documented in the standard haplogroup nomenclature), and in which the sample will in general feature the characteristic challenges inherent to ancient DNA chemical damage, namely deamination and fragmentation.

We have first demonstrated that single-source samples affected by aDNA damage can be confidently placed on the reference tree whether the source in question is a modern human or an archaic hominin haplotype, even when its mitogenome is not part of our reference and must be placed relative to its closest relatives. As one would expect, we have demonstrated in these cases that the posterior distribution grows more concentrated around the true parameter values as coverage increases, with a reliable placement from about 0.1X to 0.5X onwards.

As pertains to mtDNA mixtures, we have shown that TrailMix correctly infers, to an acceptable degree of precision, both phylogenetic placements of contributing sources as well as relative proportions per source for many samples. This is the case for in-silico-generated mixtures comprising two ancient mitogenomes, as for example may be found in sediments, in which both sources are placed in the correct clade for nearly all samples from about 4X onwards, provided that the minor source contributes at least a quarter of the reads. We also see results that largely agree with those of the original publications when run on empirical sediment samples from Denisova Cave, Chagyrskaya Cave and the Galería de las Estatuas. Of course, all models will break under some conditions, but we have demonstrated that even for highly imbalanced mixtures, archaic contributors representing as little as 1–5% of the reads are in most cases detected and placed, and their proportions correctly estimated, given sufficient coverage. In general, we recommend at least 0.5X for single sources but 1-2X for two sources.

As with any tool, it is also important to understand the conditions under which the program will fail to converge to the correct result, and we indeed see this for TrailMix when the mixture proportions are highly imbalanced: small proportions tend to be overestimated (a 1% source is estimated at about 4% even at 16X), a modern human contributing a small fraction of the reads is often misplaced, and in mixtures of two modern humans - a spurious archaic component of a few percent can appear. Moreover, the credible intervals over Θ narrow faster and converge towards the simulated values, so that they are overconfident at high coverage. Broadly speaking, for mixture samples we find that mixture proportion estimation is more reliable and robust to low coverage than the phylogenetic placement estimation. We cannot say for certain why this is the case, but we speculate that it has to do with the size of the parameter space on *β* being much larger and therefore more difficult to search than the parameter space on Θ. Further work is needed to better explain and understand this effect, and of course, to make the program even more robust to the most challenging situations that might arise, including mixtures of more than two sources, for which the likelihood is implemented but which we have not evaluated in this work.

There are a handful of competing tools for handling such mitochondrial aDNA samples, but they come with significant limitations of scope, as we have laid out in the introduction; in particular, schmutzi and contamMix assume a modern contaminant and do not apply to mixtures of two ancient sources. In this manuscript, we have therefore benchmarked TrailMix against the kallisto-based approach that has been used to quantify hominin mtDNA in sediments [44], as well as against its own RPVG initialization. TrailMix was the most accurate of the three methods at every coverage depth. On the other hand, summing the kallisto estimates over predefined groups of reference sequences yields accurate archaic fractions even at low coverage, but provides neither a phylogenetic placement of the contributors nor a measure of uncertainty. It is therefore fair to say that at the moment neither TrailMix nor the summed kallisto approach is to be preferred for all use cases. If the user only wishes to quantify predefined groups (e.g. Neanderthal versus modern human) in a low-coverage sample, the latter approach is to be preferred; if the placement of each contributor within its phylogenetic context, possibly on a lineage ancestral to those of the reference, and the uncertainty of the estimates are of interest, then the former should be used instead.

To summarize, the results shown here suggest that TrailMix constitutes a significant advance in our ability to shed light on technically challenging aDNA samples involving mixtures of hominin provenance. In light of these findings, while key limitations doubtless remain, the tool may enable researchers to gain insights into the human past and present. In addition, it could be extended to other species or clades of species with mitogenomes whose similarity is on par with humans.

## 5 Availability of Data and Materials

Data and scripts related to the benchmarking experiments (the simulation Snakefile, the TrailMix command lists and the plotting configurations) will be deposited on Zenodo (DOI to be added upon publication). The code can be found at https://github.com/g renaud/vgan.

## 6 Software Versions

Our vgan version was the development version of 3.1.0, built against vg v1.75.0 (commit 6193310e3), with reads mapped by SAFARI v1.0.0-6-g667d8f7 (based on vg v1.44). Our samtools version was 1.13. Our Snakemake version was 5.10.0. Our gargammel version was 1.1.4. Our ART version was 2.5.8. Our leeHom version was 1.2.16. Our SHRiMP version was 2.2.2. Our kallisto version was 0.51.1. Our Python version was 3.8.8. Our seqtk version was 1.3-r106. Our RPVG version was 1.0.

## 7 Author Contributions

J.B. developed parts of the software. V.S. and E.I.Z. helped with the manuscript and the interpretation of the results. P.W.S. helped with the computational aspects. N.A.V. helped with the mathematical modeling and coding. G.R. conceived and supervised the study.

## 8 Acknowledgements

We would like to thank Joshua D. Rubin, who developed the initial version of TrailMix and of this manuscript. We would like to acknowledge the help of ChatGPT in creating the subfigure for the graphical abstract. Claude was used for generating Snakemake files for simulations and empirical samples, for debugging and extending the TrailMix code and its plotting scripts, and for revising the manuscript. J.B. would like to acknowledge funding from the Natural Sciences and Engineering Research Council of Canada ref. 757749, G.R. would like to acknowledge funding from the Novo Nordisk Data Science Investigator grant number NNF20OC0062491, V.S. would like to acknowledge funding from the Israel Science Foundation (Grant #2075/22.), E.I.Z. would like to acknowledge funding from the Novo Nordisk Hallas Møller Emerging Investigator grant (#125359).

## References

[1] Richard M Andrews, et al. “Reanalysis and Revision of the Cambridge Reference Sequence for Human Mitochondrial DNA”. In: Nature Genetics 23.2 (1999), pp. 147–147.

[2] María C Ávila-Arcos, Maanasa Raghavan, and Carina Schlebusch. “Going local with ancient DNA: A review of human histories from regional perspectives”. In: Science 382.6666 (2023), pp. 53–58.

[3] Nicolas L Bray et al. “Near-optimal probabilistic RNA-seq quantification”. In: Nature Biotechnology 34.5 (2016), pp. 525–527. doi: 10.1038/nbt.3519.

[4] Samantha Brown, et al. “The earliest Denisovans and their cultural adaptation”. In: Nature Ecology & Evolution 6.1 (2022), pp. 28–35.

[5] Matei David, et al. “SHRiMP2: sensitive yet practical short read mapping”. In: Bioinformatics 27.7 (2011), pp. 1011–1012.

[6] Miroslava Derenko, et al. “Origin and post-glacial dispersal of mitochondrial DNA haplogroups C and D in northern Asia”. In: PloS ONE 5.12 (2010), e15214.

[7] Arne Dür, et al. “Post Hoc Deconvolution of Human Mitochondrial DNA Mixtures by EMMA 2 using Fine-tuned Phylotree Nomenclature”. In: Computational and Structural Biotechnology Journal 20 (2022), pp. 3630–3638.

[8] Brent Ewing and Phil Green. “Base-calling of Automated Sequencer Traces Using PHRED. II. Error Probabilities”. In: Genome Research 8.3 (1998), pp. 186–194.

[9] Joseph Felsenstein. Newick tree format. Accessed: 2024-07-18. 2005. url: https://phylipweb.github.io/phylip/newick_doc.html.

[10] Rosa Fregel, et al. “Mitogenomes Illuminate the Origin and Migration Patterns of the Indigenous People of the Canary Islands”. In: PloS One 14.3 (2019), e0209125.

[11] Qiaomei Fu, et al. “A Revised Timescale for Human Evolution Based on Ancient Mitochondrial Genomes”. In: Current Biology 23.7 (2013), pp. 553–559.

[12] Erik Garrison, et al. “Building Pangenome Graphs”. In: Nature Methods (2024), pp. 1–5.

[13] Pere Gelabert, et al. “Early Gravettian genomic data from Western Europe reveal population movements before the LGM”. In: Science Advances 12.43 (2026), eaeh6128.

[14] Pere Gelabert, et al. “Genome-scale sequencing and analysis of human, wolf, and bison DNA from 25,000-year-old sediment”. In: Current Biology 31.16 (2021), pp. 3564–3574.

[15] Torsten Günther, et al. “Ancient Genomes Link Early Farmers from Atapuerca in Spain to Modern-day Basques”. In: Proceedings of the National Academy of Sciences 112.38 (2015), pp. 11917–11922.

[16] Henrik B Hansen, et al. “Comparing Ancient DNA Preservation in Petrous Bone and Tooth Cementum”. In: PloS One 12.1 (2017), e0170940.

[17] Masami Hasegawa, Hirohisa Kishino, and Taka-aki Yano. “Dating of the Human-ape Splitting by a Molecular Clock of Mitochondrial DNA”. In: Journal of Molecular Evolution 22 (1985), pp. 160–174.

[18] Kazutaka Katoh, et al. “MAFFT: a Novel Method for Rapid Multiple Sequence Alignment Based on Fast Fourier Transform”. In: Nucleic Acids Research 30.14 (2002), pp. 3059–3066.

[19] Johannes Köster and Sven Rahmann. “Snakemake—a Scalable Bioinformatics Workflow Engine”. In: Bioinformatics 28.19 (2012), pp. 2520–2522.

[20] Louis Kraft, et al. “AdDeam: a fast and scalable tool for estimating and clustering reference-level damage profiles”. In: Bioinformatics 41.8 (2025), btaf407.

[21] Marie Lacan, et al. “Ancient DNA suggests the leading role played by men in the Neolithic dissemination”. In: Proceedings of the National Academy of Sciences 108.45 (2011), pp. 18255–18259.

[22] Nicole Maca-Meyer, et al. “Major genomic mitochondrial lineages delineate early human expansions”. In: BMC Genetics 2.1 (2001), p. 13.

[23] Fabrizio Mafessoni, et al. “A high-coverage Neandertal genome from Chagyrskaya Cave”. In: Proceedings of the National Academy of Sciences 117.26 (2020), pp. 15132–15136.

[24] Boris Malyarchuk, et al. “Mitogenomic diversity in Russians and Poles”. In: Forensic Science International: Genetics 30 (2017), pp. 51–56.

[25] Rui Martiniano, et al. “Placing ancient DNA sequences into reference phylogenies”. In: Molecular Biology and Evolution 39.2 (2022), msac017.

[26] Frederick A Matsen, Robin B Kodner, and E Virginia Armbrust. “pplacer: linear time maximum-likelihood and Bayesian phylogenetic placement of sequences onto a fixed reference tree”. In: BMC Bioinformatics 11 (2010), pp. 1–16.

[27] Nicholas Metropolis et al. “Equation of State Calculations by Fast Computing Machines”. In: The Journal of Chemical Physics 21.6 (1953), pp. 1087–1092.

[28] Nathan Nakatsuka, et al. “ContamLD: Estimation of Ancient Nuclear DNA Contamination Using Breakdown of Linkage Disequilibrium”. In: Genome Biology 21 (2020), pp. 1–22.

[29] Lam-Tung Nguyen, et al. “IQ-TREE: a fast and effective stochastic algorithm for estimating maximum-likelihood phylogenies”. In: Molecular Biology and Evolution 32.1 (2015), pp. 268–274.

[30] Sanni Översti, et al. “Human Mitochondrial DNA Lineages in Iron-Age Fennoscandia Suggest Incipient Admixture and Eastern Introduction of Farming-related Maternal Ancestry”. In: Scientific Reports 9.1 (2019), p. 16883.

[31] Christopher Pockrandt, et al. “GenMap: Ultra-fast Computation of Genome Mappability”. In: Bioinformatics 36.12 (2020), pp. 3687–3692.

[32] Matthew D. Rasmussen and Manolis Kellis. “A Bayesian Approach for Fast and Accurate Gene-tree Reconstruction”. In: Molecular Biology and Evolution 28.1 (Jan. 2011). Epub 2010 Jul 25, pp. 273–290. doi: 10.1093/molbev/msq189. url: http://compbio.mit.edu/spimap/.

[33] Gabriel Renaud, et al. “gargammel: a Sequence Simulator for Ancient DNA”. In: Bioinformatics 33.4 (2017), pp. 577–579.

[34] Gabriel Renaud, et al. “Joint Estimates of Heterozygosity and Runs of Homozygosity for Modern and Ancient Samples”. In: Genetics 212.3 (2019), pp. 587–614.

[35] Gabriel Renaud, et al. “Schmutzi: Estimation of Contamination and Endogenous Mitochondrial Consensus Calling for Ancient DNA”. In: Genome Biology 16.1 (2015), pp. 1–18.

[36] Harald Ringbauer, et al. “Accurate detection of identity-by-descent segments in human ancient DNA”. In: Nature Genetics (2023), pp. 1–9.

[37] Joshua Daniel Rubin, et al. “HaploCart: Human mtDNA Haplogroup Classification Using a Pangenomic Reference Graph”. In: PLOS Computational Biology 19.6 (2023), e1011148.

[38] Jonas A Sibbesen, et al. “Haplotype-aware pantranscriptome analyses using spliced pangenome graphs”. In: Nature Methods 20.2 (2023), pp. 239–247.

[39] Martin Sikora, et al. “Ancient genomes show social and reproductive behavior of early Upper Paleolithic foragers”. In: Science 358.6363 (2017), pp. 659–662.

[40] Viviane Slon, et al. “Neandertal and Denisovan DNA from Pleistocene sediments”. In: Science 356.6338 (2017), pp. 605–608.

[41] James B Stewart and Patrick F Chinnery. “Extreme Heterogeneity of Human Mitochondrial DNA from Organelles to Populations”. In: Nature Reviews Genetics 22.2 (2021), pp. 106–118.

[42] Arev P. Sümer, et al. “A Method to Analyze Low-Quality Archaic Human Genomes and its Application to the Teshik-Tash 1 Neandertal”. In: bioRxiv (2026). Preprint. doi: 10.64898/2026.08.10.743885.

[43] Stefania Vai et al. “A Genetic Perspective on Longobard-Era Migrations”. In: European Journal of Human Genetics 27.4 (2019), pp. 647–656.

[44] Benjamin Vernot, et al. “Unearthing Neanderthal population history using nuclear and mitochondrial DNA from cave sediments”. In: Science 372.6542 (2021), eabf1667.

[45] Nicola Alexandra Vogel, et al. “soibean: High-resolution Taxonomic Identification of Ancient Environmental DNA Using Mitochondrial Pangenome Graphs”. In: Molecular Biology and Evolution (2024), msae203.

[46] Samuel H Vohr, et al. “A Phylogenetic Approach for Haplotype Analysis of Sequence Data from Complex Mitochondrial Mixtures”. In: Forensic Science International: Genetics 30 (2017), pp. 93–105.

[47] Hansi Weissensteiner, et al. “Contamination Detection in Sequencing Studies Using the Mitochondrial Phylogeny”. In: Genome Research 31.2 (2021), pp. 309–316.

[48] Elena I Zavala, et al. “Pleistocene sediment DNA reveals hominin and faunal turnovers at Denisova Cave”. In: Nature 595.7867 (2021), pp. 399–403.

[49] Dongju Zhang, et al. “Denisovan DNA in late pleistocene sediments from Baishiya Karst Cave on the Tibetan Plateau”. In: Science 370.6516 (2020), pp. 584–587.

