## Supplemental Information for "TrailMix: Mixture Proportion and Placement Estimation of Ancient Hominin mtDNA Samples using a Pangenomic Reference"

November 20, 2026

**1** Department of Health Technology, Section for Bioinformatics, Technical University of Denmark, Kongens Lyngby, Denmark

**2** Department of Forensic Medicine, University of Copenhagen, Copenhagen, Denmark **3** The Globe Institute, University of Copenhagen, Copenhagen, Denmark **4** Department of Anatomy and Biological Anthropology, and Department of Human Genetics and Computational Medicine, Gray School of Medicine, Gray Faculty of Medical & Health Sciences, Tel Aviv University, Tel Aviv, Israel **5** The Dan David Center for Human Evolution and Biohistory Research, Tel Aviv University, Tel Aviv, Israel **6** Department of Computer Science and Software Engineering, Université Laval, Québec City, Québec, Canada

#### Contents

**S1 *in silico* Sample Generation** **2**

**S2 Reference Database and Run Settings** **5**

**S3 Initialization of the Markov State** **7**

**S4 Alignment Details** **8**

**S5 Single-source Simulated Samples** **10**

**S6 Two-source Simulated Mixtures** **21**

**S7 Estimated versus Simulated Mixture Proportions** **127**

#### S1 *in silico* Sample Generation

Simulated samples for our experiments were generated using the `Snakemake` workflow manager [11]. The Snakefile can be found at `sims_diff_deam_profiles/Snakefile`, adapted from `MT_contamination/Snakefile`, together with the data and scripts listed in the main text (Availability of Data and Materials).

We simulated reads from six published mitogenomes, none of which is part of our reference graph (Table S1). For each mitogenome, we first generated 1,000,000 fragments using the `fragSim` subcommand of `gargammel` [19], with fragment lengths drawn from the empirical length distribution of the mitochondrial reads of the ATP2 sample of Günther *et al.* [8] (35–189 bp, median 59 bp; Supplementary Figure S1A). aDNA damage was simulated using the `deamSim` subcommand of `gargammel` [19] with the damage rates measured on the same sample, a double-stranded pattern with a C→T rate of 0.33 at the first 5' position and a G→A rate of 0.33 at the last 3' position (Supplementary Figure S1B). For the U5a2a mitogenome, which serves as the second source of every two-source mixture, these rates were scaled by 0.75 (Supplementary Figure S1C), so that the two sources of a mixture carry different levels of damage. Adapters were added using the `adptSim` subcommand of `gargammel`. Illumina HiSeq 2500 paired-end reads were then simulated using `ART` [9] at a 100 bp read length. Adapters were trimmed and these reads were merged using `leeHom` with the `--ancientdna` flag [18], and the merged reads were aligned to the rCRS using `SHRiMP` [3]. No other DNA (e.g. NuMTs or bacterial DNA) was added to the simulated samples.

For the single-source samples, a random subset of 5% of the reads of each source (about 50,000 reads, a mean coverage depth of about 195X) was downsampled with `samtools` [13] to nominal mean coverage depths of 0.05X, 0.1X, 0.5X, 1X, 2X and 4X. For mixture samples, we merged a random subset of $N$% of the reads of one of the five focal sources with a random subset of $(100 - N)$% of the reads of U5a2a, for $N \in \{1, 5, 10, 15, \ldots, 90, 95, 99\}$, and downsampled each of these 105 mixtures to nominal mean coverage depths of 0.5X, 1X, 2X, 4X, 8X, 16X and 32X, giving 735 two-source samples. Since both sources share the same fragment length distribution, the proportions of reads and of nucleotides from each source are the same up to sampling noise. For each sample, the damage rates passed to `TrailMix` (`--deam5p`, `--deam3p`) were re-estimated from the downsampled reads using `bam2prof` [12].

Two technical points should be noted. First, because the downsampling selects reads by a hash of their name and the names produced by `fragSim` are not unique for fragments with identical coordinates, the FASTQ files analyzed by `TrailMix` contain about 15–30% fewer reads than the nominal coverage depth implies (e.g. 0.72X–0.86X instead of 1X for the single-source samples). Second, at 0.05X (and 0.1X for Chagyrskaya 8) some positions of the re-estimated damage profiles were undefined because no C or G was observed there; `TrailMix` sets such undefined rates to zero. We report the nominal coverage depths throughout.

Exact command and parameter settings can be found in the Snakefile.

| Source | Accession | Haplogroup | Closest relative in the reference |
|---|---|---|---|
| Chagyrskaya 8 | MK388903.1 [15] | Neanderthal | Neanderthal_Lescottes (Les Cottés) |
| Denisova 21 | MT576653.1 [2] | Denisovan | Denisova20 |
| L2 | AF381981.1 [14] | L2 | L2a1d |
| D4a1d | FJ951517.1 [4] | D4a1d (macrohaplogroup M) | D4a3a |
| T2a1 | KY671094.1 [16] | T2a1 (macrohaplogroup N) | T1a1a1 |
| U5a2a | HM229344.1 | U5a2a (macrohaplogroup N) | U clade (ancestral nodes; tips e.g. U2a1a, U6a6a) |

Table S1: **Mitogenomes used to simulate samples.** The first five were used as single sources and as the first (focal) source of the two-source mixtures; U5a2a was the second source of every mixture and carries 0.75 times the damage rates of the other five. None of these mitogenomes is part of the `TrailMix` reference; the last column gives the closest reference sequence(s) on the reference tree.

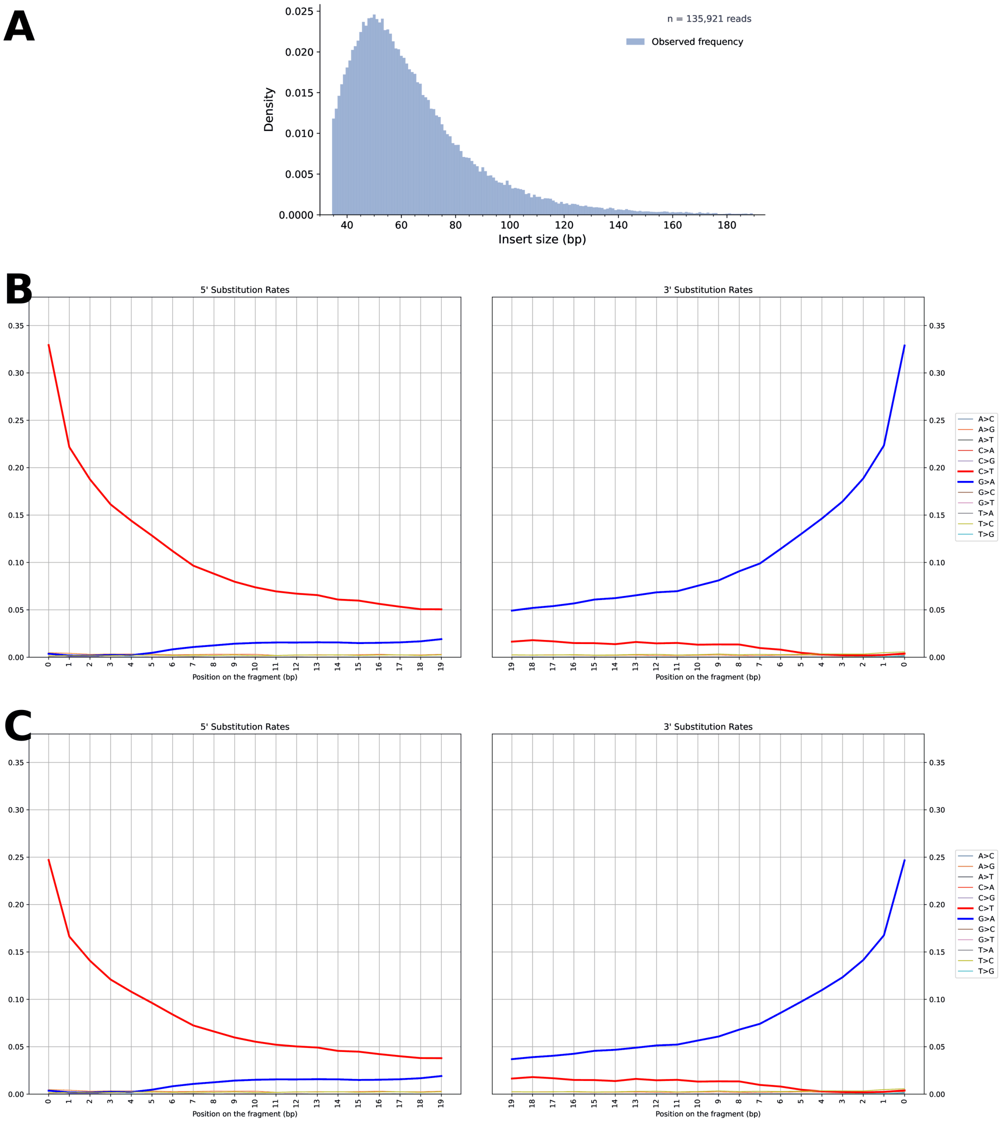


Figure S1: **Simulation parameters.** (A) Fragment length distribution of the mitochondrial reads of the ATP2 sample from Günther *et al.* [8] ($n$ = 135,921 reads), used by `fragSim` for all simulated sources. (B) Substitution rates as a function of the position from the 5' end (left) and the 3' end (right) of the fragments measured on the same sample, used by `deamSim` for the five focal sources (Chagyrskaya 8, Denisova 21, L2, D4a1d and T2a1). (C) The same rates with C$\rightarrow$T and G$\rightarrow$A scaled by 0.75, used for U5a2a.

#### S2 Reference Database and Run Settings

All runs used the reference graph described in the main text, comprising 126 sequences (64 mitogenomes, including 4 Neanderthals and 3 Denisovans, and 62 reconstructed ancestral states), 5,425 nodes and 6,664 edges, with a reference tree of 64 tips and 125 branches. The 64 mitogenomes were aligned with `MAFFT` 7.525 [10]; the tree was inferred with `IQ-TREE` 1.6.12 under the HKY+F model [17], and the 62 ancestral sequences were obtained by marginal ancestral state reconstruction, with the presence or absence of each alignment gap in the ancestors inferred by Fitch parsimony. The 126 sequences were then combined into a graph with `PGGB` v0.2.0 [6] (`-s 5000 -l 5000 -p 95 -n 126 -k 19 -G 5000,5500`), whose nodes were split into segments of at most 6 bp; no stretch of more than 35 bp of the graph is private to a single sequence. In the tree figures, the archaic tips are labeled Neanderthal_Hohlenstein (Hohlenstein-Stadel), Neanderthal_Lescottes (Les Cottés), Neanderthal_Siberia (Denisova 17, a Neanderthal from Denisova Cave), Neanderthal_Stajnia (Stajnia S5000), Denisova2, Denisova8 and Denisova20; internal branches are named after the ancestral node they lead to, e.g. Node47 for the stem of the Neanderthal clade and Node45 for the stem of the Denisovan clade.

Table S2: Mitogenomes in the TrailMix database (the 64 tips of the tree; the ancestral sequences are inferred from them). *Name* is the name of the path in the database. The modern human sequences (57) are haplogroup consensus sequences from the HaploCart graph build; they have no GenBank accession. The archaic genomes (7) are GenBank records. Length and N are those of the sequence as it enters the database (19–31 N in the modern sequences).

| # | Name | Group | Source | Length (bp) | N |
|---|---|---|---|---|---|
| 1 | Denisova2 | Denisovan | KX663333.1 | 16 570 | 13 |
| 2 | Denisova8 | Denisovan | KT780370.1 | 16 573 | 0 |
| 3 | Denisova20 | Denisovan | MT576652.1 | 16 571 | 0 |
| 4 | Neanderthal_Hohlenstein | Neanderthal | KY751400.2 | 16 584 | 12 |
| 5 | Neanderthal_Lescottes | Neanderthal | MG025536.1 | 16 565 | 0 |
| 6 | Neanderthal_Siberia | Neanderthal | MT576650.1 | 16 584 | 19 |
| 7 | Neanderthal_Stajnia | Neanderthal | MT795654.1 | 16 567 | 14 |
| 8 | A5b1c1 | Modern human | HaploCart consensus | 16 584 | 21 |
| 9 | B5b1c1 | Modern human | HaploCart consensus | 16 584 | 29 |
| 10 | B6a1 | Modern human | HaploCart consensus | 16 584 | 29 |
| 11 | CZ | Modern human | HaploCart consensus | 16 584 | 20 |
| 12 | D4a3a | Modern human | HaploCart consensus | 16 584 | 19 |
| 13 | E1a2a3 | Modern human | HaploCart consensus | 16 584 | 20 |
| 14 | F2a1 | Modern human | HaploCart consensus | 16 584 | 20 |
| 15 | G1a1a4 | Modern human | HaploCart consensus | 16 584 | 19 |

*continued on the next page*

| # | Name | Group | Source | Length (bp) | N |
|---|---|---|---|---|---|
| 16 | HV0c | Modern human | HaploCart consensus | 16 584 | 19 |
| 17 | HV2a1*1 | Modern human | HaploCart consensus | 16 584 | 20 |
| 18 | HV4b | Modern human | HaploCart consensus | 16 584 | 19 |
| 19 | I1a1a1 | Modern human | HaploCart consensus | 16 584 | 22 |
| 20 | K1a11b | Modern human | HaploCart consensus | 16 584 | 29 |
| 21 | L0a3 | Modern human | HaploCart consensus | 16 584 | 20 |
| 22 | L0d2b1a1a | Modern human | HaploCart consensus | 16 584 | 21 |
| 23 | L0d3a | Modern human | HaploCart consensus | 16 584 | 19 |
| 24 | L1b | Modern human | HaploCart consensus | 16 584 | 20 |
| 25 | L1c5 | Modern human | HaploCart consensus | 16 584 | 31 |
| 26 | L2a1d | Modern human | HaploCart consensus | 16 584 | 19 |
| 27 | L3a | Modern human | HaploCart consensus | 16 584 | 20 |
| 28 | L3b1a11 | Modern human | HaploCart consensus | 16 584 | 20 |
| 29 | L3c | Modern human | HaploCart consensus | 16 584 | 20 |
| 30 | L3d1a1b | Modern human | HaploCart consensus | 16 584 | 19 |
| 31 | L3e1a1 | Modern human | HaploCart consensus | 16 584 | 19 |
| 32 | L3f1b4c | Modern human | HaploCart consensus | 16 584 | 21 |
| 33 | L3h1b1a | Modern human | HaploCart consensus | 16 584 | 19 |
| 34 | L4b2a2 | Modern human | HaploCart consensus | 16 584 | 19 |
| 35 | L5a1a | Modern human | HaploCart consensus | 16 584 | 21 |
| 36 | L5b1a | Modern human | HaploCart consensus | 16 584 | 21 |
| 37 | L6a | Modern human | HaploCart consensus | 16 584 | 19 |
| 38 | M5a2a1 | Modern human | HaploCart consensus | 16 584 | 19 |
| 39 | M9a1a1c1b1a1 | Modern human | HaploCart consensus | 16 584 | 21 |
| 40 | M21b1 | Modern human | HaploCart consensus | 16 584 | 19 |
| 41 | M68a | Modern human | HaploCart consensus | 16 584 | 23 |
| 42 | N3a | Modern human | HaploCart consensus | 16 584 | 19 |
| 43 | N7a1 | Modern human | HaploCart consensus | 16 584 | 21 |
| 44 | N8 | Modern human | HaploCart consensus | 16 584 | 19 |
| 45 | N9a2a1 | Modern human | HaploCart consensus | 16 584 | 19 |
| 46 | N11a1 | Modern human | HaploCart consensus | 16 584 | 19 |
| 47 | O1a | Modern human | HaploCart consensus | 16 584 | 19 |
| 48 | P9a | Modern human | HaploCart consensus | 16 584 | 19 |
| 49 | Q1e1b | Modern human | HaploCart consensus | 16 584 | 19 |
| 50 | R0a1a1 | Modern human | HaploCart consensus | 16 584 | 20 |
| 51 | R2b1 | Modern human | HaploCart consensus | 16 584 | 19 |
| 52 | R12 | Modern human | HaploCart consensus | 16 584 | 19 |
| 53 | S1a | Modern human | HaploCart consensus | 16 584 | 19 |
| 54 | S3 | Modern human | HaploCart consensus | 16 584 | 20 |
| 55 | T1a1a1 | Modern human | HaploCart consensus | 16 584 | 20 |
| 56 | U2a1a | Modern human | HaploCart consensus | 16 584 | 20 |
| 57 | U4b1b1 | Modern human | HaploCart consensus | 16 584 | 19 |
| 58 | U6a6a | Modern human | HaploCart consensus | 16 584 | 19 |
| 59 | U7a4 | Modern human | HaploCart consensus | 16 584 | 20 |

*continued on the next page*

| # | Name | Group | Source | Length (bp) | N |
|---|---|---|---|---|---|
| 60 | V2a1a | Modern human | HaploCart consensus | 16 584 | 19 |
| 61 | W3a1a | Modern human | HaploCart consensus | 16 584 | 19 |
| 62 | X2c1b | Modern human | HaploCart consensus | 16 584 | 19 |
| 63 | Y1b | Modern human | HaploCart consensus | 16 584 | 20 |
| 64 | Z4a1a1 | Modern human | HaploCart consensus | 16 584 | 22 |

`TrailMix` was run with `--iter 100000 --chains 5` and `-k 1` (single-source samples) or `-k 2` (two-source mixtures), the sample-specific damage rates (`--deam5p`, `--deam3p`), the default burn-in of 100 iterations and the default restriction of the search space to branches within 10 edges of a haplotype identified by `RPVG` (`--depth 10`; main text Section 2.2.3). Each single-source sample was run twice (replicates 1 and 2 below), with identical settings. For every run, the figures show the chain with the highest minimum effective sample size over the sources and over the placement and the mixture proportions ("best chain"); the other chains are summarized in the diagnostic files written by `TrailMix`. Figures were drawn with `plot_trailmix_trees.py` as described in the main text.

### S3 Initialization of the Markov State

Our initialization procedure makes use of a tool called `RPVG`. `RPVG` is actually not intended for DNA, but is rather a pantranscriptomics tool which was designed to perform haplotype-specific transcript (HST) expression estimation from RNA-seq data using a spliced pangenome reference graph [21]. A spliced pangenome reference graph is a special type of variation graph in which the embedded paths do not represent haplotypes, but rather transcripts, and in which edges connect either side of exon boundaries, which is critical for alignment of RNA-seq data where alternative splicing means that one cannot know which intronic regions will have been spliced in the transcripts being mapped [5]. In the intended `RPVG` pipeline, the user should first map RNA-seq data to a spliced reference graph via the `vg mpmap` subcommand of the `vg` toolkit [7, 21] to create a GAMP (graph alignment / multipath) file, which would then become the input for `RPVG` to estimate HST expression.

The `RPVG` program has several different modes, designed for different use cases. We will discuss two such modes. The first is called 'haplotypes', and it is designed to infer joint posterior probabilities over haplotype combinations (i.e. combinations of $k$ haplotypes, for organisms of ploidy $k$). This is done via Gibbs sampling (a form of Monte Carlo Markov Chain sampling). The second `RPVG` mode of interest is called "haplotype-transcripts". In this mode, the program conditionalizes over the joint posterior probabilities over k-haplotype combinations discussed above, and then uses an expectation maximization (EM) algorithm to infer haplotype-specific transcript abundances [21]. Under this mode the program will report the estimated level of abundance of each sufficiently probable "transcript", so we can obtain an initial $\Theta$ estimate by taking the relative ratios of the abundance estimates of each contributing source in the most probable k-haplotype combination.

It may seem prima facie strange to use an RNA tool as part of a DNA pipeline, but in terms of the underlying model we believe it makes sense as an initialization step. Because the goal in initialization is merely to initialize the program in a reasonable region of the parameter space, so as to increase the chance of convergence of the model, etc. For the 'haplotypes' mode of `RPVG`, in case the aligned reads are to a non-spliced graph, and the alignment is single-path rather than multipath, then from a mathematical point of view (notwithstanding the biological interpretation) the program is essentially performing the task of identifying the combination of $k$ embedded paths (however you may wish to interpret them) in a mixture which would best explain the input sample, which is very close to what `TrailMix` aims to do for the estimation of $\beta$, and the "haplotype-transcripts" mode infers relative proportions of these paths, which is again what `TrailMix` aims to do for the purpose of initializing $\Theta$. The most important distinction is that of course `RPVG` does not factor in aDNA damage, which is why the initialization may not be perfect at high levels of damage. As a secondary matter, `RPVG` also assumes the underlying sample is identical to one of the embedded paths in the reference graph, which may not be the case for us, but empirically we find that the tool is quite robust to this. To allow the "haplotype-transcripts" mode to be compatible with our software we generated artificial "spliced transcripts", one for each of the graph's embedded haplotypes. The length of each transcript is declared to be the length of the relevant mitogenome, and the transcript identifier is randomly generated.

All chains of a run start from the same initial state.

#### S4 Alignment Details

As mentioned in Section S3 and in the main manuscript, the separate pantranscriptomics program `RPVG` is used for intelligent initialization of the MCMC chains in `TrailMix`. We include the `RPVG` header files when compiling `vgan`. We created a custom modified version of the `RPVG` main function (rpvg_main.cpp) so as not to conflict with our own. We found that multithreaded output writing is for some reason not compatible with our changes, so output writing in our modified main file is single-threaded.

Alignment of the input data is conducted internally using `SAFARI`, a modified version of `vg giraffe` which takes aDNA damage into account to rescue more alignments from ancient samples [22, 20]. Since `SAFARI` is based on an older version of `vg` (v1.44) than the rest of `vgan`, it is run as a separate process, to which the damage rates provided by the user are passed as well. This produces an output GAM file (the graph-theoretic analog of a BAM file) in a named pipe. We then pass this GAM file to a call to the `RPVG haplotypes` subcommand for an initial estimate of the deconvolved haplotypes for each individual source. This same GAM file is then reused to run the separate subcommand `RPVG haplotype-transcripts` (for mixture proportion initialization) in another forked child process. The outputs of both subprocesses are then parsed, and passed as input into `TrailMix`.

The exact command to the `RPVG haplotypes` subcommand is

```
rpvg -g <prefix>.xg -p <prefix>.gbwt \
     -a rpvg_gamfilename -o tmpdir/rpvg_hap \
     -i haplotypes -u -s -t <num_threads> --filt-best-score 1e-4 \
     --min-noise-prob 1e-10 -m 60 -d 20 --path-node-cluster --use-hap-gibbs \
     [-r <rng_seed>] -y <k_value> --score-not-qual \
     --vgan-temp-dir tmpdir --prob-precision 1e-10
```

Note that the `-u` flag indicates that the alignment file is GAM (single-path) format rather than GAMP (multipath). In other words this indicates that the reference graph is not spliced (or if you prefer, it tells the program that there are never any exon boundaries crossed by these 'transcripts'). Also note that the flag -y <k_value> indicates the number of distinct sources in the sample. In the original program the intent here is to indicate the ploidy of the organism, but mathematically the meanings are the same, as discussed in Section S3.

The exact command to the `RPVG haplotype-transcripts` subcommand is

```
rpvg -g <prefix>.xg -p <prefix>.gbwt
     -a rpvg_gamfilename -o tmpdir/rpvg_ht
     -i haplotype-transcripts -f pantranscriptome.txt
     -u -s -t <num_threads> --path-node-cluster --use-hap-gibbs
     [-r <rng_seed>] -y <k_value> -m 60 -d 20 --score-not-qual
     --min-noise-prob 1e-10 --vgan-temp-dir tmpdir -b
     --prob-precision 1e-8 --filt-best-score 1e-4
```

Here `pantranscriptome.txt` is the artificial transcript file described in Section S3.

#### S5 Single-source Simulated Samples

Here we show the `TrailMix` placements of the five single-source simulated samples (Table S1) at nominal mean coverage depths of 0.05X, 0.1X, 0.5X, 1X, 2X and 4X, for two independent replicate runs with identical settings (Supplementary Figures S2–S11). The four runs shown in main Figure 3, in which clades without any placement are collapsed, are shown here with the full tree. From 0.1X onwards (0.5X for D4a1d), the best chain places at least 99% of its samples on the branch of the closest relative of the simulated mitogenome in the reference. At 0.05X, Chagyrskaya 8 is still placed on the Les Cottés branch and Denisova 21 on the stem of the Denisovan clade, whereas the placements of the three modern humans are spread over many branches, as are those of D4a1d at 0.1X. Both replicates agree on the modal branch in all cases.

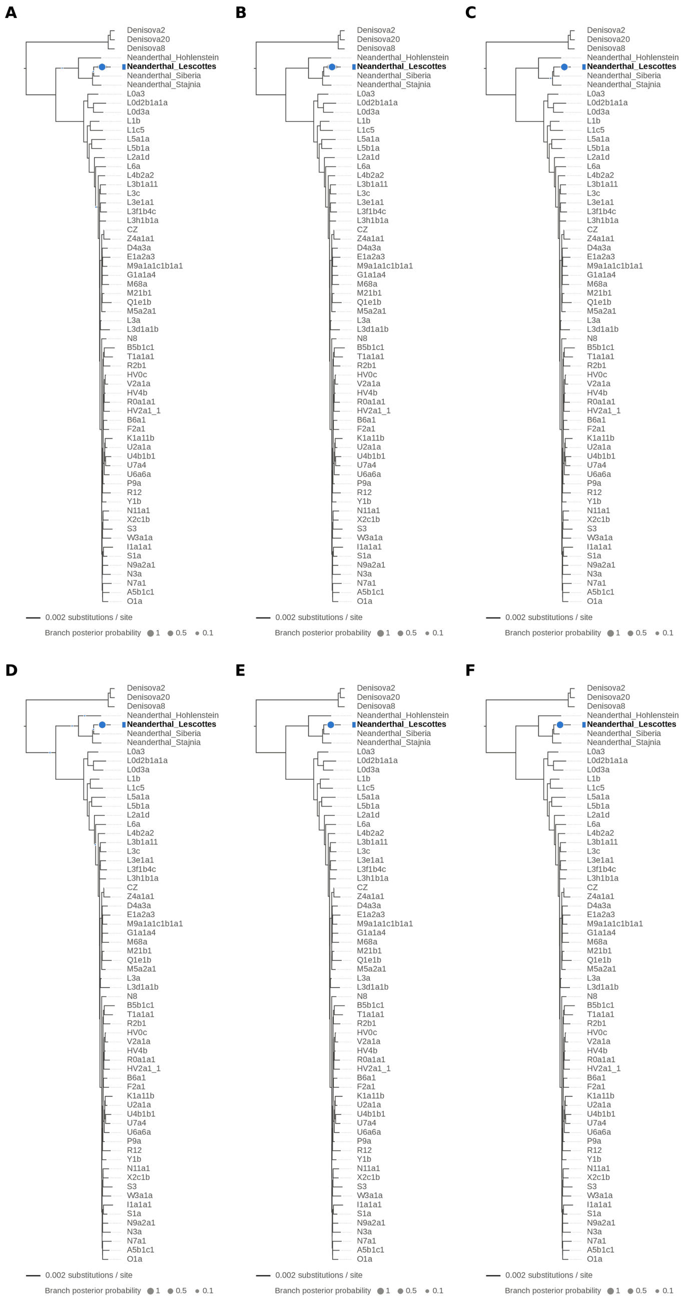


Figure S2: **Single-source simulated sample: Chagyrskaya 8 Neanderthal (MK388903.1), replicate 1, at nominal mean coverage depths of 0.05X, 0.1X, 0.5X, 1X, 2X and 4X (panels A–F).** Each panel shows the phylogenetic placement posterior of the best of five chains ($k = 1$, 100,000 iterations each) on the reference tree: the area of a dot is the posterior probability of the branch, the shaded swelling along a branch is the posterior density of the position on that branch, and the tip(s) of the highest-posterior branches are in bold. The expected placement is within the Neanderthal clade, on or near the Neanderthal_Lescottes branch (its closest relative in the reference). Replicates 1 and 2 are independent runs with identical settings. The runs at 0.5X and 4X are also shown, with collapsed trees, in main Figure 3. Modal branch of the best chain ($p$: its posterior probability): (A) 0.05X: Neanderthal_Lescottes ($p = 1.00$); (B) 0.1X: Neanderthal_Lescottes ($p = 1.00$); (C) 0.5X: Neanderthal_Lescottes ($p = 1.00$); (D) 1X: Neanderthal_Lescottes ($p = 1.00$); (E) 2X: Neanderthal_Lescottes ($p = 1.00$); (F) 4X: Neanderthal_Lescottes ($p = 1.00$).

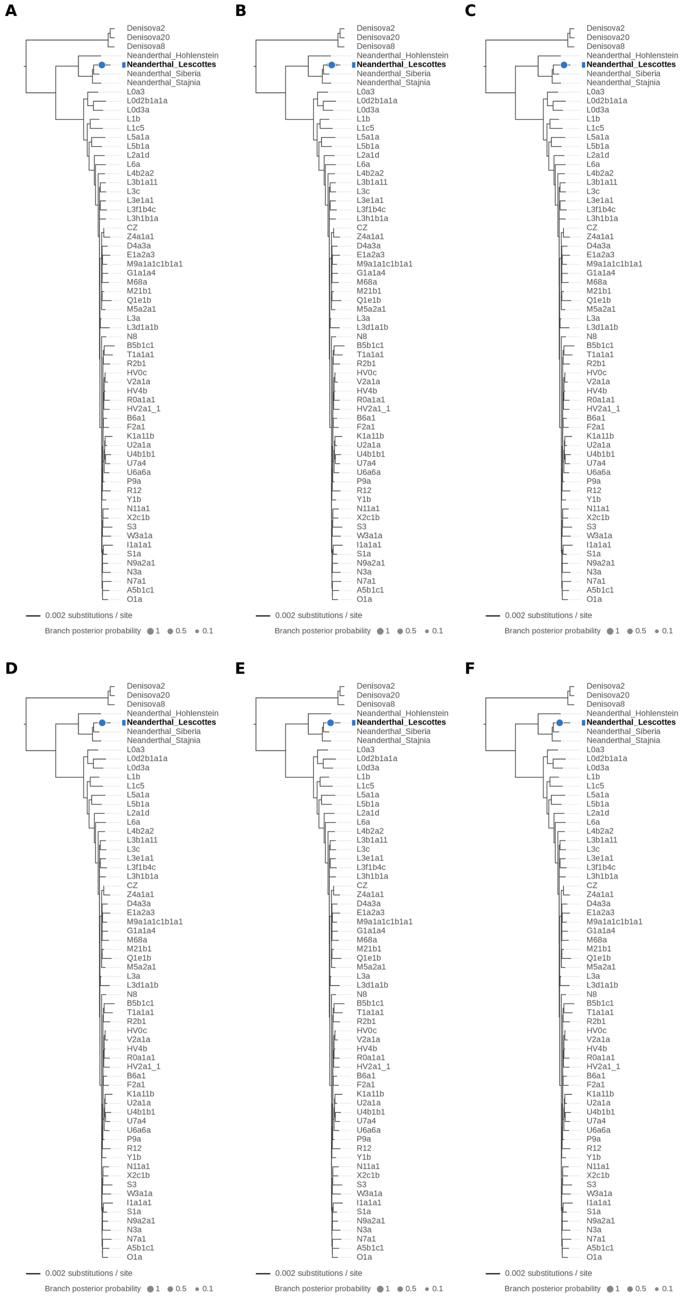


Figure S3: **Single-source simulated sample: Chagyrskaya 8 Neanderthal (MK388903.1), replicate 2, at nominal mean coverage depths of 0.05X, 0.1X, 0.5X, 1X, 2X and 4X (panels A–F).** Each panel shows the phylogenetic placement posterior of the best of five chains ($k$ = 1, 100,000 iterations each) on the reference tree: the area of a dot is the posterior probability of the branch, the shaded swelling along a branch is the posterior density of the position on that branch, and the tip(s) of the highest-posterior branches are in bold. The expected placement is within the Neanderthal clade, on or near the Neanderthal_Lescottes branch (its closest relative in the reference). Replicates 1 and 2 are independent runs with identical settings. Modal branch of the best chain ($p$: its posterior probability): (A) 0.05X: Neanderthal_Lescottes ($p = 1.00$); (B) 0.1X: Neanderthal_Lescottes ($p = 1.00$); (C) 0.5X: Neanderthal_Lescottes ($p = 1.00$); (D) 1X: Neanderthal_Lescottes ($p = 1.00$); (E) 2X: Neanderthal_Lescottes ($p = 1.00$); (F) 4X: Neanderthal_Lescottes ($p = 1.00$).

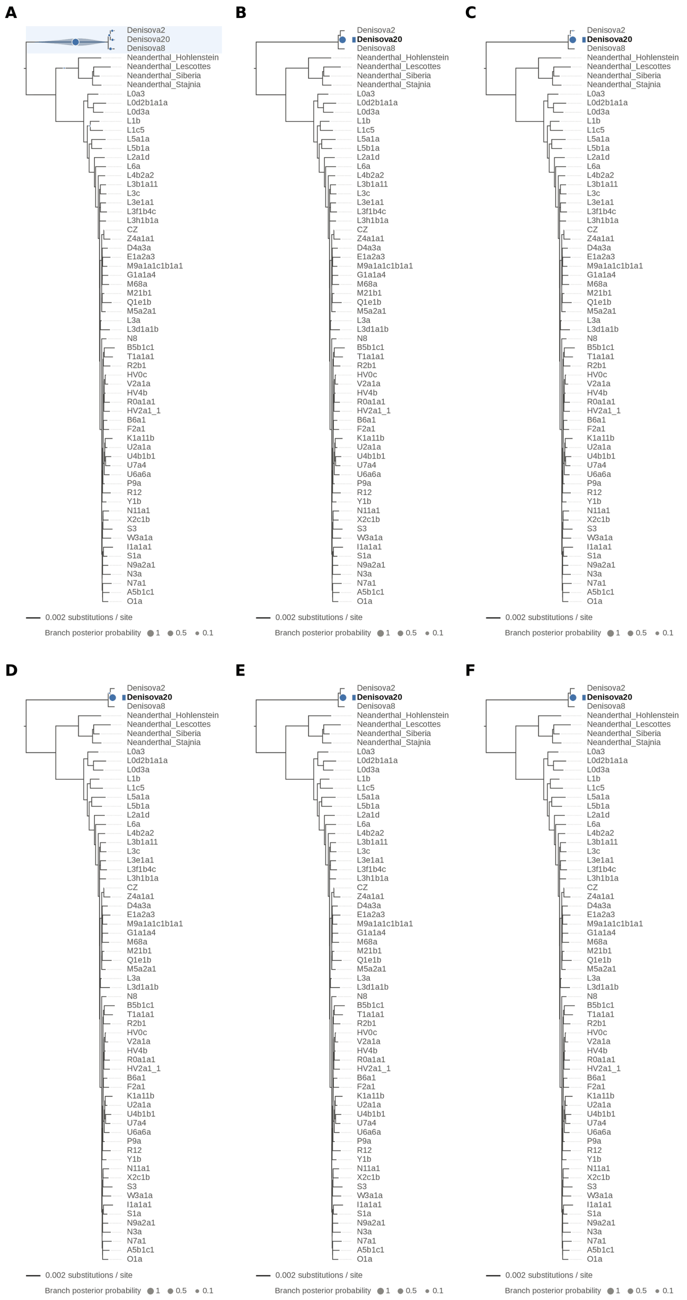


Figure S4: **Single-source simulated sample: Denisova 21 (MT576653.1), replicate 1, at nominal mean coverage depths of 0.05X, 0.1X, 0.5X, 1X, 2X and 4X (panels A–F).** Each panel shows the phylogenetic placement posterior of the best of five chains ($k = 1$, 100,000 iterations each) on the reference tree: the area of a dot is the posterior probability of the branch, the shaded swelling along a branch is the posterior density of the position on that branch, and the tip(s) of the highest-posterior branches are in bold. The expected placement is on the Denisova20 branch (its closest relative in the reference). Replicates 1 and 2 are independent runs with identical settings. Modal branch of the best chain ($p$: its posterior probability): (A) 0.05X: Node45 (stem of the Denisovan clade; $p = 0.95$); (B) 0.1X: Denisova20 ($p = 1.00$); (C) 0.5X: Denisova20 ($p = 1.00$); (D) 1X: Denisova20 ($p = 1.00$); (E) 2X: Denisova20 ($p = 1.00$); (F) 4X: Denisova20 ($p = 1.00$).

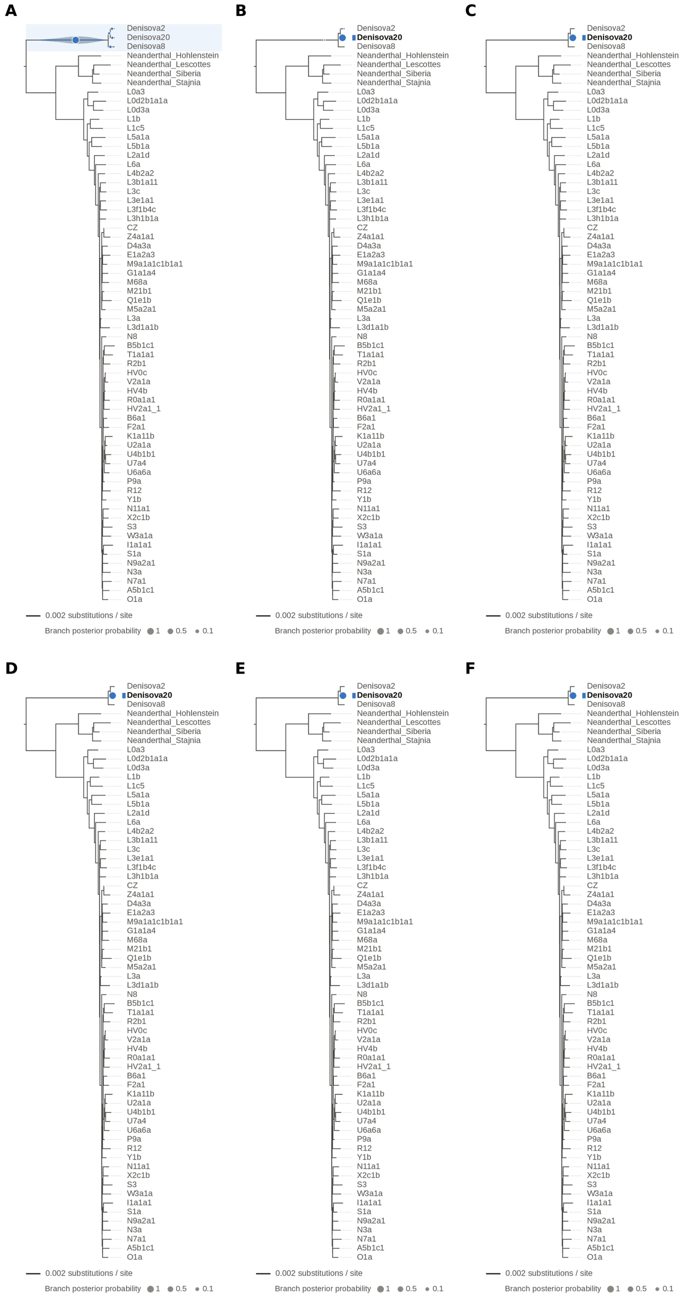


Figure S5: **Single-source simulated sample: Denisova 21 (MT576653.1), replicate 2, at nominal mean coverage depths of 0.05X, 0.1X, 0.5X, 1X, 2X and 4X (panels A–F).** Each panel shows the phylogenetic placement posterior of the best of five chains ($k = 1$, 100,000 iterations each) on the reference tree: the area of a dot is the posterior probability of the branch, the shaded swelling along a branch is the posterior density of the position on that branch, and the tip(s) of the highest-posterior branches are in bold. The expected placement is on the Denisova20 branch (its closest relative in the reference). Replicates 1 and 2 are independent runs with identical settings. Modal branch of the best chain ($p$: its posterior probability): (A) 0.05X: Node45 (stem of the Denisovan clade; $p = 0.95$); (B) 0.1X: Denisova20 ($p = 1.00$); (C) 0.5X: Denisova20 ($p = 1.00$); (D) 1X: Denisova20 ($p = 1.00$); (E) 2X: Denisova20 ($p = 1.00$); (F) 4X: Denisova20 ($p = 1.00$).

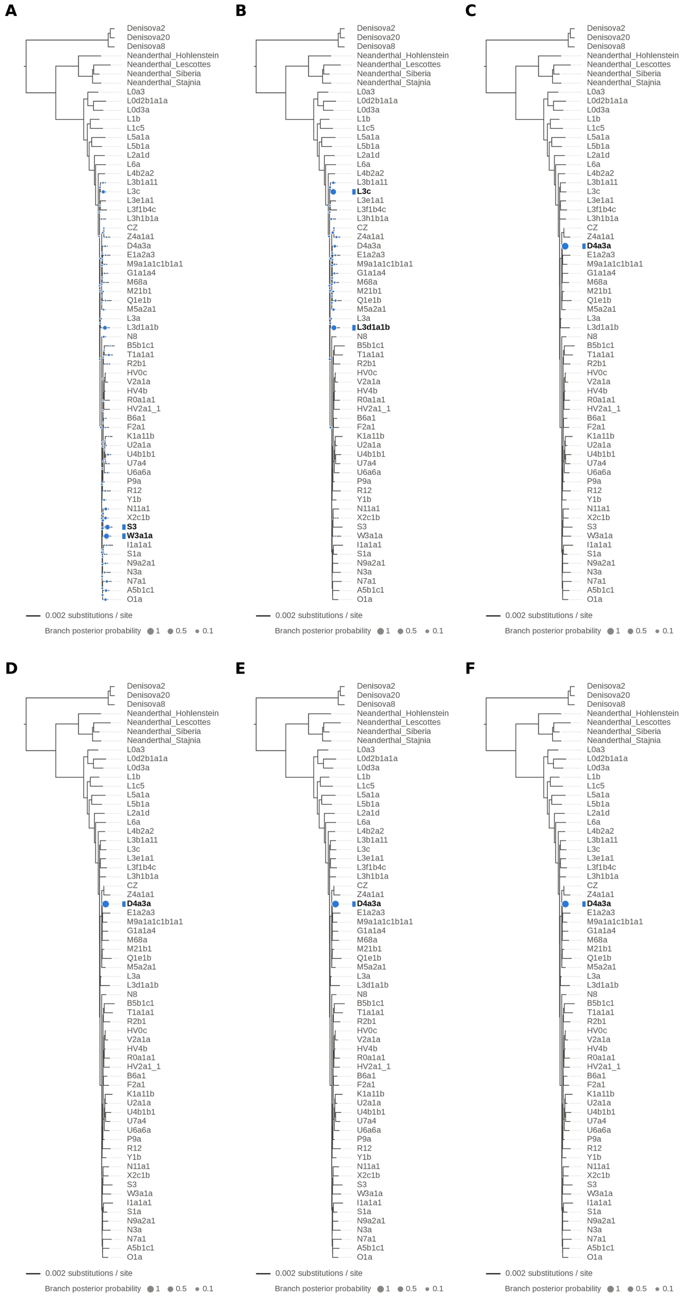


Figure S6: **Single-source simulated sample: modern human D4a1d (FJ951517.1), replicate 1, at nominal mean coverage depths of 0.05X, 0.1X, 0.5X, 1X, 2X and 4X (panels A–F).** Each panel shows the phylogenetic placement posterior of the best of five chains ($k = 1$, 100,000 iterations each) on the reference tree: the area of a dot is the posterior probability of the branch, the shaded swelling along a branch is the posterior density of the position on that branch, and the tip(s) of the highest-posterior branches are in bold. The expected placement is on the D4a3a branch (its closest relative in the reference). Replicates 1 and 2 are independent runs with identical settings. The runs at 0.1X and 1X are also shown, with collapsed trees, in main Figure 3. Modal branch of the best chain ($p$: its posterior probability): (A) 0.05X: W3a1a ($p = 0.32$), followed by S3 ($p = 0.24$) and L3d1a1b ($p = 0.11$); (B) 0.1X: L3c ($p = 0.48$), followed by L3d1a1b ($p = 0.30$) and D4a3a ($p = 0.02$); (C) 0.5X: D4a3a ($p = 1.00$); (D) 1X: D4a3a ($p = 1.00$); (E) 2X: D4a3a ($p = 1.00$); (F) 4X: D4a3a ($p = 1.00$).

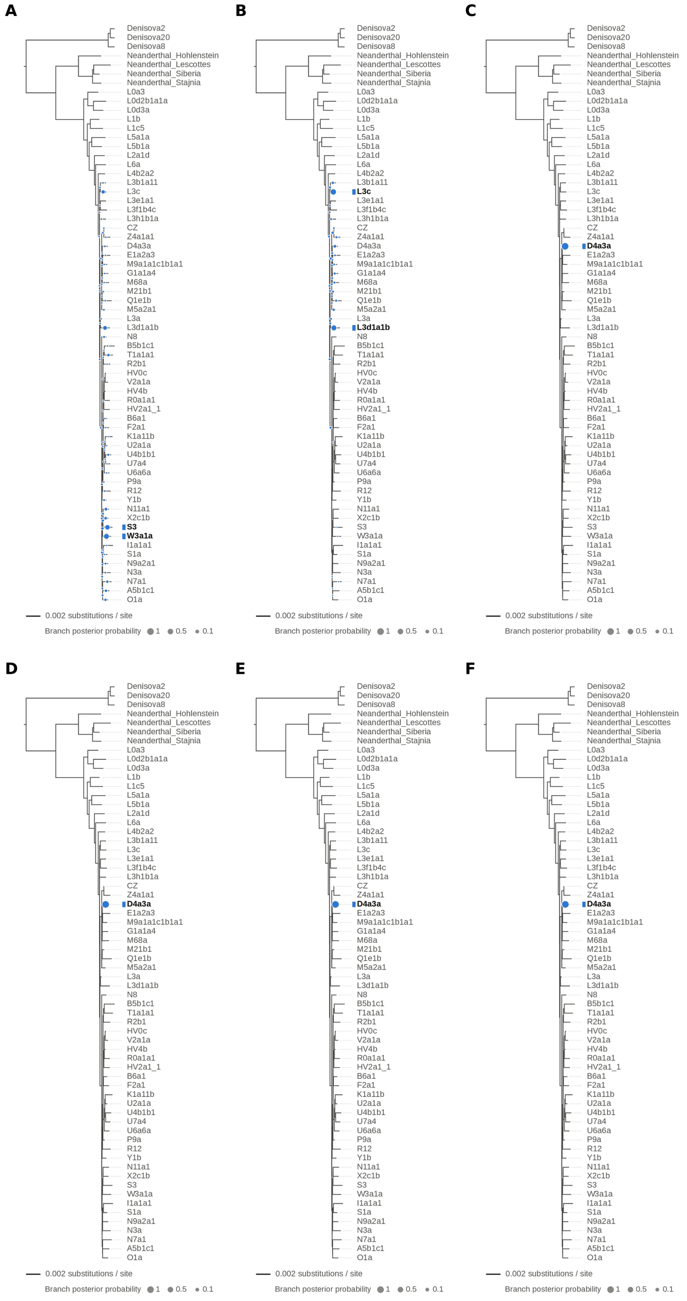


Figure S7: **Single-source simulated sample: modern human D4a1d (FJ951517.1), replicate 2, at nominal mean coverage depths of 0.05X, 0.1X, 0.5X, 1X, 2X and 4X (panels A–F).** Each panel shows the phylogenetic placement posterior of the best of five chains ($k = 1$, 100,000 iterations each) on the reference tree: the area of a dot is the posterior probability of the branch, the shaded swelling along a branch is the posterior density of the position on that branch, and the tip(s) of the highest-posterior branches are in bold. The expected placement is on the D4a3a branch (its closest relative in the reference). Replicates 1 and 2 are independent runs with identical settings. Modal branch of the best chain ($p$: its posterior probability): (A) 0.05X: W3a1a ($p = 0.31$), followed by S3 ($p = 0.22$) and L3d1a1b ($p = 0.11$); (B) 0.1X: L3c ($p = 0.44$), followed by L3d1a1b ($p = 0.34$) and D4a3a ($p = 0.03$); (C) 0.5X: D4a3a ($p = 1.00$); (D) 1X: D4a3a ($p = 1.00$); (E) 2X: D4a3a ($p = 1.00$); (F) 4X: D4a3a ($p = 1.00$).

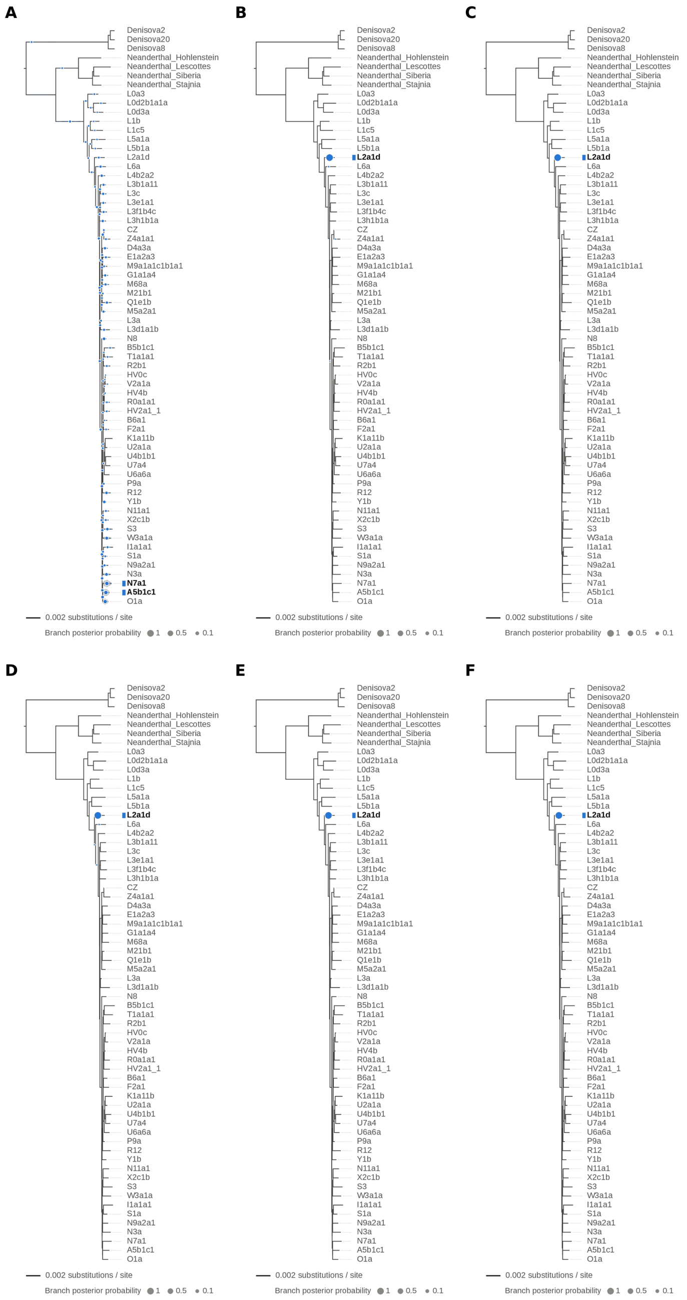


Figure S8: **Single-source simulated sample: modern human L2 (AF381981.1), replicate 1, at nominal mean coverage depths of 0.05X, 0.1X, 0.5X, 1X, 2X and 4X (panels A–F).** Each panel shows the phylogenetic placement posterior of the best of five chains ($k = 1$, 100,000 iterations each) on the reference tree: the area of a dot is the posterior probability of the branch, the shaded swelling along a branch is the posterior density of the position on that branch, and the tip(s) of the highest-posterior branches are in bold. The expected placement is on the L2a1d branch, the only L2 haplotype in the reference. Replicates 1 and 2 are independent runs with identical settings. Modal branch of the best chain ($p$: its posterior probability): (A) 0.05X: A5b1c1 ($p = 0.08$), followed by N7a1 ($p = 0.08$) and O1a ($p = 0.07$); (B) 0.1X: L2a1d ($p = 1.00$); (C) 0.5X: L2a1d ($p = 1.00$); (D) 1X: L2a1d ($p = 1.00$); (E) 2X: L2a1d ($p = 1.00$); (F) 4X: L2a1d ($p = 1.00$).

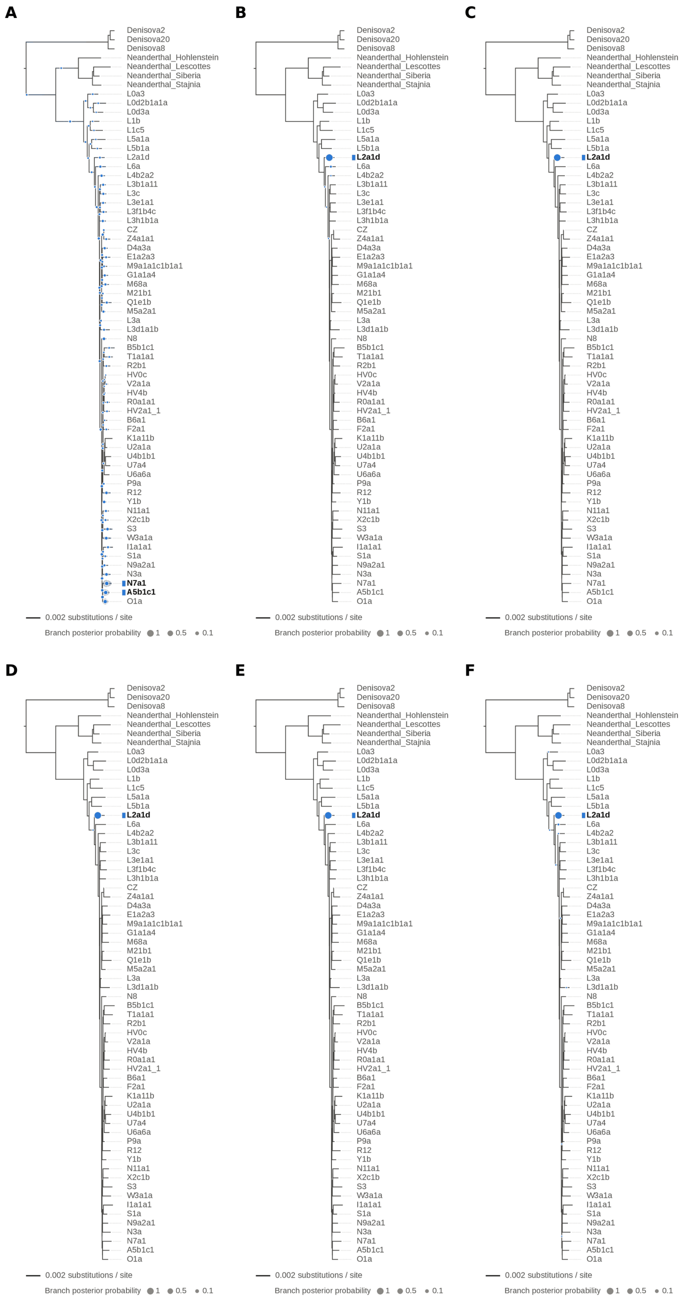

Figure S9: **Single-source simulated sample: modern human L2 (AF381981.1), replicate 2, at nominal mean coverage depths of 0.05X, 0.1X, 0.5X, 1X, 2X and 4X (panels A–F).** Each panel shows the phylogenetic placement posterior of the best of five chains ($k = 1$, 100,000 iterations each) on the reference tree: the area of a dot is the posterior probability of the branch, the shaded swelling along a branch is the posterior density of the position on that branch, and the tip(s) of the highest-posterior branches are in bold. The expected placement is on the L2a1d branch, the only L2 haplotype in the reference. Replicates 1 and 2 are independent runs with identical settings. Modal branch of the best chain ($p$: its posterior probability): (A) 0.05X: N7a1 ($p = 0.08$), followed by A5b1c1 ($p = 0.07$) and O1a ($p = 0.07$); (B) 0.1X: L2a1d ($p = 0.99$); (C) 0.5X: L2a1d ($p = 1.00$); (D) 1X: L2a1d ($p = 1.00$); (E) 2X: L2a1d ($p = 1.00$); (F) 4X: L2a1d ($p = 1.00$).

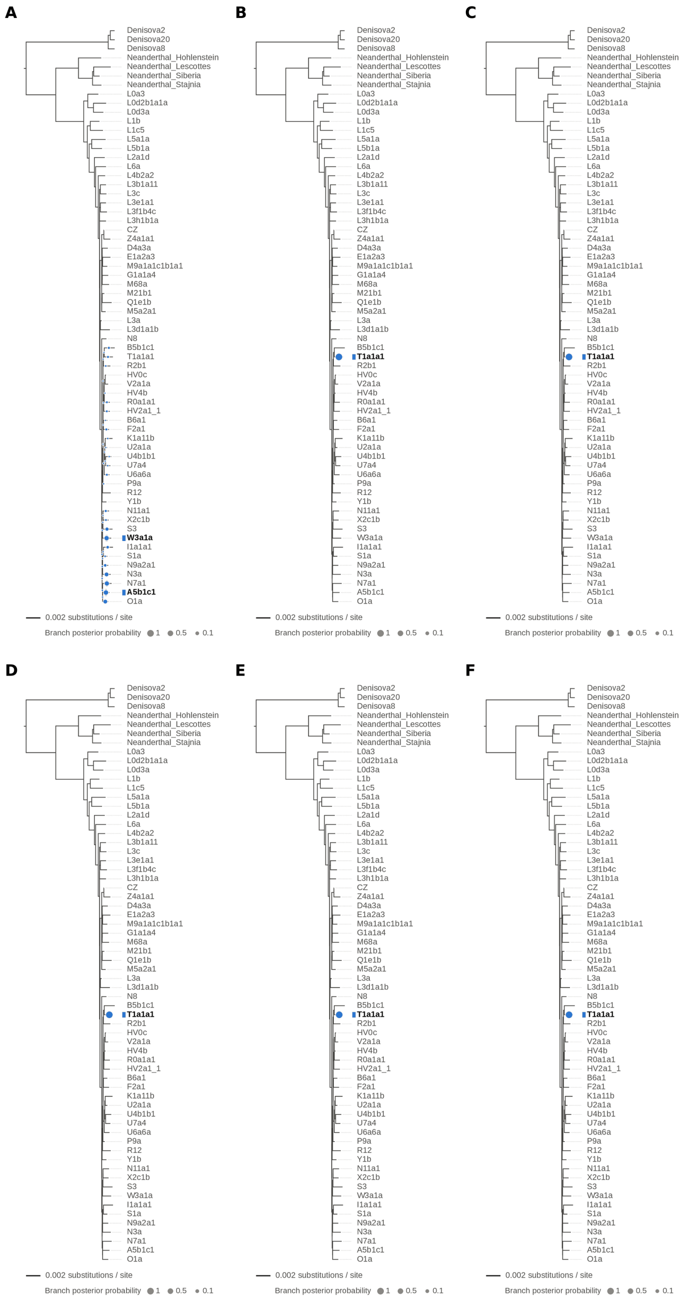


Figure S10: **Single-source simulated sample: modern human T2a1 (KY671094.1), replicate 1, at nominal mean coverage depths of 0.05X, 0.1X, 0.5X, 1X, 2X and 4X (panels A–F).** Each panel shows the phylogenetic placement posterior of the best of five chains ($k = 1$, 100,000 iterations each) on the reference tree: the area of a dot is the posterior probability of the branch, the shaded swelling along a branch is the posterior density of the position on that branch, and the tip(s) of the highest-posterior branches are in bold. The expected placement is on the T1a1a1 branch, the only haplogroup T haplotype in the reference. Replicates 1 and 2 are independent runs with identical settings. Modal branch of the best chain ($p$: its posterior probability): (A) 0.05X: A5b1c1 ($p = 0.22$), followed by W3a1a ($p = 0.19$) and N7a1 ($p = 0.18$); (B) 0.1X: T1a1a1 ($p = 1.00$); (C) 0.5X: T1a1a1 ($p = 1.00$); (D) 1X: T1a1a1 ($p = 1.00$); (E) 2X: T1a1a1 ($p = 1.00$); (F) 4X: T1a1a1 ($p = 1.00$).

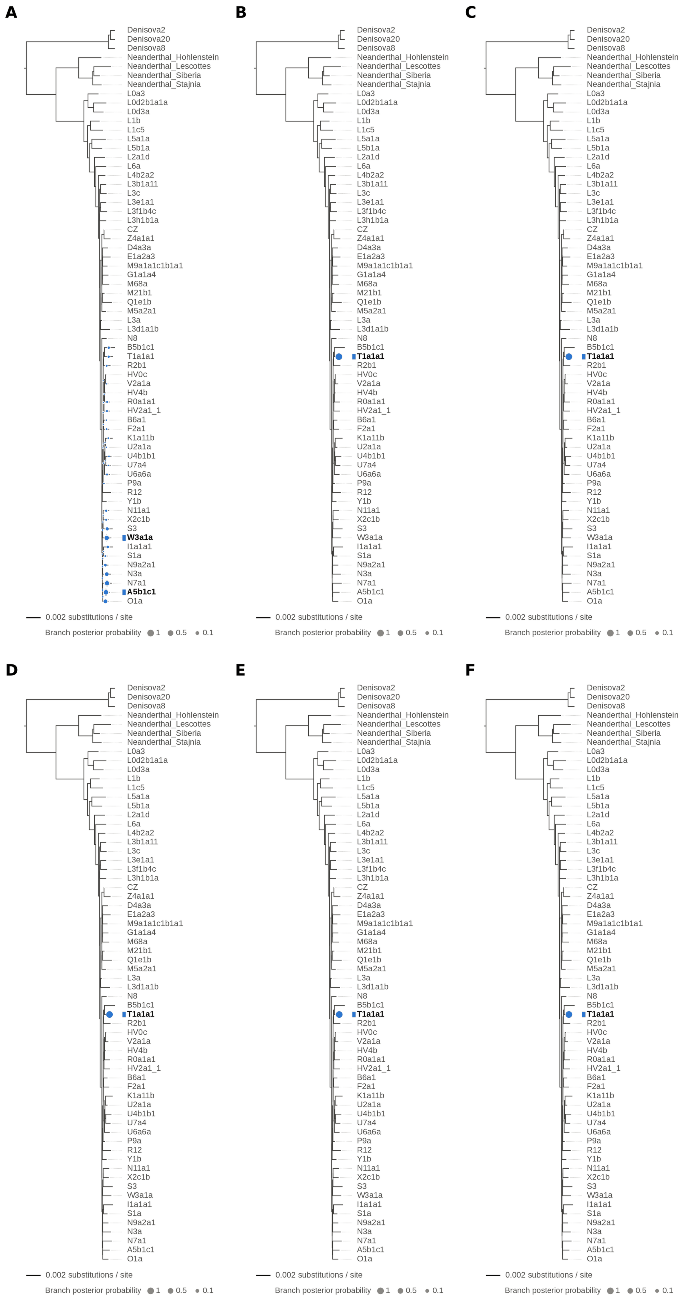


Figure S11: **Single-source simulated sample: modern human T2a1 (KY671094.1), replicate 2, at nominal mean coverage depths of 0.05X, 0.1X, 0.5X, 1X, 2X and 4X (panels A–F).** Each panel shows the phylogenetic placement posterior of the best of five chains ($k = 1$, 100,000 iterations each) on the reference tree: the area of a dot is the posterior probability of the branch, the shaded swelling along a branch is the posterior density of the position on that branch, and the tip(s) of the highest-posterior branches are in bold. The expected placement is on the T1a1a1 branch, the only haplogroup T haplotype in the reference. Replicates 1 and 2 are independent runs with identical settings. Modal branch of the best chain ($p$: its posterior probability): (A) 0.05X: A5b1c1 ($p = 0.22$), followed by W3a1a ($p = 0.19$) and N7a1 ($p = 0.19$); (B) 0.1X: T1a1a1 ($p = 1.00$); (C) 0.5X: T1a1a1 ($p = 1.00$); (D) 1X: T1a1a1 ($p = 1.00$); (E) 2X: T1a1a1 ($p = 1.00$); (F) 4X: T1a1a1 ($p = 1.00$).

#### S6 Two-source Simulated Mixtures

Here we show the `TrailMix` placements and mixture proportions for all 735 two-source mixtures: each of the five focal sources (Chagyrskaya 8, Denisova 21, D4a1d, L2 and T2a1) mixed with U5a2a at 21 proportions and 7 nominal mean coverage depths (Supplementary Figures S12–S116; one figure per mixture, with all its coverage depths). The runs of the 30% Denisova 21 mixture shown in main Figure 4, with collapsed trees, are shown here with the full tree. Table S3 summarizes how often each source was placed in its expected clade.

Note that the two sources of a mixture are exchangeable in the model, so the colors of the sources in the figures follow their estimated proportion (the most abundant source is shown in blue) rather than their identity.

| | 0.5X | 1X | 2X | 4X | 8X | 16X | 32X |
|---|---|---|---|---|---|---|---|
| *(a) Focal source placed in its expected clade or branch* | | | | | | | |
| Chagyrskaya 8 (Neanderthal clade) | 1.00 | 1.00 | 1.00 | 1.00 | 1.00 | 0.95 | 0.95 |
| Denisova 21 (Denisovan clade) | 0.95 | 0.95 | 0.95 | 1.00 | 0.95 | 0.95 | 0.95 |
| D4a1d (D4a3a branch) | 0.62 | 0.71 | 0.71 | 0.86 | 0.86 | 0.86 | 0.86 |
| L2 (L2a1d branch) | 0.52 | 0.71 | 0.81 | 0.71 | 0.76 | 0.71 | 0.76 |
| T2a1 (T1a1a1 branch) | 0.62 | 0.71 | 0.71 | 0.71 | 0.81 | 0.90 | 0.95 |
| *(b) U5a2a placed in the U clade, by focal source* | | | | | | | |
| Chagyrskaya 8 | 0.00 | 0.05 | 0.43 | 0.86 | 0.81 | 0.81 | 0.81 |
| Denisova 21 | 0.00 | 0.00 | 0.33 | 0.76 | 0.86 | 0.86 | 0.90 |
| D4a1d | 0.00 | 0.05 | 0.48 | 0.86 | 0.86 | 0.86 | 0.86 |
| L2 | 0.00 | 0.00 | 0.33 | 0.76 | 0.76 | 0.76 | 0.76 |
| T2a1 | 0.00 | 0.05 | 0.38 | 0.86 | 0.81 | 0.81 | 0.90 |
| *(c) Both sources placed correctly, pooled over focal sources, by share of the minor source* | | | | | | | |
| ≤5% ($n = 20$) | 0.00 | 0.00 | 0.15 | 0.20 | 0.15 | 0.10 | 0.15 |
| 10–20% ($n = 30$) | 0.00 | 0.00 | 0.27 | 0.53 | 0.50 | 0.57 | 0.67 |
| 25–50% ($n = 55$) | 0.00 | 0.05 | 0.27 | 0.93 | 1.00 | 0.98 | 1.00 |

Table S3: **Placement accuracy for the 735 simulated two-source mixtures.** Fraction of the mixtures (21 per source and coverage depth in (a) and (b)) in which the modal branch of one of the two sources in the best chain lies in the expected clade or on the expected branch. In (a), the expected placement is the clade or the branch of the closest relative in the reference (Table S1); in (b), the U clade. Since the sources are exchangeable, a source is counted as correctly placed if either of the two inferred sources is placed as expected; in (c), one inferred source must be placed as expected for the focal source and the other in the U clade. $n$: number of mixtures per coverage depth.

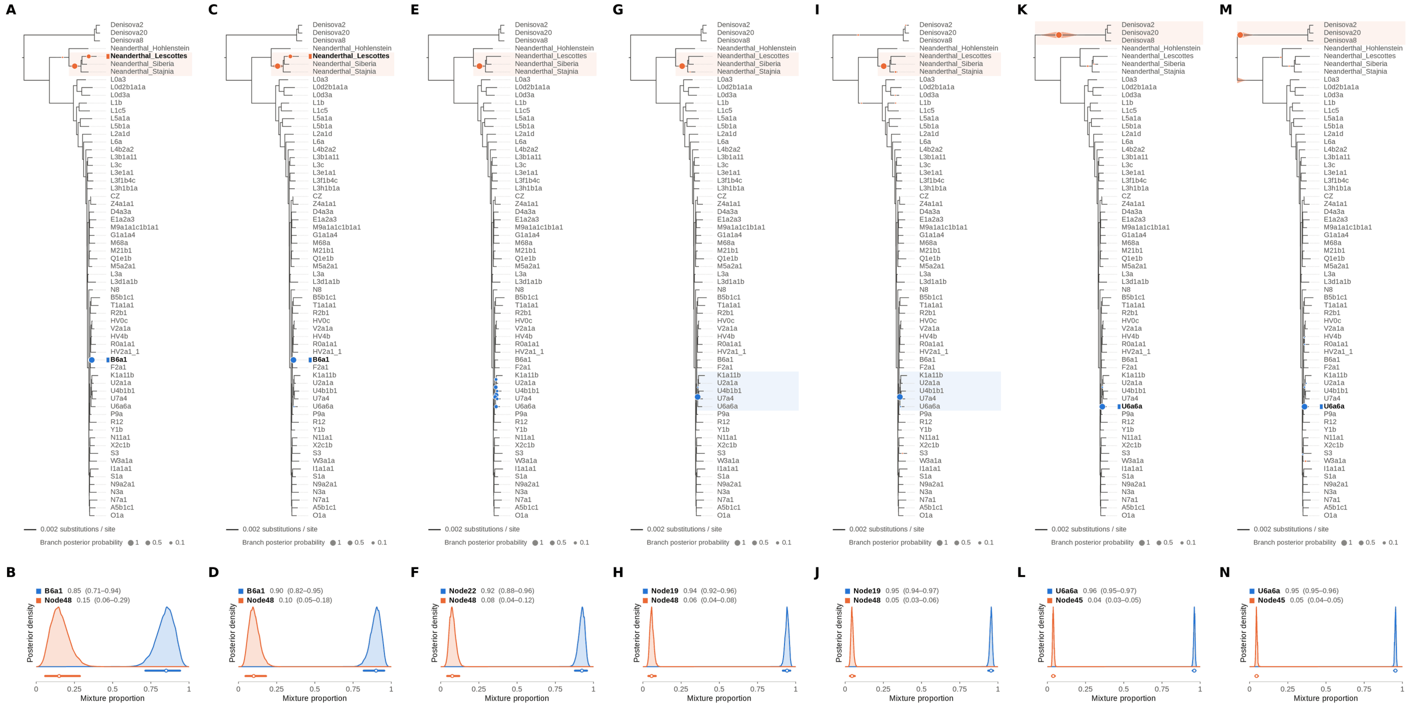


Figure S12: **Two-source simulated mixture of Chagyrskaya 8 Neanderthal (MK388903.1) (1% of reads) and modern human U5a2a (HM229344.1; 99%) at nominal mean coverage depths of 0.5X, 1X, 2X, 4X, 8X, 16X, 32X.** Both sources carry ancient DNA damage (U5a2a with the damage rates scaled by 0.75). For each coverage depth, the upper panel shows the phylogenetic placement posterior of both sources for the best of five chains ($k = 2$, 100,000 iterations each), colored by source (dot area: branch posterior probability; swelling: posterior density of the position along the branch; bold tips: highest-posterior branches), and the lower panel the posterior densities of the mixture proportions $\Theta$ with median and 95% credible interval. Chagyrskaya 8 is expected within the Neanderthal clade, on or near the Neanderthal_Lescottes branch (its closest relative in the reference); U5a2a, for which no U5 haplotype is present in the reference, is expected within the U clade. Modal branch of each source in the best chain ($p$: its posterior probability) and $\Theta$ median [95% CI]: (A,B) 0.5X: B6a1 ($p = 1.00$), $\Theta = 0.85$ [0.71, 0.94]; Node48 (internal branch ancestral to Neanderthal_Lescottes, Neanderthal_Siberia and Neanderthal_Stajnia; $p = 0.76$), $\Theta = 0.15$ [0.06, 0.29]; (C,D) 1X: B6a1 ($p = 1.00$), $\Theta = 0.90$ [0.82, 0.95]; Node48 (internal branch ancestral to Neanderthal_Lescottes, Neanderthal_Siberia and Neanderthal_Stajnia; $p = 0.81$), $\Theta = 0.10$ [0.05, 0.18]; (E,F) 2X: Node22 (internal branch ancestral to U4b1b1 and U7a4; $p = 0.33$), $\Theta = 0.92$ [0.88, 0.96]; Node48 (internal branch ancestral to Neanderthal_Lescottes, Neanderthal_Siberia and Neanderthal_Stajnia; $p = 1.00$), $\Theta = 0.08$ [0.04, 0.12]; (G,H) 4X: Node19 (stem of the U clade; $p = 1.00$), $\Theta = 0.94$ [0.92, 0.96]; Node48 (internal branch ancestral to Neanderthal_Lescottes, Neanderthal_Siberia and Neanderthal_Stajnia; $p = 1.00$), $\Theta = 0.06$ [0.04, 0.08]; (I,J) 8X: Node19 (stem of the U clade; $p = 1.00$), $\Theta = 0.95$ [0.94, 0.97]; Node48 (internal branch ancestral to Neanderthal_Lescottes, Neanderthal_Siberia and Neanderthal_Stajnia; $p = 0.99$), $\Theta = 0.05$ [0.03, 0.06]; (K,L) 16X: U6a6a ($p = 1.00$), $\Theta = 0.96$ [0.95, 0.97]; Node45 (stem of the Denisovan clade; $p = 1.00$), $\Theta = 0.04$ [0.03, 0.05]; (M,N) 32X: U6a6a ($p = 1.00$), $\Theta = 0.95$ [0.95, 0.96]; Node45 (stem of the Denisovan clade; $p = 1.00$), $\Theta = 0.05$ [0.04, 0.05].

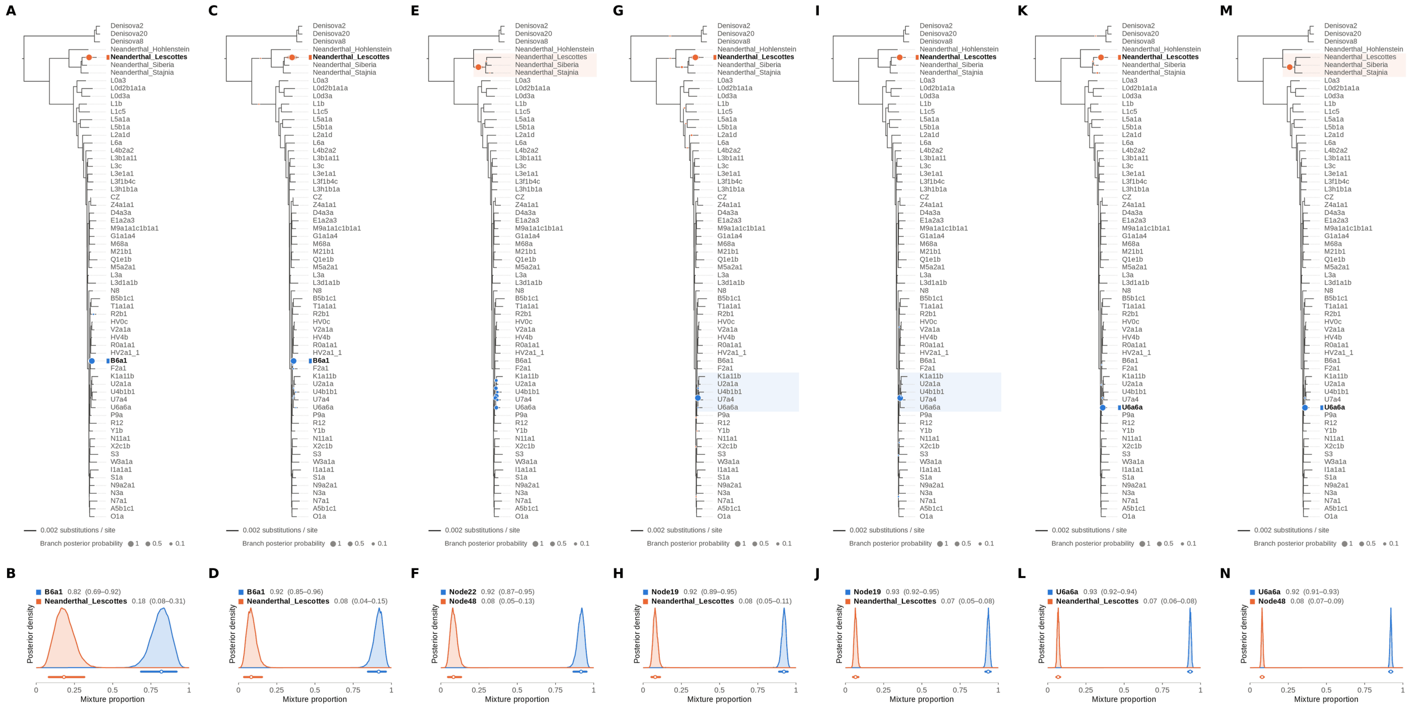

Figure S13: **Two-source simulated mixture of Chagyrskaya 8 Neanderthal (MK388903.1) (5% of reads) and modern human U5a2a (HM229344.1; 95%) at nominal mean coverage depths of 0.5X, 1X, 2X, 4X, 8X, 16X, 32X.** Both sources carry ancient DNA damage (U5a2a with the damage rates scaled by 0.75). For each coverage depth, the upper panel shows the phylogenetic placement posterior of both sources for the best of five chains ($k = 2$, 100,000 iterations each), colored by source (dot area: branch posterior probability; swelling: posterior density of the position along the branch; bold tips: highest-posterior branches), and the lower panel the posterior densities of the mixture proportions $\Theta$ with median and 95% credible interval. Chagyrskaya 8 is expected within the Neanderthal clade, on or near the Neanderthal_Lescottes branch (its closest relative in the reference); U5a2a, for which no U5 haplotype is present in the reference, is expected within the U clade. Modal branch of each source in the best chain ($p$: its posterior probability) and $\Theta$ median [95% CI]: (A,B) 0.5X: B6a1 ($p = 1.00$), $\Theta = 0.82$ [0.69, 0.92]; Neanderthal_Lescottes ($p = 1.00$), $\Theta = 0.18$ [0.08, 0.31]; (C,D) 1X: B6a1 ($p = 1.00$), $\Theta = 0.92$ [0.85, 0.96]; Neanderthal_Lescottes ($p = 1.00$), $\Theta = 0.08$ [0.04, 0.15]; (E,F) 2X: Node22 (internal branch ancestral to U4b1b1 and U7a4; $p = 0.31$), $\Theta = 0.92$ [0.87, 0.95]; Node48 (internal branch ancestral to Neanderthal_Lescottes, Neanderthal_Siberia and Neanderthal_Stajnia; $p = 1.00$), $\Theta = 0.08$ [0.05, 0.13]; (G,H) 4X: Node19 (stem of the U clade; $p = 1.00$), $\Theta = 0.92$ [0.89, 0.95]; Neanderthal_Lescottes ($p = 0.98$), $\Theta = 0.08$ [0.05, 0.11]; (I,J) 8X: Node19 (stem of the U clade; $p = 1.00$), $\Theta = 0.93$ [0.92, 0.95]; Neanderthal_Lescottes ($p = 1.00$), $\Theta = 0.07$ [0.05, 0.08]; (K,L) 16X: U6a6a ($p = 1.00$), $\Theta = 0.93$ [0.92, 0.94]; Neanderthal_Lescottes ($p = 1.00$), $\Theta = 0.07$ [0.06, 0.08]; (M,N) 32X: U6a6a ($p = 1.00$), $\Theta = 0.92$ [0.91, 0.93]; Node48 (internal branch ancestral to Neanderthal_Lescottes, Neanderthal_Siberia and Neanderthal_Stajnia; $p = 1.00$), $\Theta = 0.08$ [0.07, 0.09].

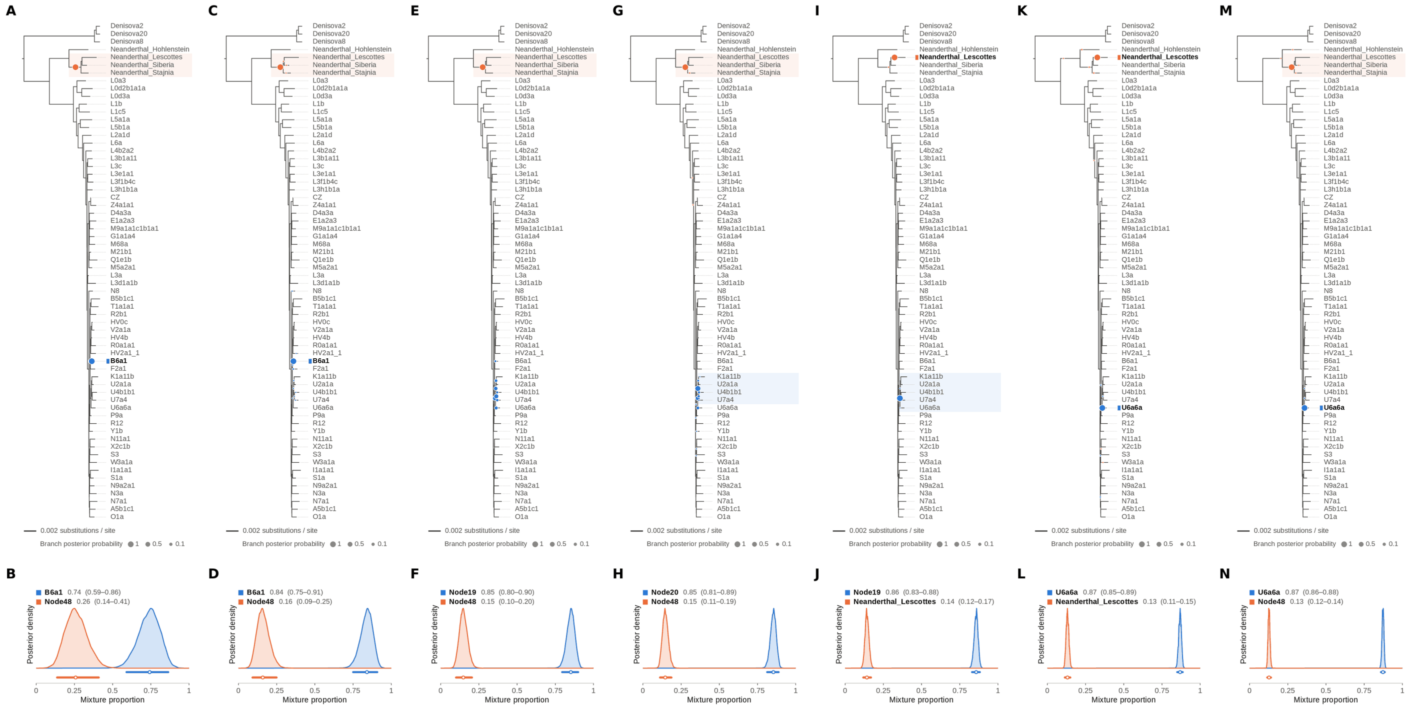

Figure S14: **Two-source simulated mixture of Chagyrskaya 8 Neanderthal (MK388903.1) (10% of reads) and modern human U5a2a (HM229344.1; 90%) at nominal mean coverage depths of 0.5X, 1X, 2X, 4X, 8X, 16X, 32X.** Both sources carry ancient DNA damage (U5a2a with the damage rates scaled by 0.75). For each coverage depth, the upper panel shows the phylogenetic placement posterior of both sources for the best of five chains ($k = 2$, 100,000 iterations each), colored by source (dot area: branch posterior probability; swelling: posterior density of the position along the branch; bold tips: highest-posterior branches), and the lower panel the posterior densities of the mixture proportions Θ with median and 95% credible interval. Chagyrskaya 8 is expected within the Neanderthal clade, on or near the Neanderthal_Lescottes branch (its closest relative in the reference); U5a2a, for which no U5 haplotype is present in the reference, is expected within the U clade. Modal branch of each source in the best chain ($p$: its posterior probability) and Θ median [95% CI]: (A,B) 0.5X: B6a1 ($p = 1.00$), $\Theta = 0.74$ [0.59, 0.86]; Node48 (internal branch ancestral to Neanderthal_Lescottes, Neanderthal_Siberia and Neanderthal_Stajnia; $p = 1.00$), $\Theta = 0.26$ [0.14, 0.41]; (C,D) 1X: B6a1 ($p = 0.99$), $\Theta = 0.84$ [0.75, 0.91]; Node48 (internal branch ancestral to Neanderthal_Lescottes, Neanderthal_Siberia and Neanderthal_Stajnia; $p = 1.00$), $\Theta = 0.16$ [0.09, 0.25]; (E,F) 2X: Node19 (stem of the U clade; $p = 0.38$), $\Theta = 0.85$ [0.80, 0.90]; Node48 (internal branch ancestral to Neanderthal_Lescottes, Neanderthal_Siberia and Neanderthal_Stajnia; $p = 1.00$), $\Theta = 0.15$ [0.10, 0.20]; (G,H) 4X: Node20 (internal branch within the U clade; $p = 0.62$), $\Theta = 0.85$ [0.81, 0.89]; Node48 (internal branch ancestral to Neanderthal_Lescottes, Neanderthal_Siberia and Neanderthal_Stajnia; $p = 1.00$), $\Theta = 0.15$ [0.11, 0.19]; (I,J) 8X: Node19 (stem of the U clade; $p = 1.00$), $\Theta = 0.86$ [0.83, 0.88]; Neanderthal_Lescottes ($p = 1.00$), $\Theta = 0.14$ [0.12, 0.17]; (K,L) 16X: U6a6a ($p = 1.00$), $\Theta = 0.87$ [0.85, 0.89]; Neanderthal_Lescottes ($p = 0.99$), $\Theta = 0.13$ [0.11, 0.15]; (M,N) 32X: U6a6a ($p = 1.00$), $\Theta = 0.87$ [0.86, 0.88]; Node48 (internal branch ancestral to Neanderthal_Lescottes, Neanderthal_Siberia and Neanderthal_Stajnia; $p = 1.00$), $\Theta = 0.13$ [0.12, 0.14].

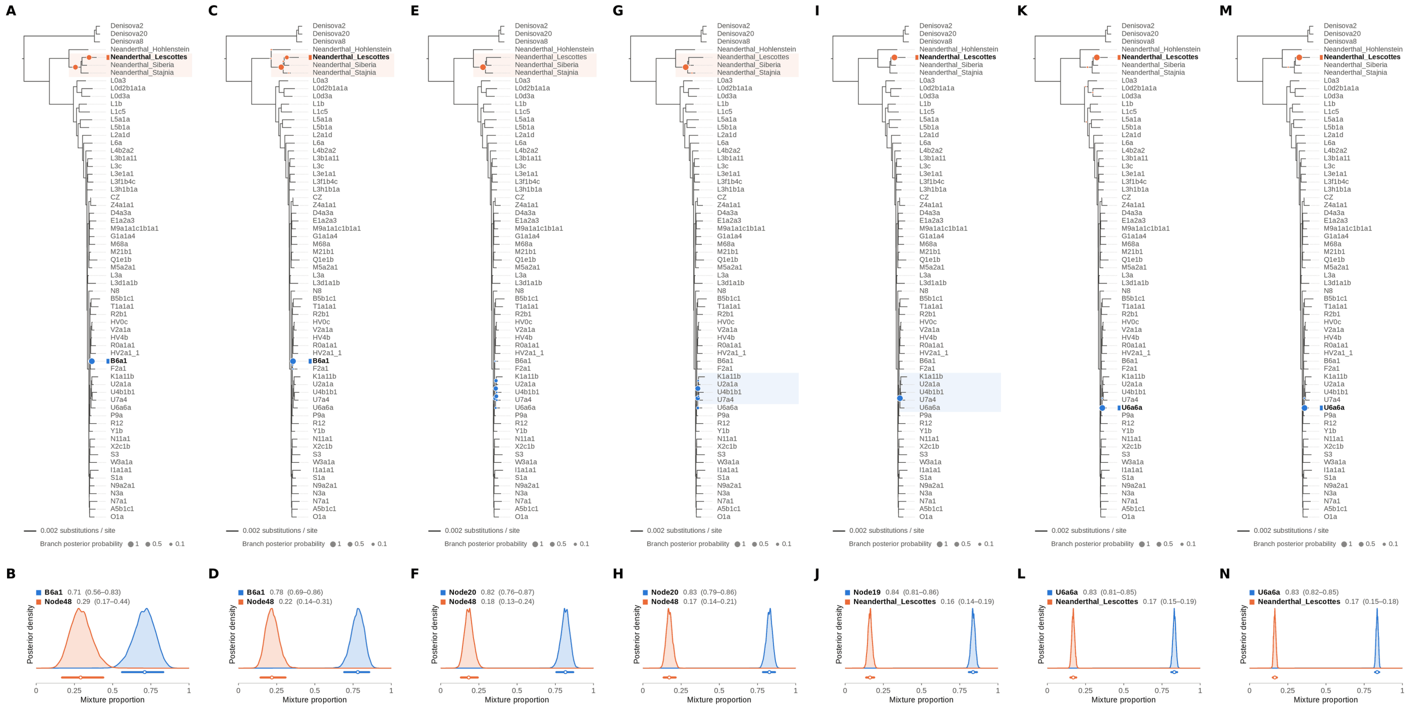


Figure S15: **Two-source simulated mixture of Chagyrskaya 8 Neanderthal (MK388903.1) (15% of reads) and modern human U5a2a (HM229344.1; 85%) at nominal mean coverage depths of 0.5X, 1X, 2X, 4X, 8X, 16X, 32X.** Both sources carry ancient DNA damage (U5a2a with the damage rates scaled by 0.75). For each coverage depth, the upper panel shows the phylogenetic placement posterior of both sources for the best of five chains ($k = 2$, 100,000 iterations each), colored by source (dot area: branch posterior probability; swelling: posterior density of the position along the branch; bold tips: highest-posterior branches), and the lower panel the posterior densities of the mixture proportions $\Theta$ with median and 95% credible interval. Chagyrskaya 8 is expected within the Neanderthal clade, on or near the Neanderthal_Lescottes branch (its closest relative in the reference); U5a2a, for which no U5 haplotype is present in the reference, is expected within the U clade. Modal branch of each source in the best chain ($p$: its posterior probability) and $\Theta$ median [95% CI]: (A,B) 0.5X: B6a1 ($p = 1.00$), $\Theta = 0.71$ [0.56, 0.83]; Node48 (internal branch ancestral to Neanderthal_Lescottes, Neanderthal_Siberia and Neanderthal_Stajnia; $p = 0.51$), $\Theta = 0.29$ [0.17, 0.44]; (C,D) 1X: B6a1 ($p = 1.00$), $\Theta = 0.78$ [0.69, 0.86]; Node48 (internal branch ancestral to Neanderthal_Lescottes, Neanderthal_Siberia and Neanderthal_Stajnia; $p = 0.77$), $\Theta = 0.22$ [0.14, 0.31]; (E,F) 2X: Node20 (internal branch within the U clade; $p = 0.35$), $\Theta = 0.82$ [0.76, 0.87]; Node48 (internal branch ancestral to Neanderthal_Lescottes, Neanderthal_Siberia and Neanderthal_Stajnia; $p = 1.00$), $\Theta = 0.18$ [0.13, 0.24]; (G,H) 4X: Node20 (internal branch within the U clade; $p = 0.59$), $\Theta = 0.83$ [0.79, 0.86]; Node48 (internal branch ancestral to Neanderthal_Lescottes, Neanderthal_Siberia and Neanderthal_Stajnia; $p = 1.00$), $\Theta = 0.17$ [0.14, 0.21]; (I,J) 8X: Node19 (stem of the U clade; $p = 1.00$), $\Theta = 0.84$ [0.81, 0.86]; Neanderthal_Lescottes ($p = 1.00$), $\Theta = 0.16$ [0.14, 0.19]; (K,L) 16X: U6a6a ($p = 1.00$), $\Theta = 0.83$ [0.81, 0.85]; Neanderthal_Lescottes ($p = 1.00$), $\Theta = 0.17$ [0.15, 0.19]; (M,N) 32X: U6a6a ($p = 1.00$), $\Theta = 0.83$ [0.82, 0.85]; Neanderthal_Lescottes ($p = 1.00$), $\Theta = 0.17$ [0.15, 0.18].

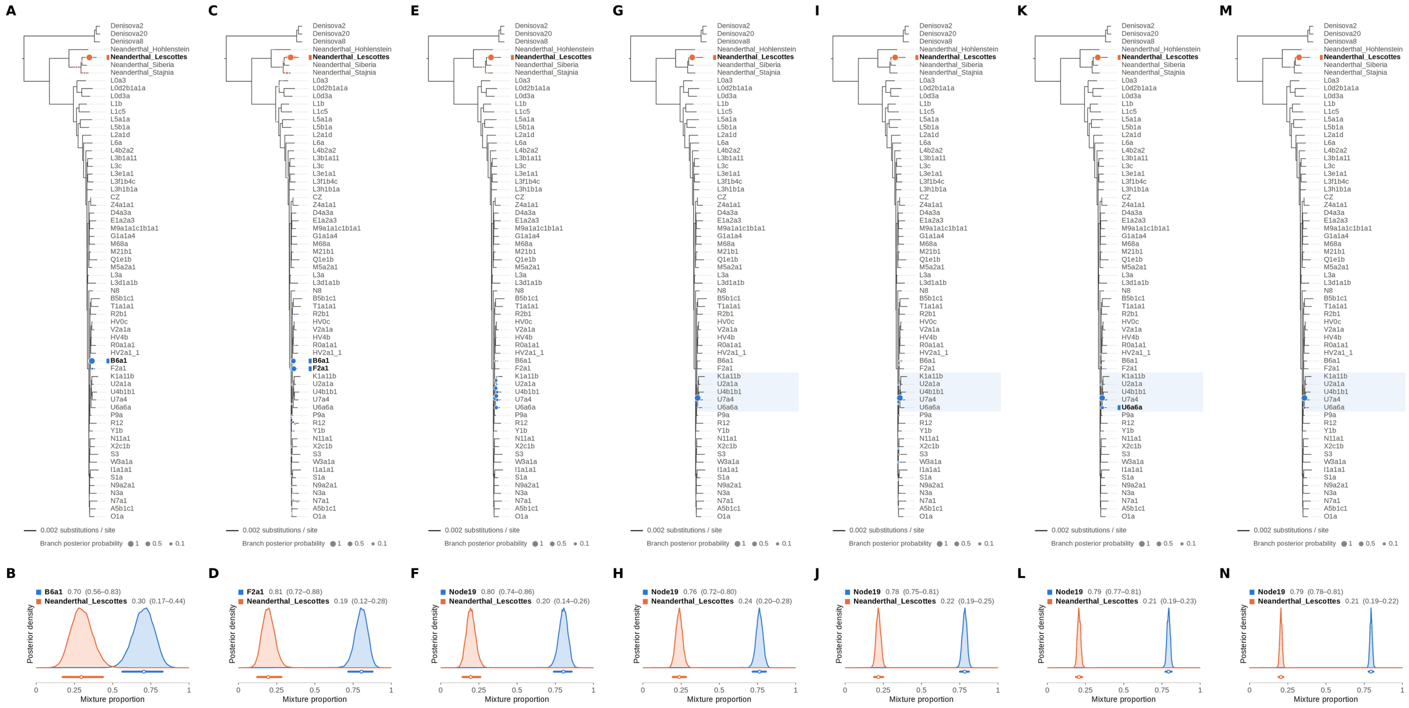


Figure S16: **Two-source simulated mixture of Chagyrskaya 8 Neanderthal (MK388903.1) (20% of reads) and modern human U5a2a (HM229344.1; 80%) at nominal mean coverage depths of 0.5X, 1X, 2X, 4X, 8X, 16X, 32X.** Both sources carry ancient DNA damage (U5a2a with the damage rates scaled by 0.75). For each coverage depth, the upper panel shows the phylogenetic placement posterior of both sources for the best of five chains ($k = 2$, 100,000 iterations each), colored by source (dot area: branch posterior probability; swelling: posterior density of the position along the branch; bold tips: highest-posterior branches), and the lower panel the posterior densities of the mixture proportions $\Theta$ with median and 95% credible interval. Chagyrskaya 8 is expected within the Neanderthal clade, on or near the Neanderthal_Lescottes branch (its closest relative in the reference); U5a2a, for which no U5 haplotype is present in the reference, is expected within the U clade. Modal branch of each source in the best chain ($p$: its posterior probability) and $\Theta$ median [95% CI]: (A,B) 0.5X: B6a1 ($p = 0.99$), $\Theta = 0.70$ [0.56, 0.83]; Neanderthal_Lescottes ($p = 0.99$), $\Theta = 0.30$ [0.17, 0.44]; (C,D) 1X: F2a1 ($p = 0.52$), $\Theta = 0.81$ [0.72, 0.88]; Neanderthal_Lescottes ($p = 0.98$), $\Theta = 0.19$ [0.12, 0.28]; (E,F) 2X: Node19 (stem of the U clade; $p = 0.35$), $\Theta = 0.80$ [0.74, 0.86]; Neanderthal_Lescottes ($p = 1.00$), $\Theta = 0.20$ [0.14, 0.26]; (G,H) 4X: Node19 (stem of the U clade; $p = 1.00$), $\Theta = 0.76$ [0.72, 0.80]; Neanderthal_Lescottes ($p = 1.00$), $\Theta = 0.24$ [0.20, 0.28]; (I,J) 8X: Node19 (stem of the U clade; $p = 1.00$), $\Theta = 0.78$ [0.75, 0.81]; Neanderthal_Lescottes ($p = 1.00$), $\Theta = 0.22$ [0.19, 0.25]; (K,L) 16X: Node19 (stem of the U clade; $p = 0.90$), $\Theta = 0.79$ [0.77, 0.81]; Neanderthal_Lescottes ($p = 1.00$), $\Theta = 0.21$ [0.19, 0.23]; (M,N) 32X: Node19 (stem of the U clade; $p = 0.98$), $\Theta = 0.79$ [0.78, 0.81]; Neanderthal_Lescottes ($p = 1.00$), $\Theta = 0.21$ [0.19, 0.22].

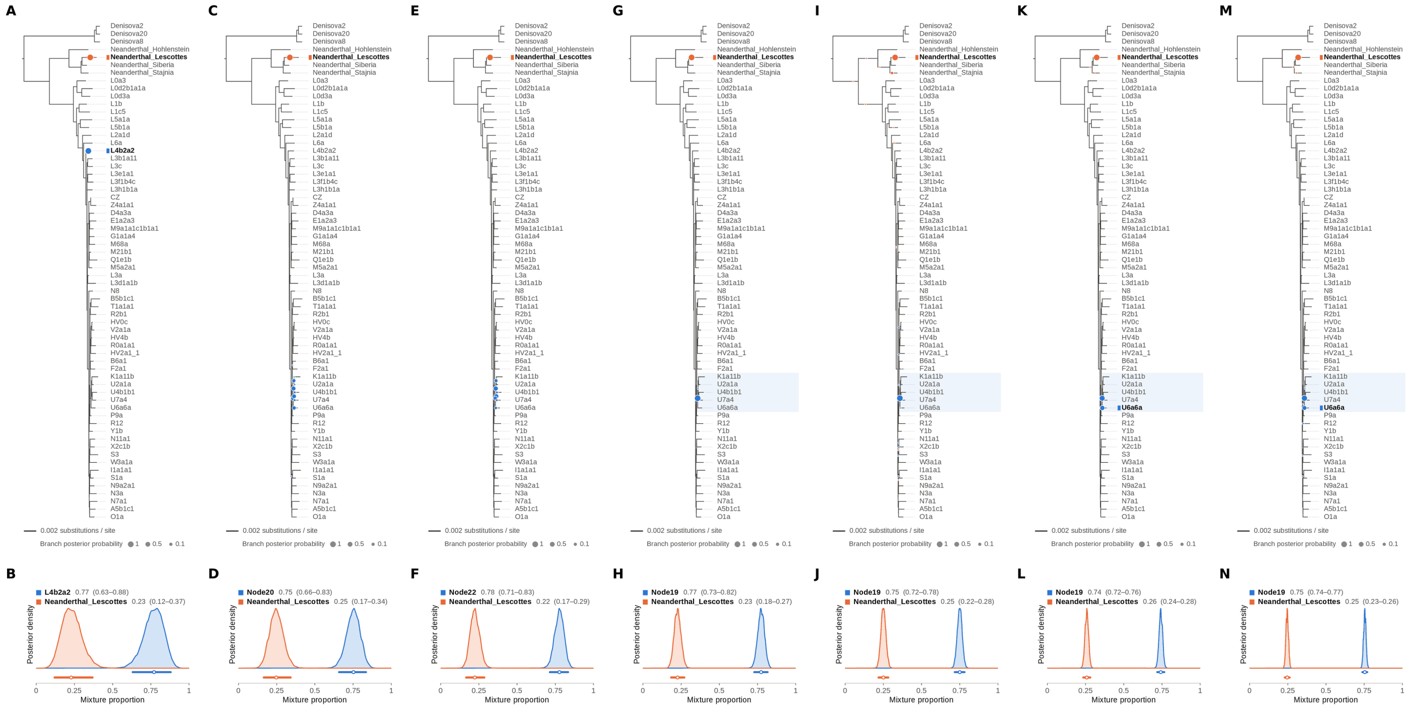


Figure S17: **Two-source simulated mixture of Chagyrskaya 8 Neanderthal (MK388903.1) (25% of reads) and modern human U5a2a (HM229344.1; 75%) at nominal mean coverage depths of 0.5X, 1X, 2X, 4X, 8X, 16X, 32X.** Both sources carry ancient DNA damage (U5a2a with the damage rates scaled by 0.75). For each coverage depth, the upper panel shows the phylogenetic placement posterior of both sources for the best of five chains ($k = 2$, 100,000 iterations each), colored by source (dot area: branch posterior probability; swelling: posterior density of the position along the branch; bold tips: highest-posterior branches), and the lower panel the posterior densities of the mixture proportions $\Theta$ with median and 95% credible interval. Chagyrskaya 8 is expected within the Neanderthal clade, on or near the Neanderthal_Lescottes branch (its closest relative in the reference); U5a2a, for which no U5 haplotype is present in the reference, is expected within the U clade. Modal branch of each source in the best chain ($p$: its posterior probability) and $\Theta$ median [95% CI]: (A,B) 0.5X: L4b2a2 ($p = 1.00$), $\Theta = 0.77$ [0.63, 0.88]; Neanderthal_Lescottes ($p = 1.00$), $\Theta = 0.23$ [0.12, 0.37]; (C,D) 1X: Node20 (internal branch within the U clade; $p = 0.26$), $\Theta = 0.75$ [0.66, 0.83]; Neanderthal_Lescottes ($p = 1.00$), $\Theta = 0.25$ [0.17, 0.34]; (E,F) 2X: Node22 (internal branch ancestral to U4b1b1 and U7a4; $p = 0.34$), $\Theta = 0.78$ [0.71, 0.83]; Neanderthal_Lescottes ($p = 1.00$), $\Theta = 0.22$ [0.17, 0.29]; (G,H) 4X: Node19 (stem of the U clade; $p = 1.00$), $\Theta = 0.77$ [0.73, 0.82]; Neanderthal_Lescottes ($p = 1.00$), $\Theta = 0.23$ [0.18, 0.27]; (I,J) 8X: Node19 (stem of the U clade; $p = 1.00$), $\Theta = 0.75$ [0.72, 0.78]; Neanderthal_Lescottes ($p = 0.96$), $\Theta = 0.25$ [0.22, 0.28]; (K,L) 16X: Node19 (stem of the U clade; $p = 0.65$), $\Theta = 0.74$ [0.72, 0.76]; Neanderthal_Lescottes ($p = 1.00$), $\Theta = 0.26$ [0.24, 0.28]; (M,N) 32X: Node19 (stem of the U clade; $p = 0.74$), $\Theta = 0.75$ [0.74, 0.77]; Neanderthal_Lescottes ($p = 1.00$), $\Theta = 0.25$ [0.23, 0.26].

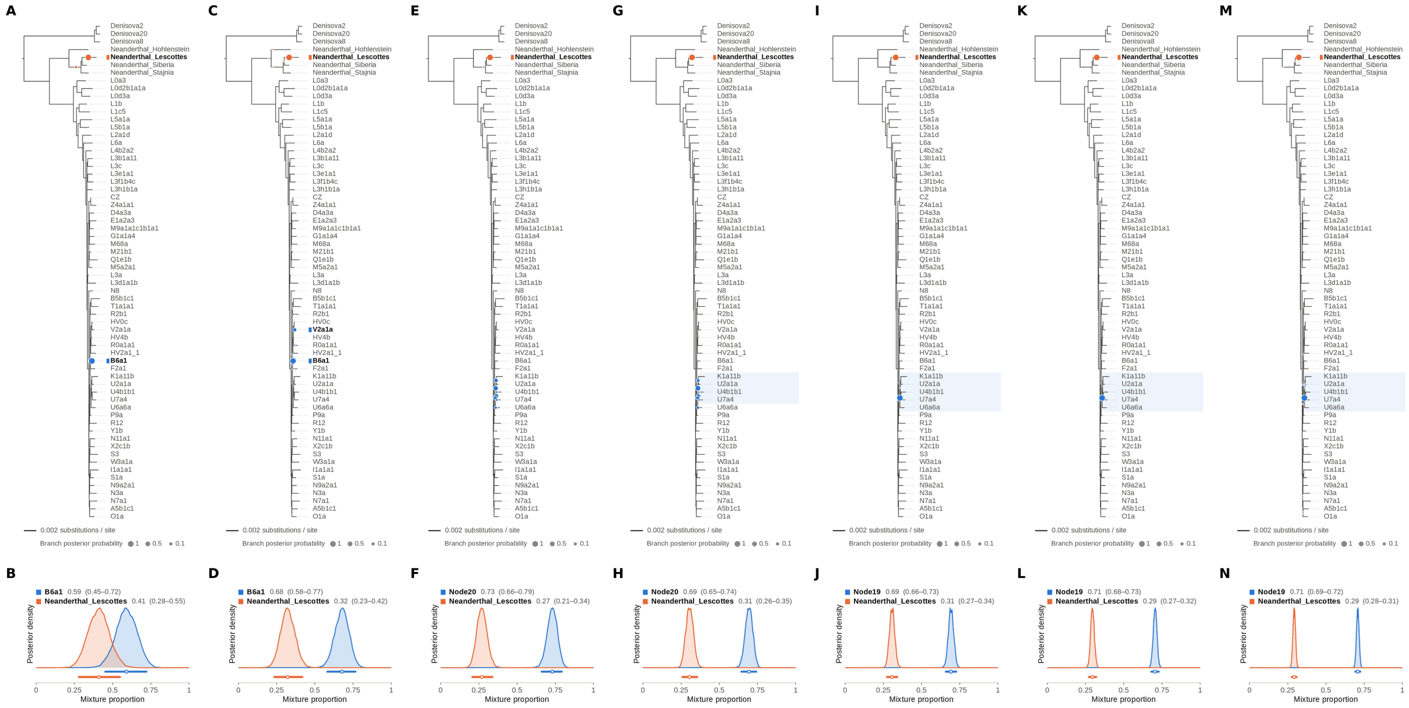


Figure S18: **Two-source simulated mixture of Chagyrskaya 8 Neanderthal (MK388903.1) (30% of reads) and modern human U5a2a (HM229344.1; 70%) at nominal mean coverage depths of 0.5X, 1X, 2X, 4X, 8X, 16X, 32X.** Both sources carry ancient DNA damage (U5a2a with the damage rates scaled by 0.75). For each coverage depth, the upper panel shows the phylogenetic placement posterior of both sources for the best of five chains ($k = 2$, 100,000 iterations each), colored by source (dot area: branch posterior probability; swelling: posterior density of the position along the branch; bold tips: highest-posterior branches), and the lower panel the posterior densities of the mixture proportions $\Theta$ with median and 95% credible interval. Chagyrskaya 8 is expected within the Neanderthal clade, on or near the Neanderthal_Lescottes branch (its closest relative in the reference); U5a2a, for which no U5 haplotype is present in the reference, is expected within the U clade. Modal branch of each source in the best chain ($p$: its posterior probability) and $\Theta$ median [95% CI]: (A,B) 0.5X: B6a1 ($p = 1.00$), $\Theta = 0.59$ [0.45, 0.72]; Neanderthal_Lescottes ($p = 0.98$), $\Theta = 0.41$ [0.28, 0.55]; (C,D) 1X: B6a1 ($p = 0.86$), $\Theta = 0.68$ [0.58, 0.77]; Neanderthal_Lescottes ($p = 1.00$), $\Theta = 0.32$ [0.23, 0.42]; (E,F) 2X: Node20 (internal branch within the U clade; $p = 0.38$), $\Theta = 0.73$ [0.66, 0.79]; Neanderthal_Lescottes ($p = 1.00$), $\Theta = 0.27$ [0.21, 0.34]; (G,H) 4X: Node20 (internal branch within the U clade; $p = 0.55$), $\Theta = 0.69$ [0.65, 0.74]; Neanderthal_Lescottes ($p = 1.00$), $\Theta = 0.31$ [0.26, 0.35]; (I,J) 8X: Node19 (stem of the U clade; $p = 1.00$), $\Theta = 0.69$ [0.66, 0.73]; Neanderthal_Lescottes ($p = 1.00$), $\Theta = 0.31$ [0.27, 0.34]; (K,L) 16X: Node19 (stem of the U clade; $p = 1.00$), $\Theta = 0.71$ [0.68, 0.73]; Neanderthal_Lescottes ($p = 1.00$), $\Theta = 0.29$ [0.27, 0.32]; (M,N) 32X: Node19 (stem of the U clade; $p = 1.00$), $\Theta = 0.71$ [0.69, 0.72]; Neanderthal_Lescottes ($p = 1.00$), $\Theta = 0.29$ [0.28, 0.31].

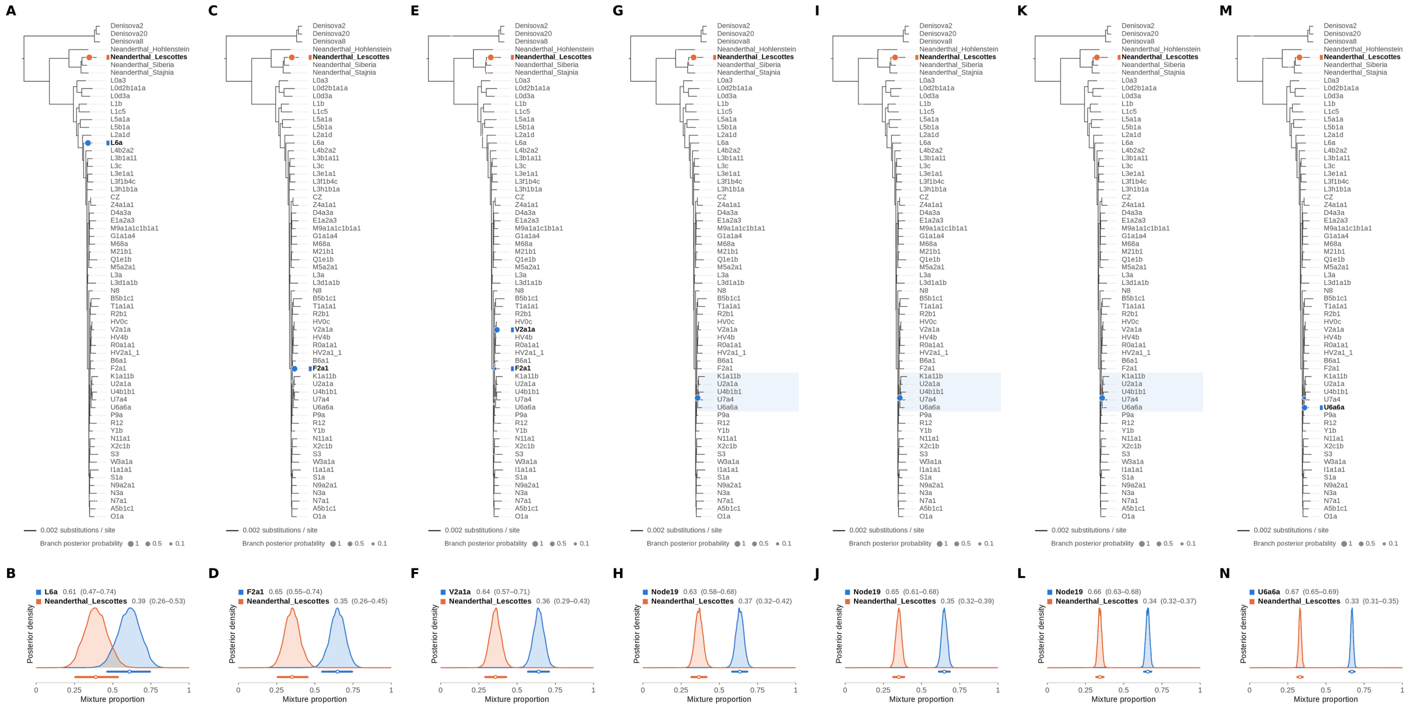


Figure S19: **Two-source simulated mixture of Chagyrskaya 8 Neanderthal (MK388903.1) (35% of reads) and modern human U5a2a (HM229344.1; 65%) at nominal mean coverage depths of 0.5X, 1X, 2X, 4X, 8X, 16X, 32X.** Both sources carry ancient DNA damage (U5a2a with the damage rates scaled by 0.75). For each coverage depth, the upper panel shows the phylogenetic placement posterior of both sources for the best of five chains ($k = 2$, 100,000 iterations each), colored by source (dot area: branch posterior probability; swelling: posterior density of the position along the branch; bold tips: highest-posterior branches), and the lower panel the posterior densities of the mixture proportions $\Theta$ with median and 95% credible interval. Chagyrskaya 8 is expected within the Neanderthal clade, on or near the Neanderthal_Lescottes branch (its closest relative in the reference); U5a2a, for which no U5 haplotype is present in the reference, is expected within the U clade. Modal branch of each source in the best chain ($p$: its posterior probability) and $\Theta$ median [95% CI]: (A,B) 0.5X: L6a ($p = 1.00$), $\Theta = 0.61$ [0.47, 0.74]; Neanderthal_Lescottes ($p = 1.00$), $\Theta = 0.39$ [0.26, 0.53]; (C,D) 1X: F2a1 ($p = 1.00$), $\Theta = 0.65$ [0.55, 0.74]; Neanderthal_Lescottes ($p = 1.00$), $\Theta = 0.35$ [0.26, 0.45]; (E,F) 2X: V2a1a ($p = 0.94$), $\Theta = 0.64$ [0.57, 0.71]; Neanderthal_Lescottes ($p = 0.99$), $\Theta = 0.36$ [0.29, 0.43]; (G,H) 4X: Node19 (stem of the U clade; $p = 1.00$), $\Theta = 0.63$ [0.58, 0.68]; Neanderthal_Lescottes ($p = 1.00$), $\Theta = 0.37$ [0.32, 0.42]; (I,J) 8X: Node19 (stem of the U clade; $p = 1.00$), $\Theta = 0.65$ [0.61, 0.68]; Neanderthal_Lescottes ($p = 1.00$), $\Theta = 0.35$ [0.32, 0.39]; (K,L) 16X: Node19 (stem of the U clade; $p = 0.99$), $\Theta = 0.66$ [0.63, 0.68]; Neanderthal_Lescottes ($p = 1.00$), $\Theta = 0.34$ [0.32, 0.37]; (M,N) 32X: U6a6a ($p = 0.88$), $\Theta = 0.67$ [0.65, 0.69]; Neanderthal_Lescottes ($p = 1.00$), $\Theta = 0.33$ [0.31, 0.35].

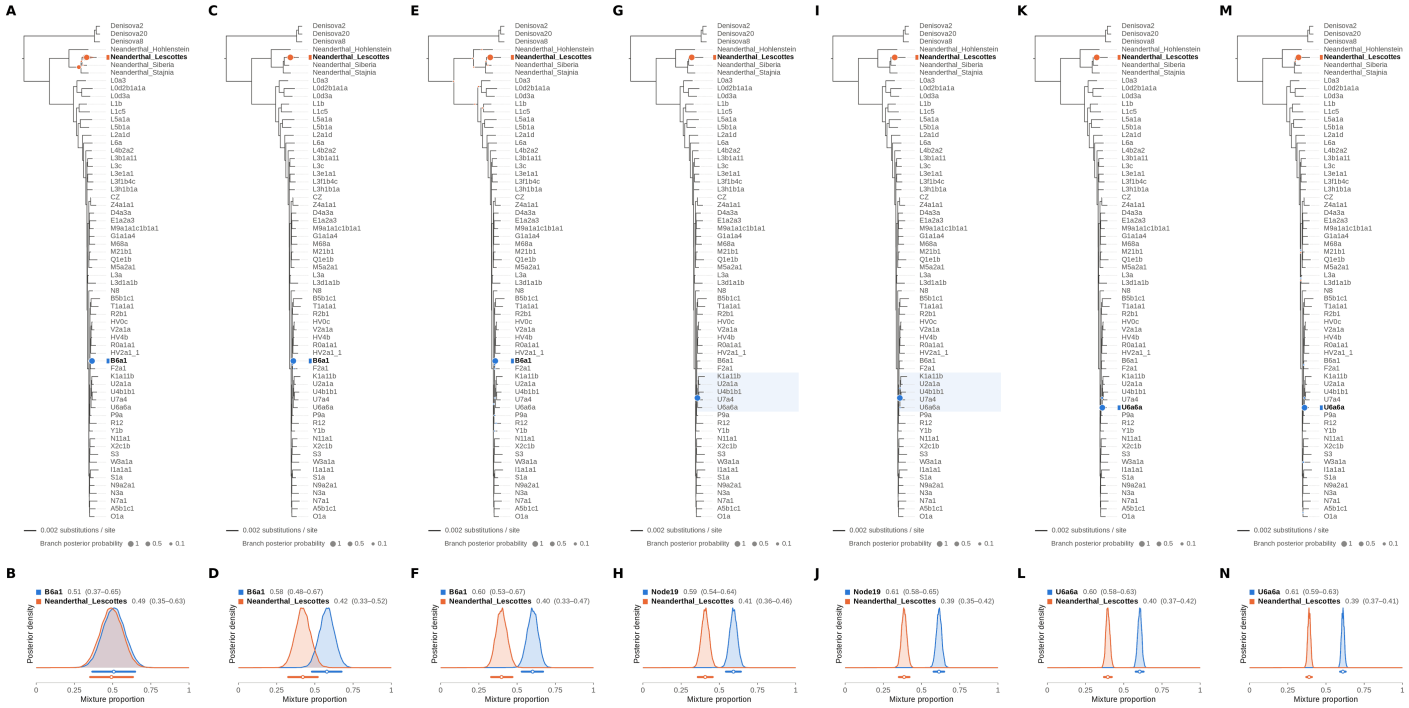


Figure S20: **Two-source simulated mixture of Chagyrskaya 8 Neanderthal (MK388903.1) (40% of reads) and modern human U5a2a (HM229344.1; 60%) at nominal mean coverage depths of 0.5X, 1X, 2X, 4X, 8X, 16X, 32X.** Both sources carry ancient DNA damage (U5a2a with the damage rates scaled by 0.75). For each coverage depth, the upper panel shows the phylogenetic placement posterior of both sources for the best of five chains ($k = 2$, 100,000 iterations each), colored by source (dot area: branch posterior probability; swelling: posterior density of the position along the branch; bold tips: highest-posterior branches), and the lower panel the posterior densities of the mixture proportions $\Theta$ with median and 95% credible interval. Chagyrskaya 8 is expected within the Neanderthal clade, on or near the Neanderthal_Lescottes branch (its closest relative in the reference); U5a2a, for which no U5 haplotype is present in the reference, is expected within the U clade. Modal branch of each source in the best chain ($p$: its posterior probability) and $\Theta$ median [95% CI]: (A,B) 0.5X: B6a1 ($p = 1.00$), $\Theta = 0.51$ [0.37, 0.65]; Neanderthal_Lescottes ($p = 0.78$), $\Theta = 0.49$ [0.35, 0.63]; (C,D) 1X: B6a1 ($p = 1.00$), $\Theta = 0.58$ [0.48, 0.67]; Neanderthal_Lescottes ($p = 1.00$), $\Theta = 0.42$ [0.33, 0.52]; (E,F) 2X: B6a1 ($p = 1.00$), $\Theta = 0.60$ [0.53, 0.67]; Neanderthal_Lescottes ($p = 1.00$), $\Theta = 0.40$ [0.33, 0.47]; (G,H) 4X: Node19 (stem of the U clade; $p = 1.00$), $\Theta = 0.59$ [0.54, 0.64]; Neanderthal_Lescottes ($p = 1.00$), $\Theta = 0.41$ [0.36, 0.46]; (I,J) 8X: Node19 (stem of the U clade; $p = 1.00$), $\Theta = 0.61$ [0.58, 0.65]; Neanderthal_Lescottes ($p = 1.00$), $\Theta = 0.39$ [0.35, 0.42]; (K,L) 16X: U6a6a ($p = 0.99$), $\Theta = 0.60$ [0.58, 0.63]; Neanderthal_Lescottes ($p = 1.00$), $\Theta = 0.40$ [0.37, 0.42]; (M,N) 32X: U6a6a ($p = 0.99$), $\Theta = 0.61$ [0.59, 0.63]; Neanderthal_Lescottes ($p = 1.00$), $\Theta = 0.39$ [0.37, 0.41].

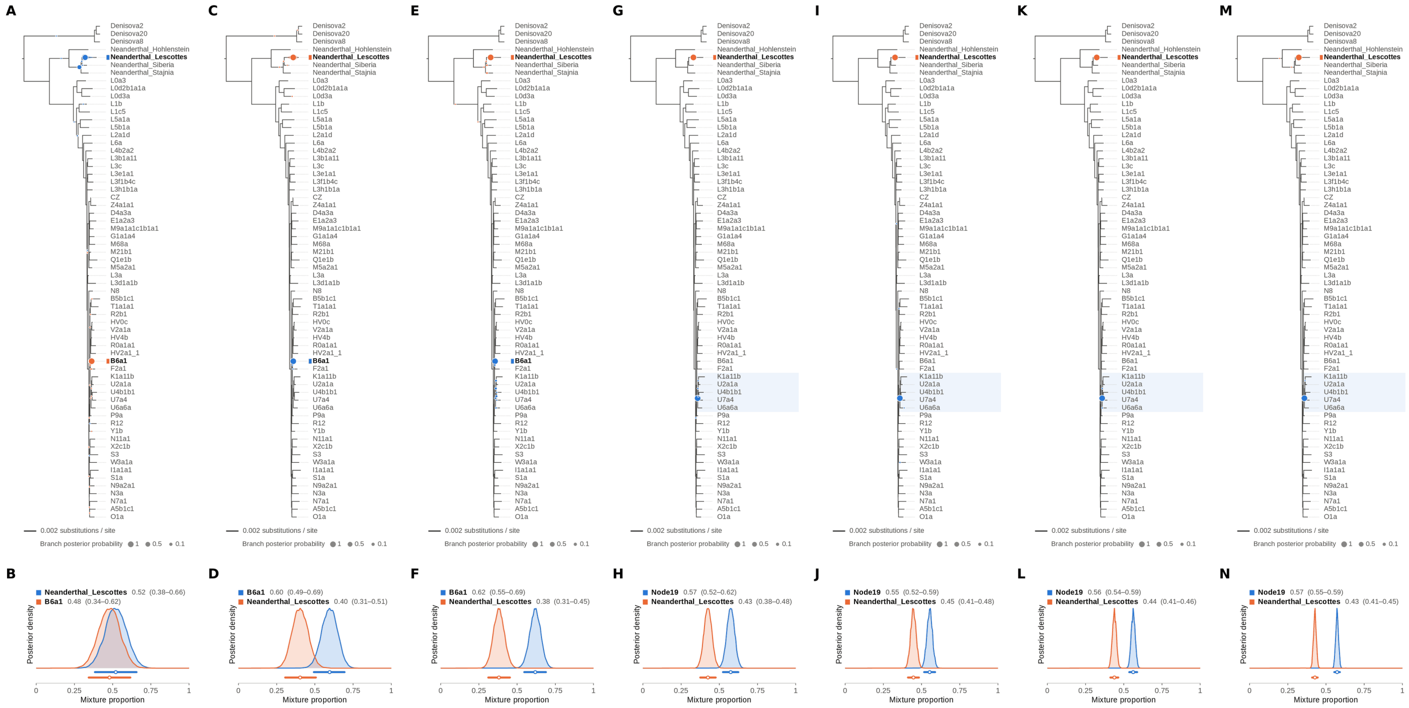


Figure S21: **Two-source simulated mixture of Chagyrskaya 8 Neanderthal (MK388903.1) (45% of reads) and modern human U5a2a (HM229344.1; 55%) at nominal mean coverage depths of 0.5X, 1X, 2X, 4X, 8X, 16X, 32X.** Both sources carry ancient DNA damage (U5a2a with the damage rates scaled by 0.75). For each coverage depth, the upper panel shows the phylogenetic placement posterior of both sources for the best of five chains ($k = 2$, 100,000 iterations each), colored by source (dot area: branch posterior probability; swelling: posterior density of the position along the branch; bold tips: highest-posterior branches), and the lower panel the posterior densities of the mixture proportions Θ with median and 95% credible interval. Chagyrskaya 8 is expected within the Neanderthal clade, on or near the Neanderthal_Lescottes branch (its closest relative in the reference); U5a2a, for which no U5 haplotype is present in the reference, is expected within the U clade. Modal branch of each source in the best chain ($p$: its posterior probability) and Θ median [95% CI]: (A,B) 0.5X: Neanderthal_Lescottes ($p = 0.73$), $\Theta = 0.52$ [0.38, 0.66]; B6a1 ($p = 0.99$), $\Theta = 0.48$ [0.34, 0.62]; (C,D) 1X: B6a1 ($p = 1.00$), $\Theta = 0.60$ [0.49, 0.69]; Neanderthal_Lescottes ($p = 1.00$), $\Theta = 0.40$ [0.31, 0.51]; (E,F) 2X: B6a1 ($p = 0.94$), $\Theta = 0.62$ [0.55, 0.69]; Neanderthal_Lescottes ($p = 0.99$), $\Theta = 0.38$ [0.31, 0.45]; (G,H) 4X: Node19 (stem of the U clade; $p = 1.00$), $\Theta = 0.57$ [0.52, 0.62]; Neanderthal_Lescottes ($p = 1.00$), $\Theta = 0.43$ [0.38, 0.48]; (I,J) 8X: Node19 (stem of the U clade; $p = 1.00$), $\Theta = 0.55$ [0.52, 0.59]; Neanderthal_Lescottes ($p = 1.00$), $\Theta = 0.45$ [0.41, 0.48]; (K,L) 16X: Node19 (stem of the U clade; $p = 1.00$), $\Theta = 0.56$ [0.54, 0.59]; Neanderthal_Lescottes ($p = 1.00$), $\Theta = 0.44$ [0.41, 0.46]; (M,N) 32X: Node19 (stem of the U clade; $p = 1.00$), $\Theta = 0.57$ [0.55, 0.59]; Neanderthal_Lescottes ($p = 1.00$), $\Theta = 0.43$ [0.41, 0.45].

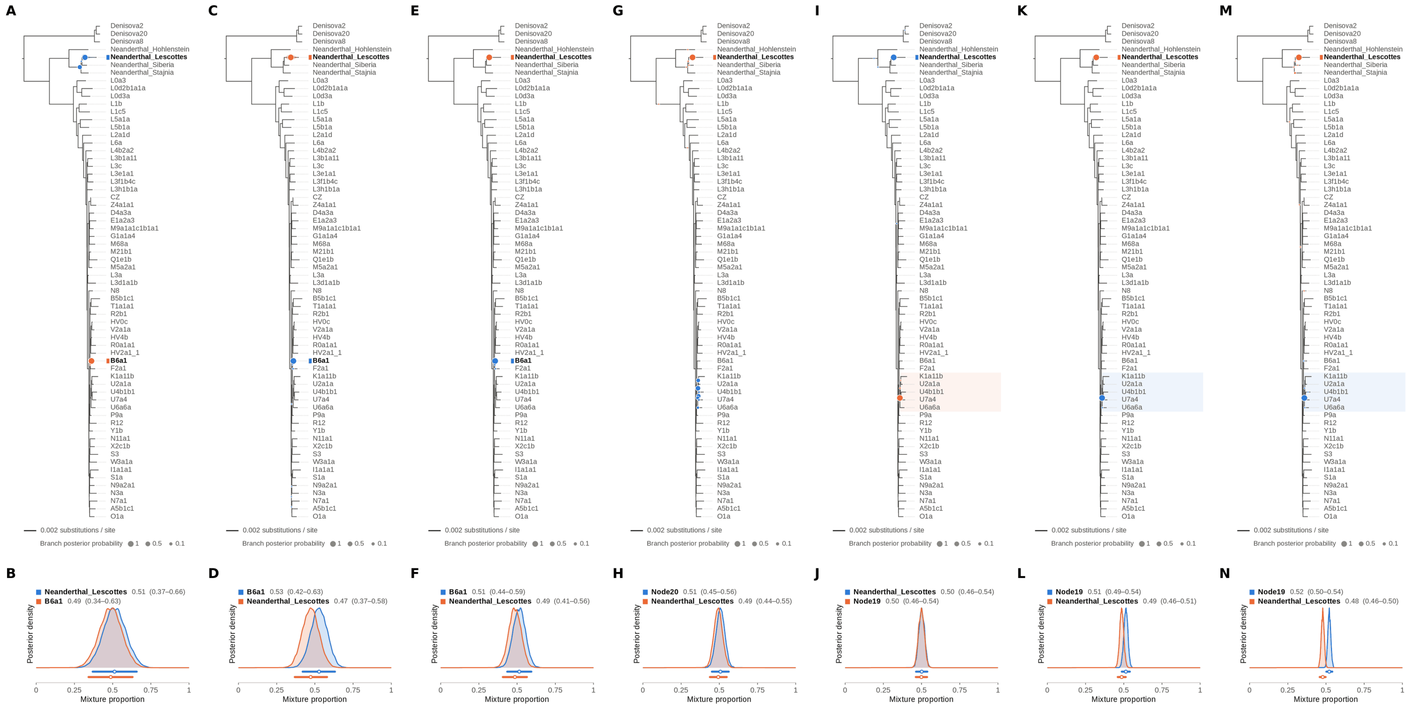


Figure S22: **Two-source simulated mixture of Chagyrskaya 8 Neanderthal (MK388903.1) (50% of reads) and modern human U5a2a (HM229344.1; 50%) at nominal mean coverage depths of 0.5X, 1X, 2X, 4X, 8X, 16X, 32X.** Both sources carry ancient DNA damage (U5a2a with the damage rates scaled by 0.75). For each coverage depth, the upper panel shows the phylogenetic placement posterior of both sources for the best of five chains ($k = 2$, 100,000 iterations each), colored by source (dot area: branch posterior probability; swelling: posterior density of the position along the branch; bold tips: highest-posterior branches), and the lower panel the posterior densities of the mixture proportions $\Theta$ with median and 95% credible interval. Chagyrskaya 8 is expected within the Neanderthal clade, on or near the Neanderthal_Lescottes branch (its closest relative in the reference); U5a2a, for which no U5 haplotype is present in the reference, is expected within the U clade. Modal branch of each source in the best chain ($p$: its posterior probability) and $\Theta$ median [95% CI]: (A,B) 0.5X: Neanderthal_Lescottes ($p = 0.70$), $\Theta = 0.51$ [0.37, 0.66]; B6a1 ($p = 1.00$), $\Theta = 0.49$ [0.34, 0.63]; (C,D) 1X: B6a1 ($p = 1.00$), $\Theta = 0.53$ [0.42, 0.63]; Neanderthal_Lescottes ($p = 1.00$), $\Theta = 0.47$ [0.37, 0.58]; (E,F) 2X: B6a1 ($p = 0.99$), $\Theta = 0.51$ [0.44, 0.59]; Neanderthal_Lescottes ($p = 1.00$), $\Theta = 0.49$ [0.41, 0.56]; (G,H) 4X: Node20 (internal branch within the U clade; $p = 0.43$), $\Theta = 0.51$ [0.45, 0.56]; Neanderthal_Lescottes ($p = 1.00$), $\Theta = 0.49$ [0.44, 0.55]; (I,J) 8X: Neanderthal_Lescottes ($p = 1.00$), $\Theta = 0.50$ [0.46, 0.54]; Node19 (stem of the U clade; $p = 1.00$), $\Theta = 0.50$ [0.46, 0.54]; (K,L) 16X: Node19 (stem of the U clade; $p = 1.00$), $\Theta = 0.51$ [0.49, 0.54]; Neanderthal_Lescottes ($p = 1.00$), $\Theta = 0.49$ [0.46, 0.51]; (M,N) 32X: Node19 (stem of the U clade; $p = 1.00$), $\Theta = 0.52$ [0.50, 0.54]; Neanderthal_Lescottes ($p = 0.99$), $\Theta = 0.48$ [0.46, 0.50].

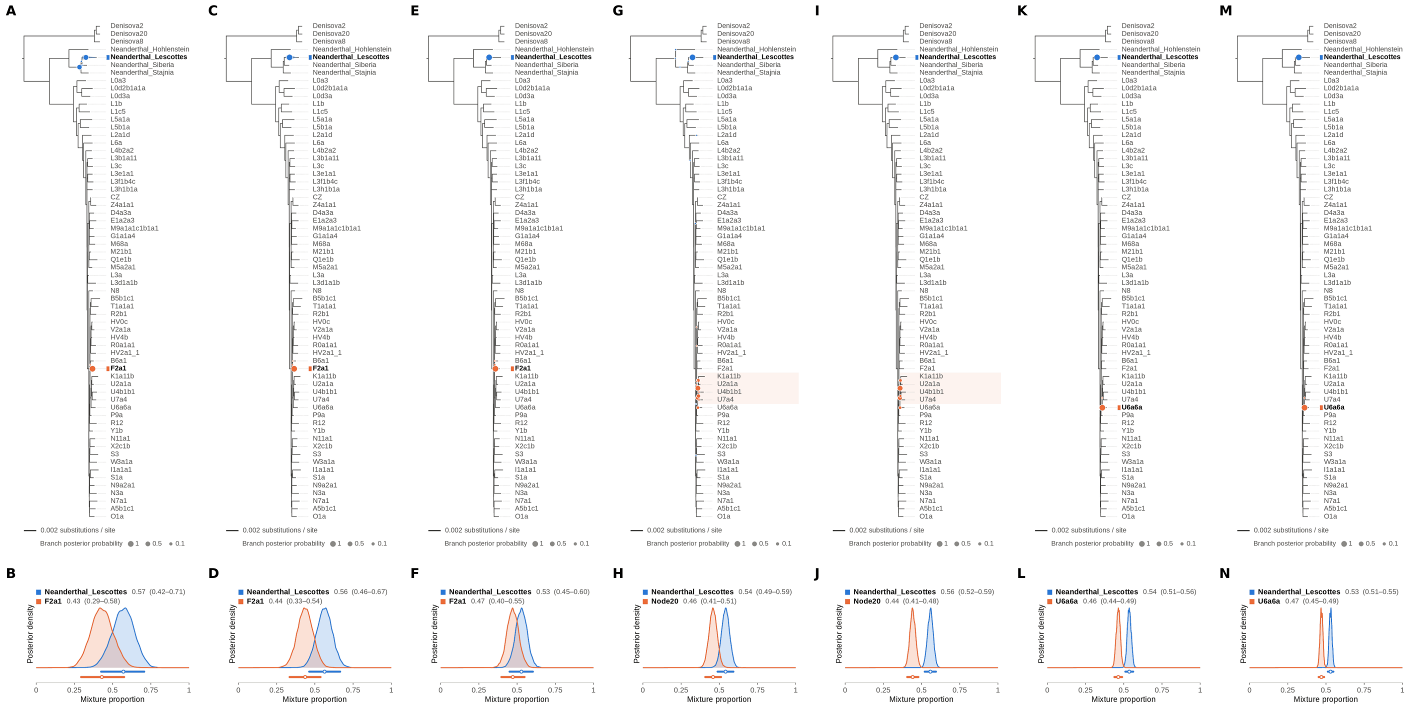

Figure S23: **Two-source simulated mixture of Chagyrskaya 8 Neanderthal (MK388903.1) (55% of reads) and modern human U5a2a (HM229344.1; 45%) at nominal mean coverage depths of 0.5X, 1X, 2X, 4X, 8X, 16X, 32X.** Both sources carry ancient DNA damage (U5a2a with the damage rates scaled by 0.75). For each coverage depth, the upper panel shows the phylogenetic placement posterior of both sources for the best of five chains ($k = 2$, 100,000 iterations each), colored by source (dot area: branch posterior probability; swelling: posterior density of the position along the branch; bold tips: highest-posterior branches), and the lower panel the posterior densities of the mixture proportions $\Theta$ with median and 95% credible interval. Chagyrskaya 8 is expected within the Neanderthal clade, on or near the Neanderthal_Lescottes branch (its closest relative in the reference); U5a2a, for which no U5 haplotype is present in the reference, is expected within the U clade. Modal branch of each source in the best chain ($p$: its posterior probability) and $\Theta$ median [95% CI]: (A,B) 0.5X: Neanderthal_Lescottes ($p = 0.59$), $\Theta = 0.57$ [0.42, 0.71]; F2a1 ($p = 1.00$), $\Theta = 0.43$ [0.29, 0.58]; (C,D) 1X: Neanderthal_Lescottes ($p = 1.00$), $\Theta = 0.56$ [0.46, 0.67]; F2a1 ($p = 1.00$), $\Theta = 0.44$ [0.33, 0.54]; (E,F) 2X: Neanderthal_Lescottes ($p = 1.00$), $\Theta = 0.53$ [0.45, 0.60]; F2a1 ($p = 0.99$), $\Theta = 0.47$ [0.40, 0.55]; (G,H) 4X: Neanderthal_Lescottes ($p = 1.00$), $\Theta = 0.54$ [0.49, 0.59]; Node20 (internal branch within the U clade; $p = 0.52$), $\Theta = 0.46$ [0.41, 0.51]; (I,J) 8X: Neanderthal_Lescottes ($p = 1.00$), $\Theta = 0.56$ [0.52, 0.59]; Node20 (internal branch within the U clade; $p = 0.57$), $\Theta = 0.44$ [0.41, 0.48]; (K,L) 16X: Neanderthal_Lescottes ($p = 1.00$), $\Theta = 0.54$ [0.51, 0.56]; U6a6a ($p = 1.00$), $\Theta = 0.46$ [0.44, 0.49]; (M,N) 32X: Neanderthal_Lescottes ($p = 1.00$), $\Theta = 0.53$ [0.51, 0.55]; U6a6a ($p = 1.00$), $\Theta = 0.47$ [0.45, 0.49].

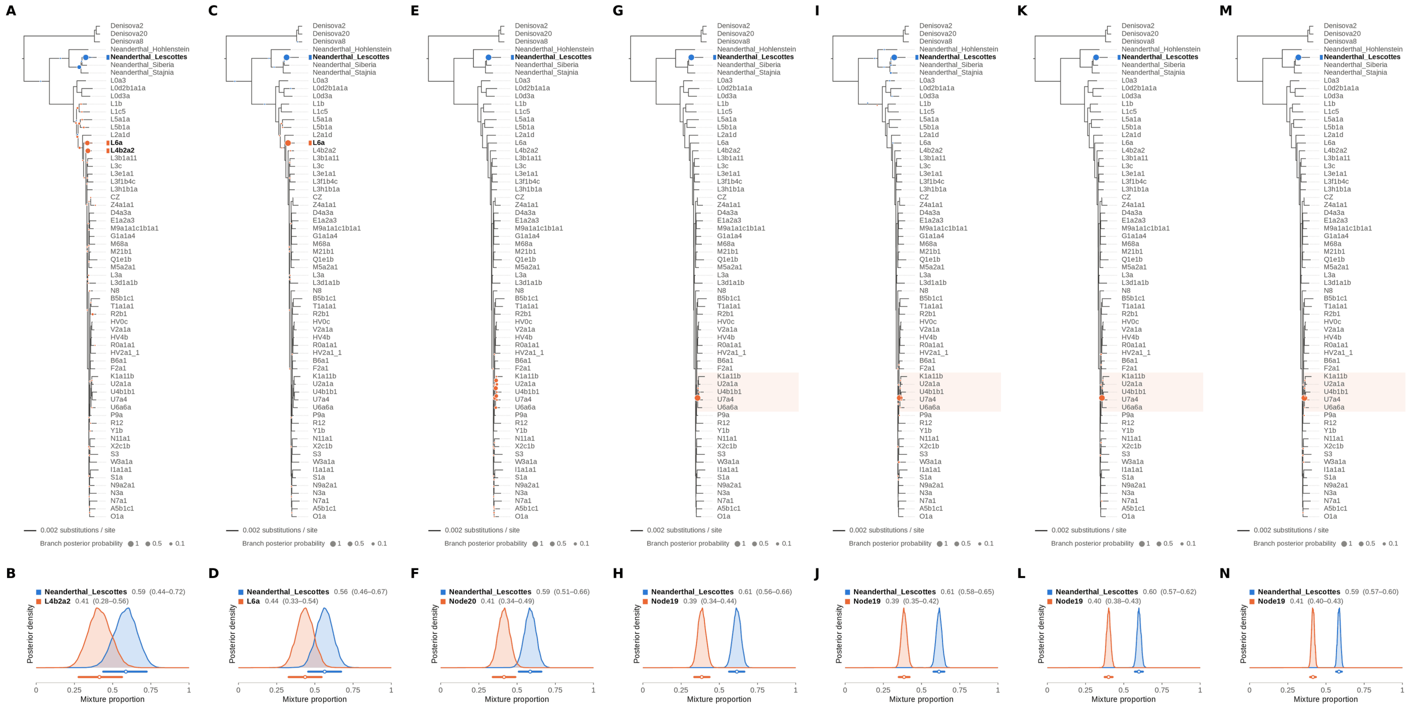

Figure S24: **Two-source simulated mixture of Chagyrskaya 8 Neanderthal (MK388903.1) (60% of reads) and modern human U5a2a (HM229344.1; 40%) at nominal mean coverage depths of 0.5X, 1X, 2X, 4X, 8X, 16X, 32X.** Both sources carry ancient DNA damage (U5a2a with the damage rates scaled by 0.75). For each coverage depth, the upper panel shows the phylogenetic placement posterior of both sources for the best of five chains ($k = 2$, 100,000 iterations each), colored by source (dot area: branch posterior probability; swelling: posterior density of the position along the branch; bold tips: highest-posterior branches), and the lower panel the posterior densities of the mixture proportions Θ with median and 95% credible interval. Chagyrskaya 8 is expected within the Neanderthal clade, on or near the Neanderthal_Lescottes branch (its closest relative in the reference); U5a2a, for which no U5 haplotype is present in the reference, is expected within the U clade. Modal branch of each source in the best chain ($p$: its posterior probability) and Θ median [95% CI]: (A,B) 0.5X: Neanderthal_Lescottes ($p = 0.80$), $\Theta = 0.59$ [0.44, 0.72]; L4b2a2 ($p = 0.49$), $\Theta = 0.41$ [0.28, 0.56]; (C,D) 1X: Neanderthal_Lescottes ($p = 1.00$), $\Theta = 0.56$ [0.46, 0.67]; L6a ($p = 0.99$), $\Theta = 0.44$ [0.33, 0.54]; (E,F) 2X: Neanderthal_Lescottes ($p = 1.00$), $\Theta = 0.59$ [0.51, 0.66]; Node20 (internal branch within the U clade; $p = 0.30$), $\Theta = 0.41$ [0.34, 0.49]; (G,H) 4X: Neanderthal_Lescottes ($p = 1.00$), $\Theta = 0.61$ [0.56, 0.66]; Node19 (stem of the U clade; $p = 1.00$), $\Theta = 0.39$ [0.34, 0.44]; (I,J) 8X: Neanderthal_Lescottes ($p = 0.99$), $\Theta = 0.61$ [0.58, 0.65]; Node19 (stem of the U clade; $p = 0.99$), $\Theta = 0.39$ [0.35, 0.42]; (K,L) 16X: Neanderthal_Lescottes ($p = 1.00$), $\Theta = 0.60$ [0.57, 0.62]; Node19 (stem of the U clade; $p = 1.00$), $\Theta = 0.40$ [0.38, 0.43]; (M,N) 32X: Neanderthal_Lescottes ($p = 1.00$), $\Theta = 0.59$ [0.57, 0.60]; Node19 (stem of the U clade; $p = 0.99$), $\Theta = 0.41$ [0.40, 0.43].

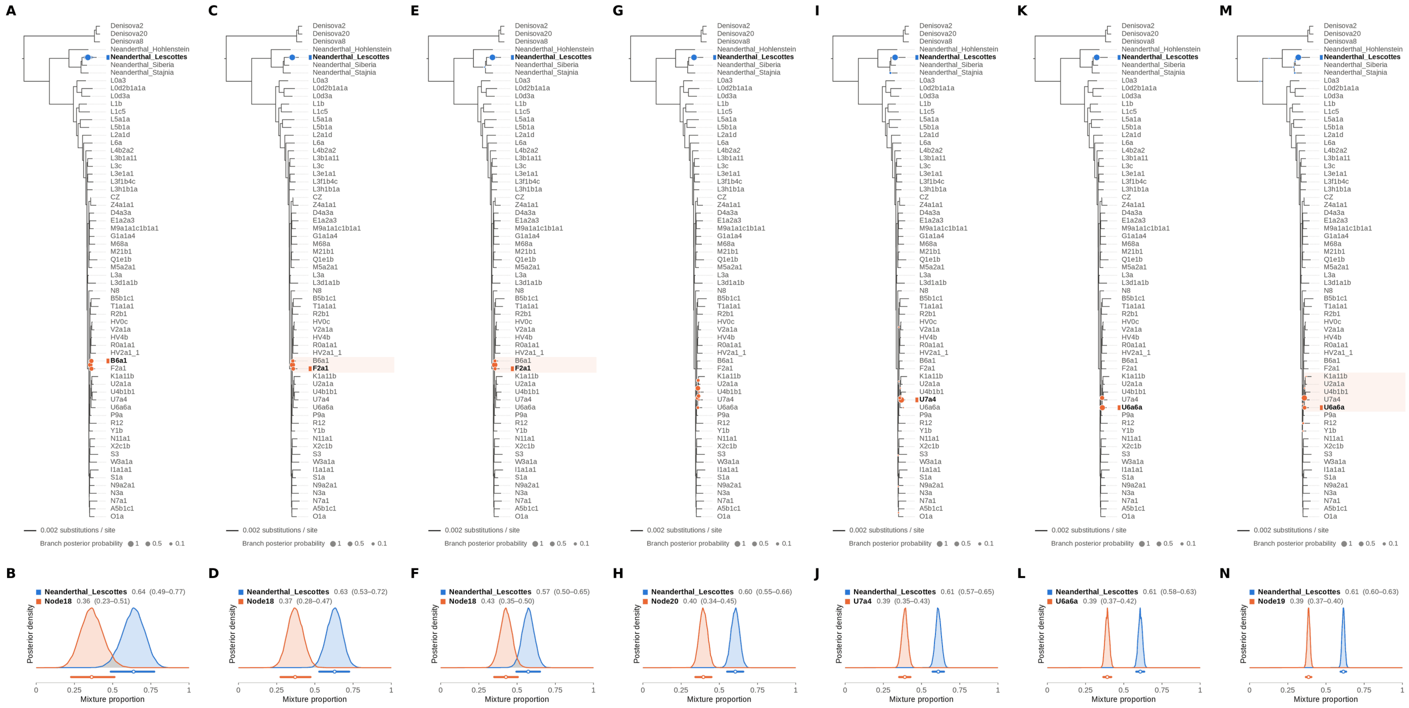


Figure S25: **Two-source simulated mixture of Chagyrskaya 8 Neanderthal (MK388903.1) (65% of reads) and modern human U5a2a (HM229344.1; 35%) at nominal mean coverage depths of 0.5X, 1X, 2X, 4X, 8X, 16X, 32X.** Both sources carry ancient DNA damage (U5a2a with the damage rates scaled by 0.75). For each coverage depth, the upper panel shows the phylogenetic placement posterior of both sources for the best of five chains ($k = 2$, 100,000 iterations each), colored by source (dot area: branch posterior probability; swelling: posterior density of the position along the branch; bold tips: highest-posterior branches), and the lower panel the posterior densities of the mixture proportions $\Theta$ with median and 95% credible interval. Chagyrskaya 8 is expected within the Neanderthal clade, on or near the Neanderthal_Lescottes branch (its closest relative in the reference); U5a2a, for which no U5 haplotype is present in the reference, is expected within the U clade. Modal branch of each source in the best chain ($p$: its posterior probability) and $\Theta$ median [95% CI]: (A,B) 0.5X: Neanderthal_Lescottes ($p = 1.00$), $\Theta = 0.64$ [0.49, 0.77]; Node18 (internal branch ancestral to B6a1 and F2a1; $p = 0.40$), $\Theta = 0.36$ [0.23, 0.51]; (C,D) 1X: Neanderthal_Lescottes ($p = 1.00$), $\Theta = 0.63$ [0.53, 0.72]; Node18 (internal branch ancestral to B6a1 and F2a1; $p = 0.66$), $\Theta = 0.37$ [0.28, 0.47]; (E,F) 2X: Neanderthal_Lescottes ($p = 1.00$), $\Theta = 0.57$ [0.50, 0.65]; Node18 (internal branch ancestral to B6a1 and F2a1; $p = 0.82$), $\Theta = 0.43$ [0.35, 0.50]; (G,H) 4X: Neanderthal_Lescottes ($p = 1.00$), $\Theta = 0.60$ [0.55, 0.66]; Node20 (internal branch within the U clade; $p = 0.47$), $\Theta = 0.40$ [0.34, 0.45]; (I,J) 8X: Neanderthal_Lescottes ($p = 0.99$), $\Theta = 0.61$ [0.57, 0.65]; U7a4 ($p = 0.92$), $\Theta = 0.39$ [0.35, 0.43]; (K,L) 16X: Neanderthal_Lescottes ($p = 1.00$), $\Theta = 0.61$ [0.58, 0.63]; U6a6a ($p = 0.69$), $\Theta = 0.39$ [0.37, 0.42]; (M,N) 32X: Neanderthal_Lescottes ($p = 0.99$), $\Theta = 0.61$ [0.60, 0.63]; Node19 (stem of the U clade; $p = 0.77$), $\Theta = 0.39$ [0.37, 0.40].

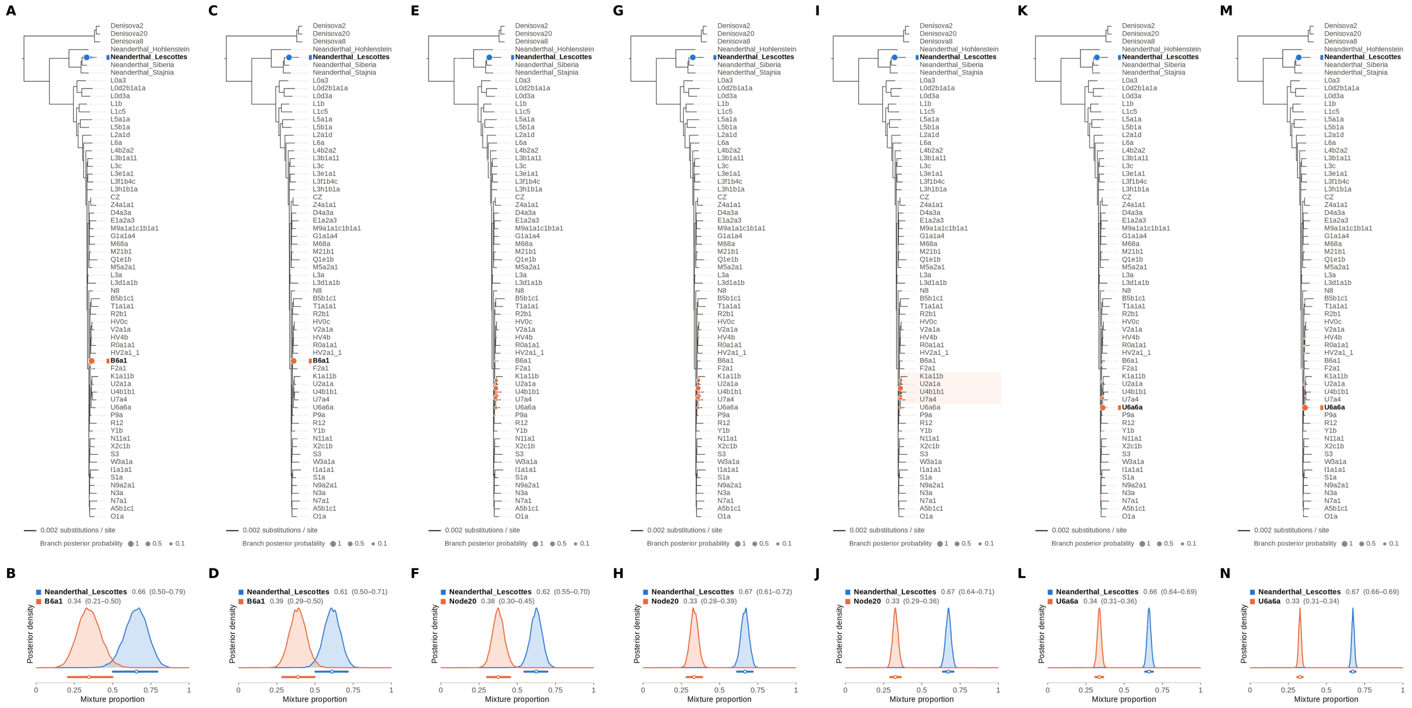


Figure S26: **Two-source simulated mixture of Chagyrskaya 8 Neanderthal (MK388903.1) (70% of reads) and modern human U5a2a (HM229344.1; 30%) at nominal mean coverage depths of 0.5X, 1X, 2X, 4X, 8X, 16X, 32X.** Both sources carry ancient DNA damage (U5a2a with the damage rates scaled by 0.75). For each coverage depth, the upper panel shows the phylogenetic placement posterior of both sources for the best of five chains ($k = 2$, 100,000 iterations each), colored by source (dot area: branch posterior probability; swelling: posterior density of the position along the branch; bold tips: highest-posterior branches), and the lower panel the posterior densities of the mixture proportions $\Theta$ with median and 95% credible interval. Chagyrskaya 8 is expected within the Neanderthal clade, on or near the Neanderthal_Lescottes branch (its closest relative in the reference); U5a2a, for which no U5 haplotype is present in the reference, is expected within the U clade. Modal branch of each source in the best chain ($p$: its posterior probability) and $\Theta$ median [95% CI]: (A,B) 0.5X: Neanderthal_Lescottes ($p = 1.00$), $\Theta = 0.66$ [0.50, 0.79]; B6a1 ($p = 1.00$), $\Theta = 0.34$ [0.21, 0.50]; (C,D) 1X: Neanderthal_Lescottes ($p = 1.00$), $\Theta = 0.61$ [0.50, 0.71]; B6a1 ($p = 1.00$), $\Theta = 0.39$ [0.29, 0.50]; (E,F) 2X: Neanderthal_Lescottes ($p = 1.00$), $\Theta = 0.62$ [0.55, 0.70]; Node20 (internal branch within the U clade; $p = 0.37$), $\Theta = 0.38$ [0.30, 0.45]; (G,H) 4X: Neanderthal_Lescottes ($p = 1.00$), $\Theta = 0.67$ [0.61, 0.72]; Node20 (internal branch within the U clade; $p = 0.42$), $\Theta = 0.33$ [0.28, 0.39]; (I,J) 8X: Neanderthal_Lescottes ($p = 1.00$), $\Theta = 0.67$ [0.64, 0.71]; Node20 (internal branch within the U clade; $p = 0.59$), $\Theta = 0.33$ [0.29, 0.36]; (K,L) 16X: Neanderthal_Lescottes ($p = 1.00$), $\Theta = 0.66$ [0.64, 0.69]; U6a6a ($p = 0.89$), $\Theta = 0.34$ [0.31, 0.36]; (M,N) 32X: Neanderthal_Lescottes ($p = 1.00$), $\Theta = 0.67$ [0.66, 0.69]; U6a6a ($p = 1.00$), $\Theta = 0.33$ [0.31, 0.34].

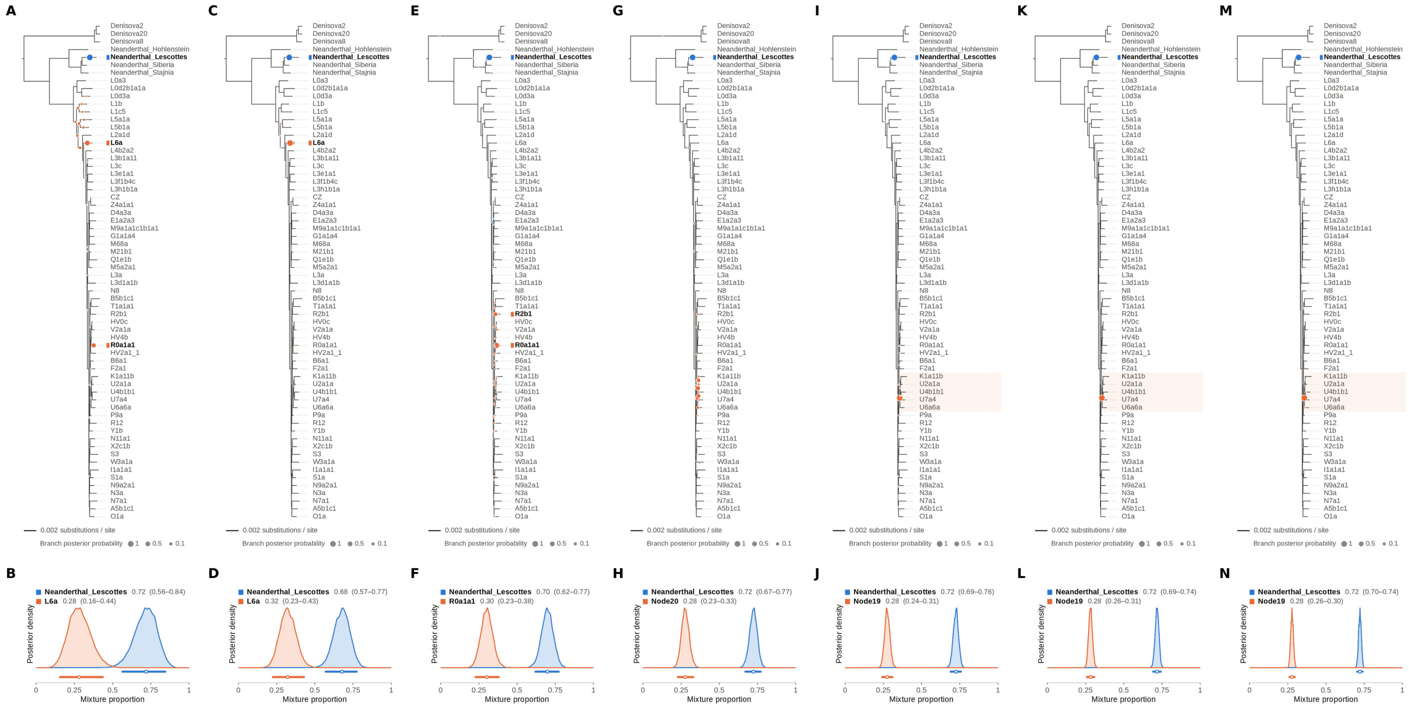


Figure S27: **Two-source simulated mixture of Chagyrskaya 8 Neanderthal (MK388903.1) (75% of reads) and modern human U5a2a (HM229344.1; 25%) at nominal mean coverage depths of 0.5X, 1X, 2X, 4X, 8X, 16X, 32X.** Both sources carry ancient DNA damage (U5a2a with the damage rates scaled by 0.75). For each coverage depth, the upper panel shows the phylogenetic placement posterior of both sources for the best of five chains ($k = 2$, 100,000 iterations each), colored by source (dot area: branch posterior probability; swelling: posterior density of the position along the branch; bold tips: highest-posterior branches), and the lower panel the posterior densities of the mixture proportions $\Theta$ with median and 95% credible interval. Chagyrskaya 8 is expected within the Neanderthal clade, on or near the Neanderthal_Lescottes branch (its closest relative in the reference); U5a2a, for which no U5 haplotype is present in the reference, is expected within the U clade. Modal branch of each source in the best chain ($p$: its posterior probability) and $\Theta$ median [95% CI]: (A,B) 0.5X: Neanderthal_Lescottes ($p = 1.00$), $\Theta = 0.72$ [0.56, 0.84]; L6a ($p = 0.47$), $\Theta = 0.28$ [0.16, 0.44]; (C,D) 1X: Neanderthal_Lescottes ($p = 1.00$), $\Theta = 0.68$ [0.57, 0.77]; L6a ($p = 1.00$), $\Theta = 0.32$ [0.23, 0.43]; (E,F) 2X: Neanderthal_Lescottes ($p = 1.00$), $\Theta = 0.70$ [0.62, 0.77]; R0a1a1 ($p = 0.73$), $\Theta = 0.30$ [0.23, 0.38]; (G,H) 4X: Neanderthal_Lescottes ($p = 0.99$), $\Theta = 0.72$ [0.67, 0.77]; Node20 (internal branch within the U clade; $p = 0.30$), $\Theta = 0.28$ [0.23, 0.33]; (I,J) 8X: Neanderthal_Lescottes ($p = 1.00$), $\Theta = 0.72$ [0.69, 0.76]; Node19 (stem of the U clade; $p = 1.00$), $\Theta = 0.28$ [0.24, 0.31]; (K,L) 16X: Neanderthal_Lescottes ($p = 1.00$), $\Theta = 0.72$ [0.69, 0.74]; Node19 (stem of the U clade; $p = 1.00$), $\Theta = 0.28$ [0.26, 0.31]; (M,N) 32X: Neanderthal_Lescottes ($p = 1.00$), $\Theta = 0.72$ [0.70, 0.74]; Node19 (stem of the U clade; $p = 1.00$), $\Theta = 0.28$ [0.26, 0.30].

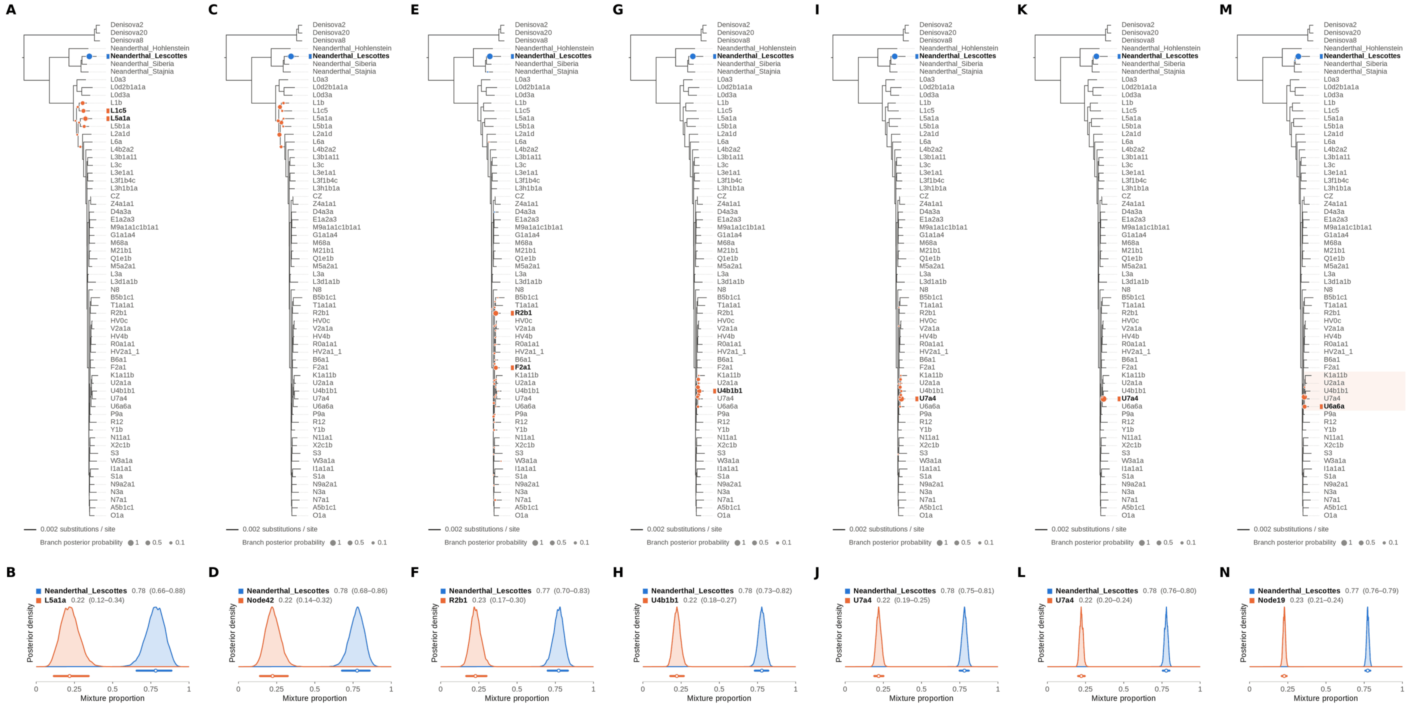

Figure S28: **Two-source simulated mixture of Chagyrskaya 8 Neanderthal (MK388903.1) (80% of reads) and modern human U5a2a (HM229344.1; 20%) at nominal mean coverage depths of 0.5X, 1X, 2X, 4X, 8X, 16X, 32X.** Both sources carry ancient DNA damage (U5a2a with the damage rates scaled by 0.75). For each coverage depth, the upper panel shows the phylogenetic placement posterior of both sources for the best of five chains ($k = 2$, 100,000 iterations each), colored by source (dot area: branch posterior probability; swelling: posterior density of the position along the branch; bold tips: highest-posterior branches), and the lower panel the posterior densities of the mixture proportions $\Theta$ with median and 95% credible interval. Chagyrskaya 8 is expected within the Neanderthal clade, on or near the Neanderthal_Lescottes branch (its closest relative in the reference); U5a2a, for which no U5 haplotype is present in the reference, is expected within the U clade. Modal branch of each source in the best chain ($p$: its posterior probability) and $\Theta$ median [95% CI]: (A,B) 0.5X: Neanderthal_Lescottes ($p = 1.00$), $\Theta = 0.78$ [0.66, 0.88]; L5a1a ($p = 0.36$), $\Theta = 0.22$ [0.12, 0.34]; (C,D) 1X: Neanderthal_Lescottes ($p = 1.00$), $\Theta = 0.78$ [0.68, 0.86]; Node42 (deep internal branch; $p = 0.29$), $\Theta = 0.22$ [0.14, 0.32]; (E,F) 2X: Neanderthal_Lescottes ($p = 1.00$), $\Theta = 0.77$ [0.70, 0.83]; R2b1 ($p = 0.59$), $\Theta = 0.23$ [0.17, 0.30]; (G,H) 4X: Neanderthal_Lescottes ($p = 1.00$), $\Theta = 0.78$ [0.73, 0.82]; U4b1b1 ($p = 0.34$), $\Theta = 0.22$ [0.18, 0.27]; (I,J) 8X: Neanderthal_Lescottes ($p = 1.00$), $\Theta = 0.78$ [0.75, 0.81]; U7a4 ($p = 0.71$), $\Theta = 0.22$ [0.19, 0.25]; (K,L) 16X: Neanderthal_Lescottes ($p = 1.00$), $\Theta = 0.78$ [0.76, 0.80]; U7a4 ($p = 1.00$), $\Theta = 0.22$ [0.20, 0.24]; (M,N) 32X: Neanderthal_Lescottes ($p = 1.00$), $\Theta = 0.77$ [0.76, 0.79]; Node19 (stem of the U clade; $p = 0.72$), $\Theta = 0.23$ [0.21, 0.24].

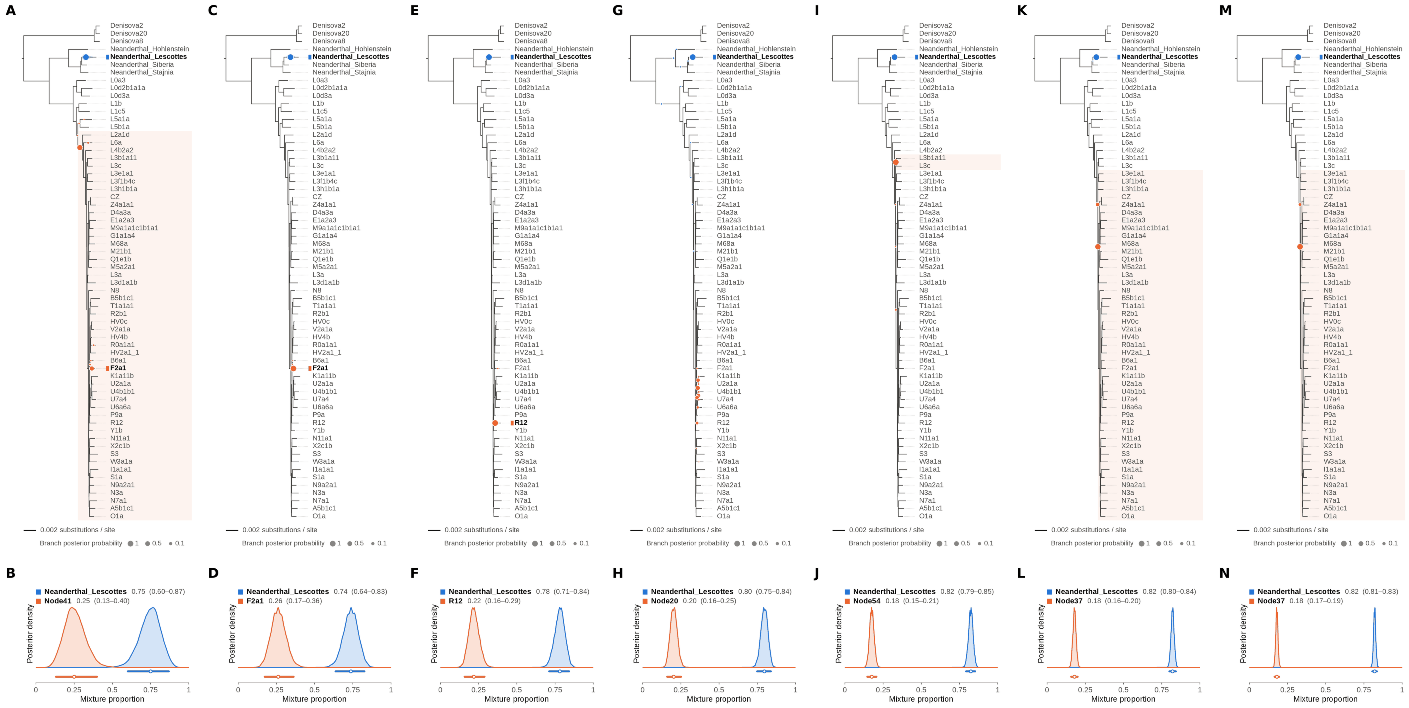


Figure S29: **Two-source simulated mixture of Chagyrskaya 8 Neanderthal (MK388903.1) (85% of reads) and modern human U5a2a (HM229344.1; 15%) at nominal mean coverage depths of 0.5X, 1X, 2X, 4X, 8X, 16X, 32X.** Both sources carry ancient DNA damage (U5a2a with the damage rates scaled by 0.75). For each coverage depth, the upper panel shows the phylogenetic placement posterior of both sources for the best of five chains ($k$ = 2, 100,000 iterations each), colored by source (dot area: branch posterior probability; swelling: posterior density of the position along the branch; bold tips: highest-posterior branches), and the lower panel the posterior densities of the mixture proportions $\Theta$ with median and 95% credible interval. Chagyrskaya 8 is expected within the Neanderthal clade, on or near the Neanderthal_Lescottes branch (its closest relative in the reference); U5a2a, for which no U5 haplotype is present in the reference, is expected within the U clade. Modal branch of each source in the best chain ($p$: its posterior probability) and $\Theta$ median [95% CI]: (A,B) 0.5X: Neanderthal_Lescottes ($p = 1.00$), $\Theta = 0.75$ [0.60, 0.87]; Node41 (deep internal branch; $p = 0.69$), $\Theta = 0.25$ [0.13, 0.40]; (C,D) 1X: Neanderthal_Lescottes ($p = 1.00$), $\Theta = 0.74$ [0.64, 0.83]; F2a1 ($p = 1.00$), $\Theta = 0.26$ [0.17, 0.36]; (E,F) 2X: Neanderthal_Lescottes ($p = 1.00$), $\Theta = 0.78$ [0.71, 0.84]; R12 ($p = 1.00$), $\Theta = 0.22$ [0.16, 0.29]; (G,H) 4X: Neanderthal_Lescottes ($p = 0.99$), $\Theta = 0.80$ [0.75, 0.84]; Node20 (internal branch within the U clade; $p = 0.31$), $\Theta = 0.20$ [0.16, 0.25]; (I,J) 8X: Neanderthal_Lescottes ($p = 1.00$), $\Theta = 0.82$ [0.79, 0.85]; Node54 (internal branch ancestral to L3b1a11 and L3c; $p = 0.99$), $\Theta = 0.18$ [0.15, 0.21]; (K,L) 16X: Neanderthal_Lescottes ($p = 1.00$), $\Theta = 0.82$ [0.80, 0.84]; Node37 (deep internal branch; $p = 0.83$), $\Theta = 0.18$ [0.16, 0.20]; (M,N) 32X: Neanderthal_Lescottes ($p = 1.00$), $\Theta = 0.82$ [0.81, 0.83]; Node37 (deep internal branch; $p = 0.94$), $\Theta = 0.18$ [0.17, 0.19].

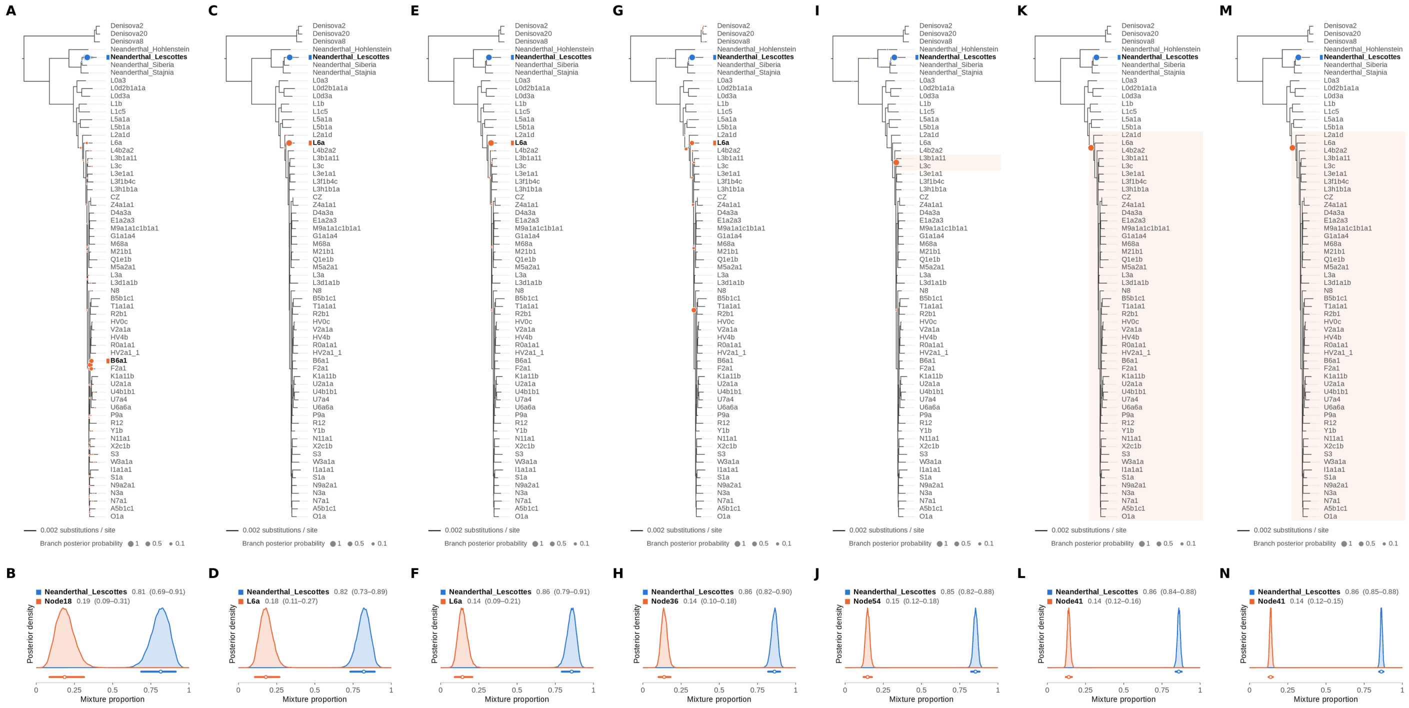

Figure S30: **Two-source simulated mixture of Chagyrskaya 8 Neanderthal (MK388903.1) (90% of reads) and modern human U5a2a (HM229344.1; 10%) at nominal mean coverage depths of 0.5X, 1X, 2X, 4X, 8X, 16X, 32X.** Both sources carry ancient DNA damage (U5a2a with the damage rates scaled by 0.75). For each coverage depth, the upper panel shows the phylogenetic placement posterior of both sources for the best of five chains ($k = 2$, 100,000 iterations each), colored by source (dot area: branch posterior probability; swelling: posterior density of the position along the branch; bold tips: highest-posterior branches), and the lower panel the posterior densities of the mixture proportions $\Theta$ with median and 95% credible interval. Chagyrskaya 8 is expected within the Neanderthal clade, on or near the Neanderthal_Lescottes branch (its closest relative in the reference); U5a2a, for which no U5 haplotype is present in the reference, is expected within the U clade. Modal branch of each source in the best chain ($p$: its posterior probability) and $\Theta$ median [95% CI]: (A,B) 0.5X: Neanderthal_Lescottes ($p = 1.00$), $\Theta = 0.81$ [0.69, 0.91]; Node18 (internal branch ancestral to B6a1 and F2a1; $p = 0.48$), $\Theta = 0.19$ [0.09, 0.31]; (C,D) 1X: Neanderthal_Lescottes ($p = 1.00$), $\Theta = 0.82$ [0.73, 0.89]; L6a ($p = 1.00$), $\Theta = 0.18$ [0.11, 0.27]; (E,F) 2X: Neanderthal_Lescottes ($p = 1.00$), $\Theta = 0.86$ [0.79, 0.91]; L6a ($p = 0.99$), $\Theta = 0.14$ [0.09, 0.21]; (G,H) 4X: Neanderthal_Lescottes ($p = 1.00$), $\Theta = 0.86$ [0.82, 0.90]; Node36 (deep internal branch; $p = 0.49$), $\Theta = 0.14$ [0.10, 0.18]; (I,J) 8X: Neanderthal_Lescottes ($p = 1.00$), $\Theta = 0.85$ [0.82, 0.88]; Node54 (internal branch ancestral to L3b1a11 and L3c; $p = 1.00$), $\Theta = 0.15$ [0.12, 0.18]; (K,L) 16X: Neanderthal_Lescottes ($p = 1.00$), $\Theta = 0.86$ [0.84, 0.88]; Node41 (deep internal branch; $p = 1.00$), $\Theta = 0.14$ [0.12, 0.16]; (M,N) 32X: Neanderthal_Lescottes ($p = 1.00$), $\Theta = 0.86$ [0.85, 0.88]; Node41 (deep internal branch; $p = 1.00$), $\Theta = 0.14$ [0.12, 0.15].

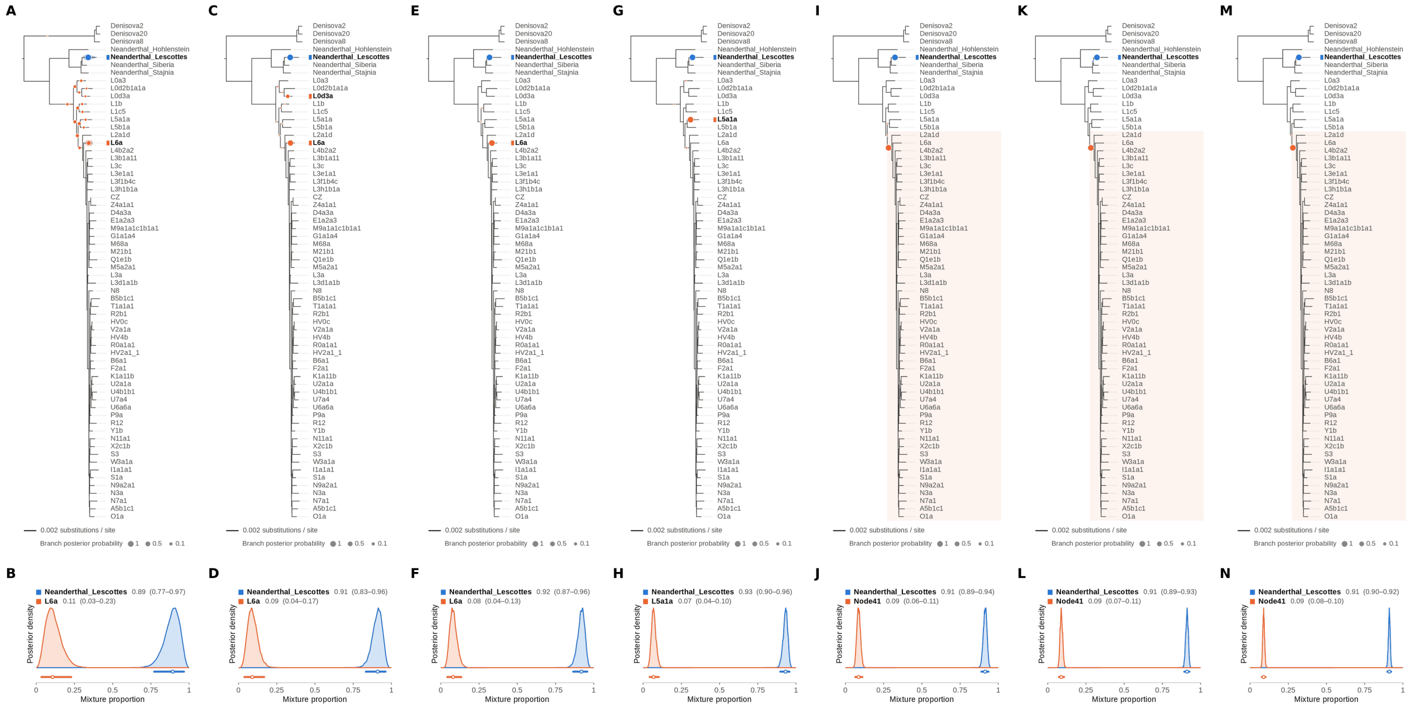


Figure S31: **Two-source simulated mixture of Chagyrskaya 8 Neanderthal (MK388903.1) (95% of reads) and modern human U5a2a (HM229344.1; 5%) at nominal mean coverage depths of 0.5X, 1X, 2X, 4X, 8X, 16X, 32X.** Both sources carry ancient DNA damage (U5a2a with the damage rates scaled by 0.75). For each coverage depth, the upper panel shows the phylogenetic placement posterior of both sources for the best of five chains ($k = 2$, 100,000 iterations each), colored by source (dot area: branch posterior probability; swelling: posterior density of the position along the branch; bold tips: highest-posterior branches), and the lower panel the posterior densities of the mixture proportions $\Theta$ with median and 95% credible interval. Chagyrskaya 8 is expected within the Neanderthal clade, on or near the Neanderthal_Lescottes branch (its closest relative in the reference); U5a2a, for which no U5 haplotype is present in the reference, is expected within the U clade. Modal branch of each source in the best chain ($p$: its posterior probability) and $\Theta$ median [95% CI]: (A,B) 0.5X: Neanderthal_Lescottes ($p = 1.00$), $\Theta = 0.89$ [0.77, 0.97]; L6a ($p = 0.25$), $\Theta = 0.11$ [0.03, 0.23]; (C,D) 1X: Neanderthal_Lescottes ($p = 1.00$), $\Theta = 0.91$ [0.83, 0.96]; L6a ($p = 0.83$), $\Theta = 0.09$ [0.04, 0.17]; (E,F) 2X: Neanderthal_Lescottes ($p = 1.00$), $\Theta = 0.92$ [0.87, 0.96]; L6a ($p = 1.00$), $\Theta = 0.08$ [0.04, 0.13]; (G,H) 4X: Neanderthal_Lescottes ($p = 1.00$), $\Theta = 0.93$ [0.90, 0.96]; L5a1a ($p = 1.00$), $\Theta = 0.07$ [0.04, 0.10]; (I,J) 8X: Neanderthal_Lescottes ($p = 1.00$), $\Theta = 0.91$ [0.89, 0.94]; Node41 (deep internal branch; $p = 1.00$), $\Theta = 0.09$ [0.06, 0.11]; (K,L) 16X: Neanderthal_Lescottes ($p = 1.00$), $\Theta = 0.91$ [0.89, 0.93]; Node41 (deep internal branch; $p = 1.00$), $\Theta = 0.09$ [0.07, 0.11]; (M,N) 32X: Neanderthal_Lescottes ($p = 1.00$), $\Theta = 0.91$ [0.90, 0.92]; Node41 (deep internal branch; $p = 1.00$), $\Theta = 0.09$ [0.08, 0.10].

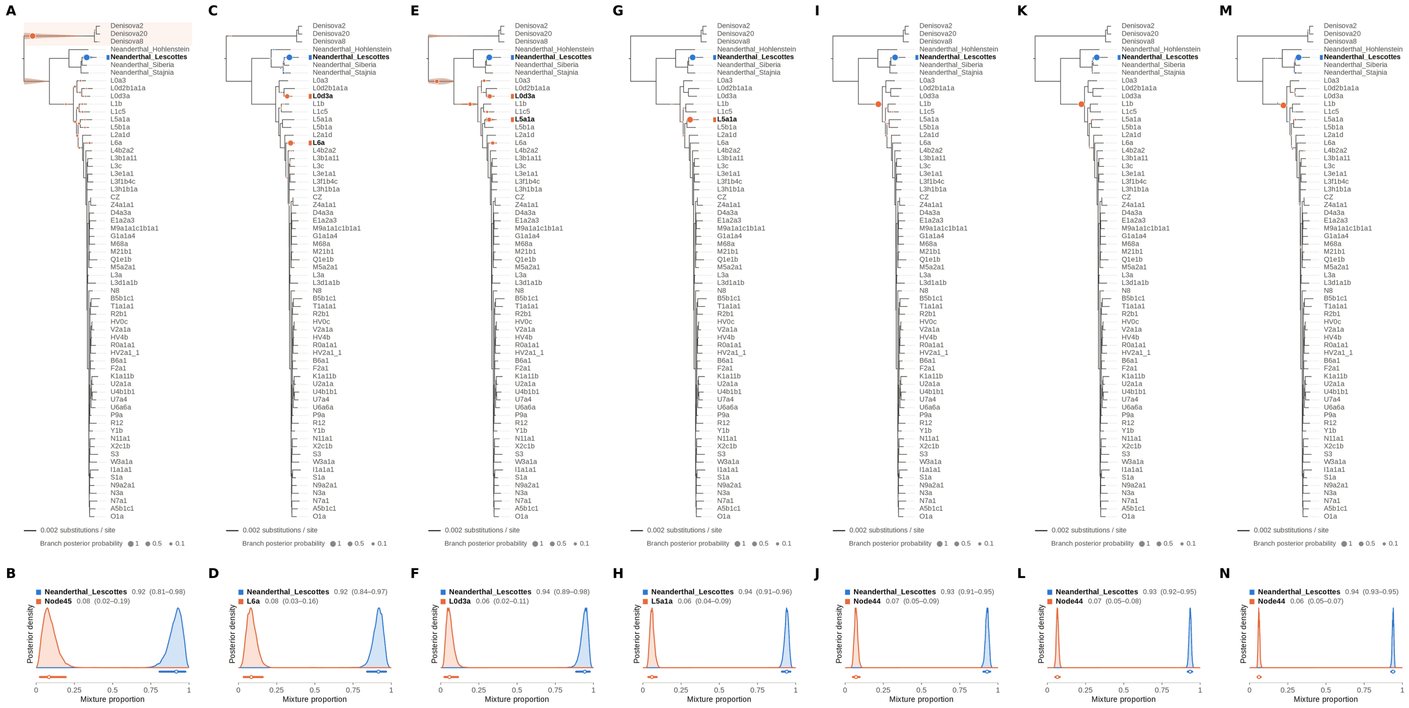


Figure S32: **Two-source simulated mixture of Chagyrskaya 8 Neanderthal (MK388903.1) (99% of reads) and modern human U5a2a (HM229344.1; 1%) at nominal mean coverage depths of 0.5X, 1X, 2X, 4X, 8X, 16X, 32X.** Both sources carry ancient DNA damage (U5a2a with the damage rates scaled by 0.75). For each coverage depth, the upper panel shows the phylogenetic placement posterior of both sources for the best of five chains ($k = 2$, 100,000 iterations each), colored by source (dot area: branch posterior probability; swelling: posterior density of the position along the branch; bold tips: highest-posterior branches), and the lower panel the posterior densities of the mixture proportions $\Theta$ with median and 95% credible interval. Chagyrskaya 8 is expected within the Neanderthal clade, on or near the Neanderthal_Lescottes branch (its closest relative in the reference); U5a2a, for which no U5 haplotype is present in the reference, is expected within the U clade. Modal branch of each source in the best chain ($p$: its posterior probability) and $\Theta$ median [95% CI]: (A,B) 0.5X: Neanderthal_Lescottes ($p = 1.00$), $\Theta = 0.92$ [0.81, 0.98]; Node45 (stem of the Denisovan clade; $p = 0.84$), $\Theta = 0.08$ [0.02, 0.19]; (C,D) 1X: Neanderthal_Lescottes ($p = 0.99$), $\Theta = 0.92$ [0.84, 0.97]; L6a ($p = 0.63$), $\Theta = 0.08$ [0.03, 0.16]; (E,F) 2X: Neanderthal_Lescottes ($p = 1.00$), $\Theta = 0.94$ [0.89, 0.98]; L0d3a ($p = 0.30$), $\Theta = 0.06$ [0.02, 0.11]; (G,H) 4X: Neanderthal_Lescottes ($p = 1.00$), $\Theta = 0.94$ [0.91, 0.96]; L5a1a ($p = 1.00$), $\Theta = 0.06$ [0.04, 0.09]; (I,J) 8X: Neanderthal_Lescottes ($p = 1.00$), $\Theta = 0.93$ [0.91, 0.95]; Node44 (stem of the archaic hominin clade; $p = 0.99$), $\Theta = 0.07$ [0.05, 0.09]; (K,L) 16X: Neanderthal_Lescottes ($p = 1.00$), $\Theta = 0.93$ [0.92, 0.95]; Node44 (stem of the archaic hominin clade; $p = 0.99$), $\Theta = 0.07$ [0.05, 0.08]; (M,N) 32X: Neanderthal_Lescottes ($p = 0.99$), $\Theta = 0.94$ [0.93, 0.95]; Node44 (stem of the archaic hominin clade; $p = 0.99$), $\Theta = 0.06$ [0.05, 0.07].

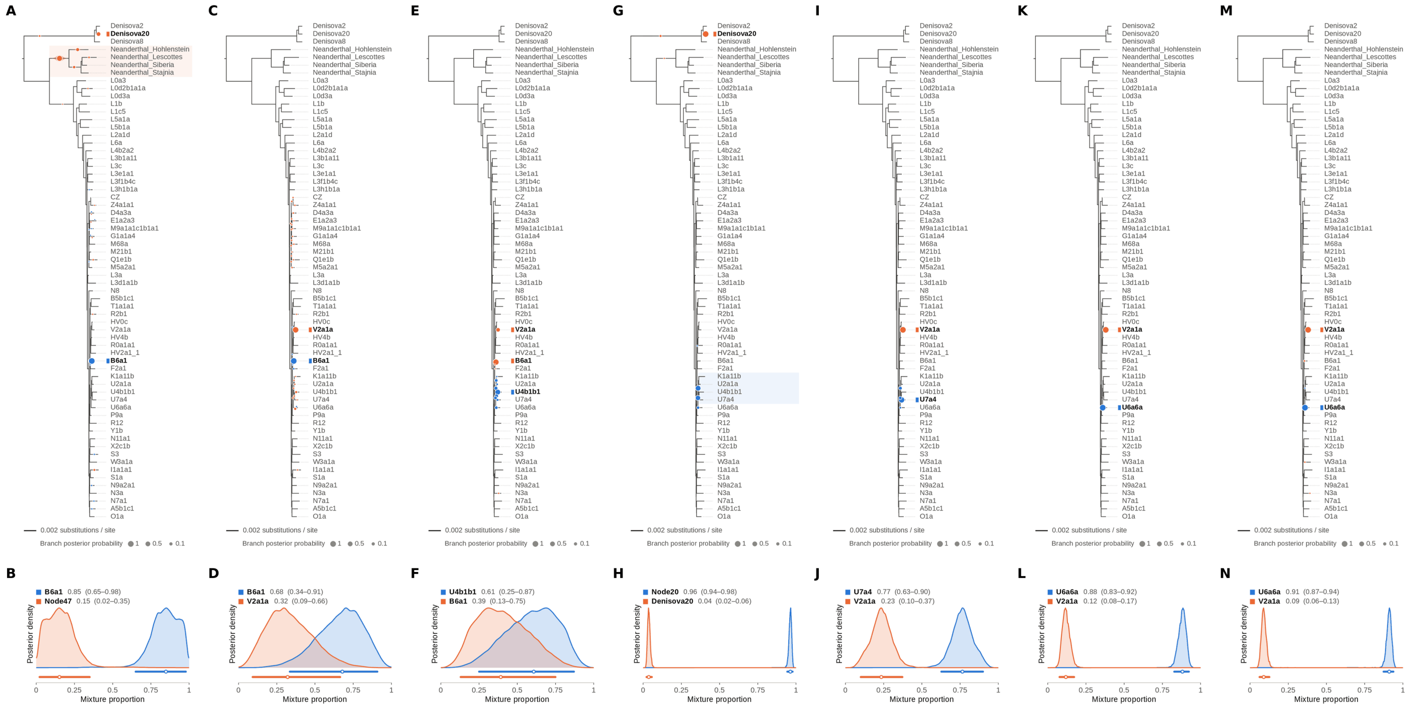


Figure S33: **Two-source simulated mixture of Denisova 21 (MT576653.1) (1% of reads) and modern human U5a2a (HM229344.1; 99%) at nominal mean coverage depths of 0.5X, 1X, 2X, 4X, 8X, 16X, 32X.** Both sources carry ancient DNA damage (U5a2a with the damage rates scaled by 0.75). For each coverage depth, the upper panel shows the phylogenetic placement posterior of both sources for the best of five chains ($k$ = 2, 100,000 iterations each), colored by source (dot area: branch posterior probability; swelling: posterior density of the position along the branch; bold tips: highest-posterior branches), and the lower panel the posterior densities of the mixture proportions $\Theta$ with median and 95% credible interval. Denisova 21 is expected on the Denisova20 branch (its closest relative in the reference); U5a2a, for which no U5 haplotype is present in the reference, is expected within the U clade. Modal branch of each source in the best chain ($p$: its posterior probability) and $\Theta$ median [95% CI]: (A,B) 0.5X: B6a1 ($p = 0.99$), $\Theta$ = 0.85 [0.65, 0.98]; Node47 (stem of the Neanderthal clade; $p = 0.66$), $\Theta$ = 0.15 [0.02, 0.35]; (C,D) 1X: B6a1 ($p = 1.00$), $\Theta$ = 0.68 [0.34, 0.91]; V2a1a ($p = 0.91$), $\Theta$ = 0.32 [0.09, 0.66]; (E,F) 2X: U4b1b1 ($p = 0.58$), $\Theta$ = 0.61 [0.25, 0.87]; B6a1 ($p = 0.82$), $\Theta$ = 0.39 [0.13, 0.75]; (G,H) 4X: Node20 (internal branch within the U clade; $p = 0.54$), $\Theta$ = 0.96 [0.94, 0.98]; Denisova20 ($p = 0.98$), $\Theta$ = 0.04 [0.02, 0.06]; (I,J) 8X: U7a4 ($p = 0.85$), $\Theta$ = 0.77 [0.63, 0.90]; V2a1a ($p = 1.00$), $\Theta$ = 0.23 [0.10, 0.37]; (K,L) 16X: U6a6a ($p = 1.00$), $\Theta$ = 0.88 [0.83, 0.92]; V2a1a ($p = 1.00$), $\Theta$ = 0.12 [0.08, 0.17]; (M,N) 32X: U6a6a ($p = 1.00$), $\Theta$ = 0.91 [0.87, 0.94]; V2a1a ($p = 0.99$), $\Theta$ = 0.09 [0.06, 0.13].

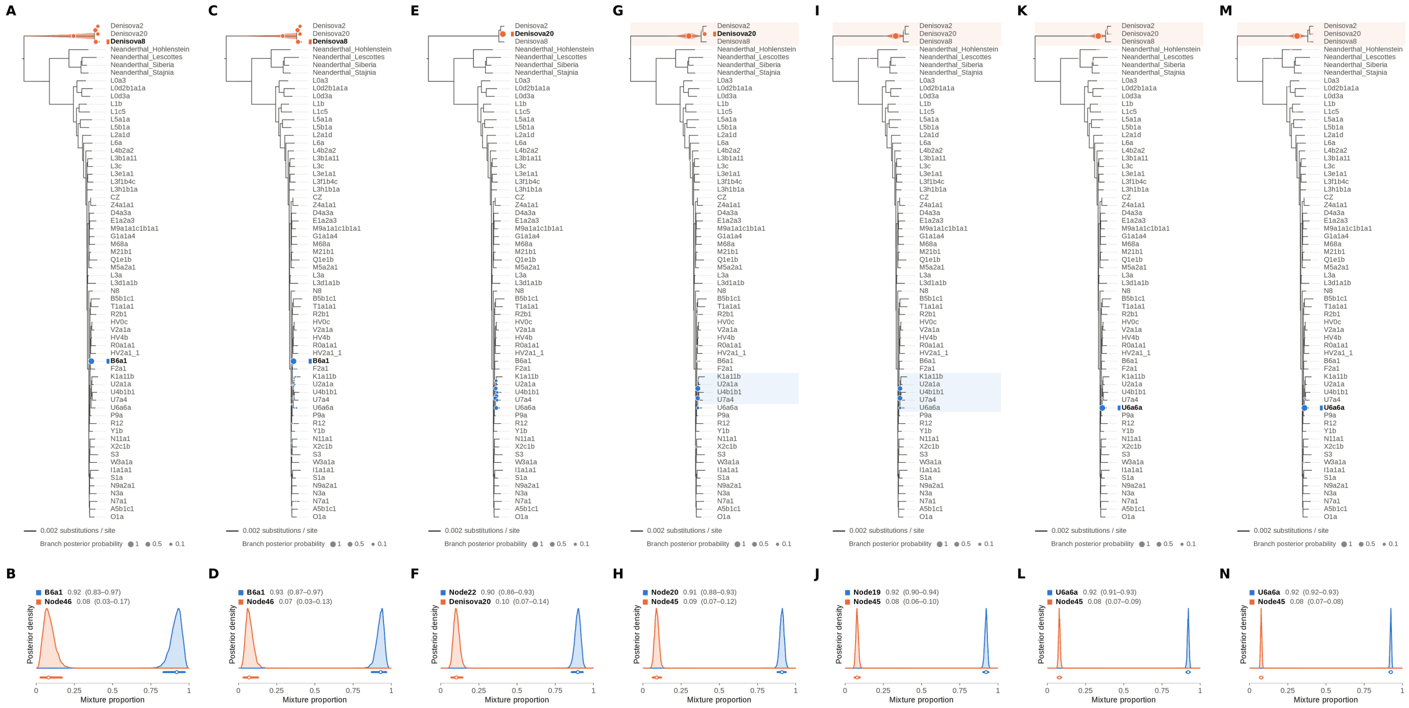


Figure S34: **Two-source simulated mixture of Denisova 21 (MT576653.1) (5% of reads) and modern human U5a2a (HM229344.1; 95%) at nominal mean coverage depths of 0.5X, 1X, 2X, 4X, 8X, 16X, 32X.** Both sources carry ancient DNA damage (U5a2a with the damage rates scaled by 0.75). For each coverage depth, the upper panel shows the phylogenetic placement posterior of both sources for the best of five chains ($k$ = 2, 100,000 iterations each), colored by source (dot area: branch posterior probability; swelling: posterior density of the position along the branch; bold tips: highest-posterior branches), and the lower panel the posterior densities of the mixture proportions $\Theta$ with median and 95% credible interval. Denisova 21 is expected on the Denisova20 branch (its closest relative in the reference); U5a2a, for which no U5 haplotype is present in the reference, is expected within the U clade. Modal branch of each source in the best chain ($p$: its posterior probability) and $\Theta$ median [95% CI]: (A,B) 0.5X: B6a1 ($p = 1.00$), $\Theta = 0.92$ [0.83, 0.97]; Node46 (internal branch ancestral to Denisova2 and Denisova20; $p = 0.27$), $\Theta = 0.08$ [0.03, 0.17]; (C,D) 1X: B6a1 ($p = 0.99$), $\Theta = 0.93$ [0.87, 0.97]; Node46 (internal branch ancestral to Denisova2 and Denisova20; $p = 0.30$), $\Theta = 0.07$ [0.03, 0.13]; (E,F) 2X: Node22 (internal branch ancestral to U4b1b1 and U7a4; $p = 0.28$), $\Theta = 0.90$ [0.86, 0.93]; Denisova20 ($p = 1.00$), $\Theta = 0.10$ [0.07, 0.14]; (G,H) 4X: Node20 (internal branch within the U clade; $p = 0.60$), $\Theta = 0.91$ [0.88, 0.93]; Node45 (stem of the Denisovan clade; $p = 0.79$), $\Theta = 0.09$ [0.07, 0.12]; (I,J) 8X: Node19 (stem of the U clade; $p = 0.51$), $\Theta = 0.92$ [0.90, 0.94]; Node45 (stem of the Denisovan clade; $p = 1.00$), $\Theta = 0.08$ [0.06, 0.10]; (K,L) 16X: U6a6a ($p = 1.00$), $\Theta = 0.92$ [0.91, 0.93]; Node45 (stem of the Denisovan clade; $p = 1.00$), $\Theta = 0.08$ [0.07, 0.09]; (M,N) 32X: U6a6a ($p = 1.00$), $\Theta = 0.92$ [0.92, 0.93]; Node45 (stem of the Denisovan clade; $p = 1.00$), $\Theta = 0.08$ [0.07, 0.08].

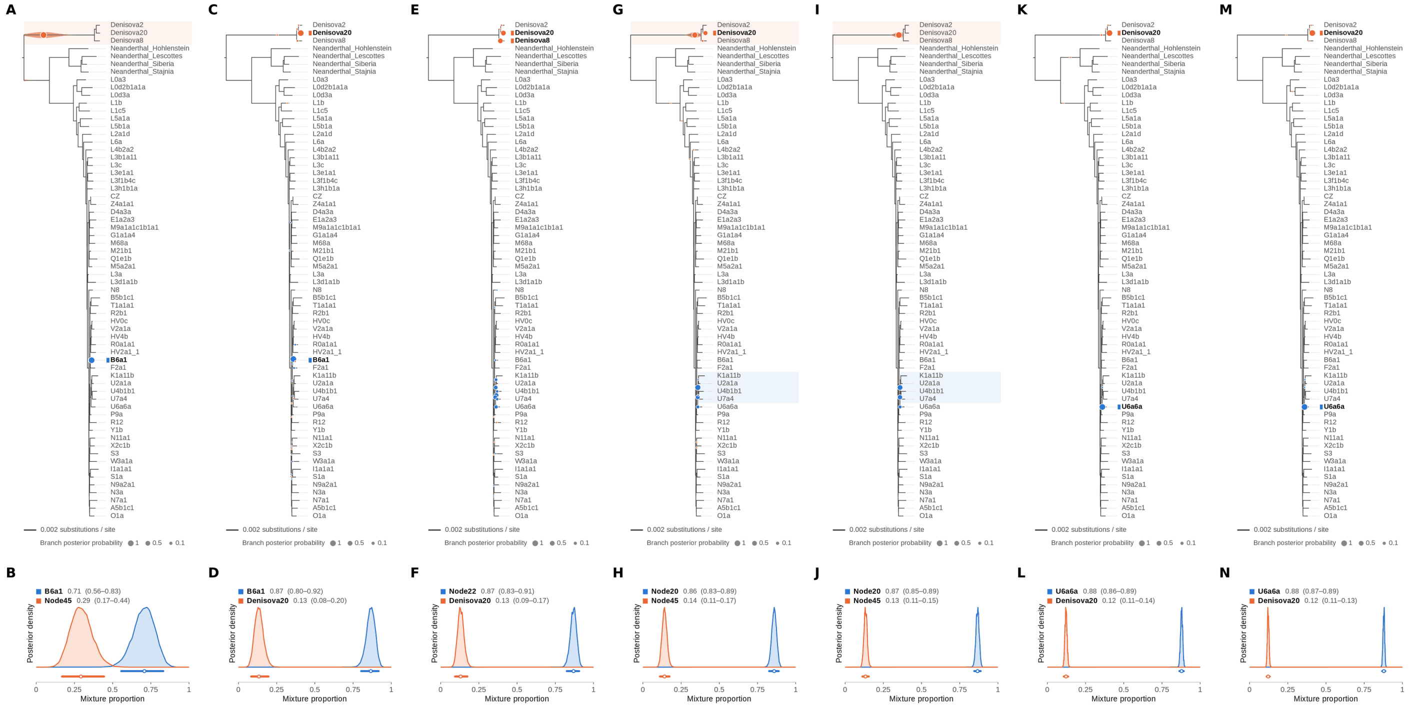


Figure S35: **Two-source simulated mixture of Denisova 21 (MT576653.1) (10% of reads) and modern human U5a2a (HM229344.1; 90%) at nominal mean coverage depths of 0.5X, 1X, 2X, 4X, 8X, 16X, 32X.** Both sources carry ancient DNA damage (U5a2a with the damage rates scaled by 0.75). For each coverage depth, the upper panel shows the phylogenetic placement posterior of both sources for the best of five chains ($k = 2$, 100,000 iterations each), colored by source (dot area: branch posterior probability; swelling: posterior density of the position along the branch; bold tips: highest-posterior branches), and the lower panel the posterior densities of the mixture proportions $\Theta$ with median and 95% credible interval. Denisova 21 is expected on the Denisova20 branch (its closest relative in the reference); U5a2a, for which no U5 haplotype is present in the reference, is expected within the U clade. Modal branch of each source in the best chain ($p$: its posterior probability) and $\Theta$ median [95% CI]: (A,B) 0.5X: B6a1 ($p = 1.00$), $\Theta = 0.71$ [0.56, 0.83]; Node45 (stem of the Denisovan clade; $p = 1.00$), $\Theta = 0.29$ [0.17, 0.44]; (C,D) 1X: B6a1 ($p = 0.98$), $\Theta = 0.87$ [0.80, 0.92]; Denisova20 ($p = 0.99$), $\Theta = 0.13$ [0.08, 0.20]; (E,F) 2X: Node22 (internal branch ancestral to U4b1b1 and U7a4; $p = 0.35$), $\Theta = 0.87$ [0.83, 0.91]; Denisova20 ($p = 0.52$), $\Theta = 0.13$ [0.09, 0.17]; (G,H) 4X: Node20 (internal branch within the U clade; $p = 0.69$), $\Theta = 0.86$ [0.83, 0.89]; Node45 (stem of the Denisovan clade; $p = 0.70$), $\Theta = 0.14$ [0.11, 0.17]; (I,J) 8X: Node20 (internal branch within the U clade; $p = 0.54$), $\Theta = 0.87$ [0.85, 0.89]; Node45 (stem of the Denisovan clade; $p = 1.00$), $\Theta = 0.13$ [0.11, 0.15]; (K,L) 16X: U6a6a ($p = 1.00$), $\Theta = 0.88$ [0.86, 0.89]; Denisova20 ($p = 0.99$), $\Theta = 0.12$ [0.11, 0.14]; (M,N) 32X: U6a6a ($p = 1.00$), $\Theta = 0.88$ [0.87, 0.89]; Denisova20 ($p = 1.00$), $\Theta = 0.12$ [0.11, 0.13].

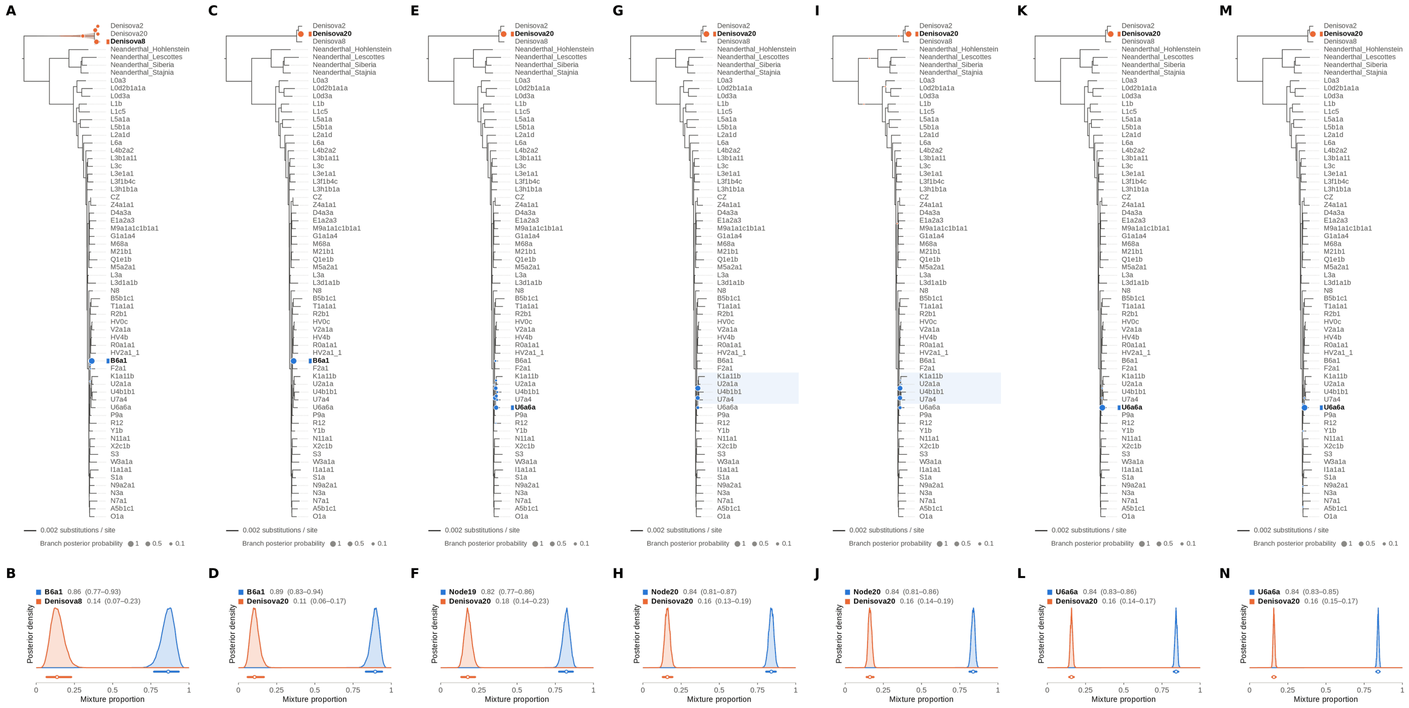


Figure S36: **Two-source simulated mixture of Denisova 21 (MT576653.1) (15% of reads) and modern human U5a2a (HM229344.1; 85%) at nominal mean coverage depths of 0.5X, 1X, 2X, 4X, 8X, 16X, 32X.** Both sources carry ancient DNA damage (U5a2a with the damage rates scaled by 0.75). For each coverage depth, the upper panel shows the phylogenetic placement posterior of both sources for the best of five chains ($k = 2$, 100,000 iterations each), colored by source (dot area: branch posterior probability; swelling: posterior density of the position along the branch; bold tips: highest-posterior branches), and the lower panel the posterior densities of the mixture proportions $\Theta$ with median and 95% credible interval. Denisova 21 is expected on the Denisova20 branch (its closest relative in the reference); U5a2a, for which no U5 haplotype is present in the reference, is expected within the U clade. Modal branch of each source in the best chain ($p$: its posterior probability) and $\Theta$ median [95% CI]: (A,B) 0.5X: B6a1 ($p = 1.00$), $\Theta = 0.86$ [0.77, 0.93]; Denisova8 ($p = 0.44$), $\Theta = 0.14$ [0.07, 0.23]; (C,D) 1X: B6a1 ($p = 1.00$), $\Theta = 0.89$ [0.83, 0.94]; Denisova20 ($p = 1.00$), $\Theta = 0.11$ [0.06, 0.17]; (E,F) 2X: Node19 (stem of the U clade; $p = 0.37$), $\Theta = 0.82$ [0.77, 0.86]; Denisova20 ($p = 1.00$), $\Theta = 0.18$ [0.14, 0.23]; (G,H) 4X: Node20 (internal branch within the U clade; $p = 0.65$), $\Theta = 0.84$ [0.81, 0.87]; Denisova20 ($p = 1.00$), $\Theta = 0.16$ [0.13, 0.19]; (I,J) 8X: Node20 (internal branch within the U clade; $p = 0.52$), $\Theta = 0.84$ [0.81, 0.86]; Denisova20 ($p = 0.99$), $\Theta = 0.16$ [0.14, 0.19]; (K,L) 16X: U6a6a ($p = 1.00$), $\Theta = 0.84$ [0.83, 0.86]; Denisova20 ($p = 1.00$), $\Theta = 0.16$ [0.14, 0.17]; (M,N) 32X: U6a6a ($p = 1.00$), $\Theta = 0.84$ [0.83, 0.85]; Denisova20 ($p = 1.00$), $\Theta = 0.16$ [0.15, 0.17].

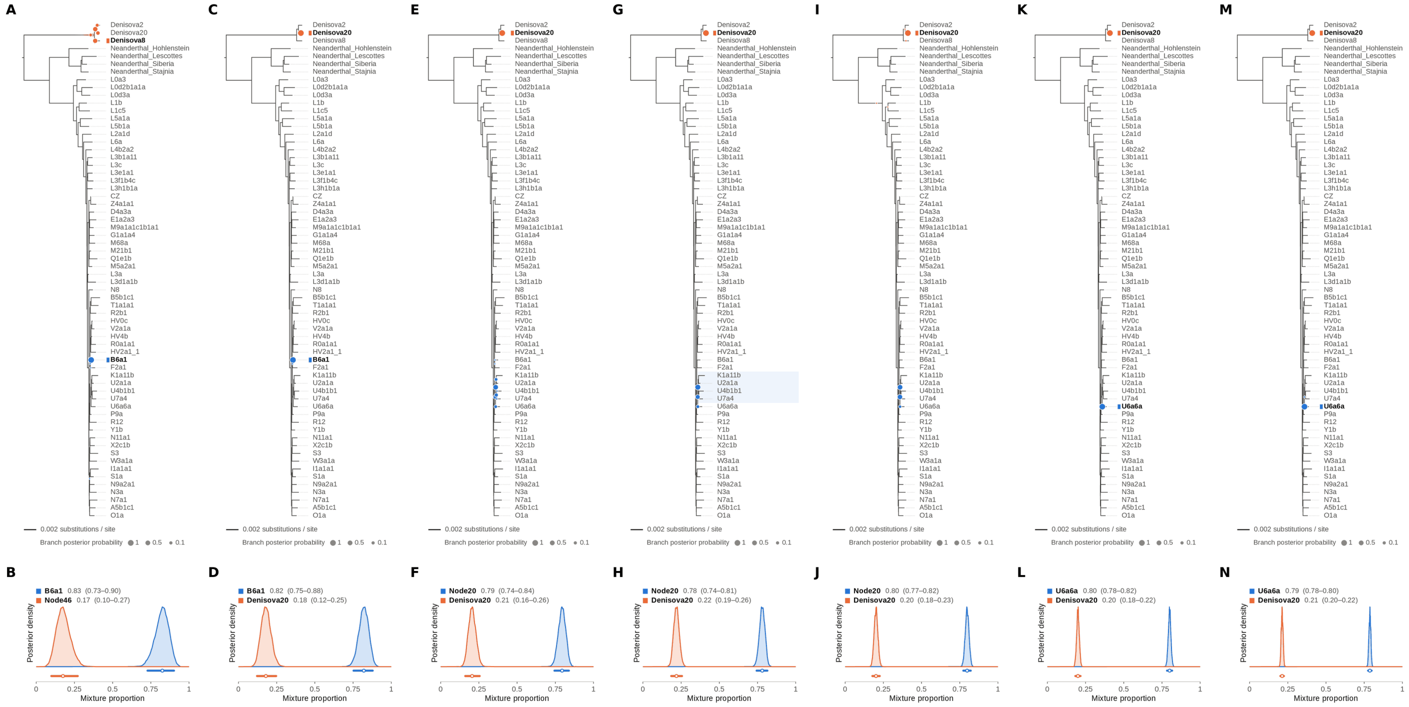

Figure S37: **Two-source simulated mixture of Denisova 21 (MT576653.1) (20% of reads) and modern human U5a2a (HM229344.1; 80%) at nominal mean coverage depths of 0.5X, 1X, 2X, 4X, 8X, 16X, 32X.** Both sources carry ancient DNA damage (U5a2a with the damage rates scaled by 0.75). For each coverage depth, the upper panel shows the phylogenetic placement posterior of both sources for the best of five chains ($k = 2$, 100,000 iterations each), colored by source (dot area: branch posterior probability; swelling: posterior density of the position along the branch; bold tips: highest-posterior branches), and the lower panel the posterior densities of the mixture proportions $\Theta$ with median and 95% credible interval. Denisova 21 is expected on the Denisova20 branch (its closest relative in the reference); U5a2a, for which no U5 haplotype is present in the reference, is expected within the U clade. Modal branch of each source in the best chain ($p$: its posterior probability) and $\Theta$ median [95% CI]: (A,B) 0.5X: B6a1 ($p = 1.00$), $\Theta = 0.83$ [0.73, 0.90]; Node46 (internal branch ancestral to Denisova2 and Denisova20; $p = 0.37$), $\Theta = 0.17$ [0.10, 0.27]; (C,D) 1X: B6a1 ($p = 1.00$), $\Theta = 0.82$ [0.75, 0.88]; Denisova20 ($p = 1.00$), $\Theta = 0.18$ [0.12, 0.25]; (E,F) 2X: Node20 (internal branch within the U clade; $p = 0.50$), $\Theta = 0.79$ [0.74, 0.84]; Denisova20 ($p = 1.00$), $\Theta = 0.21$ [0.16, 0.26]; (G,H) 4X: Node20 (internal branch within the U clade; $p = 0.64$), $\Theta = 0.78$ [0.74, 0.81]; Denisova20 ($p = 1.00$), $\Theta = 0.22$ [0.19, 0.26]; (I,J) 8X: Node20 (internal branch within the U clade; $p = 0.50$), $\Theta = 0.80$ [0.77, 0.82]; Denisova20 ($p = 1.00$), $\Theta = 0.20$ [0.18, 0.23]; (K,L) 16X: U6a6a ($p = 1.00$), $\Theta = 0.80$ [0.78, 0.82]; Denisova20 ($p = 1.00$), $\Theta = 0.20$ [0.18, 0.22]; (M,N) 32X: U6a6a ($p = 1.00$), $\Theta = 0.79$ [0.78, 0.80]; Denisova20 ($p = 1.00$), $\Theta = 0.21$ [0.20, 0.22].

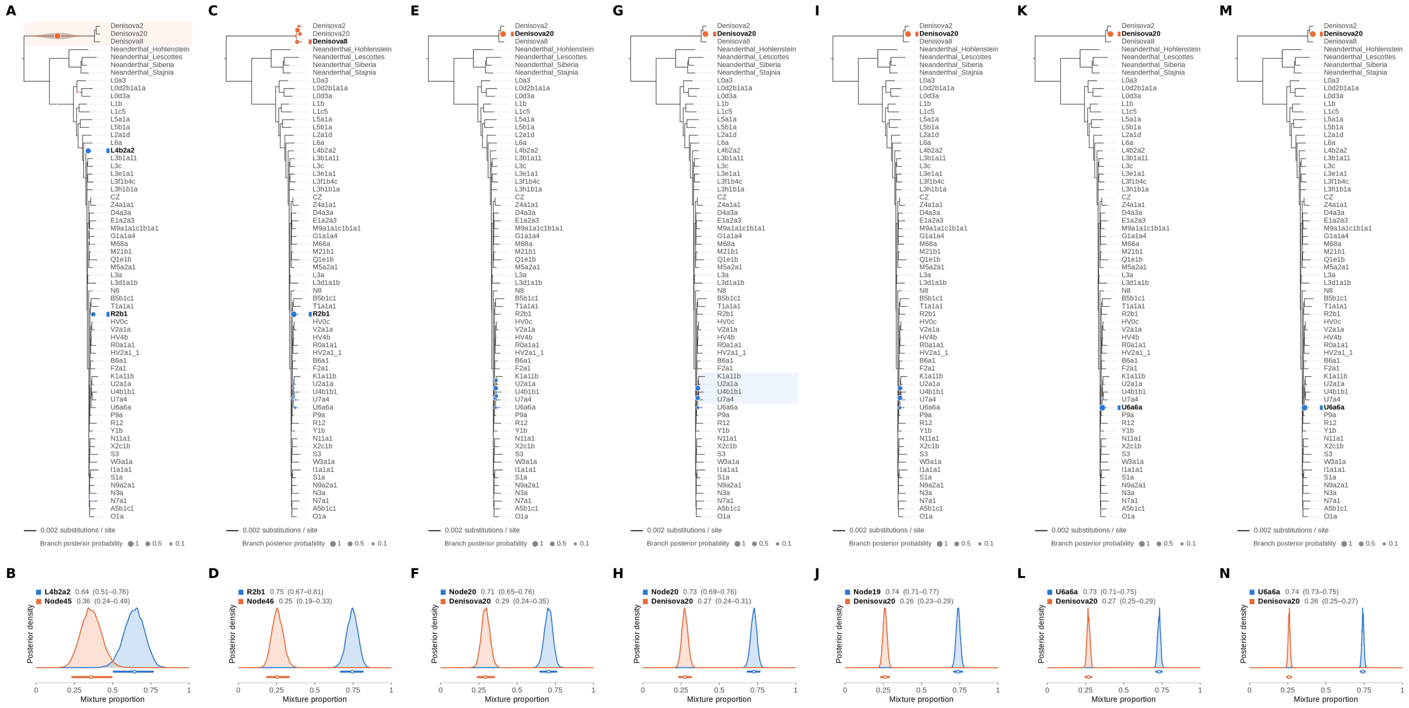


Figure S38: **Two-source simulated mixture of Denisova 21 (MT576653.1) (25% of reads) and modern human U5a2a (HM229344.1; 75%) at nominal mean coverage depths of 0.5X, 1X, 2X, 4X, 8X, 16X, 32X.** Both sources carry ancient DNA damage (U5a2a with the damage rates scaled by 0.75). For each coverage depth, the upper panel shows the phylogenetic placement posterior of both sources for the best of five chains ($k$ = 2, 100,000 iterations each), colored by source (dot area: branch posterior probability; swelling: posterior density of the position along the branch; bold tips: highest-posterior branches), and the lower panel the posterior densities of the mixture proportions $\Theta$ with median and 95% credible interval. Denisova 21 is expected on the Denisova20 branch (its closest relative in the reference); U5a2a, for which no U5 haplotype is present in the reference, is expected within the U clade. Modal branch of each source in the best chain ($p$: its posterior probability) and $\Theta$ median [95% CI]: (A,B) 0.5X: L4b2a2 ($p$ = 0.68), $\Theta$ = 0.64 [0.51, 0.76]; Node45 (stem of the Denisovan clade; $p$ = 1.00), $\Theta$ = 0.36 [0.24, 0.49]; (C,D) 1X: R2b1 ($p$ = 0.92), $\Theta$ = 0.75 [0.67, 0.81]; Node46 (internal branch ancestral to Denisova2 and Denisova20; $p$ = 0.45), $\Theta$ = 0.25 [0.19, 0.33]; (E,F) 2X: Node20 (internal branch within the U clade; $p$ = 0.43), $\Theta$ = 0.71 [0.65, 0.76]; Denisova20 ($p$ = 1.00), $\Theta$ = 0.29 [0.24, 0.35]; (G,H) 4X: Node20 (internal branch within the U clade; $p$ = 0.55), $\Theta$ = 0.73 [0.69, 0.76]; Denisova20 ($p$ = 1.00), $\Theta$ = 0.27 [0.24, 0.31]; (I,J) 8X: Node19 (stem of the U clade; $p$ = 0.48), $\Theta$ = 0.74 [0.71, 0.77]; Denisova20 ($p$ = 1.00), $\Theta$ = 0.26 [0.23, 0.29]; (K,L) 16X: U6a6a ($p$ = 1.00), $\Theta$ = 0.73 [0.71, 0.75]; Denisova20 ($p$ = 1.00), $\Theta$ = 0.27 [0.25, 0.29]; (M,N) 32X: U6a6a ($p$ = 1.00), $\Theta$ = 0.74 [0.73, 0.75]; Denisova20 ($p$ = 1.00), $\Theta$ = 0.26 [0.25, 0.27].

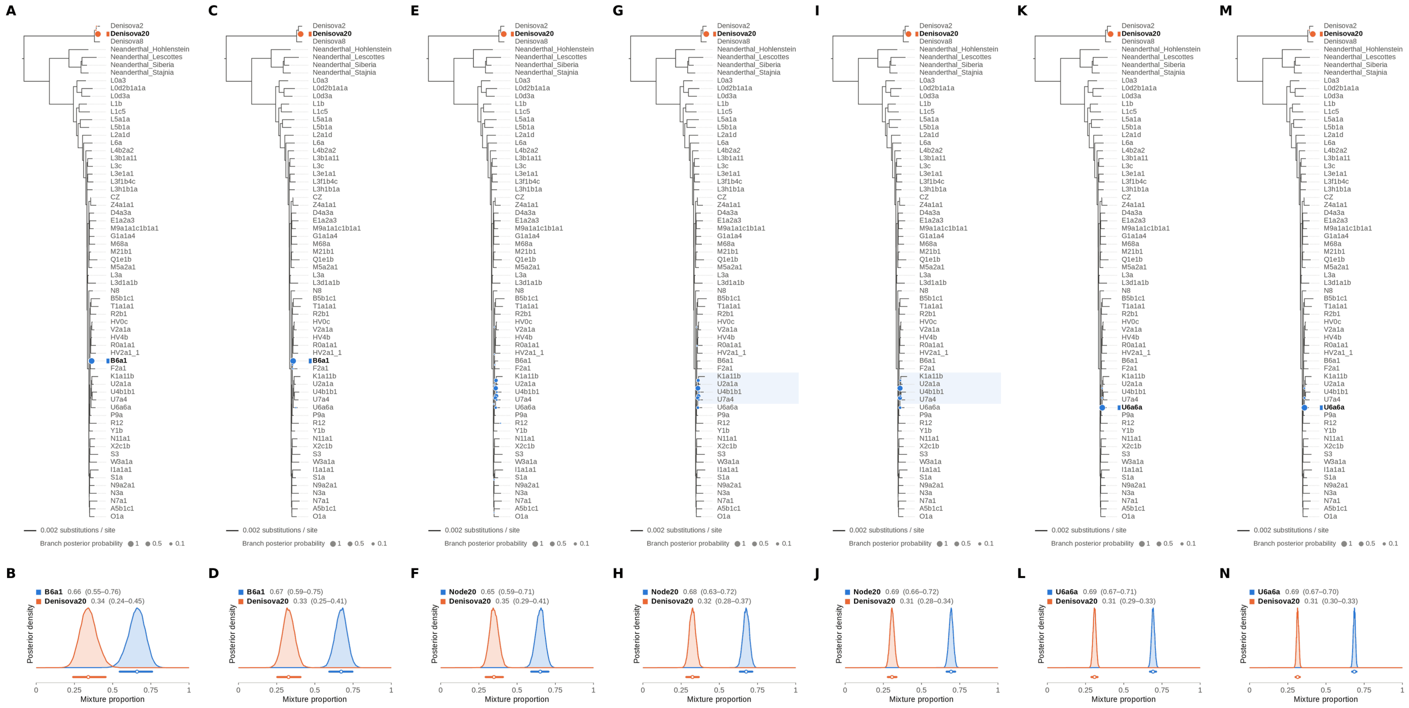


Figure S39: **Two-source simulated mixture of Denisova 21 (MT576653.1) (30% of reads) and modern human U5a2a (HM229344.1; 70%) at nominal mean coverage depths of 0.5X, 1X, 2X, 4X, 8X, 16X, 32X.** The runs of this mixture at 0.5X, 1X, 2X and 8X are also shown, with collapsed trees, in main Figure 4. Both sources carry ancient DNA damage (U5a2a with the damage rates scaled by 0.75). For each coverage depth, the upper panel shows the phylogenetic placement posterior of both sources for the best of five chains ($k = 2$, 100,000 iterations each), colored by source (dot area: branch posterior probability; swelling: posterior density of the position along the branch; bold tips: highest-posterior branches), and the lower panel the posterior densities of the mixture proportions $\Theta$ with median and 95% credible interval. Denisova 21 is expected on the Denisova20 branch (its closest relative in the reference); U5a2a, for which no U5 haplotype is present in the reference, is expected within the U clade. Modal branch of each source in the best chain ($p$: its posterior probability) and $\Theta$ median [95% CI]: (A,B) 0.5X: B6a1 ($p = 1.00$), $\Theta = 0.66$ [0.55, 0.76]; Denisova20 ($p = 1.00$), $\Theta = 0.34$ [0.24, 0.45]; (C,D) 1X: B6a1 ($p = 0.99$), $\Theta = 0.67$ [0.59, 0.75]; Denisova20 ($p = 1.00$), $\Theta = 0.33$ [0.25, 0.41]; (E,F) 2X: Node20 (internal branch within the U clade; $p = 0.37$), $\Theta = 0.65$ [0.59, 0.71]; Denisova20 ($p = 1.00$), $\Theta = 0.35$ [0.29, 0.41]; (G,H) 4X: Node20 (internal branch within the U clade; $p = 0.51$), $\Theta = 0.68$ [0.63, 0.72]; Denisova20 ($p = 1.00$), $\Theta = 0.32$ [0.28, 0.37]; (I,J) 8X: Node20 (internal branch within the U clade; $p = 0.67$), $\Theta = 0.69$ [0.66, 0.72]; Denisova20 ($p = 1.00$), $\Theta = 0.31$ [0.28, 0.34]; (K,L) 16X: U6a6a ($p = 1.00$), $\Theta = 0.69$ [0.67, 0.71]; Denisova20 ($p = 1.00$), $\Theta = 0.31$ [0.29, 0.33]; (M,N) 32X: U6a6a ($p = 0.98$), $\Theta = 0.69$ [0.67, 0.70]; Denisova20 ($p = 1.00$), $\Theta = 0.31$ [0.30, 0.33].

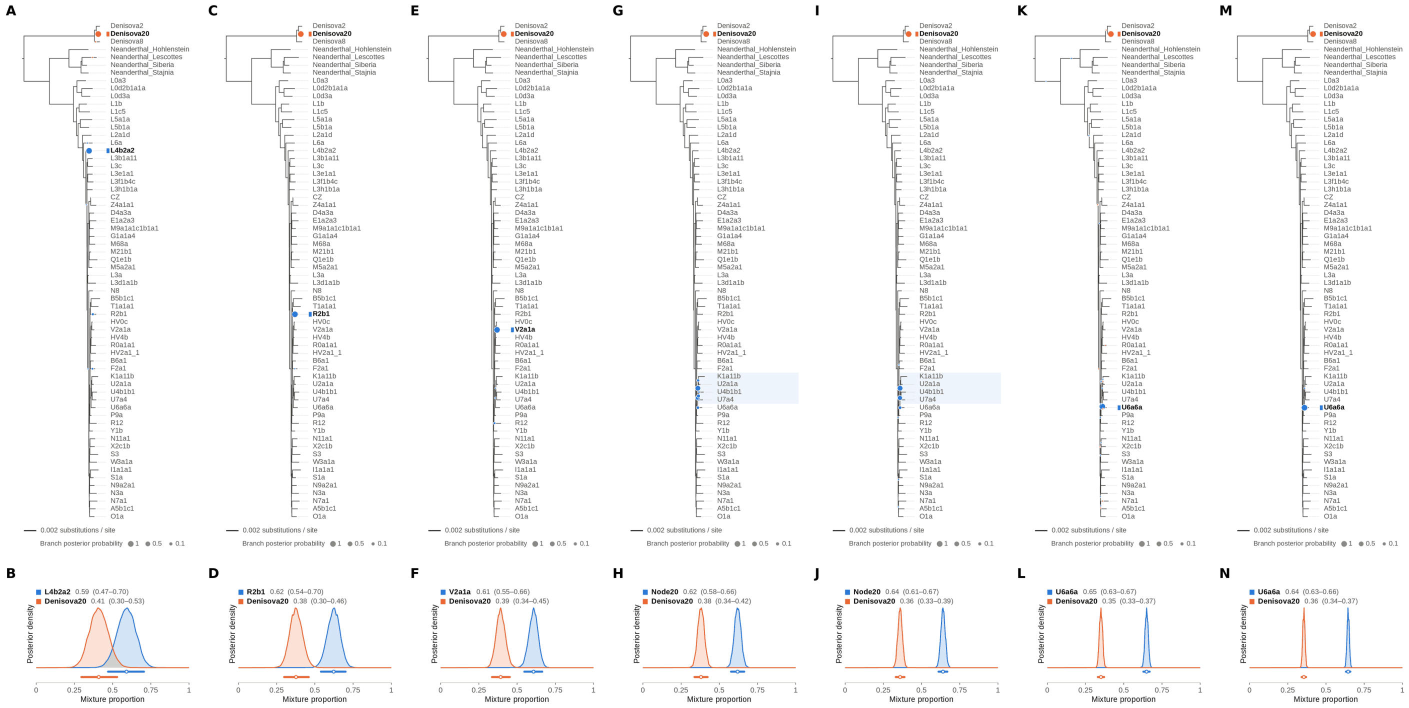

Figure S40: **Two-source simulated mixture of Denisova 21 (MT576653.1) (35% of reads) and modern human U5a2a (HM229344.1; 65%) at nominal mean coverage depths of 0.5X, 1X, 2X, 4X, 8X, 16X, 32X.** Both sources carry ancient DNA damage (U5a2a with the damage rates scaled by 0.75). For each coverage depth, the upper panel shows the phylogenetic placement posterior of both sources for the best of five chains ($k$ = 2, 100,000 iterations each), colored by source (dot area: branch posterior probability; swelling: posterior density of the position along the branch; bold tips: highest-posterior branches), and the lower panel the posterior densities of the mixture proportions $\Theta$ with median and 95% credible interval. Denisova 21 is expected on the Denisova20 branch (its closest relative in the reference); U5a2a, for which no U5 haplotype is present in the reference, is expected within the U clade. Modal branch of each source in the best chain ($p$: its posterior probability) and $\Theta$ median [95% CI]: (A,B) 0.5X: L4b2a2 ($p$ = 0.97), $\Theta$ = 0.59 [0.47, 0.70]; Denisova20 ($p$ = 1.00), $\Theta$ = 0.41 [0.30, 0.53]; (C,D) 1X: R2b1 ($p$ = 1.00), $\Theta$ = 0.62 [0.54, 0.70]; Denisova20 ($p$ = 1.00), $\Theta$ = 0.38 [0.30, 0.46]; (E,F) 2X: V2a1a ($p$ = 0.99), $\Theta$ = 0.61 [0.55, 0.66]; Denisova20 ($p$ = 1.00), $\Theta$ = 0.39 [0.34, 0.45]; (G,H) 4X: Node20 (internal branch within the U clade; $p$ = 0.55), $\Theta$ = 0.62 [0.58, 0.66]; Denisova20 ($p$ = 1.00), $\Theta$ = 0.38 [0.34, 0.42]; (I,J) 8X: Node20 (internal branch within the U clade; $p$ = 0.59), $\Theta$ = 0.64 [0.61, 0.67]; Denisova20 ($p$ = 1.00), $\Theta$ = 0.36 [0.33, 0.39]; (K,L) 16X: U6a6a ($p$ = 1.00), $\Theta$ = 0.65 [0.63, 0.67]; Denisova20 ($p$ = 1.00), $\Theta$ = 0.35 [0.33, 0.37]; (M,N) 32X: U6a6a ($p$ = 1.00), $\Theta$ = 0.64 [0.63, 0.66]; Denisova20 ($p$ = 1.00), $\Theta$ = 0.36 [0.34, 0.37].

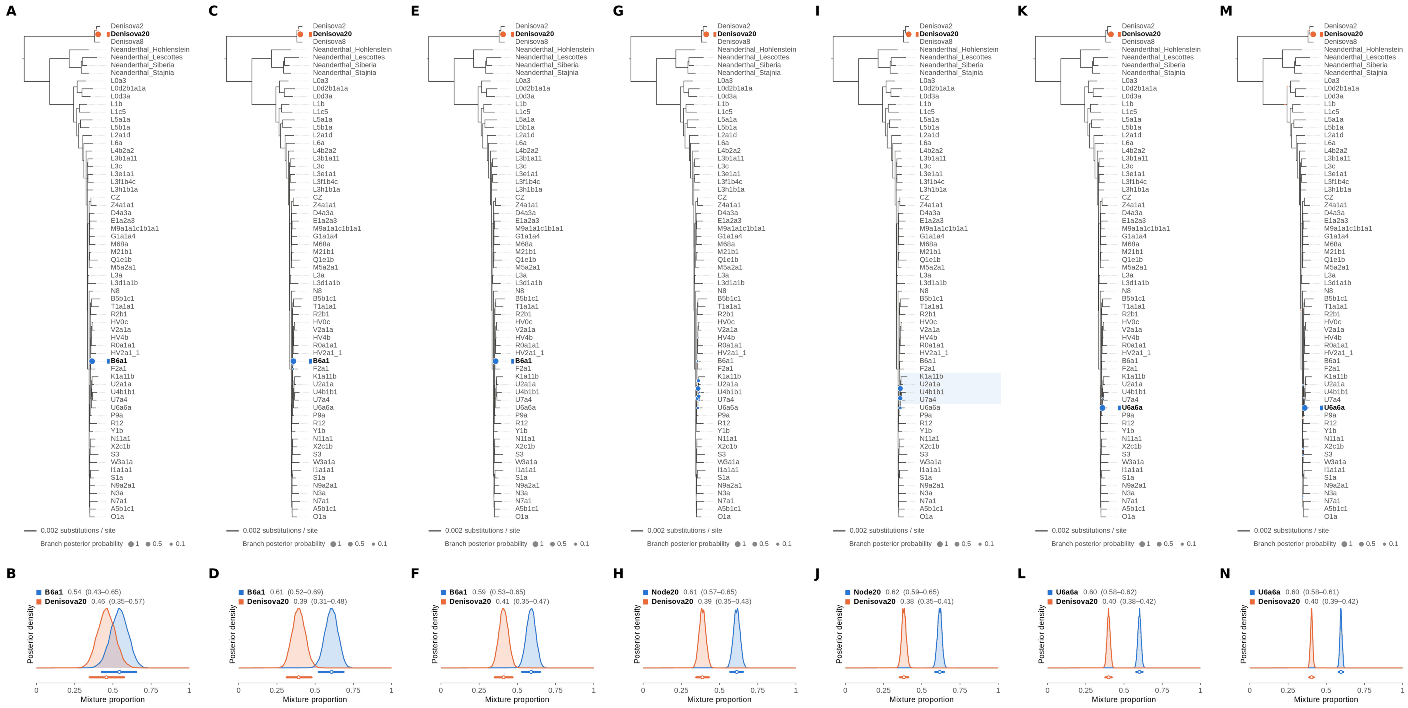

Figure S41: **Two-source simulated mixture of Denisova 21 (MT576653.1) (40% of reads) and modern human U5a2a (HM229344.1; 60%) at nominal mean coverage depths of 0.5X, 1X, 2X, 4X, 8X, 16X, 32X.** Both sources carry ancient DNA damage (U5a2a with the damage rates scaled by 0.75). For each coverage depth, the upper panel shows the phylogenetic placement posterior of both sources for the best of five chains ($k = 2$, 100,000 iterations each), colored by source (dot area: branch posterior probability; swelling: posterior density of the position along the branch; bold tips: highest-posterior branches), and the lower panel the posterior densities of the mixture proportions $\Theta$ with median and 95% credible interval. Denisova 21 is expected on the Denisova20 branch (its closest relative in the reference); U5a2a, for which no U5 haplotype is present in the reference, is expected within the U clade. Modal branch of each source in the best chain ($p$: its posterior probability) and $\Theta$ median [95% CI]: (A,B) 0.5X: B6a1 ($p = 1.00$), $\Theta = 0.54$ [0.43, 0.65]; Denisova20 ($p = 1.00$), $\Theta = 0.46$ [0.35, 0.57]; (C,D) 1X: B6a1 ($p = 1.00$), $\Theta = 0.61$ [0.52, 0.69]; Denisova20 ($p = 1.00$), $\Theta = 0.39$ [0.31, 0.48]; (E,F) 2X: B6a1 ($p = 1.00$), $\Theta = 0.59$ [0.53, 0.65]; Denisova20 ($p = 1.00$), $\Theta = 0.41$ [0.35, 0.47]; (G,H) 4X: Node20 (internal branch within the U clade; $p = 0.44$), $\Theta = 0.61$ [0.57, 0.65]; Denisova20 ($p = 1.00$), $\Theta = 0.39$ [0.35, 0.43]; (I,J) 8X: Node20 (internal branch within the U clade; $p = 0.60$), $\Theta = 0.62$ [0.59, 0.65]; Denisova20 ($p = 1.00$), $\Theta = 0.38$ [0.35, 0.41]; (K,L) 16X: U6a6a ($p = 1.00$), $\Theta = 0.60$ [0.58, 0.62]; Denisova20 ($p = 1.00$), $\Theta = 0.40$ [0.38, 0.42]; (M,N) 32X: U6a6a ($p = 1.00$), $\Theta = 0.60$ [0.58, 0.61]; Denisova20 ($p = 1.00$), $\Theta = 0.40$ [0.39, 0.42].

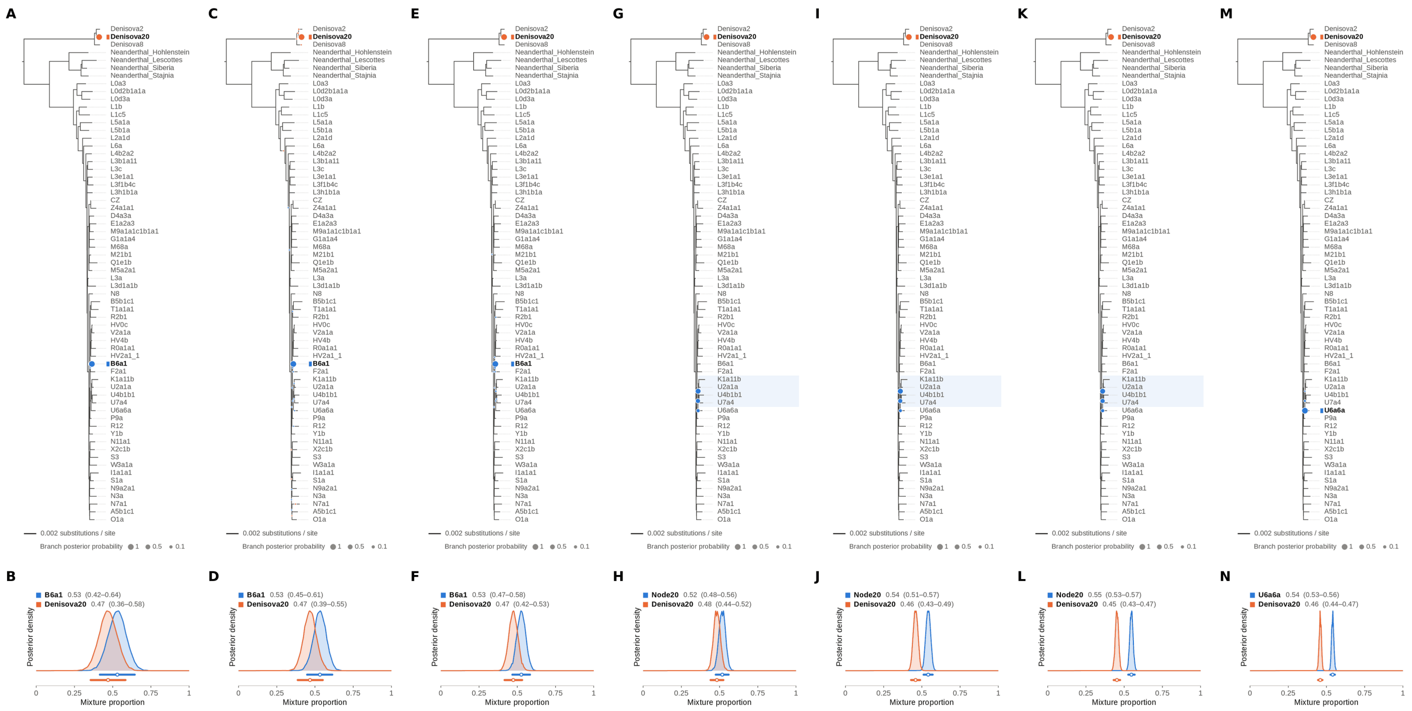


Figure S42: **Two-source simulated mixture of Denisova 21 (MT576653.1) (45% of reads) and modern human U5a2a (HM229344.1; 55%) at nominal mean coverage depths of 0.5X, 1X, 2X, 4X, 8X, 16X, 32X.** Both sources carry ancient DNA damage (U5a2a with the damage rates scaled by 0.75). For each coverage depth, the upper panel shows the phylogenetic placement posterior of both sources for the best of five chains ($k = 2$, 100,000 iterations each), colored by source (dot area: branch posterior probability; swelling: posterior density of the position along the branch; bold tips: highest-posterior branches), and the lower panel the posterior densities of the mixture proportions $\Theta$ with median and 95% credible interval. Denisova 21 is expected on the Denisova20 branch (its closest relative in the reference); U5a2a, for which no U5 haplotype is present in the reference, is expected within the U clade. Modal branch of each source in the best chain ($p$: its posterior probability) and $\Theta$ median [95% CI]: (A,B) 0.5X: B6a1 ($p = 1.00$), $\Theta = 0.53$ [0.42, 0.64]; Denisova20 ($p = 1.00$), $\Theta = 0.47$ [0.36, 0.58]; (C,D) 1X: B6a1 ($p = 1.00$), $\Theta = 0.53$ [0.45, 0.61]; Denisova20 ($p = 1.00$), $\Theta = 0.47$ [0.39, 0.55]; (E,F) 2X: B6a1 ($p = 0.99$), $\Theta = 0.53$ [0.47, 0.58]; Denisova20 ($p = 1.00$), $\Theta = 0.47$ [0.42, 0.53]; (G,H) 4X: Node20 (internal branch within the U clade; $p = 0.56$), $\Theta = 0.52$ [0.48, 0.56]; Denisova20 ($p = 1.00$), $\Theta = 0.48$ [0.44, 0.52]; (I,J) 8X: Node20 (internal branch within the U clade; $p = 0.61$), $\Theta = 0.54$ [0.51, 0.57]; Denisova20 ($p = 1.00$), $\Theta = 0.46$ [0.43, 0.49]; (K,L) 16X: Node20 (internal branch within the U clade; $p = 0.57$), $\Theta = 0.55$ [0.53, 0.57]; Denisova20 ($p = 1.00$), $\Theta = 0.45$ [0.43, 0.47]; (M,N) 32X: U6a6a ($p = 0.99$), $\Theta = 0.54$ [0.53, 0.56]; Denisova20 ($p = 1.00$), $\Theta = 0.46$ [0.44, 0.47].

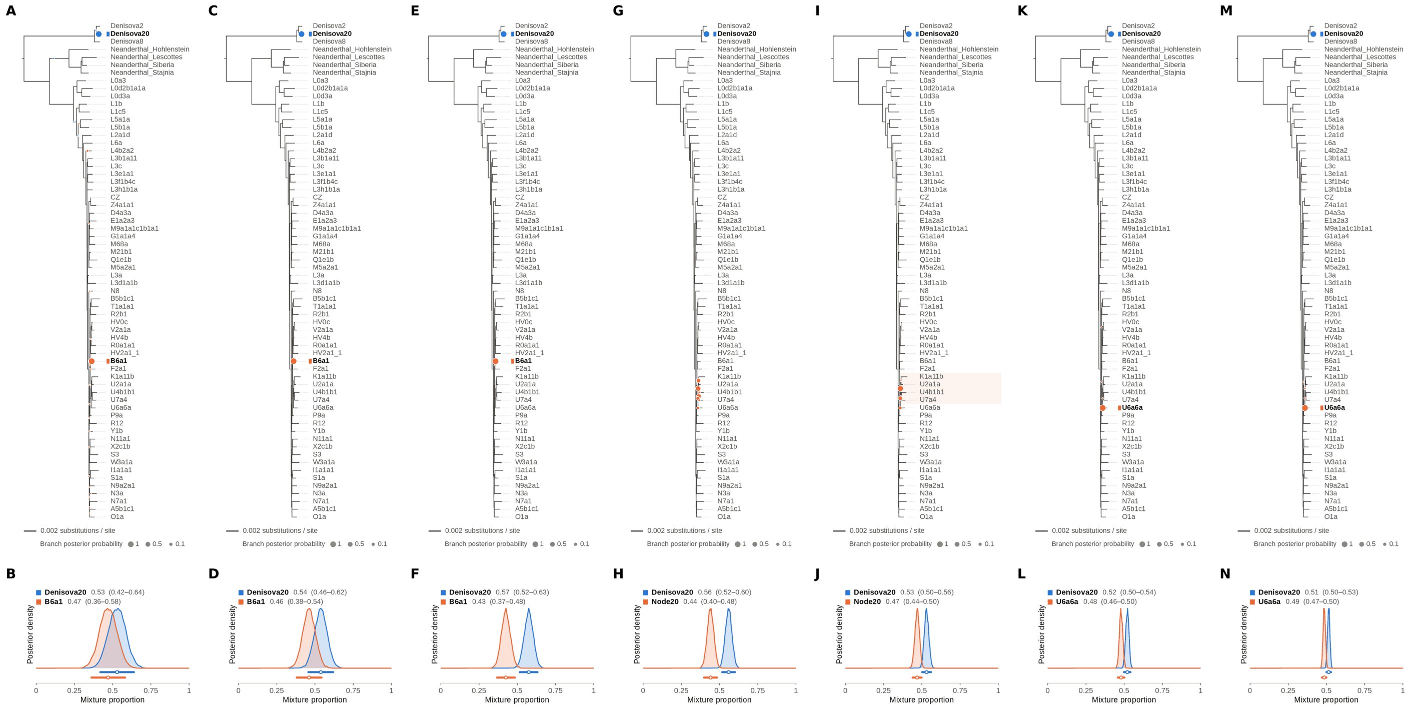

Figure S43: **Two-source simulated mixture of Denisova 21 (MT576653.1) (50% of reads) and modern human U5a2a (HM229344.1; 50%) at nominal mean coverage depths of 0.5X, 1X, 2X, 4X, 8X, 16X, 32X.** Both sources carry ancient DNA damage (U5a2a with the damage rates scaled by 0.75). For each coverage depth, the upper panel shows the phylogenetic placement posterior of both sources for the best of five chains ($k = 2$, 100,000 iterations each), colored by source (dot area: branch posterior probability; swelling: posterior density of the position along the branch; bold tips: highest-posterior branches), and the lower panel the posterior densities of the mixture proportions $\Theta$ with median and 95% credible interval. Denisova 21 is expected on the Denisova20 branch (its closest relative in the reference); U5a2a, for which no U5 haplotype is present in the reference, is expected within the U clade. Modal branch of each source in the best chain ($p$: its posterior probability) and $\Theta$ median [95% CI]: (A,B) 0.5X: Denisova20 ($p = 1.00$), $\Theta = 0.53$ [0.42, 0.64]; B6a1 ($p = 0.99$), $\Theta = 0.47$ [0.36, 0.58]; (C,D) 1X: Denisova20 ($p = 1.00$), $\Theta = 0.54$ [0.46, 0.62]; B6a1 ($p = 1.00$), $\Theta = 0.46$ [0.38, 0.54]; (E,F) 2X: Denisova20 ($p = 1.00$), $\Theta = 0.57$ [0.52, 0.63]; B6a1 ($p = 1.00$), $\Theta = 0.43$ [0.37, 0.48]; (G,H) 4X: Denisova20 ($p = 1.00$), $\Theta = 0.56$ [0.52, 0.60]; Node20 (internal branch within the U clade; $p = 0.40$), $\Theta = 0.44$ [0.40, 0.48]; (I,J) 8X: Denisova20 ($p = 1.00$), $\Theta = 0.53$ [0.50, 0.56]; Node20 (internal branch within the U clade; $p = 0.73$), $\Theta = 0.47$ [0.44, 0.50]; (K,L) 16X: Denisova20 ($p = 1.00$), $\Theta = 0.52$ [0.50, 0.54]; U6a6a ($p = 1.00$), $\Theta = 0.48$ [0.46, 0.50]; (M,N) 32X: Denisova20 ($p = 1.00$), $\Theta = 0.51$ [0.50, 0.53]; U6a6a ($p = 1.00$), $\Theta = 0.49$ [0.47, 0.50].

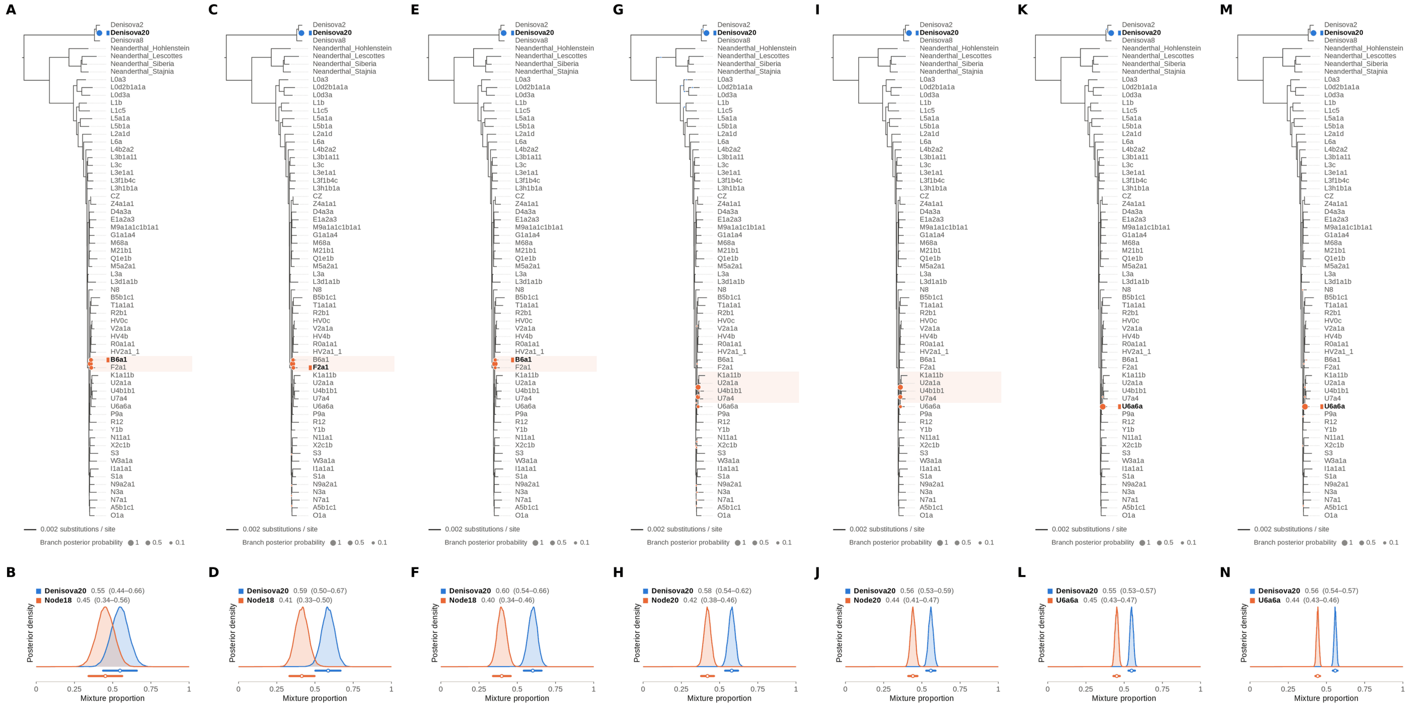


Figure S44: **Two-source simulated mixture of Denisova 21 (MT576653.1) (55% of reads) and modern human U5a2a (HM229344.1; 45%) at nominal mean coverage depths of 0.5X, 1X, 2X, 4X, 8X, 16X, 32X.** Both sources carry ancient DNA damage (U5a2a with the damage rates scaled by 0.75). For each coverage depth, the upper panel shows the phylogenetic placement posterior of both sources for the best of five chains ($k = 2$, 100,000 iterations each), colored by source (dot area: branch posterior probability; swelling: posterior density of the position along the branch; bold tips: highest-posterior branches), and the lower panel the posterior densities of the mixture proportions $\Theta$ with median and 95% credible interval. Denisova 21 is expected on the Denisova20 branch (its closest relative in the reference); U5a2a, for which no U5 haplotype is present in the reference, is expected within the U clade. Modal branch of each source in the best chain ($p$: its posterior probability) and $\Theta$ median [95% CI]: (A,B) 0.5X: Denisova20 ($p = 1.00$), $\Theta = 0.55$ [0.44, 0.66]; Node18 (internal branch ancestral to B6a1 and F2a1; $p = 0.56$), $\Theta = 0.45$ [0.34, 0.56]; (C,D) 1X: Denisova20 ($p = 1.00$), $\Theta = 0.59$ [0.50, 0.67]; Node18 (internal branch ancestral to B6a1 and F2a1; $p = 0.66$), $\Theta = 0.41$ [0.33, 0.50]; (E,F) 2X: Denisova20 ($p = 1.00$), $\Theta = 0.60$ [0.54, 0.66]; Node18 (internal branch ancestral to B6a1 and F2a1; $p = 0.79$), $\Theta = 0.40$ [0.34, 0.46]; (G,H) 4X: Denisova20 ($p = 1.00$), $\Theta = 0.58$ [0.54, 0.62]; Node20 (internal branch within the U clade; $p = 0.58$), $\Theta = 0.42$ [0.38, 0.46]; (I,J) 8X: Denisova20 ($p = 1.00$), $\Theta = 0.56$ [0.53, 0.59]; Node20 (internal branch within the U clade; $p = 0.61$), $\Theta = 0.44$ [0.41, 0.47]; (K,L) 16X: Denisova20 ($p = 1.00$), $\Theta = 0.55$ [0.53, 0.57]; U6a6a ($p = 1.00$), $\Theta = 0.45$ [0.43, 0.47]; (M,N) 32X: Denisova20 ($p = 1.00$), $\Theta = 0.56$ [0.54, 0.57]; U6a6a ($p = 1.00$), $\Theta = 0.44$ [0.43, 0.46].

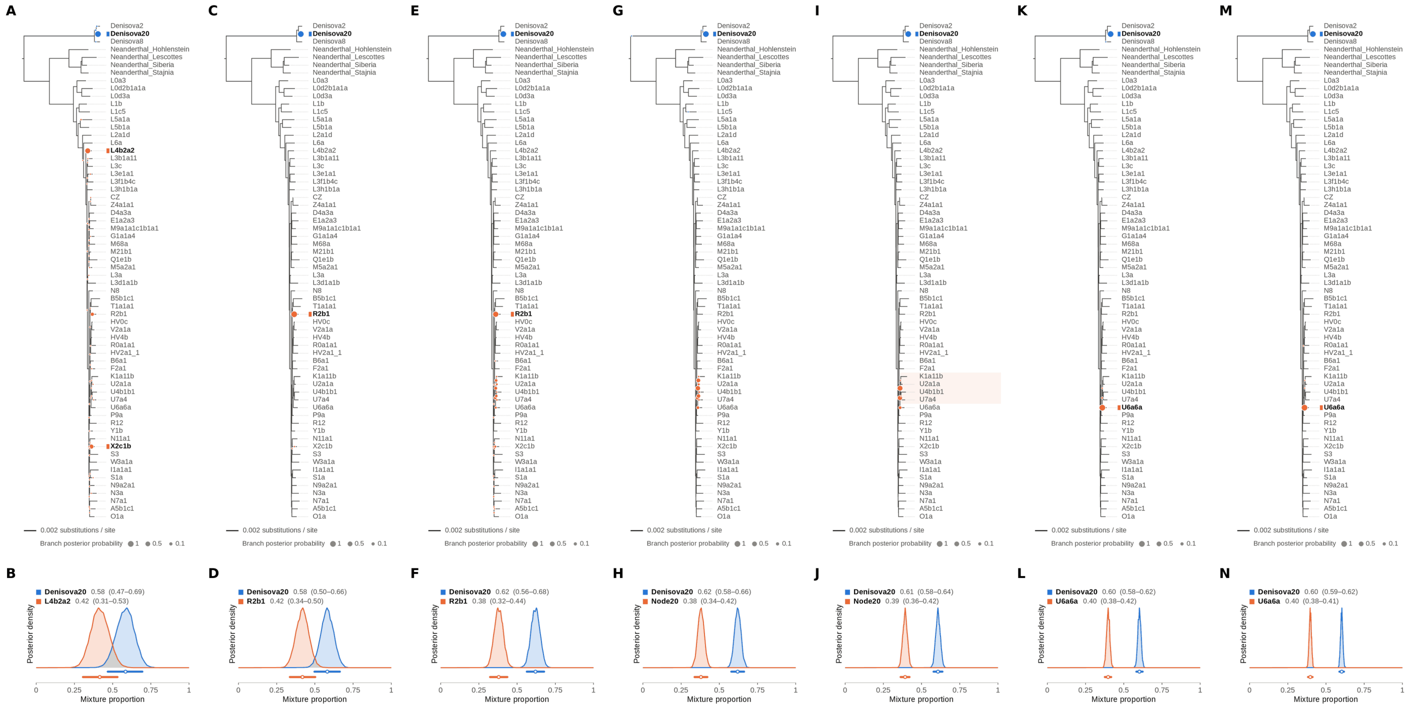


Figure S45: **Two-source simulated mixture of Denisova 21 (MT576653.1) (60% of reads) and modern human U5a2a (HM229344.1; 40%) at nominal mean coverage depths of 0.5X, 1X, 2X, 4X, 8X, 16X, 32X.** Both sources carry ancient DNA damage (U5a2a with the damage rates scaled by 0.75). For each coverage depth, the upper panel shows the phylogenetic placement posterior of both sources for the best of five chains ($k$ = 2, 100,000 iterations each), colored by source (dot area: branch posterior probability; swelling: posterior density of the position along the branch; bold tips: highest-posterior branches), and the lower panel the posterior densities of the mixture proportions $\Theta$ with median and 95% credible interval. Denisova 21 is expected on the Denisova20 branch (its closest relative in the reference); U5a2a, for which no U5 haplotype is present in the reference, is expected within the U clade. Modal branch of each source in the best chain ($p$: its posterior probability) and $\Theta$ median [95% CI]: (A,B) 0.5X: Denisova20 ($p$ = 1.00), $\Theta$ = 0.58 [0.47, 0.69]; L4b2a2 ($p$ = 0.60), $\Theta$ = 0.42 [0.31, 0.53]; (C,D) 1X: Denisova20 ($p$ = 1.00), $\Theta$ = 0.58 [0.50, 0.66]; R2b1 ($p$ = 1.00), $\Theta$ = 0.42 [0.34, 0.50]; (E,F) 2X: Denisova20 ($p$ = 1.00), $\Theta$ = 0.62 [0.56, 0.68]; R2b1 ($p$ = 0.71), $\Theta$ = 0.38 [0.32, 0.44]; (G,H) 4X: Denisova20 ($p$ = 1.00), $\Theta$ = 0.62 [0.58, 0.66]; Node20 (internal branch within the U clade; $p$ = 0.39), $\Theta$ = 0.38 [0.34, 0.42]; (I,J) 8X: Denisova20 ($p$ = 1.00), $\Theta$ = 0.61 [0.58, 0.64]; Node20 (internal branch within the U clade; $p$ = 0.57), $\Theta$ = 0.39 [0.36, 0.42]; (K,L) 16X: Denisova20 ($p$ = 1.00), $\Theta$ = 0.60 [0.58, 0.62]; U6a6a ($p$ = 0.99), $\Theta$ = 0.40 [0.38, 0.42]; (M,N) 32X: Denisova20 ($p$ = 1.00), $\Theta$ = 0.60 [0.59, 0.62]; U6a6a ($p$ = 1.00), $\Theta$ = 0.40 [0.38, 0.41].

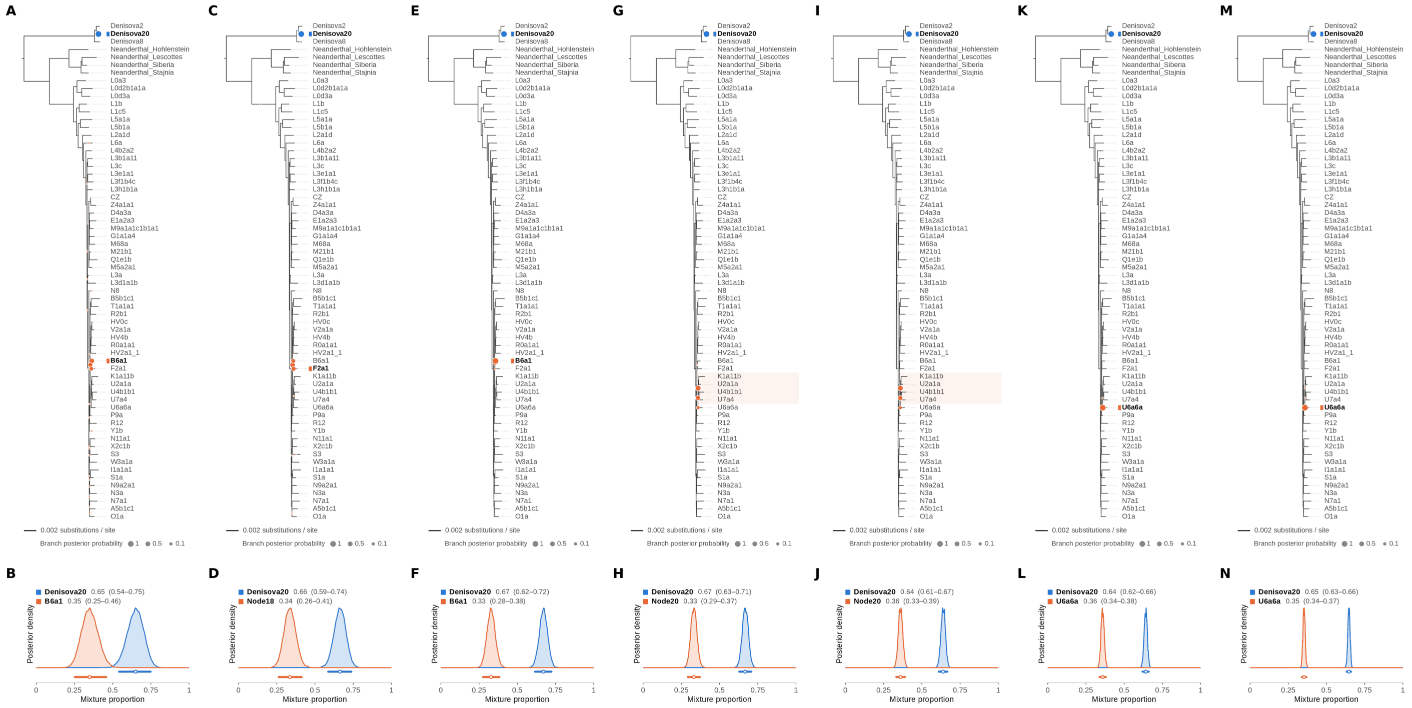


Figure S46: **Two-source simulated mixture of Denisova 21 (MT576653.1) (65% of reads) and modern human U5a2a (HM229344.1; 35%) at nominal mean coverage depths of 0.5X, 1X, 2X, 4X, 8X, 16X, 32X.** Both sources carry ancient DNA damage (U5a2a with the damage rates scaled by 0.75). For each coverage depth, the upper panel shows the phylogenetic placement posterior of both sources for the best of five chains ($k = 2$, 100,000 iterations each), colored by source (dot area: branch posterior probability; swelling: posterior density of the position along the branch; bold tips: highest-posterior branches), and the lower panel the posterior densities of the mixture proportions $\Theta$ with median and 95% credible interval. Denisova 21 is expected on the Denisova20 branch (its closest relative in the reference); U5a2a, for which no U5 haplotype is present in the reference, is expected within the U clade. Modal branch of each source in the best chain ($p$: its posterior probability) and $\Theta$ median [95% CI]: (A,B) 0.5X: Denisova20 ($p = 1.00$), $\Theta = 0.65$ [0.54, 0.75]; B6a1 ($p = 0.50$), $\Theta = 0.35$ [0.25, 0.46]; (C,D) 1X: Denisova20 ($p = 1.00$), $\Theta = 0.66$ [0.59, 0.74]; Node18 (internal branch ancestral to B6a1 and F2a1; $p = 0.48$), $\Theta = 0.34$ [0.26, 0.41]; (E,F) 2X: Denisova20 ($p = 1.00$), $\Theta = 0.67$ [0.62, 0.72]; B6a1 ($p = 1.00$), $\Theta = 0.33$ [0.28, 0.38]; (G,H) 4X: Denisova20 ($p = 1.00$), $\Theta = 0.67$ [0.63, 0.71]; Node20 (internal branch within the U clade; $p = 0.57$), $\Theta = 0.33$ [0.29, 0.37]; (I,J) 8X: Denisova20 ($p = 1.00$), $\Theta = 0.64$ [0.61, 0.67]; Node20 (internal branch within the U clade; $p = 0.57$), $\Theta = 0.36$ [0.33, 0.39]; (K,L) 16X: Denisova20 ($p = 1.00$), $\Theta = 0.64$ [0.62, 0.66]; U6a6a ($p = 1.00$), $\Theta = 0.36$ [0.34, 0.38]; (M,N) 32X: Denisova20 ($p = 1.00$), $\Theta = 0.65$ [0.63, 0.66]; U6a6a ($p = 1.00$), $\Theta = 0.35$ [0.34, 0.37].

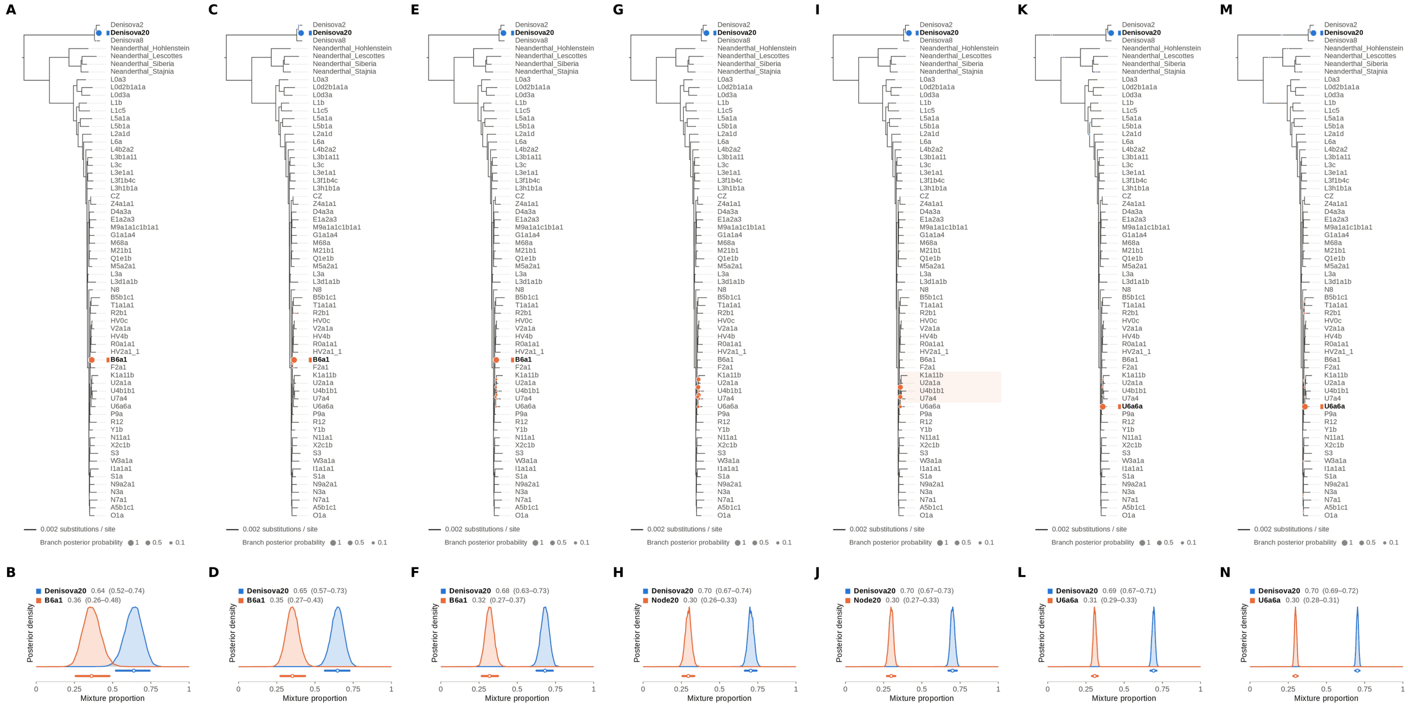


Figure S47: **Two-source simulated mixture of Denisova 21 (MT576653.1) (70% of reads) and modern human U5a2a (HM229344.1; 30%) at nominal mean coverage depths of 0.5X, 1X, 2X, 4X, 8X, 16X, 32X.** Both sources carry ancient DNA damage (U5a2a with the damage rates scaled by 0.75). For each coverage depth, the upper panel shows the phylogenetic placement posterior of both sources for the best of five chains ($k = 2$, 100,000 iterations each), colored by source (dot area: branch posterior probability; swelling: posterior density of the position along the branch; bold tips: highest-posterior branches), and the lower panel the posterior densities of the mixture proportions $\Theta$ with median and 95% credible interval. Denisova 21 is expected on the Denisova20 branch (its closest relative in the reference); U5a2a, for which no U5 haplotype is present in the reference, is expected within the U clade. Modal branch of each source in the best chain ($p$: its posterior probability) and $\Theta$ median [95% CI]: (A,B) 0.5X: Denisova20 ($p = 1.00$), $\Theta = 0.64$ [0.52, 0.74]; B6a1 ($p = 1.00$), $\Theta = 0.36$ [0.26, 0.48]; (C,D) 1X: Denisova20 ($p = 1.00$), $\Theta = 0.65$ [0.57, 0.73]; B6a1 ($p = 1.00$), $\Theta = 0.35$ [0.27, 0.43]; (E,F) 2X: Denisova20 ($p = 1.00$), $\Theta = 0.68$ [0.63, 0.73]; B6a1 ($p = 0.91$), $\Theta = 0.32$ [0.27, 0.37]; (G,H) 4X: Denisova20 ($p = 1.00$), $\Theta = 0.70$ [0.67, 0.74]; Node20 (internal branch within the U clade; $p = 0.33$), $\Theta = 0.30$ [0.26, 0.33]; (I,J) 8X: Denisova20 ($p = 1.00$), $\Theta = 0.70$ [0.67, 0.73]; Node20 (internal branch within the U clade; $p = 0.60$), $\Theta = 0.30$ [0.27, 0.33]; (K,L) 16X: Denisova20 ($p = 1.00$), $\Theta = 0.69$ [0.67, 0.71]; U6a6a ($p = 1.00$), $\Theta = 0.31$ [0.29, 0.33]; (M,N) 32X: Denisova20 ($p = 1.00$), $\Theta = 0.70$ [0.69, 0.72]; U6a6a ($p = 0.99$), $\Theta = 0.30$ [0.28, 0.31].

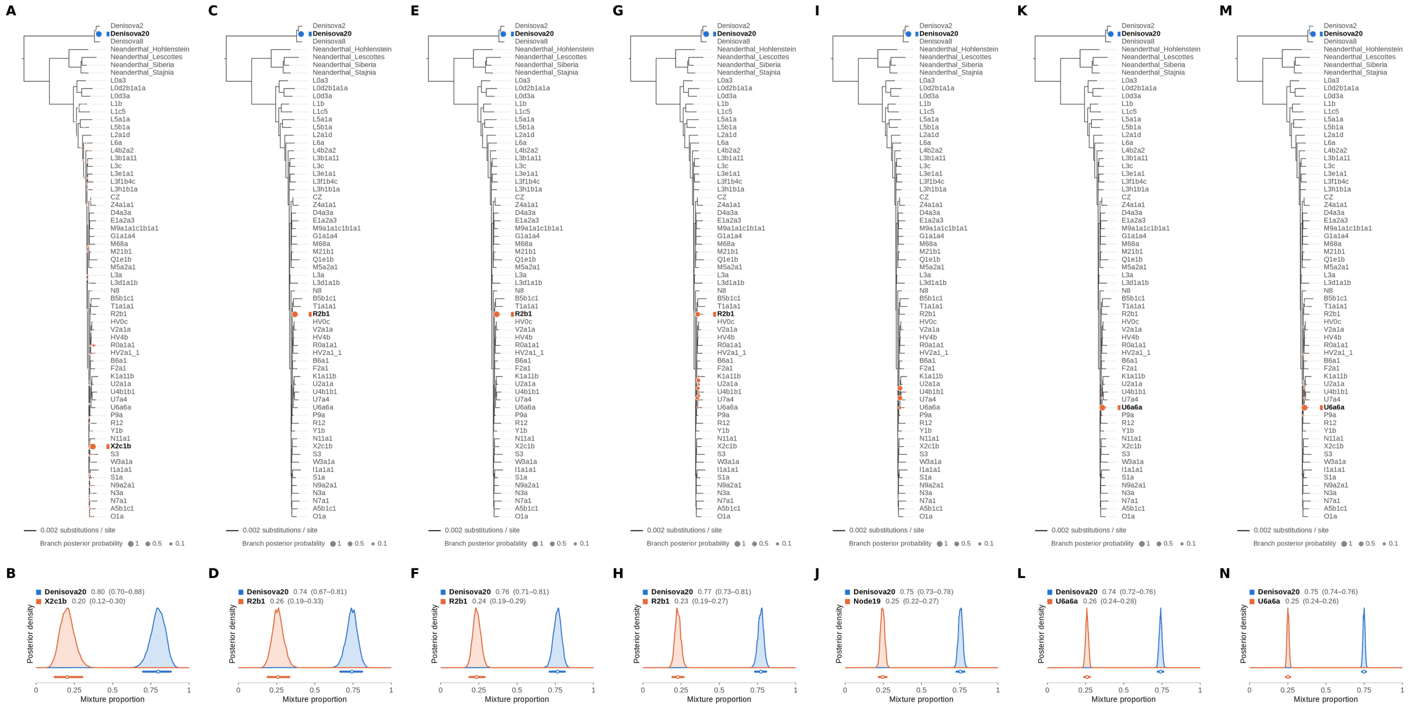


Figure S48: **Two-source simulated mixture of Denisova 21 (MT576653.1) (75% of reads) and modern human U5a2a (HM229344.1; 25%) at nominal mean coverage depths of 0.5X, 1X, 2X, 4X, 8X, 16X, 32X.** Both sources carry ancient DNA damage (U5a2a with the damage rates scaled by 0.75). For each coverage depth, the upper panel shows the phylogenetic placement posterior of both sources for the best of five chains ($k = 2$, 100,000 iterations each), colored by source (dot area: branch posterior probability; swelling: posterior density of the position along the branch; bold tips: highest-posterior branches), and the lower panel the posterior densities of the mixture proportions $\Theta$ with median and 95% credible interval. Denisova 21 is expected on the Denisova20 branch (its closest relative in the reference); U5a2a, for which no U5 haplotype is present in the reference, is expected within the U clade. Modal branch of each source in the best chain ($p$: its posterior probability) and $\Theta$ median [95% CI]: (A,B) 0.5X: Denisova20 ($p = 1.00$), $\Theta = 0.80$ [0.70, 0.88]; X2c1b ($p = 0.95$), $\Theta = 0.20$ [0.12, 0.30]; (C,D) 1X: Denisova20 ($p = 1.00$), $\Theta = 0.74$ [0.67, 0.81]; R2b1 ($p = 1.00$), $\Theta = 0.26$ [0.19, 0.33]; (E,F) 2X: Denisova20 ($p = 1.00$), $\Theta = 0.76$ [0.71, 0.81]; R2b1 ($p = 1.00$), $\Theta = 0.24$ [0.19, 0.29]; (G,H) 4X: Denisova20 ($p = 1.00$), $\Theta = 0.77$ [0.73, 0.81]; R2b1 ($p = 0.41$), $\Theta = 0.23$ [0.19, 0.27]; (I,J) 8X: Denisova20 ($p = 1.00$), $\Theta = 0.75$ [0.73, 0.78]; Node19 (stem of the U clade; $p = 0.48$), $\Theta = 0.25$ [0.22, 0.27]; (K,L) 16X: Denisova20 ($p = 1.00$), $\Theta = 0.74$ [0.72, 0.76]; U6a6a ($p = 1.00$), $\Theta = 0.26$ [0.24, 0.28]; (M,N) 32X: Denisova20 ($p = 1.00$), $\Theta = 0.75$ [0.74, 0.76]; U6a6a ($p = 0.99$), $\Theta = 0.25$ [0.24, 0.26].

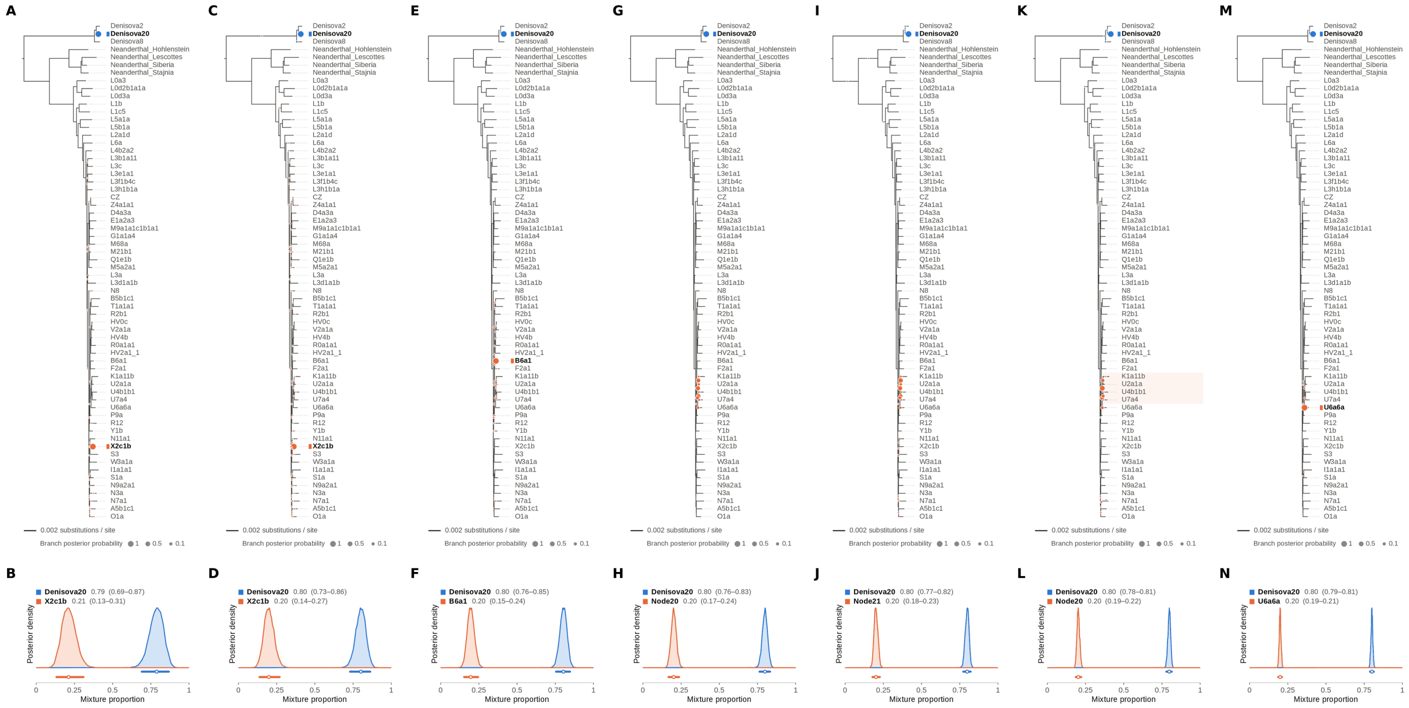


Figure S49: **Two-source simulated mixture of Denisova 21 (MT576653.1) (80% of reads) and modern human U5a2a (HM229344.1; 20%) at nominal mean coverage depths of 0.5X, 1X, 2X, 4X, 8X, 16X, 32X.** Both sources carry ancient DNA damage (U5a2a with the damage rates scaled by 0.75). For each coverage depth, the upper panel shows the phylogenetic placement posterior of both sources for the best of five chains ($k$ = 2, 100,000 iterations each), colored by source (dot area: branch posterior probability; swelling: posterior density of the position along the branch; bold tips: highest-posterior branches), and the lower panel the posterior densities of the mixture proportions Θ with median and 95% credible interval. Denisova 21 is expected on the Denisova20 branch (its closest relative in the reference); U5a2a, for which no U5 haplotype is present in the reference, is expected within the U clade. Modal branch of each source in the best chain ($p$: its posterior probability) and Θ median [95% CI]: (A,B) 0.5X: Denisova20 ($p$ = 1.00), Θ = 0.79 [0.69, 0.87]; X2c1b ($p$ = 0.99), Θ = 0.21 [0.13, 0.31]; (C,D) 1X: Denisova20 ($p$ = 1.00), Θ = 0.80 [0.73, 0.86]; X2c1b ($p$ = 0.98), Θ = 0.20 [0.14, 0.27]; (E,F) 2X: Denisova20 ($p$ = 1.00), Θ = 0.80 [0.76, 0.85]; B6a1 ($p$ = 0.96), Θ = 0.20 [0.15, 0.24]; (G,H) 4X: Denisova20 ($p$ = 1.00), Θ = 0.80 [0.76, 0.83]; Node20 (internal branch within the U clade; $p$ = 0.29), Θ = 0.20 [0.17, 0.24]; (I,J) 8X: Denisova20 ($p$ = 1.00), Θ = 0.80 [0.77, 0.82]; Node21 (internal branch ancestral to K1a11b and U2a1a; $p$ = 0.37), Θ = 0.20 [0.18, 0.23]; (K,L) 16X: Denisova20 ($p$ = 1.00), Θ = 0.80 [0.78, 0.81]; Node20 (internal branch within the U clade; $p$ = 0.53), Θ = 0.20 [0.19, 0.22]; (M,N) 32X: Denisova20 ($p$ = 1.00), Θ = 0.80 [0.79, 0.81]; U6a6a ($p$ = 0.99), Θ = 0.20 [0.19, 0.21].

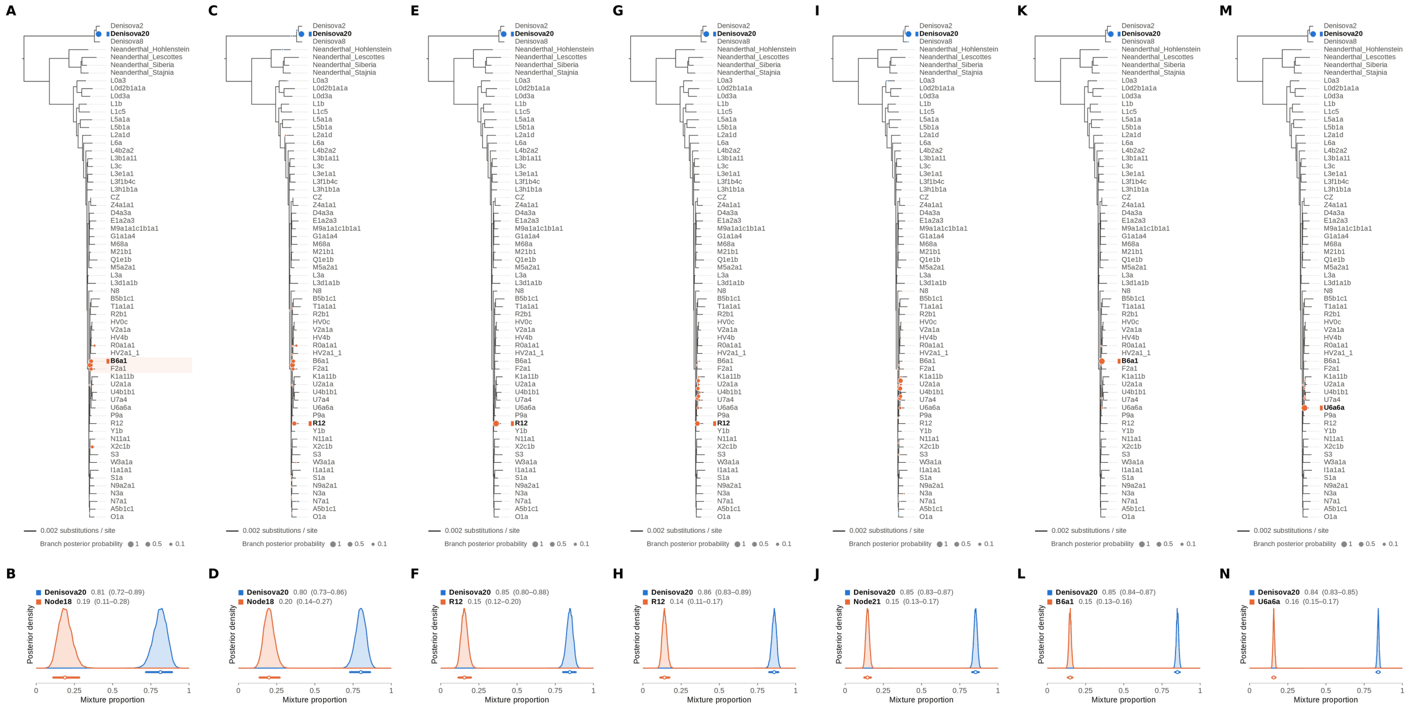


Figure S50: **Two-source simulated mixture of Denisova 21 (MT576653.1) (85% of reads) and modern human U5a2a (HM229344.1; 15%) at nominal mean coverage depths of 0.5X, 1X, 2X, 4X, 8X, 16X, 32X.** Both sources carry ancient DNA damage (U5a2a with the damage rates scaled by 0.75). For each coverage depth, the upper panel shows the phylogenetic placement posterior of both sources for the best of five chains ($k = 2$, 100,000 iterations each), colored by source (dot area: branch posterior probability; swelling: posterior density of the position along the branch; bold tips: highest-posterior branches), and the lower panel the posterior densities of the mixture proportions $\Theta$ with median and 95% credible interval. Denisova 21 is expected on the Denisova20 branch (its closest relative in the reference); U5a2a, for which no U5 haplotype is present in the reference, is expected within the U clade. Modal branch of each source in the best chain ($p$: its posterior probability) and $\Theta$ median [95% CI]: (A,B) 0.5X: Denisova20 ($p = 1.00$), $\Theta = 0.81$ [0.72, 0.89]; Node18 (internal branch ancestral to B6a1 and F2a1; $p = 0.55$), $\Theta = 0.19$ [0.11, 0.28]; (C,D) 1X: Denisova20 ($p = 1.00$), $\Theta = 0.80$ [0.73, 0.86]; Node18 (internal branch ancestral to B6a1 and F2a1; $p = 0.45$), $\Theta = 0.20$ [0.14, 0.27]; (E,F) 2X: Denisova20 ($p = 1.00$), $\Theta = 0.85$ [0.80, 0.88]; R12 ($p = 1.00$), $\Theta = 0.15$ [0.12, 0.20]; (G,H) 4X: Denisova20 ($p = 1.00$), $\Theta = 0.86$ [0.83, 0.89]; R12 ($p = 0.43$), $\Theta = 0.14$ [0.11, 0.17]; (I,J) 8X: Denisova20 ($p = 1.00$), $\Theta = 0.85$ [0.83, 0.87]; Node21 (internal branch ancestral to K1a11b and U2a1a; $p = 0.38$), $\Theta = 0.15$ [0.13, 0.17]; (K,L) 16X: Denisova20 ($p = 1.00$), $\Theta = 0.85$ [0.84, 0.87]; B6a1 ($p = 1.00$), $\Theta = 0.15$ [0.13, 0.16]; (M,N) 32X: Denisova20 ($p = 1.00$), $\Theta = 0.84$ [0.83, 0.85]; U6a6a ($p = 0.99$), $\Theta = 0.16$ [0.15, 0.17].

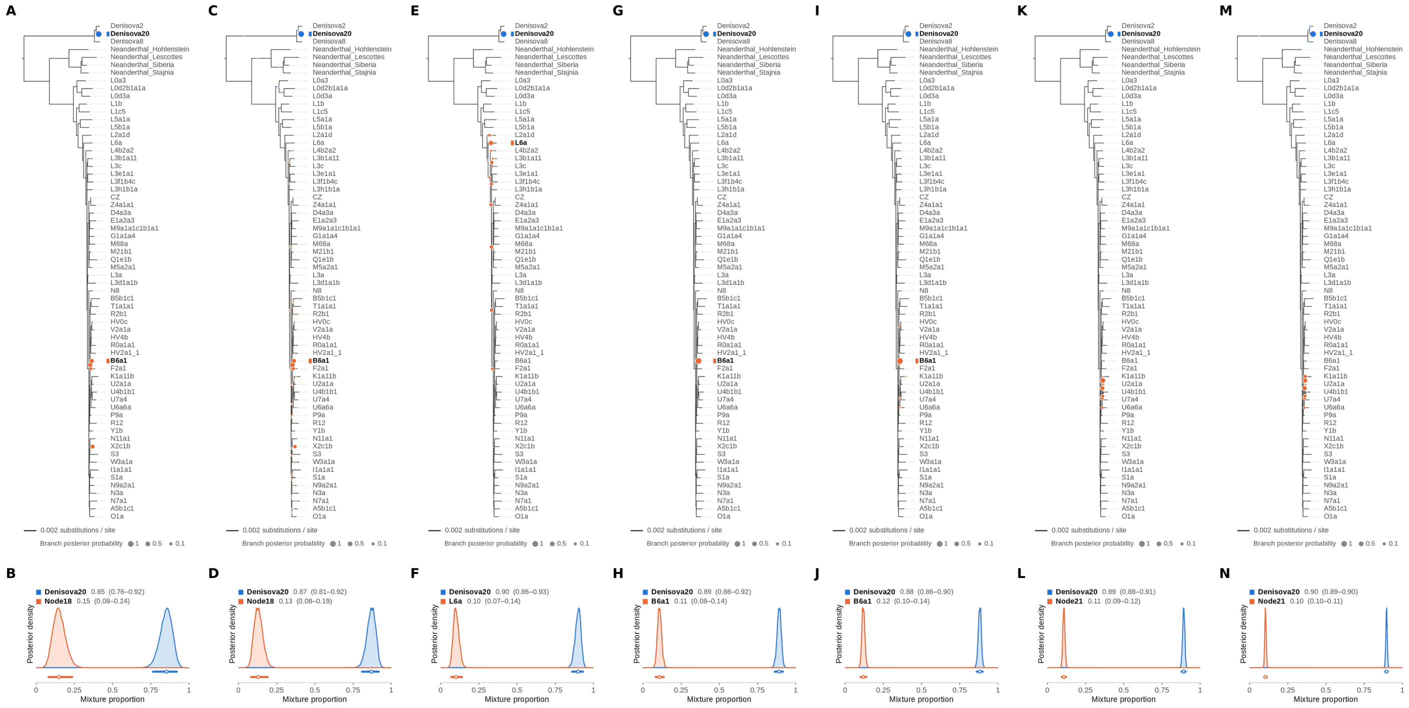

Figure S51: **Two-source simulated mixture of Denisova 21 (MT576653.1) (90% of reads) and modern human U5a2a (HM229344.1; 10%) at nominal mean coverage depths of 0.5X, 1X, 2X, 4X, 8X, 16X, 32X.** Both sources carry ancient DNA damage (U5a2a with the damage rates scaled by 0.75). For each coverage depth, the upper panel shows the phylogenetic placement posterior of both sources for the best of five chains ($k = 2$, 100,000 iterations each), colored by source (dot area: branch posterior probability; swelling: posterior density of the position along the branch; bold tips: highest-posterior branches), and the lower panel the posterior densities of the mixture proportions $\Theta$ with median and 95% credible interval. Denisova 21 is expected on the Denisova20 branch (its closest relative in the reference); U5a2a, for which no U5 haplotype is present in the reference, is expected within the U clade. Modal branch of each source in the best chain ($p$: its posterior probability) and $\Theta$ median [95% CI]: (A,B) 0.5X: Denisova20 ($p = 1.00$), $\Theta = 0.85$ [0.76, 0.92]; Node18 (internal branch ancestral to B6a1 and F2a1; $p = 0.37$), $\Theta = 0.15$ [0.08, 0.24]; (C,D) 1X: Denisova20 ($p = 1.00$), $\Theta = 0.87$ [0.81, 0.92]; Node18 (internal branch ancestral to B6a1 and F2a1; $p = 0.47$), $\Theta = 0.13$ [0.08, 0.19]; (E,F) 2X: Denisova20 ($p = 1.00$), $\Theta = 0.90$ [0.86, 0.93]; L6a ($p = 0.29$), $\Theta = 0.10$ [0.07, 0.14]; (G,H) 4X: Denisova20 ($p = 1.00$), $\Theta = 0.89$ [0.86, 0.92]; B6a1 ($p = 1.00$), $\Theta = 0.11$ [0.08, 0.14]; (I,J) 8X: Denisova20 ($p = 1.00$), $\Theta = 0.88$ [0.86, 0.90]; B6a1 ($p = 0.99$), $\Theta = 0.12$ [0.10, 0.14]; (K,L) 16X: Denisova20 ($p = 1.00$), $\Theta = 0.89$ [0.88, 0.91]; Node21 (internal branch ancestral to K1a11b and U2a1a; $p = 0.47$), $\Theta = 0.11$ [0.09, 0.12]; (M,N) 32X: Denisova20 ($p = 1.00$), $\Theta = 0.90$ [0.89, 0.90]; Node21 (internal branch ancestral to K1a11b and U2a1a; $p = 0.36$), $\Theta = 0.10$ [0.10, 0.11].

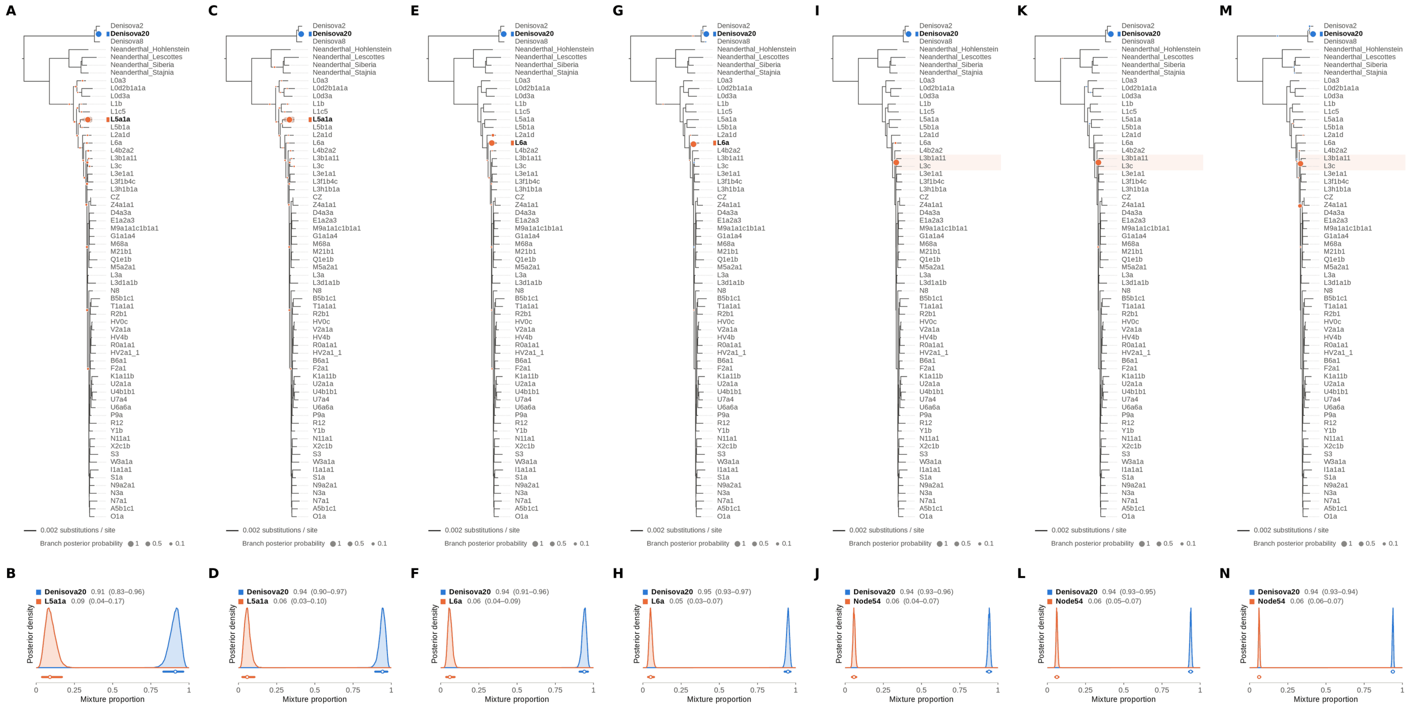


Figure S52: **Two-source simulated mixture of Denisova 21 (MT576653.1) (95% of reads) and modern human U5a2a (HM229344.1; 5%) at nominal mean coverage depths of 0.5X, 1X, 2X, 4X, 8X, 16X, 32X.** Both sources carry ancient DNA damage (U5a2a with the damage rates scaled by 0.75). For each coverage depth, the upper panel shows the phylogenetic placement posterior of both sources for the best of five chains ($k = 2$, 100,000 iterations each), colored by source (dot area: branch posterior probability; swelling: posterior density of the position along the branch; bold tips: highest-posterior branches), and the lower panel the posterior densities of the mixture proportions $\Theta$ with median and 95% credible interval. Denisova 21 is expected on the Denisova20 branch (its closest relative in the reference); U5a2a, for which no U5 haplotype is present in the reference, is expected within the U clade. Modal branch of each source in the best chain ($p$: its posterior probability) and $\Theta$ median [95% CI]: (A,B) 0.5X: Denisova20 ($p = 1.00$), $\Theta = 0.91$ [0.83, 0.96]; L5a1a ($p = 0.83$), $\Theta = 0.09$ [0.04, 0.17]; (C,D) 1X: Denisova20 ($p = 1.00$), $\Theta = 0.94$ [0.90, 0.97]; L5a1a ($p = 0.95$), $\Theta = 0.06$ [0.03, 0.10]; (E,F) 2X: Denisova20 ($p = 1.00$), $\Theta = 0.94$ [0.91, 0.96]; L6a ($p = 0.95$), $\Theta = 0.06$ [0.04, 0.09]; (G,H) 4X: Denisova20 ($p = 1.00$), $\Theta = 0.95$ [0.93, 0.97]; L6a ($p = 1.00$), $\Theta = 0.05$ [0.03, 0.07]; (I,J) 8X: Denisova20 ($p = 1.00$), $\Theta = 0.94$ [0.93, 0.96]; Node54 (internal branch ancestral to L3b1a11 and L3c; $p = 0.99$), $\Theta = 0.06$ [0.04, 0.07]; (K,L) 16X: Denisova20 ($p = 1.00$), $\Theta = 0.94$ [0.93, 0.95]; Node54 (internal branch ancestral to L3b1a11 and L3c; $p = 1.00$), $\Theta = 0.06$ [0.05, 0.07]; (M,N) 32X: Denisova20 ($p = 1.00$), $\Theta = 0.94$ [0.93, 0.94]; Node54 (internal branch ancestral to L3b1a11 and L3c; $p = 0.82$), $\Theta = 0.06$ [0.06, 0.07].

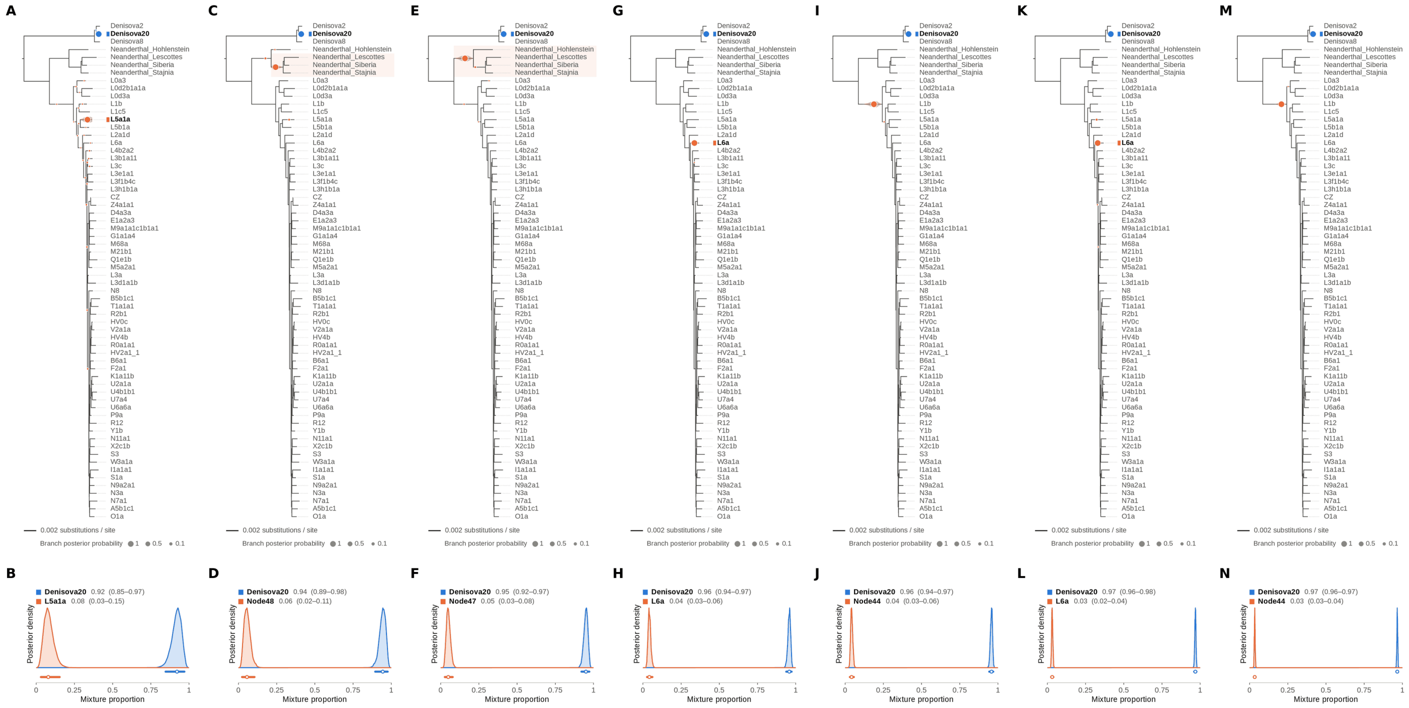


Figure S53: **Two-source simulated mixture of Denisova 21 (MT576653.1) (99% of reads) and modern human U5a2a (HM229344.1; 1%) at nominal mean coverage depths of 0.5X, 1X, 2X, 4X, 8X, 16X, 32X.** Both sources carry ancient DNA damage (U5a2a with the damage rates scaled by 0.75). For each coverage depth, the upper panel shows the phylogenetic placement posterior of both sources for the best of five chains ($k = 2$, 100,000 iterations each), colored by source (dot area: branch posterior probability; swelling: posterior density of the position along the branch; bold tips: highest-posterior branches), and the lower panel the posterior densities of the mixture proportions $\Theta$ with median and 95% credible interval. Denisova 21 is expected on the Denisova20 branch (its closest relative in the reference); U5a2a, for which no U5 haplotype is present in the reference, is expected within the U clade. Modal branch of each source in the best chain ($p$: its posterior probability) and $\Theta$ median [95% CI]: (A,B) 0.5X: Denisova20 ($p = 1.00$), $\Theta = 0.92$ [0.85, 0.97]; L5a1a ($p = 0.96$), $\Theta = 0.08$ [0.03, 0.15]; (C,D) 1X: Denisova20 ($p = 1.00$), $\Theta = 0.94$ [0.89, 0.98]; Node48 (internal branch ancestral to Neanderthal_Lescottes, Neanderthal_Siberia and Neanderthal_Stajnia; $p = 0.98$), $\Theta = 0.06$ [0.02, 0.11]; (E,F) 2X: Denisova20 ($p = 1.00$), $\Theta = 0.95$ [0.92, 0.97]; Node47 (stem of the Neanderthal clade; $p = 0.99$), $\Theta = 0.05$ [0.03, 0.08]; (G,H) 4X: Denisova20 ($p = 1.00$), $\Theta = 0.96$ [0.94, 0.97]; L6a ($p = 1.00$), $\Theta = 0.04$ [0.03, 0.06]; (I,J) 8X: Denisova20 ($p = 1.00$), $\Theta = 0.96$ [0.94, 0.97]; Node44 (stem of the archaic hominin clade; $p = 1.00$), $\Theta = 0.04$ [0.03, 0.06]; (K,L) 16X: Denisova20 ($p = 1.00$), $\Theta = 0.97$ [0.96, 0.98]; L6a ($p = 0.99$), $\Theta = 0.03$ [0.02, 0.04]; (M,N) 32X: Denisova20 ($p = 1.00$), $\Theta = 0.97$ [0.96, 0.97]; Node44 (stem of the archaic hominin clade; $p = 1.00$), $\Theta = 0.03$ [0.03, 0.04].

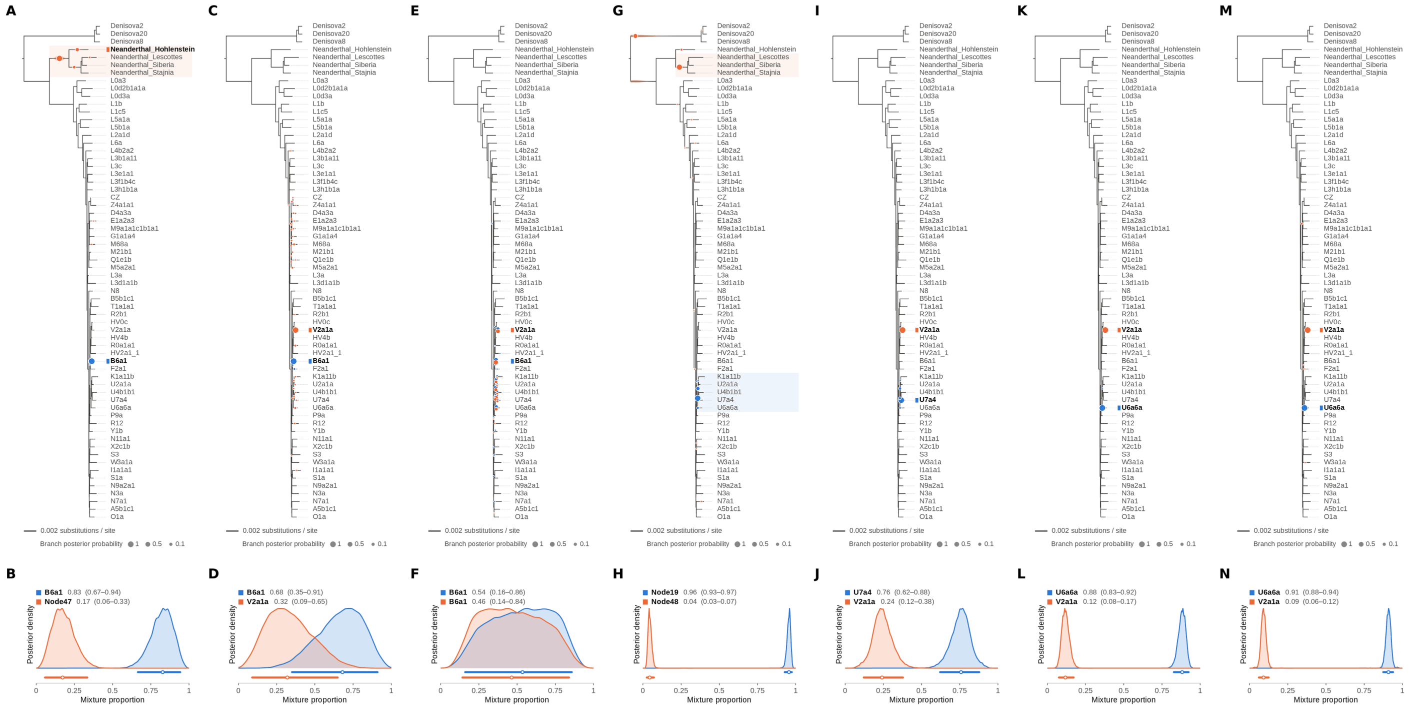

Figure S54: **Two-source simulated mixture of modern human D4a1d (FJ951517.1) (1% of reads) and modern human U5a2a (HM229344.1; 99%) at nominal mean coverage depths of 0.5X, 1X, 2X, 4X, 8X, 16X, 32X.** Both sources carry ancient DNA damage (U5a2a with the damage rates scaled by 0.75). For each coverage depth, the upper panel shows the phylogenetic placement posterior of both sources for the best of five chains ($k$ = 2, 100,000 iterations each), colored by source (dot area: branch posterior probability; swelling: posterior density of the position along the branch; bold tips: highest-posterior branches), and the lower panel the posterior densities of the mixture proportions $\Theta$ with median and 95% credible interval. D4a1d is expected on the D4a3a branch (its closest relative in the reference); U5a2a, for which no U5 haplotype is present in the reference, is expected within the U clade. Modal branch of each source in the best chain ($p$: its posterior probability) and $\Theta$ median [95% CI]: (A,B) 0.5X: B6a1 ($p$ = 1.00), $\Theta$ = 0.83 [0.67, 0.94]; Node47 (stem of the Neanderthal clade; $p$ = 0.83), $\Theta$ = 0.17 [0.06, 0.33]; (C,D) 1X: B6a1 ($p$ = 0.99), $\Theta$ = 0.68 [0.35, 0.91]; V2a1a ($p$ = 0.88), $\Theta$ = 0.32 [0.09, 0.65]; (E,F) 2X: B6a1 ($p$ = 0.28), $\Theta$ = 0.54 [0.16, 0.86]; B6a1 ($p$ = 0.39), $\Theta$ = 0.46 [0.14, 0.84]; (G,H) 4X: Node19 (stem of the U clade; $p$ = 0.85), $\Theta$ = 0.96 [0.93, 0.97]; Node48 (internal branch ancestral to Neanderthal_Lescottes, Neanderthal_Siberia and Neanderthal_Stajnia; $p$ = 0.71), $\Theta$ = 0.04 [0.03, 0.07]; (I,J) 8X: U7a4 ($p$ = 0.94), $\Theta$ = 0.76 [0.62, 0.88]; V2a1a ($p$ = 1.00), $\Theta$ = 0.24 [0.12, 0.38]; (K,L) 16X: U6a6a ($p$ = 1.00), $\Theta$ = 0.88 [0.83, 0.92]; V2a1a ($p$ = 1.00), $\Theta$ = 0.12 [0.08, 0.17]; (M,N) 32X: U6a6a ($p$ = 1.00), $\Theta$ = 0.91 [0.88, 0.94]; V2a1a ($p$ = 0.99), $\Theta$ = 0.09 [0.06, 0.12].

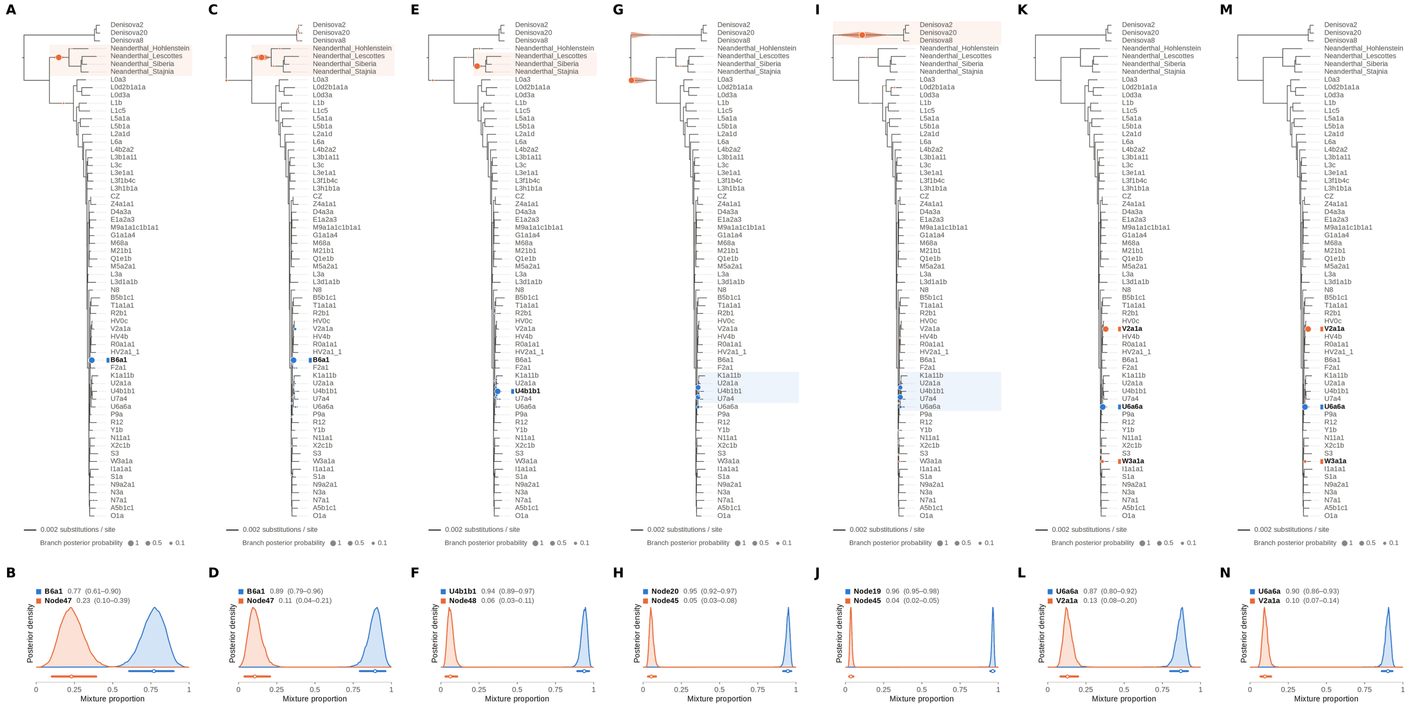


Figure S55: **Two-source simulated mixture of modern human D4a1d (FJ951517.1) (5% of reads) and modern human U5a2a (HM229344.1; 95%) at nominal mean coverage depths of 0.5X, 1X, 2X, 4X, 8X, 16X, 32X.** Both sources carry ancient DNA damage (U5a2a with the damage rates scaled by 0.75). For each coverage depth, the upper panel shows the phylogenetic placement posterior of both sources for the best of five chains ($k = 2$, 100,000 iterations each), colored by source (dot area: branch posterior probability; swelling: posterior density of the position along the branch; bold tips: highest-posterior branches), and the lower panel the posterior densities of the mixture proportions $\Theta$ with median and 95% credible interval. D4a1d is expected on the D4a3a branch (its closest relative in the reference); U5a2a, for which no U5 haplotype is present in the reference, is expected within the U clade. Modal branch of each source in the best chain ($p$: its posterior probability) and $\Theta$ median [95% CI]: (A,B) 0.5X: B6a1 ($p = 1.00$), $\Theta = 0.77$ [0.61, 0.90]; Node47 (stem of the Neanderthal clade; $p = 1.00$), $\Theta = 0.23$ [0.10, 0.39]; (C,D) 1X: B6a1 ($p = 0.96$), $\Theta = 0.89$ [0.79, 0.96]; Node47 (stem of the Neanderthal clade; $p = 0.97$), $\Theta = 0.11$ [0.04, 0.21]; (E,F) 2X: U4b1b1 ($p = 0.96$), $\Theta = 0.94$ [0.89, 0.97]; Node48 (internal branch ancestral to Neanderthal_Lescottes, Neanderthal_Siberia and Neanderthal_Stajnia; $p = 0.98$), $\Theta = 0.06$ [0.03, 0.11]; (G,H) 4X: Node20 (internal branch within the U clade; $p = 0.56$), $\Theta = 0.95$ [0.92, 0.97]; Node45 (stem of the Denisovan clade; $p = 1.00$), $\Theta = 0.05$ [0.03, 0.08]; (I,J) 8X: Node19 (stem of the U clade; $p = 0.66$), $\Theta = 0.96$ [0.95, 0.98]; Node45 (stem of the Denisovan clade; $p = 0.99$), $\Theta = 0.04$ [0.02, 0.05]; (K,L) 16X: U6a6a ($p = 1.00$), $\Theta = 0.87$ [0.80, 0.92]; V2a1a ($p = 0.91$), $\Theta = 0.13$ [0.08, 0.20]; (M,N) 32X: U6a6a ($p = 1.00$), $\Theta = 0.90$ [0.86, 0.93]; V2a1a ($p = 0.94$), $\Theta = 0.10$ [0.07, 0.14].

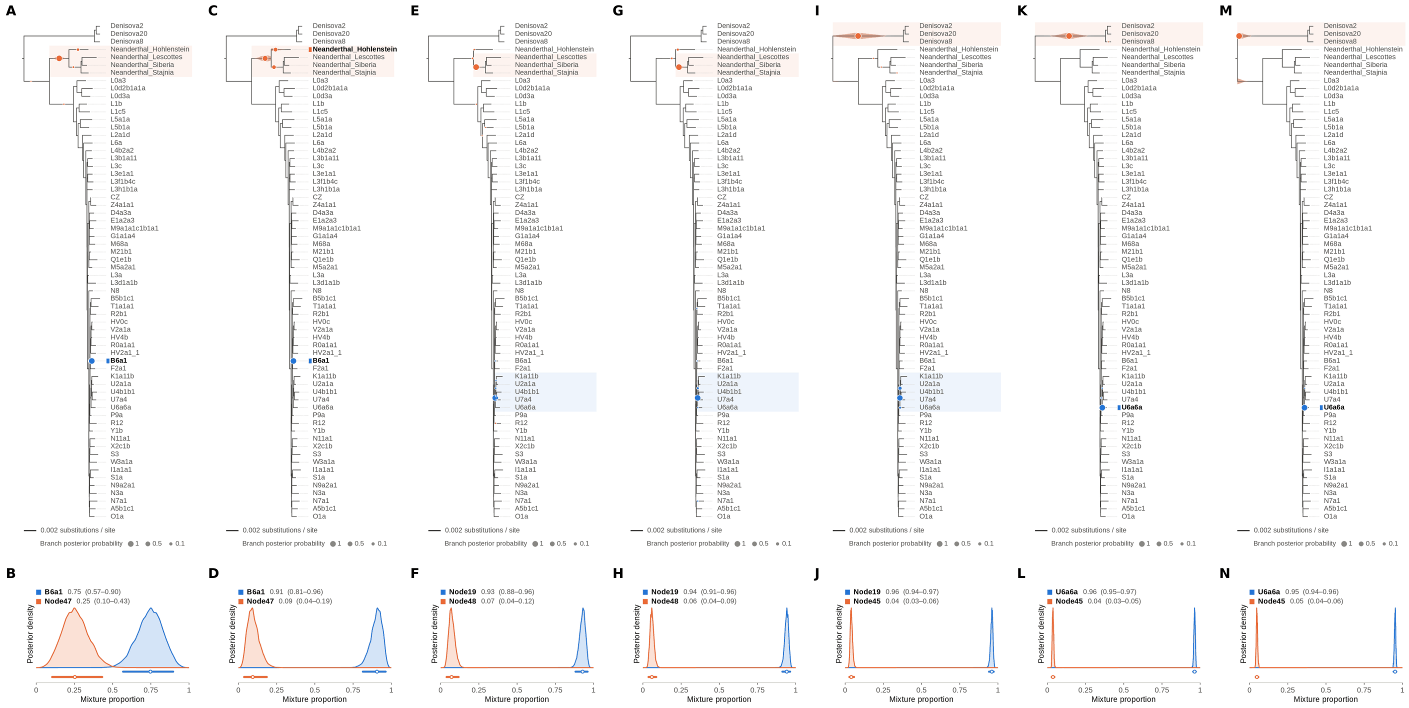


Figure S56: **Two-source simulated mixture of modern human D4a1d (FJ951517.1) (10% of reads) and modern human U5a2a (HM229344.1; 90%) at nominal mean coverage depths of 0.5X, 1X, 2X, 4X, 8X, 16X, 32X.** Both sources carry ancient DNA damage (U5a2a with the damage rates scaled by 0.75). For each coverage depth, the upper panel shows the phylogenetic placement posterior of both sources for the best of five chains ($k = 2$, 100,000 iterations each), colored by source (dot area: branch posterior probability; swelling: posterior density of the position along the branch; bold tips: highest-posterior branches), and the lower panel the posterior densities of the mixture proportions $\Theta$ with median and 95% credible interval. D4a1d is expected on the D4a3a branch (its closest relative in the reference); U5a2a, for which no U5 haplotype is present in the reference, is expected within the U clade. Modal branch of each source in the best chain ($p$: its posterior probability) and $\Theta$ median [95% CI]: (A,B) 0.5X: B6a1 ($p = 1.00$), $\Theta = 0.75$ [0.57, 0.90]; Node47 (stem of the Neanderthal clade; $p = 0.96$), $\Theta = 0.25$ [0.10, 0.43]; (C,D) 1X: B6a1 ($p = 1.00$), $\Theta = 0.91$ [0.81, 0.96]; Node47 (stem of the Neanderthal clade; $p = 0.62$), $\Theta = 0.09$ [0.04, 0.19]; (E,F) 2X: Node19 (stem of the U clade; $p = 0.99$), $\Theta = 0.93$ [0.88, 0.96]; Node48 (internal branch ancestral to Neanderthal_Lescottes, Neanderthal_Siberia and Neanderthal_Stajnia; $p = 0.99$), $\Theta = 0.07$ [0.04, 0.12]; (G,H) 4X: Node19 (stem of the U clade; $p = 0.98$), $\Theta = 0.94$ [0.91, 0.96]; Node48 (internal branch ancestral to Neanderthal_Lescottes, Neanderthal_Siberia and Neanderthal_Stajnia; $p = 0.98$), $\Theta = 0.06$ [0.04, 0.09]; (I,J) 8X: Node19 (stem of the U clade; $p = 0.87$), $\Theta = 0.96$ [0.94, 0.97]; Node45 (stem of the Denisovan clade; $p = 1.00$), $\Theta = 0.04$ [0.03, 0.06]; (K,L) 16X: U6a6a ($p = 1.00$), $\Theta = 0.96$ [0.95, 0.97]; Node45 (stem of the Denisovan clade; $p = 1.00$), $\Theta = 0.04$ [0.03, 0.05]; (M,N) 32X: U6a6a ($p = 1.00$), $\Theta = 0.95$ [0.94, 0.96]; Node45 (stem of the Denisovan clade; $p = 1.00$), $\Theta = 0.05$ [0.04, 0.06].

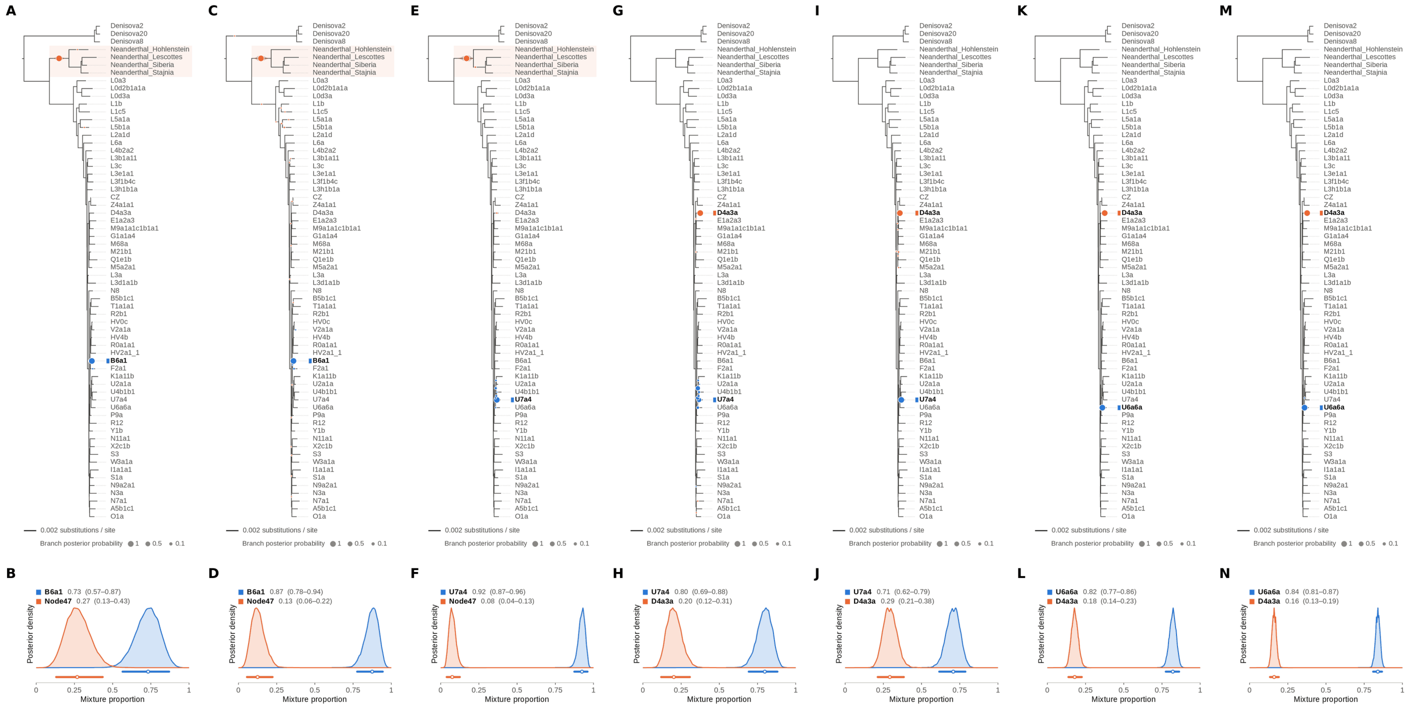


Figure S57: **Two-source simulated mixture of modern human D4a1d (FJ951517.1) (15% of reads) and modern human U5a2a (HM229344.1; 85%) at nominal mean coverage depths of 0.5X, 1X, 2X, 4X, 8X, 16X, 32X.** Both sources carry ancient DNA damage (U5a2a with the damage rates scaled by 0.75). For each coverage depth, the upper panel shows the phylogenetic placement posterior of both sources for the best of five chains ($k = 2$, 100,000 iterations each), colored by source (dot area: branch posterior probability; swelling: posterior density of the position along the branch; bold tips: highest-posterior branches), and the lower panel the posterior densities of the mixture proportions $\Theta$ with median and 95% credible interval. D4a1d is expected on the D4a3a branch (its closest relative in the reference); U5a2a, for which no U5 haplotype is present in the reference, is expected within the U clade. Modal branch of each source in the best chain ($p$: its posterior probability) and $\Theta$ median [95% CI]: (A,B) 0.5X: B6a1 ($p = 1.00$), $\Theta = 0.73$ [0.57, 0.87]; Node47 (stem of the Neanderthal clade; $p = 1.00$), $\Theta = 0.27$ [0.13, 0.43]; (C,D) 1X: B6a1 ($p = 0.99$), $\Theta = 0.87$ [0.78, 0.94]; Node47 (stem of the Neanderthal clade; $p = 0.99$), $\Theta = 0.13$ [0.06, 0.22]; (E,F) 2X: U7a4 ($p = 0.94$), $\Theta = 0.92$ [0.87, 0.96]; Node47 (stem of the Neanderthal clade; $p = 1.00$), $\Theta = 0.08$ [0.04, 0.13]; (G,H) 4X: U7a4 ($p = 0.41$), $\Theta = 0.80$ [0.69, 0.88]; D4a3a ($p = 1.00$), $\Theta = 0.20$ [0.12, 0.31]; (I,J) 8X: U7a4 ($p = 1.00$), $\Theta = 0.71$ [0.62, 0.79]; D4a3a ($p = 1.00$), $\Theta = 0.29$ [0.21, 0.38]; (K,L) 16X: U6a6a ($p = 1.00$), $\Theta = 0.82$ [0.77, 0.86]; D4a3a ($p = 1.00$), $\Theta = 0.18$ [0.14, 0.23]; (M,N) 32X: U6a6a ($p = 1.00$), $\Theta = 0.84$ [0.81, 0.87]; D4a3a ($p = 1.00$), $\Theta = 0.16$ [0.13, 0.19].

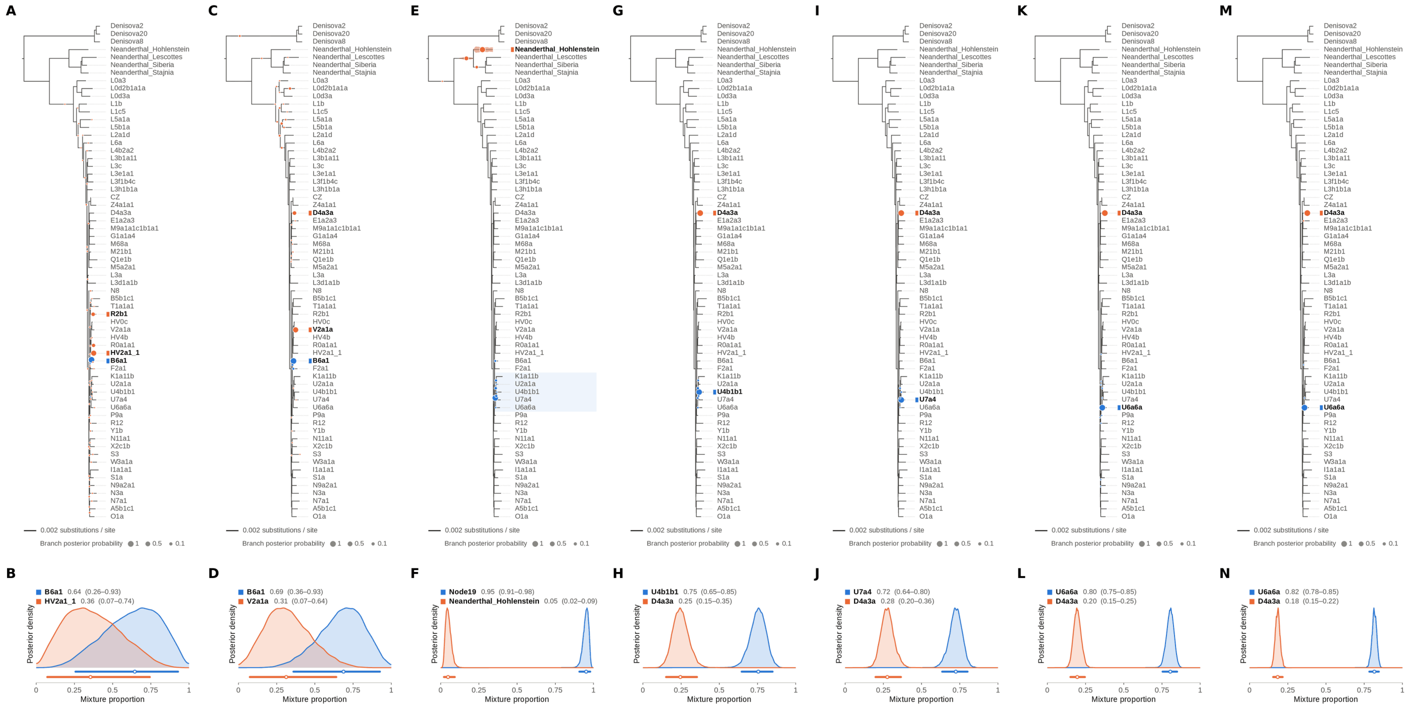


Figure S58: **Two-source simulated mixture of modern human D4a1d (FJ951517.1) (20% of reads) and modern human U5a2a (HM229344.1; 80%) at nominal mean coverage depths of 0.5X, 1X, 2X, 4X, 8X, 16X, 32X.** Both sources carry ancient DNA damage (U5a2a with the damage rates scaled by 0.75). For each coverage depth, the upper panel shows the phylogenetic placement posterior of both sources for the best of five chains ($k = 2$, 100,000 iterations each), colored by source (dot area: branch posterior probability; swelling: posterior density of the position along the branch; bold tips: highest-posterior branches), and the lower panel the posterior densities of the mixture proportions $\Theta$ with median and 95% credible interval. D4a1d is expected on the D4a3a branch (its closest relative in the reference); U5a2a, for which no U5 haplotype is present in the reference, is expected within the U clade. Modal branch of each source in the best chain ($p$: its posterior probability) and $\Theta$ median [95% CI]: (A,B) 0.5X: B6a1 ($p = 1.00$), $\Theta = 0.64$ [0.26, 0.93]; HV2a1_1 ($p = 0.67$), $\Theta = 0.36$ [0.07, 0.74]; (C,D) 1X: B6a1 ($p = 0.97$), $\Theta = 0.69$ [0.36, 0.93]; V2a1a ($p = 0.72$), $\Theta = 0.31$ [0.07, 0.64]; (E,F) 2X: Node19 (stem of the U clade; $p = 0.89$), $\Theta = 0.95$ [0.91, 0.98]; Neanderthal_Hohlenstein ($p = 0.73$), $\Theta = 0.05$ [0.02, 0.09]; (G,H) 4X: U4b1b1 ($p = 0.93$), $\Theta = 0.75$ [0.65, 0.85]; D4a3a ($p = 1.00$), $\Theta = 0.25$ [0.15, 0.35]; (I,J) 8X: U7a4 ($p = 0.98$), $\Theta = 0.72$ [0.64, 0.80]; D4a3a ($p = 1.00$), $\Theta = 0.28$ [0.20, 0.36]; (K,L) 16X: U6a6a ($p = 1.00$), $\Theta = 0.80$ [0.75, 0.85]; D4a3a ($p = 1.00$), $\Theta = 0.20$ [0.15, 0.25]; (M,N) 32X: U6a6a ($p = 1.00$), $\Theta = 0.82$ [0.78, 0.85]; D4a3a ($p = 1.00$), $\Theta = 0.18$ [0.15, 0.22].

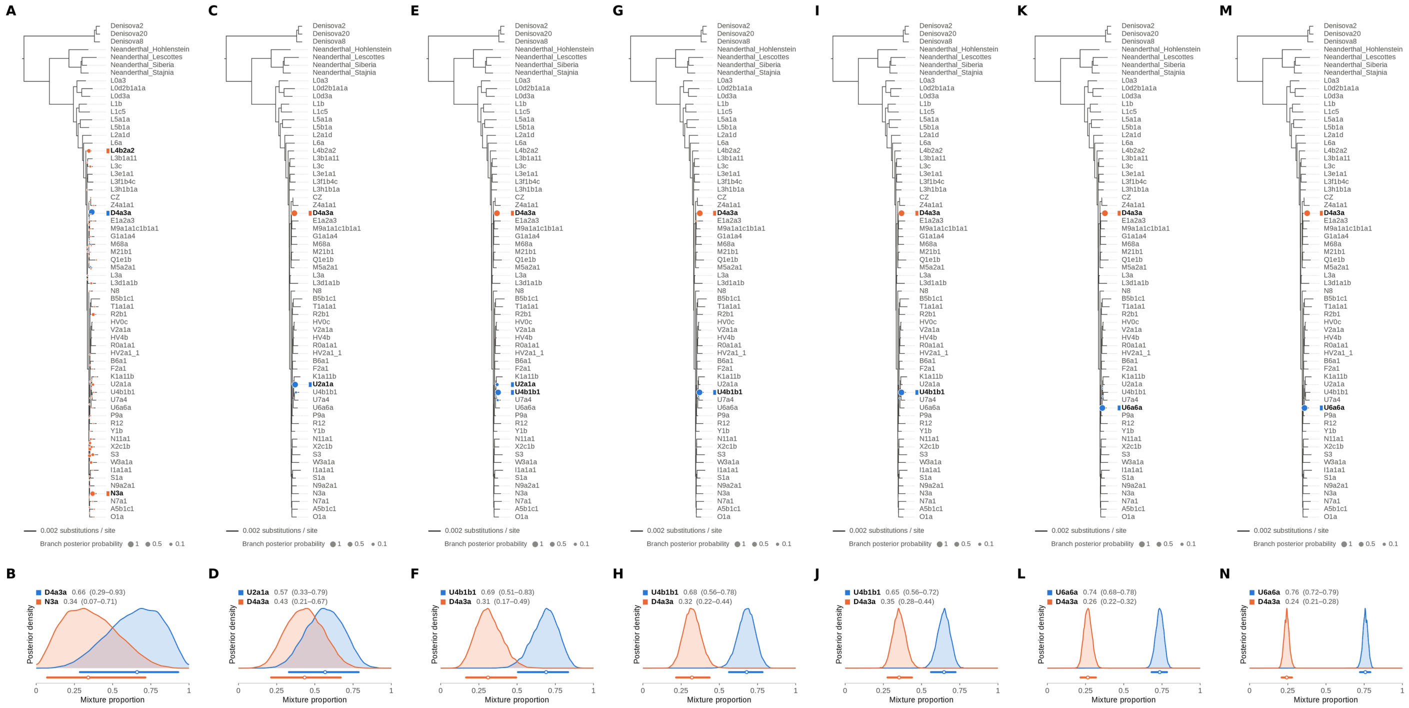

Figure S59: **Two-source simulated mixture of modern human D4a1d (FJ951517.1) (25% of reads) and modern human U5a2a (HM229344.1; 75%) at nominal mean coverage depths of 0.5X, 1X, 2X, 4X, 8X, 16X, 32X.** Both sources carry ancient DNA damage (U5a2a with the damage rates scaled by 0.75). For each coverage depth, the upper panel shows the phylogenetic placement posterior of both sources for the best of five chains ($k = 2$, 100,000 iterations each), colored by source (dot area: branch posterior probability; swelling: posterior density of the position along the branch; bold tips: highest-posterior branches), and the lower panel the posterior densities of the mixture proportions $\Theta$ with median and 95% credible interval. D4a1d is expected on the D4a3a branch (its closest relative in the reference); U5a2a, for which no U5 haplotype is present in the reference, is expected within the U clade. Modal branch of each source in the best chain ($p$: its posterior probability) and $\Theta$ median [95% CI]: (A,B) 0.5X: D4a3a ($p = 1.00$), $\Theta = 0.66$ [0.29, 0.93]; N3a ($p = 0.37$), $\Theta = 0.34$ [0.07, 0.71]; (C,D) 1X: U2a1a ($p = 0.99$), $\Theta = 0.57$ [0.33, 0.79]; D4a3a ($p = 1.00$), $\Theta = 0.43$ [0.21, 0.67]; (E,F) 2X: U4b1b1 ($p = 0.90$), $\Theta = 0.69$ [0.51, 0.83]; D4a3a ($p = 1.00$), $\Theta = 0.31$ [0.17, 0.49]; (G,H) 4X: U4b1b1 ($p = 1.00$), $\Theta = 0.68$ [0.56, 0.78]; D4a3a ($p = 1.00$), $\Theta = 0.32$ [0.22, 0.44]; (I,J) 8X: U4b1b1 ($p = 1.00$), $\Theta = 0.65$ [0.56, 0.72]; D4a3a ($p = 1.00$), $\Theta = 0.35$ [0.28, 0.44]; (K,L) 16X: U6a6a ($p = 1.00$), $\Theta = 0.74$ [0.68, 0.78]; D4a3a ($p = 1.00$), $\Theta = 0.26$ [0.22, 0.32]; (M,N) 32X: U6a6a ($p = 1.00$), $\Theta = 0.76$ [0.72, 0.79]; D4a3a ($p = 1.00$), $\Theta = 0.24$ [0.21, 0.28].

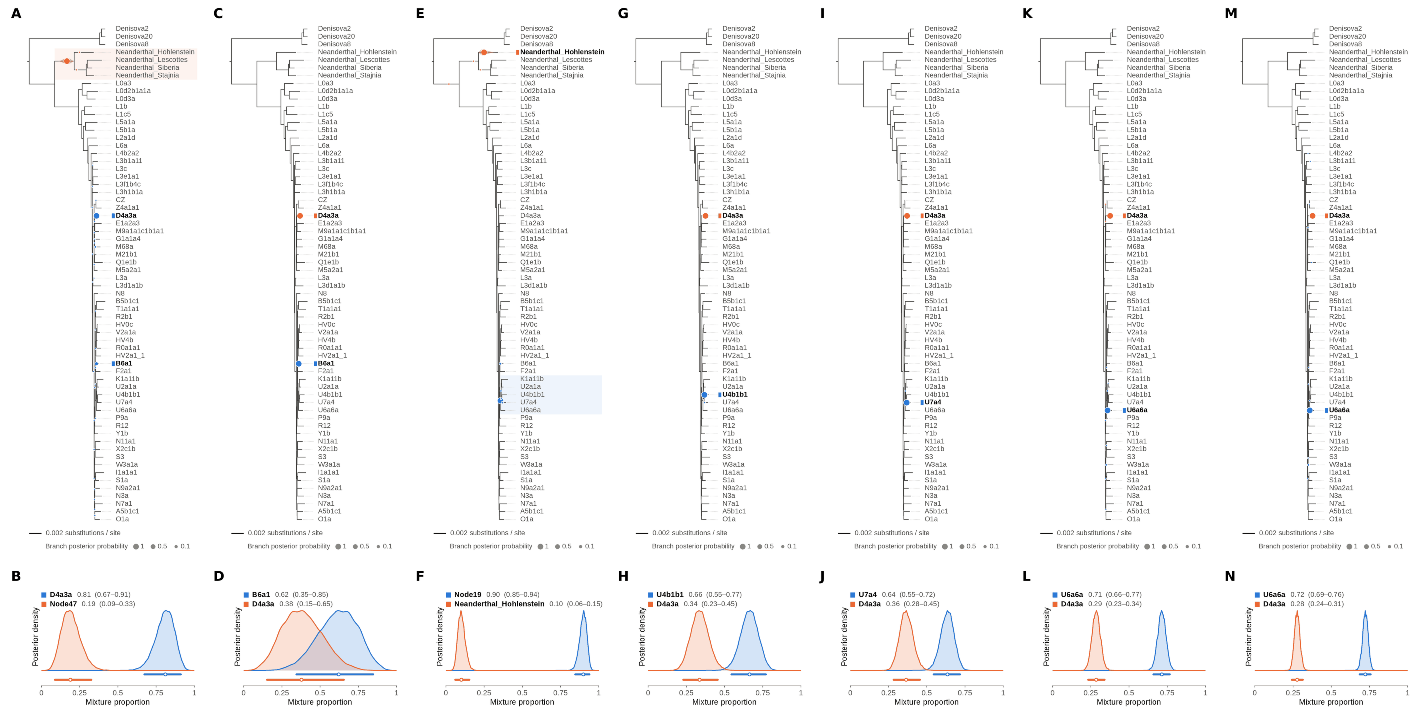


Figure S60: **Two-source simulated mixture of modern human D4a1d (FJ951517.1) (30% of reads) and modern human U5a2a (HM229344.1; 70%) at nominal mean coverage depths of 0.5X, 1X, 2X, 4X, 8X, 16X, 32X.** Both sources carry ancient DNA damage (U5a2a with the damage rates scaled by 0.75). For each coverage depth, the upper panel shows the phylogenetic placement posterior of both sources for the best of five chains ($k = 2$, 100,000 iterations each), colored by source (dot area: branch posterior probability; swelling: posterior density of the position along the branch; bold tips: highest-posterior branches), and the lower panel the posterior densities of the mixture proportions $\Theta$ with median and 95% credible interval. D4a1d is expected on the D4a3a branch (its closest relative in the reference); U5a2a, for which no U5 haplotype is present in the reference, is expected within the U clade. Modal branch of each source in the best chain ($p$: its posterior probability) and $\Theta$ median [95% CI]: (A,B) 0.5X: D4a3a ($p = 0.91$), $\Theta = 0.81$ [0.67, 0.91]; Node47 (stem of the Neanderthal clade; $p = 1.00$), $\Theta = 0.19$ [0.09, 0.33]; (C,D) 1X: B6a1 ($p = 1.00$), $\Theta = 0.62$ [0.35, 0.85]; D4a3a ($p = 1.00$), $\Theta = 0.38$ [0.15, 0.65]; (E,F) 2X: Node19 (stem of the U clade; $p = 0.98$), $\Theta = 0.90$ [0.85, 0.94]; Neanderthal_Hohlenstein ($p = 1.00$), $\Theta = 0.10$ [0.06, 0.15]; (G,H) 4X: U4b1b1 ($p = 1.00$), $\Theta = 0.66$ [0.55, 0.77]; D4a3a ($p = 1.00$), $\Theta = 0.34$ [0.23, 0.45]; (I,J) 8X: U7a4 ($p = 1.00$), $\Theta = 0.64$ [0.55, 0.72]; D4a3a ($p = 1.00$), $\Theta = 0.36$ [0.28, 0.45]; (K,L) 16X: U6a6a ($p = 0.99$), $\Theta = 0.71$ [0.66, 0.77]; D4a3a ($p = 1.00$), $\Theta = 0.29$ [0.23, 0.34]; (M,N) 32X: U6a6a ($p = 0.99$), $\Theta = 0.72$ [0.69, 0.76]; D4a3a ($p = 1.00$), $\Theta = 0.28$ [0.24, 0.31].

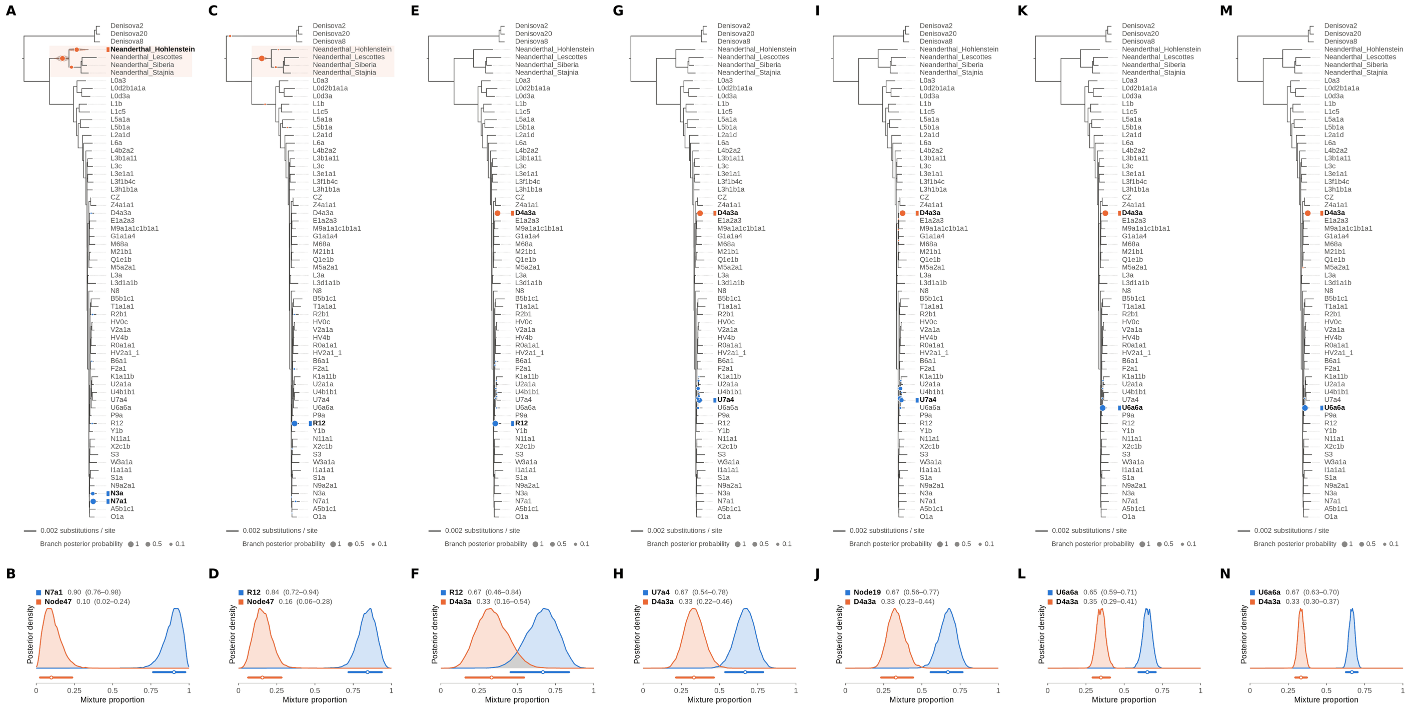

Figure S61: **Two-source simulated mixture of modern human D4a1d (FJ951517.1) (35% of reads) and modern human U5a2a (HM229344.1; 65%) at nominal mean coverage depths of 0.5X, 1X, 2X, 4X, 8X, 16X, 32X.** Both sources carry ancient DNA damage (U5a2a with the damage rates scaled by 0.75). For each coverage depth, the upper panel shows the phylogenetic placement posterior of both sources for the best of five chains ($k = 2$, 100,000 iterations each), colored by source (dot area: branch posterior probability; swelling: posterior density of the position along the branch; bold tips: highest-posterior branches), and the lower panel the posterior densities of the mixture proportions $\Theta$ with median and 95% credible interval. D4a1d is expected on the D4a3a branch (its closest relative in the reference); U5a2a, for which no U5 haplotype is present in the reference, is expected within the U clade. Modal branch of each source in the best chain ($p$: its posterior probability) and $\Theta$ median [95% CI]: (A,B) 0.5X: N7a1 ($p = 0.82$), $\Theta = 0.90$ [0.76, 0.98]; Node47 (stem of the Neanderthal clade; $p = 0.54$), $\Theta = 0.10$ [0.02, 0.24]; (C,D) 1X: R12 ($p = 0.99$), $\Theta = 0.84$ [0.72, 0.94]; Node47 (stem of the Neanderthal clade; $p = 0.96$), $\Theta = 0.16$ [0.06, 0.28]; (E,F) 2X: R12 ($p = 1.00$), $\Theta = 0.67$ [0.46, 0.84]; D4a3a ($p = 1.00$), $\Theta = 0.33$ [0.16, 0.54]; (G,H) 4X: U7a4 ($p = 0.69$), $\Theta = 0.67$ [0.54, 0.78]; D4a3a ($p = 1.00$), $\Theta = 0.33$ [0.22, 0.46]; (I,J) 8X: Node19 (stem of the U clade; $p = 0.49$), $\Theta = 0.67$ [0.56, 0.77]; D4a3a ($p = 1.00$), $\Theta = 0.33$ [0.23, 0.44]; (K,L) 16X: U6a6a ($p = 1.00$), $\Theta = 0.65$ [0.59, 0.71]; D4a3a ($p = 1.00$), $\Theta = 0.35$ [0.29, 0.41]; (M,N) 32X: U6a6a ($p = 1.00$), $\Theta = 0.67$ [0.63, 0.70]; D4a3a ($p = 1.00$), $\Theta = 0.33$ [0.30, 0.37].

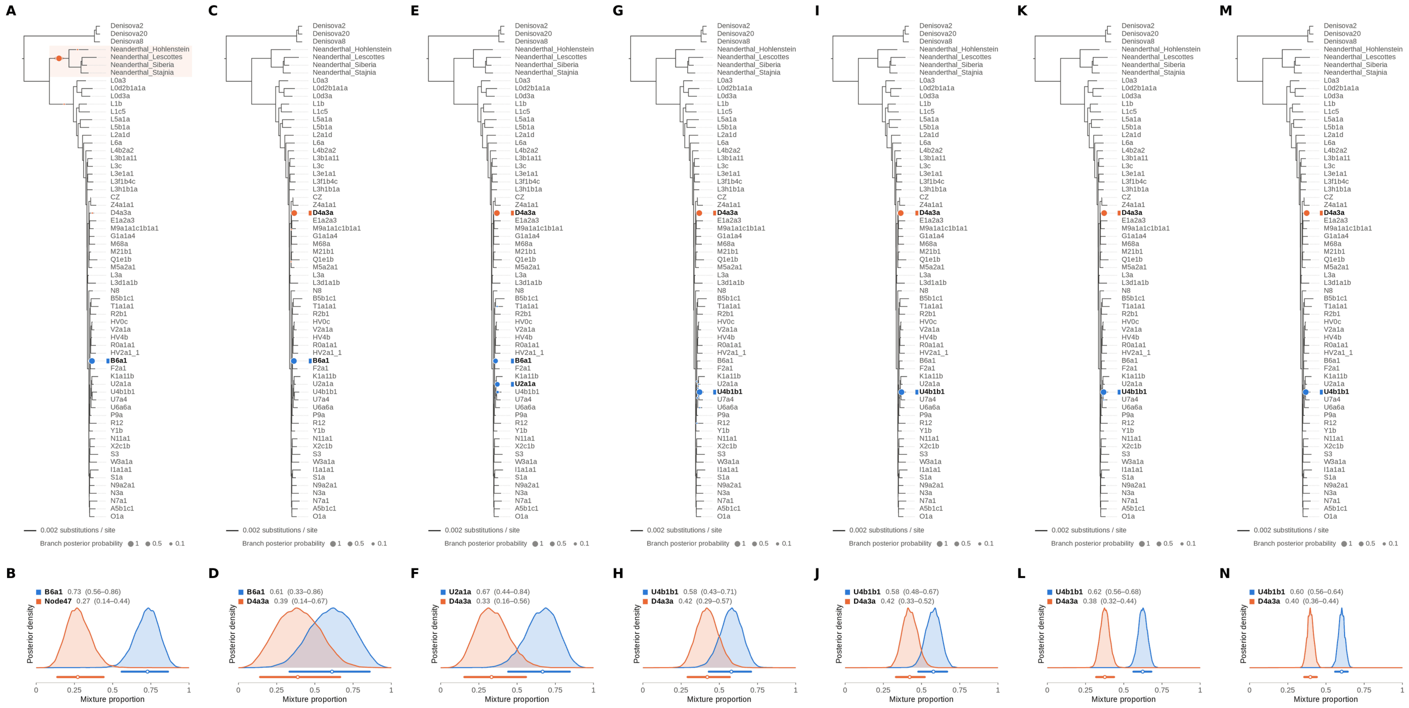


Figure S62: **Two-source simulated mixture of modern human D4a1d (FJ951517.1) (40% of reads) and modern human U5a2a (HM229344.1; 60%) at nominal mean coverage depths of 0.5X, 1X, 2X, 4X, 8X, 16X, 32X.** Both sources carry ancient DNA damage (U5a2a with the damage rates scaled by 0.75). For each coverage depth, the upper panel shows the phylogenetic placement posterior of both sources for the best of five chains ($k = 2$, 100,000 iterations each), colored by source (dot area: branch posterior probability; swelling: posterior density of the position along the branch; bold tips: highest-posterior branches), and the lower panel the posterior densities of the mixture proportions $\Theta$ with median and 95% credible interval. D4a1d is expected on the D4a3a branch (its closest relative in the reference); U5a2a, for which no U5 haplotype is present in the reference, is expected within the U clade. Modal branch of each source in the best chain ($p$: its posterior probability) and $\Theta$ median [95% CI]: (A,B) 0.5X: B6a1 ($p = 1.00$), $\Theta = 0.73$ [0.56, 0.86]; Node47 (stem of the Neanderthal clade; $p = 1.00$), $\Theta = 0.27$ [0.14, 0.44]; (C,D) 1X: B6a1 ($p = 1.00$), $\Theta = 0.61$ [0.33, 0.86]; D4a3a ($p = 1.00$), $\Theta = 0.39$ [0.14, 0.67]; (E,F) 2X: U2a1a ($p = 0.48$), $\Theta = 0.67$ [0.44, 0.84]; D4a3a ($p = 1.00$), $\Theta = 0.33$ [0.16, 0.56]; (G,H) 4X: U4b1b1 ($p = 0.99$), $\Theta = 0.58$ [0.43, 0.71]; D4a3a ($p = 1.00$), $\Theta = 0.42$ [0.29, 0.57]; (I,J) 8X: U4b1b1 ($p = 1.00$), $\Theta = 0.58$ [0.48, 0.67]; D4a3a ($p = 1.00$), $\Theta = 0.42$ [0.33, 0.52]; (K,L) 16X: U4b1b1 ($p = 1.00$), $\Theta = 0.62$ [0.56, 0.68]; D4a3a ($p = 1.00$), $\Theta = 0.38$ [0.32, 0.44]; (M,N) 32X: U4b1b1 ($p = 1.00$), $\Theta = 0.60$ [0.56, 0.64]; D4a3a ($p = 1.00$), $\Theta = 0.40$ [0.36, 0.44].

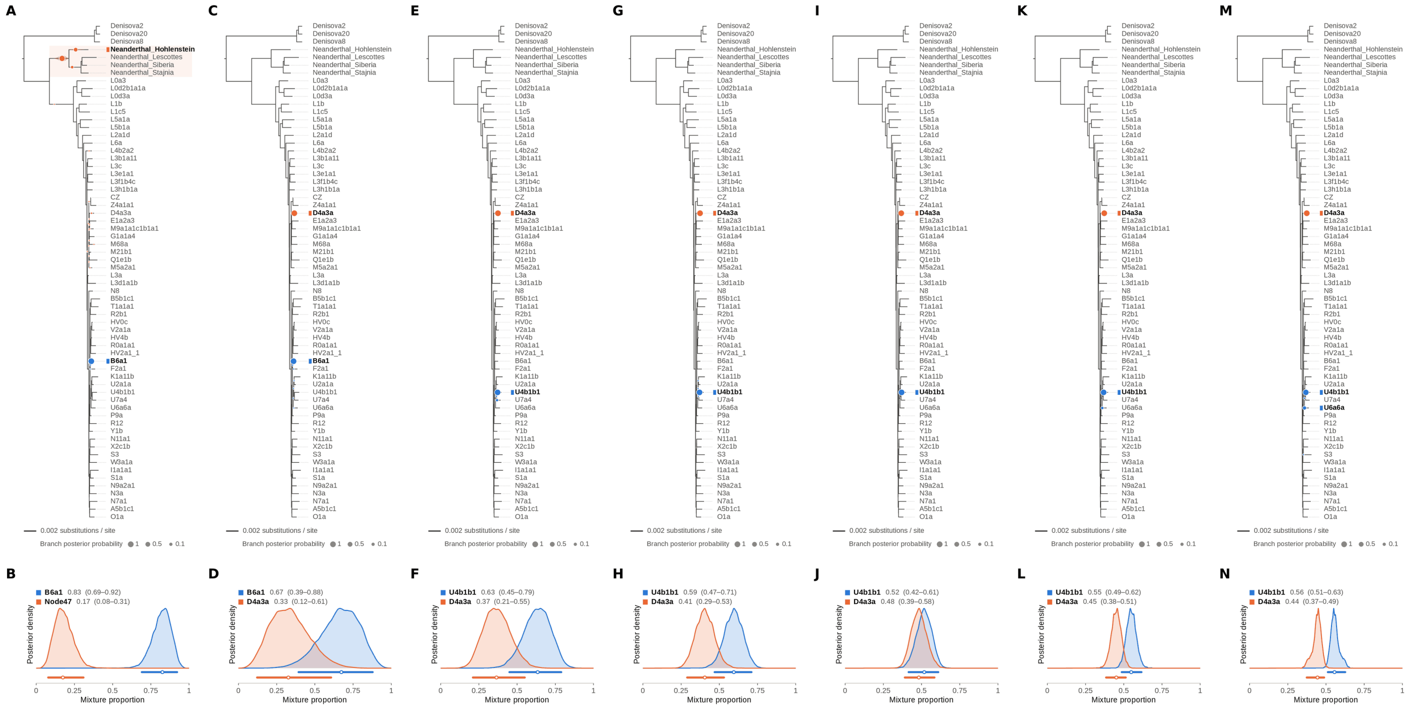


Figure S63: **Two-source simulated mixture of modern human D4a1d (FJ951517.1) (45% of reads) and modern human U5a2a (HM229344.1; 55%) at nominal mean coverage depths of 0.5X, 1X, 2X, 4X, 8X, 16X, 32X.** Both sources carry ancient DNA damage (U5a2a with the damage rates scaled by 0.75). For each coverage depth, the upper panel shows the phylogenetic placement posterior of both sources for the best of five chains ($k = 2$, 100,000 iterations each), colored by source (dot area: branch posterior probability; swelling: posterior density of the position along the branch; bold tips: highest-posterior branches), and the lower panel the posterior densities of the mixture proportions $\Theta$ with median and 95% credible interval. D4a1d is expected on the D4a3a branch (its closest relative in the reference); U5a2a, for which no U5 haplotype is present in the reference, is expected within the U clade. Modal branch of each source in the best chain ($p$: its posterior probability) and $\Theta$ median [95% CI]: (A,B) 0.5X: B6a1 ($p = 1.00$), $\Theta = 0.83$ [0.69, 0.92]; Node47 (stem of the Neanderthal clade; $p = 0.76$), $\Theta = 0.17$ [0.08, 0.31]; (C,D) 1X: B6a1 ($p = 1.00$), $\Theta = 0.67$ [0.39, 0.88]; D4a3a ($p = 1.00$), $\Theta = 0.33$ [0.12, 0.61]; (E,F) 2X: U4b1b1 ($p = 0.97$), $\Theta = 0.63$ [0.45, 0.79]; D4a3a ($p = 1.00$), $\Theta = 0.37$ [0.21, 0.55]; (G,H) 4X: U4b1b1 ($p = 1.00$), $\Theta = 0.59$ [0.47, 0.71]; D4a3a ($p = 1.00$), $\Theta = 0.41$ [0.29, 0.53]; (I,J) 8X: U4b1b1 ($p = 1.00$), $\Theta = 0.52$ [0.42, 0.61]; D4a3a ($p = 1.00$), $\Theta = 0.48$ [0.39, 0.58]; (K,L) 16X: U4b1b1 ($p = 0.96$), $\Theta = 0.55$ [0.49, 0.62]; D4a3a ($p = 1.00$), $\Theta = 0.45$ [0.38, 0.51]; (M,N) 32X: U4b1b1 ($p = 0.87$), $\Theta = 0.56$ [0.51, 0.63]; D4a3a ($p = 1.00$), $\Theta = 0.44$ [0.37, 0.49].

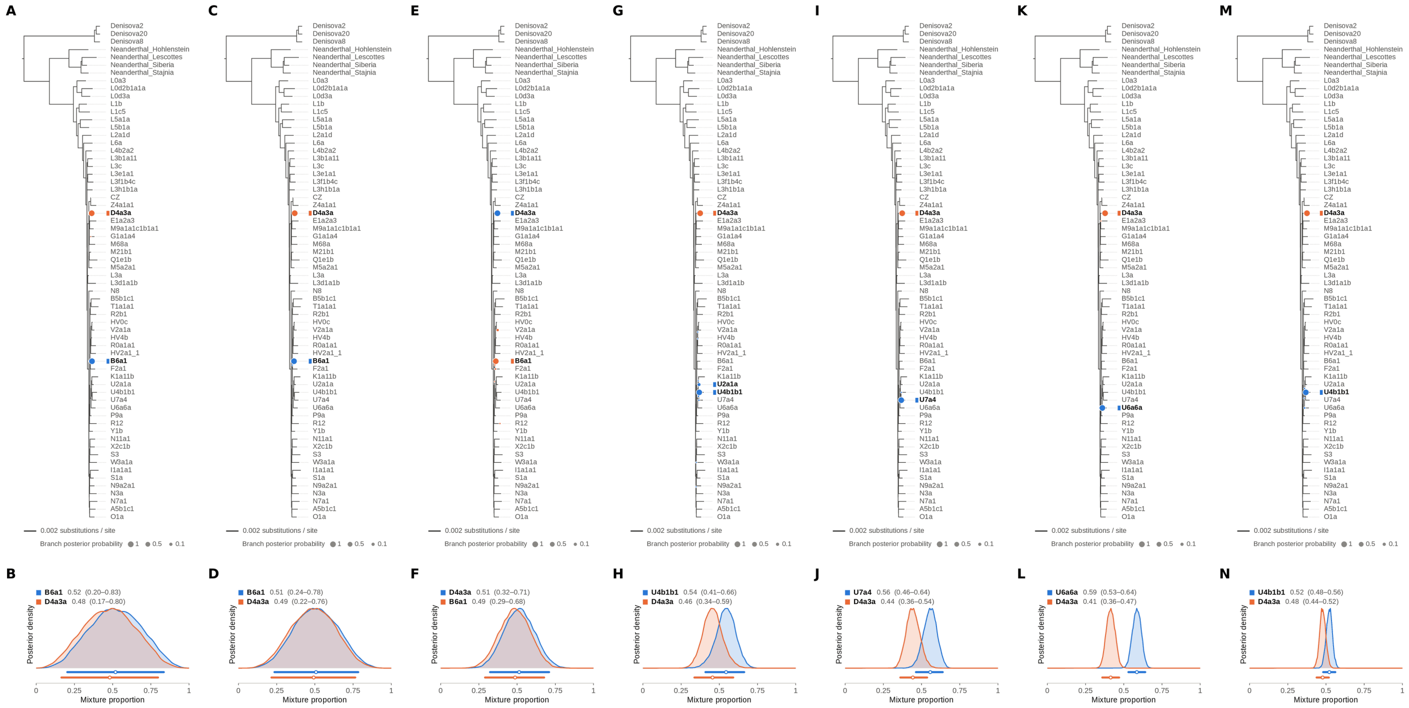


Figure S64: **Two-source simulated mixture of modern human D4a1d (FJ951517.1) (50% of reads) and modern human U5a2a (HM229344.1; 50%) at nominal mean coverage depths of 0.5X, 1X, 2X, 4X, 8X, 16X, 32X.** Both sources carry ancient DNA damage (U5a2a with the damage rates scaled by 0.75). For each coverage depth, the upper panel shows the phylogenetic placement posterior of both sources for the best of five chains ($k = 2$, 100,000 iterations each), colored by source (dot area: branch posterior probability; swelling: posterior density of the position along the branch; bold tips: highest-posterior branches), and the lower panel the posterior densities of the mixture proportions $\Theta$ with median and 95% credible interval. D4a1d is expected on the D4a3a branch (its closest relative in the reference); U5a2a, for which no U5 haplotype is present in the reference, is expected within the U clade. Modal branch of each source in the best chain ($p$: its posterior probability) and $\Theta$ median [95% CI]: (A,B) 0.5X: B6a1 ($p = 1.00$), $\Theta = 0.52$ [0.20, 0.83]; D4a3a ($p = 1.00$), $\Theta = 0.48$ [0.17, 0.80]; (C,D) 1X: B6a1 ($p = 1.00$), $\Theta = 0.51$ [0.24, 0.78]; D4a3a ($p = 1.00$), $\Theta = 0.49$ [0.22, 0.76]; (E,F) 2X: D4a3a ($p = 1.00$), $\Theta = 0.51$ [0.32, 0.71]; B6a1 ($p = 0.97$), $\Theta = 0.49$ [0.29, 0.68]; (G,H) 4X: U4b1b1 ($p = 0.92$), $\Theta = 0.54$ [0.41, 0.66]; D4a3a ($p = 1.00$), $\Theta = 0.46$ [0.34, 0.59]; (I,J) 8X: U7a4 ($p = 1.00$), $\Theta = 0.56$ [0.46, 0.64]; D4a3a ($p = 1.00$), $\Theta = 0.44$ [0.36, 0.54]; (K,L) 16X: U6a6a ($p = 1.00$), $\Theta = 0.59$ [0.53, 0.64]; D4a3a ($p = 1.00$), $\Theta = 0.41$ [0.36, 0.47]; (M,N) 32X: U4b1b1 ($p = 0.99$), $\Theta = 0.52$ [0.48, 0.56]; D4a3a ($p = 1.00$), $\Theta = 0.48$ [0.44, 0.52].

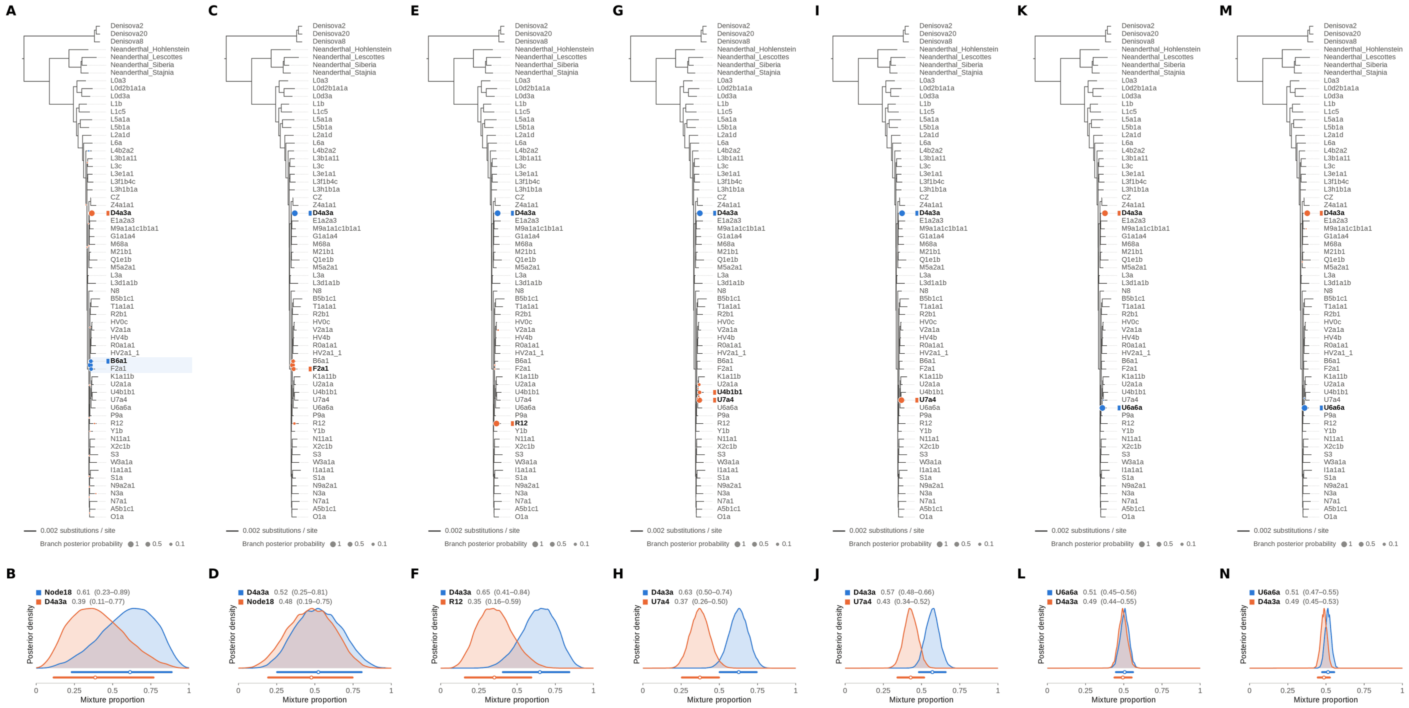


Figure S65: **Two-source simulated mixture of modern human D4a1d (FJ951517.1) (55% of reads) and modern human U5a2a (HM229344.1; 45%) at nominal mean coverage depths of 0.5X, 1X, 2X, 4X, 8X, 16X, 32X.** Both sources carry ancient DNA damage (U5a2a with the damage rates scaled by 0.75). For each coverage depth, the upper panel shows the phylogenetic placement posterior of both sources for the best of five chains ($k = 2$, 100,000 iterations each), colored by source (dot area: branch posterior probability; swelling: posterior density of the position along the branch; bold tips: highest-posterior branches), and the lower panel the posterior densities of the mixture proportions $\Theta$ with median and 95% credible interval. D4a1d is expected on the D4a3a branch (its closest relative in the reference); U5a2a, for which no U5 haplotype is present in the reference, is expected within the U clade. Modal branch of each source in the best chain ($p$: its posterior probability) and $\Theta$ median [95% CI]: (A,B) 0.5X: Node18 (internal branch ancestral to B6a1 and F2a1; $p = 0.58$), $\Theta = 0.61$ [0.23, 0.89]; D4a3a ($p = 1.00$), $\Theta = 0.39$ [0.11, 0.77]; (C,D) 1X: D4a3a ($p = 1.00$), $\Theta = 0.52$ [0.25, 0.81]; Node18 (internal branch ancestral to B6a1 and F2a1; $p = 0.49$), $\Theta = 0.48$ [0.19, 0.75]; (E,F) 2X: D4a3a ($p = 1.00$), $\Theta = 0.65$ [0.41, 0.84]; R12 ($p = 1.00$), $\Theta = 0.35$ [0.16, 0.59]; (G,H) 4X: D4a3a ($p = 1.00$), $\Theta = 0.63$ [0.50, 0.74]; U7a4 ($p = 0.77$), $\Theta = 0.37$ [0.26, 0.50]; (I,J) 8X: D4a3a ($p = 1.00$), $\Theta = 0.57$ [0.48, 0.66]; U7a4 ($p = 1.00$), $\Theta = 0.43$ [0.34, 0.52]; (K,L) 16X: U6a6a ($p = 1.00$), $\Theta = 0.51$ [0.45, 0.56]; D4a3a ($p = 1.00$), $\Theta = 0.49$ [0.44, 0.55]; (M,N) 32X: U6a6a ($p = 1.00$), $\Theta = 0.51$ [0.47, 0.55]; D4a3a ($p = 1.00$), $\Theta = 0.49$ [0.45, 0.53].

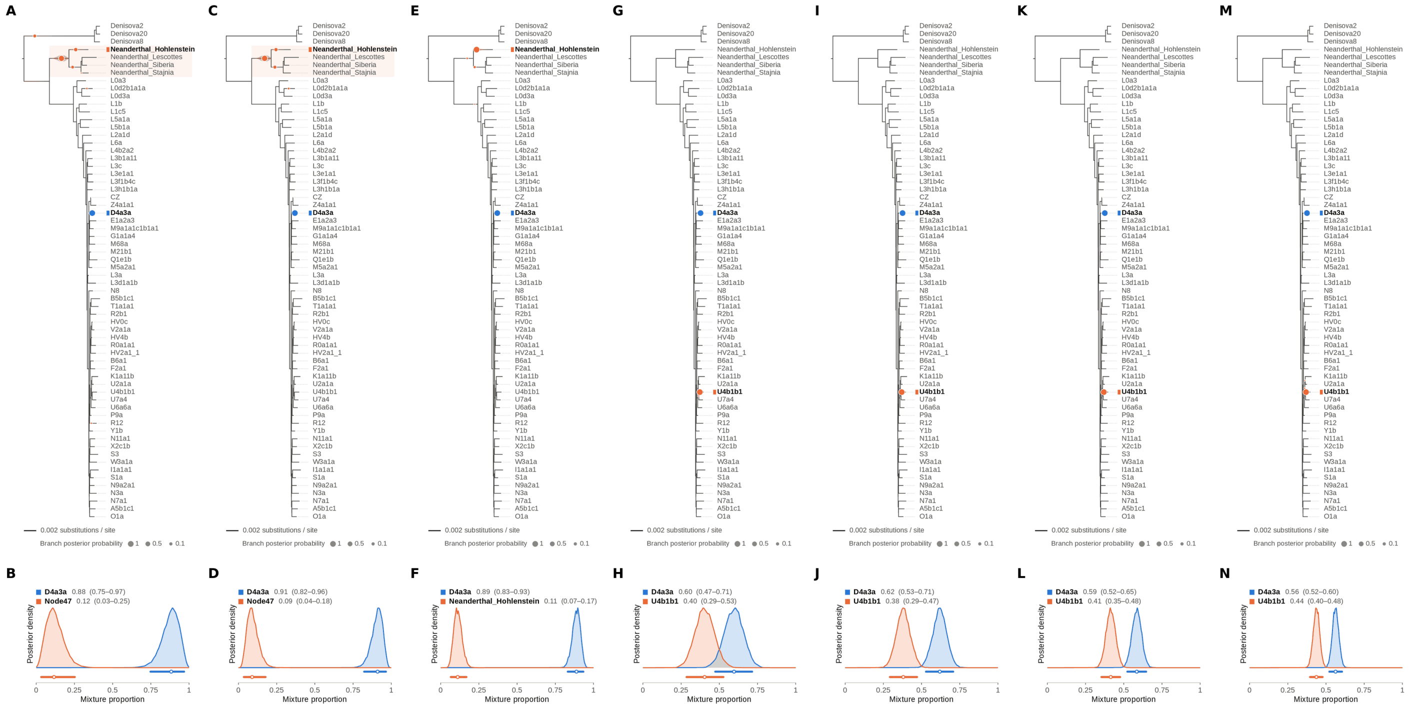

Figure S66: **Two-source simulated mixture of modern human D4a1d (FJ951517.1) (60% of reads) and modern human U5a2a (HM229344.1; 40%) at nominal mean coverage depths of 0.5X, 1X, 2X, 4X, 8X, 16X, 32X.** Both sources carry ancient DNA damage (U5a2a with the damage rates scaled by 0.75). For each coverage depth, the upper panel shows the phylogenetic placement posterior of both sources for the best of five chains ($k = 2$, 100,000 iterations each), colored by source (dot area: branch posterior probability; swelling: posterior density of the position along the branch; bold tips: highest-posterior branches), and the lower panel the posterior densities of the mixture proportions Θ with median and 95% credible interval. D4a1d is expected on the D4a3a branch (its closest relative in the reference); U5a2a, for which no U5 haplotype is present in the reference, is expected within the U clade. Modal branch of each source in the best chain ($p$: its posterior probability) and Θ median [95% CI]: (A,B) 0.5X: D4a3a ($p = 1.00$), $\Theta = 0.88$ [0.75, 0.97]; Node47 (stem of the Neanderthal clade; $p = 0.72$), $\Theta = 0.12$ [0.03, 0.25]; (C,D) 1X: D4a3a ($p = 1.00$), $\Theta = 0.91$ [0.82, 0.96]; Node47 (stem of the Neanderthal clade; $p = 0.79$), $\Theta = 0.09$ [0.04, 0.18]; (E,F) 2X: D4a3a ($p = 1.00$), $\Theta = 0.89$ [0.83, 0.93]; Neanderthal_Hohlenstein ($p = 0.97$), $\Theta = 0.11$ [0.07, 0.17]; (G,H) 4X: D4a3a ($p = 1.00$), $\Theta = 0.60$ [0.47, 0.71]; U4b1b1 ($p = 1.00$), $\Theta = 0.40$ [0.29, 0.53]; (I,J) 8X: D4a3a ($p = 1.00$), $\Theta = 0.62$ [0.53, 0.71]; U4b1b1 ($p = 1.00$), $\Theta = 0.38$ [0.29, 0.47]; (K,L) 16X: D4a3a ($p = 1.00$), $\Theta = 0.59$ [0.52, 0.65]; U4b1b1 ($p = 1.00$), $\Theta = 0.41$ [0.35, 0.48]; (M,N) 32X: D4a3a ($p = 1.00$), $\Theta = 0.56$ [0.52, 0.60]; U4b1b1 ($p = 1.00$), $\Theta = 0.44$ [0.40, 0.48].

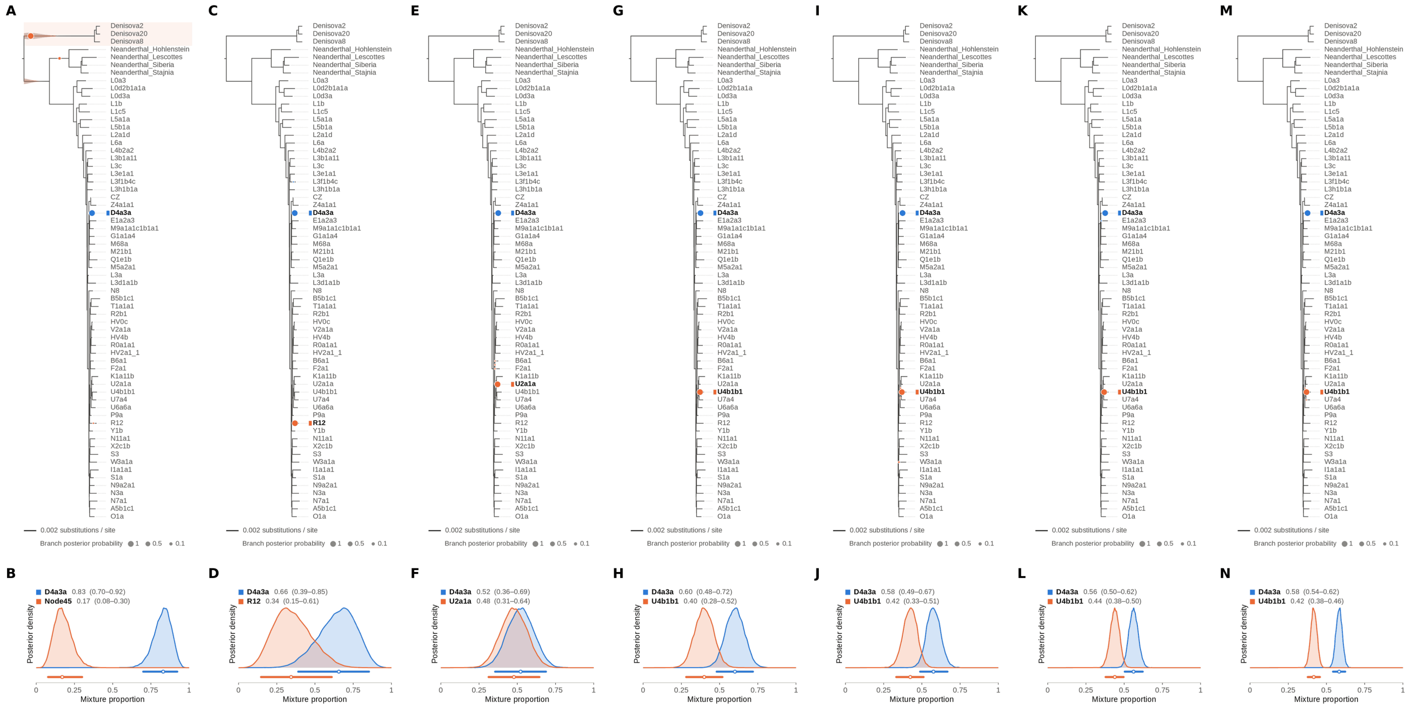

Figure S67: **Two-source simulated mixture of modern human D4a1d (FJ951517.1) (65% of reads) and modern human U5a2a (HM229344.1; 35%) at nominal mean coverage depths of 0.5X, 1X, 2X, 4X, 8X, 16X, 32X.** Both sources carry ancient DNA damage (U5a2a with the damage rates scaled by 0.75). For each coverage depth, the upper panel shows the phylogenetic placement posterior of both sources for the best of five chains ($k = 2$, 100,000 iterations each), colored by source (dot area: branch posterior probability; swelling: posterior density of the position along the branch; bold tips: highest-posterior branches), and the lower panel the posterior densities of the mixture proportions $\Theta$ with median and 95% credible interval. D4a1d is expected on the D4a3a branch (its closest relative in the reference); U5a2a, for which no U5 haplotype is present in the reference, is expected within the U clade. Modal branch of each source in the best chain ($p$: its posterior probability) and $\Theta$ median [95% CI]: (A,B) 0.5X: D4a3a ($p = 1.00$), $\Theta = 0.83$ [0.70, 0.92]; Node45 (stem of the Denisovan clade; $p = 0.96$), $\Theta = 0.17$ [0.08, 0.30]; (C,D) 1X: D4a3a ($p = 1.00$), $\Theta = 0.66$ [0.39, 0.85]; R12 ($p = 1.00$), $\Theta = 0.34$ [0.15, 0.61]; (E,F) 2X: D4a3a ($p = 1.00$), $\Theta = 0.52$ [0.36, 0.69]; U2a1a ($p = 1.00$), $\Theta = 0.48$ [0.31, 0.64]; (G,H) 4X: D4a3a ($p = 1.00$), $\Theta = 0.60$ [0.48, 0.72]; U4b1b1 ($p = 1.00$), $\Theta = 0.40$ [0.28, 0.52]; (I,J) 8X: D4a3a ($p = 1.00$), $\Theta = 0.58$ [0.49, 0.67]; U4b1b1 ($p = 0.99$), $\Theta = 0.42$ [0.33, 0.51]; (K,L) 16X: D4a3a ($p = 1.00$), $\Theta = 0.56$ [0.50, 0.62]; U4b1b1 ($p = 1.00$), $\Theta = 0.44$ [0.38, 0.50]; (M,N) 32X: D4a3a ($p = 1.00$), $\Theta = 0.58$ [0.54, 0.62]; U4b1b1 ($p = 1.00$), $\Theta = 0.42$ [0.38, 0.46].

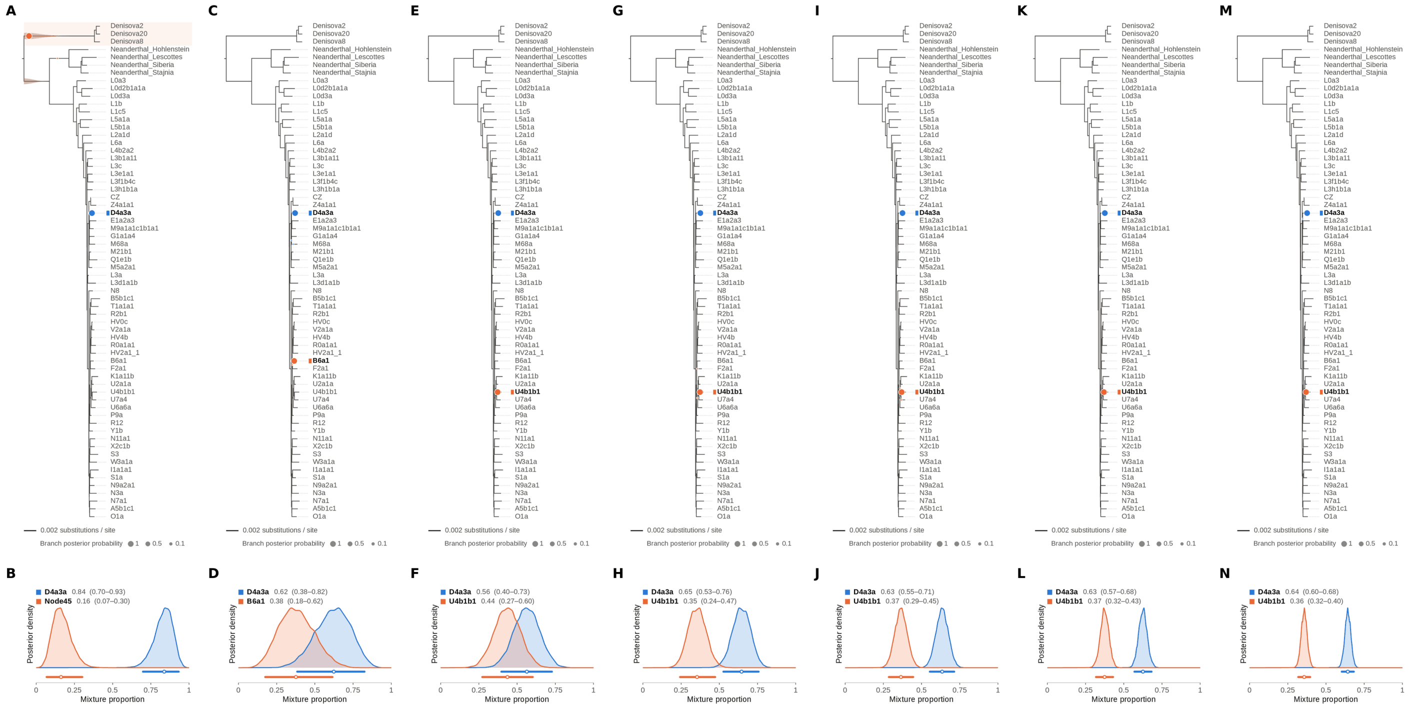

Figure S68: **Two-source simulated mixture of modern human D4a1d (FJ951517.1) (70% of reads) and modern human U5a2a (HM229344.1; 30%) at nominal mean coverage depths of 0.5X, 1X, 2X, 4X, 8X, 16X, 32X.** Both sources carry ancient DNA damage (U5a2a with the damage rates scaled by 0.75). For each coverage depth, the upper panel shows the phylogenetic placement posterior of both sources for the best of five chains ($k = 2$, 100,000 iterations each), colored by source (dot area: branch posterior probability; swelling: posterior density of the position along the branch; bold tips: highest-posterior branches), and the lower panel the posterior densities of the mixture proportions $\Theta$ with median and 95% credible interval. D4a1d is expected on the D4a3a branch (its closest relative in the reference); U5a2a, for which no U5 haplotype is present in the reference, is expected within the U clade. Modal branch of each source in the best chain ($p$: its posterior probability) and $\Theta$ median [95% CI]: (A,B) 0.5X: D4a3a ($p = 1.00$), $\Theta = 0.84$ [0.70, 0.93]; Node45 (stem of the Denisovan clade; $p = 1.00$), $\Theta = 0.16$ [0.07, 0.30]; (C,D) 1X: D4a3a ($p = 1.00$), $\Theta = 0.62$ [0.38, 0.82]; B6a1 ($p = 1.00$), $\Theta = 0.38$ [0.18, 0.62]; (E,F) 2X: D4a3a ($p = 1.00$), $\Theta = 0.56$ [0.40, 0.73]; U4b1b1 ($p = 1.00$), $\Theta = 0.44$ [0.27, 0.60]; (G,H) 4X: D4a3a ($p = 1.00$), $\Theta = 0.65$ [0.53, 0.76]; U4b1b1 ($p = 1.00$), $\Theta = 0.35$ [0.24, 0.47]; (I,J) 8X: D4a3a ($p = 1.00$), $\Theta = 0.63$ [0.55, 0.71]; U4b1b1 ($p = 1.00$), $\Theta = 0.37$ [0.29, 0.45]; (K,L) 16X: D4a3a ($p = 1.00$), $\Theta = 0.63$ [0.57, 0.68]; U4b1b1 ($p = 1.00$), $\Theta = 0.37$ [0.32, 0.43]; (M,N) 32X: D4a3a ($p = 1.00$), $\Theta = 0.64$ [0.60, 0.68]; U4b1b1 ($p = 1.00$), $\Theta = 0.36$ [0.32, 0.40].

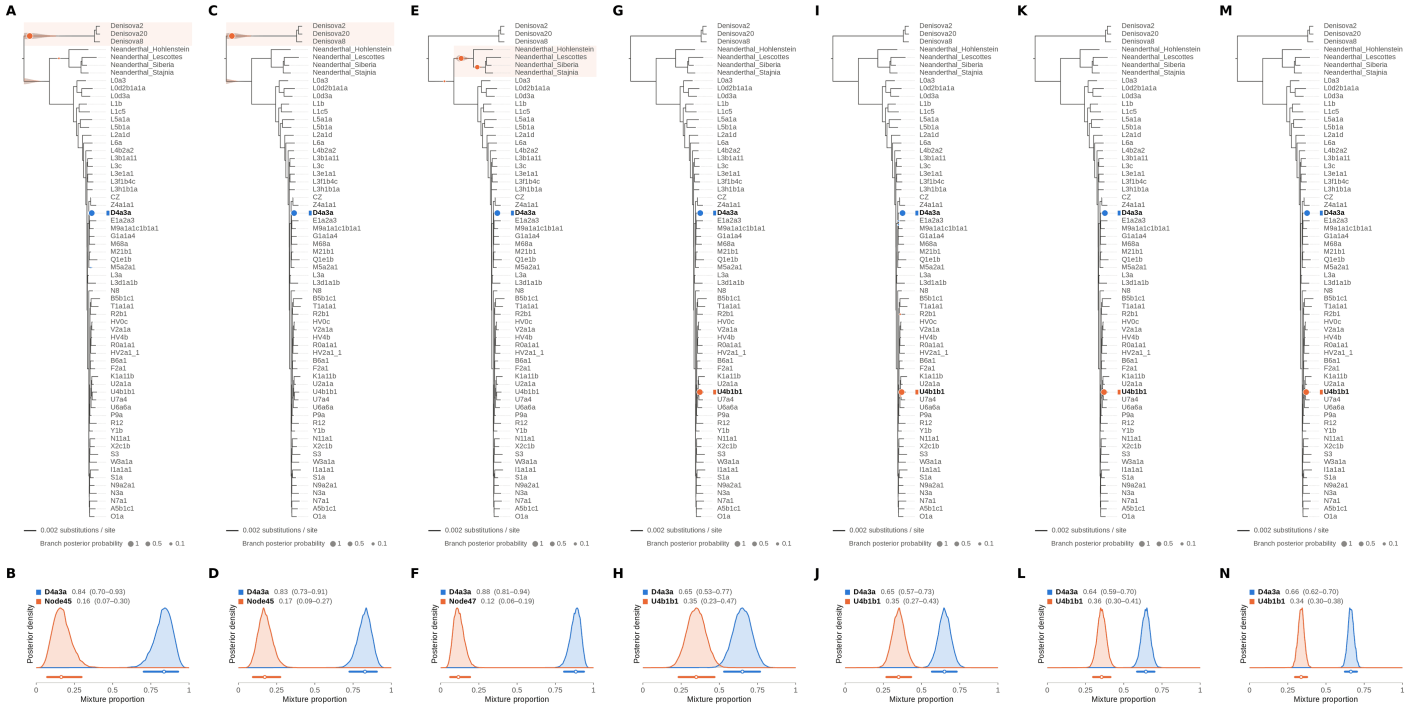


Figure S69: **Two-source simulated mixture of modern human D4a1d (FJ951517.1) (75% of reads) and modern human U5a2a (HM229344.1; 25%) at nominal mean coverage depths of 0.5X, 1X, 2X, 4X, 8X, 16X, 32X.** Both sources carry ancient DNA damage (U5a2a with the damage rates scaled by 0.75). For each coverage depth, the upper panel shows the phylogenetic placement posterior of both sources for the best of five chains ($k = 2$, 100,000 iterations each), colored by source (dot area: branch posterior probability; swelling: posterior density of the position along the branch; bold tips: highest-posterior branches), and the lower panel the posterior densities of the mixture proportions $\Theta$ with median and 95% credible interval. D4a1d is expected on the D4a3a branch (its closest relative in the reference); U5a2a, for which no U5 haplotype is present in the reference, is expected within the U clade. Modal branch of each source in the best chain ($p$: its posterior probability) and $\Theta$ median [95% CI]: (A,B) 0.5X: D4a3a ($p = 1.00$), $\Theta = 0.84$ [0.70, 0.93]; Node45 (stem of the Denisovan clade; $p = 1.00$), $\Theta = 0.16$ [0.07, 0.30]; (C,D) 1X: D4a3a ($p = 1.00$), $\Theta = 0.83$ [0.73, 0.91]; Node45 (stem of the Denisovan clade; $p = 1.00$), $\Theta = 0.17$ [0.09, 0.27]; (E,F) 2X: D4a3a ($p = 1.00$), $\Theta = 0.88$ [0.81, 0.94]; Node47 (stem of the Neanderthal clade; $p = 0.61$), $\Theta = 0.12$ [0.06, 0.19]; (G,H) 4X: D4a3a ($p = 1.00$), $\Theta = 0.65$ [0.53, 0.77]; U4b1b1 ($p = 1.00$), $\Theta = 0.35$ [0.23, 0.47]; (I,J) 8X: D4a3a ($p = 1.00$), $\Theta = 0.65$ [0.57, 0.73]; U4b1b1 ($p = 1.00$), $\Theta = 0.35$ [0.27, 0.43]; (K,L) 16X: D4a3a ($p = 1.00$), $\Theta = 0.64$ [0.59, 0.70]; U4b1b1 ($p = 1.00$), $\Theta = 0.36$ [0.30, 0.41]; (M,N) 32X: D4a3a ($p = 1.00$), $\Theta = 0.66$ [0.62, 0.70]; U4b1b1 ($p = 1.00$), $\Theta = 0.34$ [0.30, 0.38].

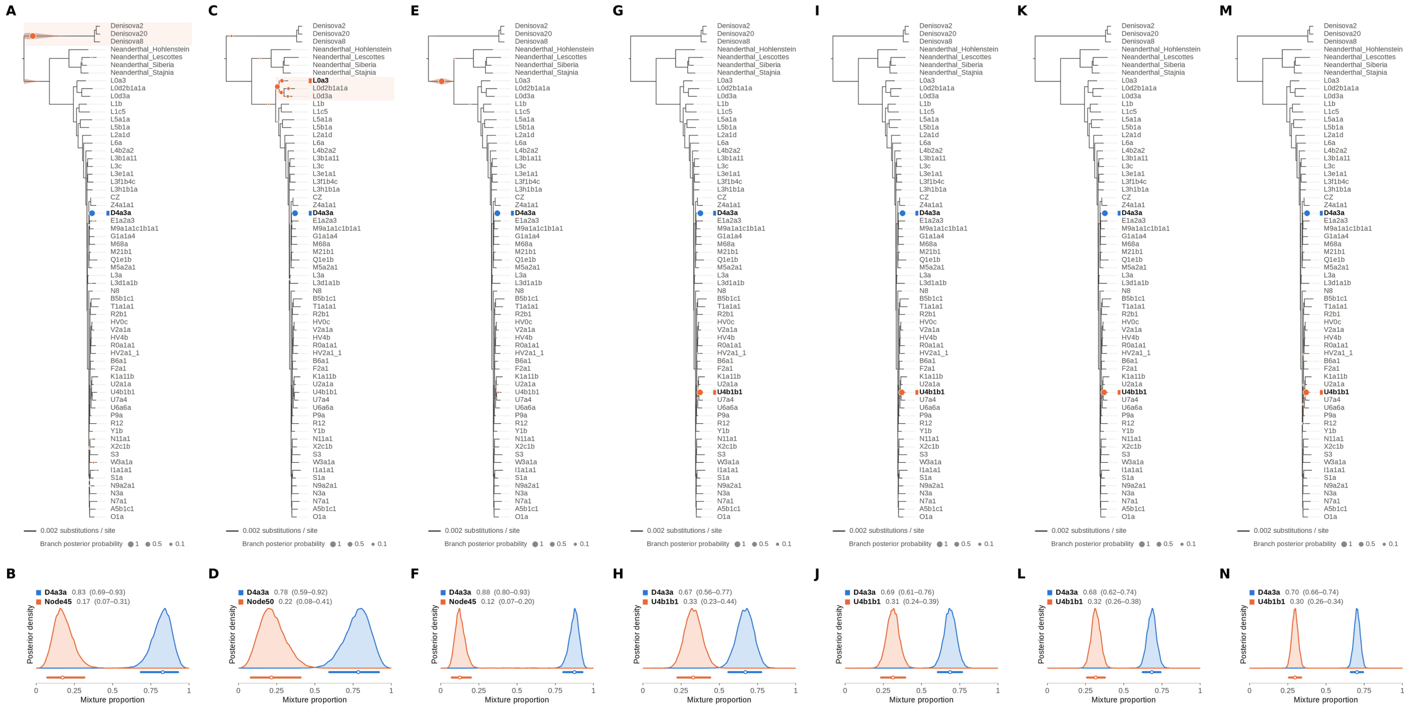

Figure S70: **Two-source simulated mixture of modern human D4a1d (FJ951517.1) (80% of reads) and modern human U5a2a (HM229344.1; 20%) at nominal mean coverage depths of 0.5X, 1X, 2X, 4X, 8X, 16X, 32X.** Both sources carry ancient DNA damage (U5a2a with the damage rates scaled by 0.75). For each coverage depth, the upper panel shows the phylogenetic placement posterior of both sources for the best of five chains ($k = 2$, 100,000 iterations each), colored by source (dot area: branch posterior probability; swelling: posterior density of the position along the branch; bold tips: highest-posterior branches), and the lower panel the posterior densities of the mixture proportions $\Theta$ with median and 95% credible interval. D4a1d is expected on the D4a3a branch (its closest relative in the reference); U5a2a, for which no U5 haplotype is present in the reference, is expected within the U clade. Modal branch of each source in the best chain ($p$: its posterior probability) and $\Theta$ median [95% CI]: (A,B) 0.5X: D4a3a ($p = 1.00$), $\Theta = 0.83$ [0.69, 0.93]; Node45 (stem of the Denisovan clade; $p = 0.99$), $\Theta = 0.17$ [0.07, 0.31]; (C,D) 1X: D4a3a ($p = 1.00$), $\Theta = 0.78$ [0.59, 0.92]; Node50 (internal branch ancestral to L0a3, L0d2b1a1a and L0d3a; $p = 0.55$), $\Theta = 0.22$ [0.08, 0.41]; (E,F) 2X: D4a3a ($p = 1.00$), $\Theta = 0.88$ [0.80, 0.93]; Node45 (stem of the Denisovan clade; $p = 0.99$), $\Theta = 0.12$ [0.07, 0.20]; (G,H) 4X: D4a3a ($p = 1.00$), $\Theta = 0.67$ [0.56, 0.77]; U4b1b1 ($p = 1.00$), $\Theta = 0.33$ [0.23, 0.44]; (I,J) 8X: D4a3a ($p = 1.00$), $\Theta = 0.69$ [0.61, 0.76]; U4b1b1 ($p = 1.00$), $\Theta = 0.31$ [0.24, 0.39]; (K,L) 16X: D4a3a ($p = 1.00$), $\Theta = 0.68$ [0.62, 0.74]; U4b1b1 ($p = 1.00$), $\Theta = 0.32$ [0.26, 0.38]; (M,N) 32X: D4a3a ($p = 1.00$), $\Theta = 0.70$ [0.66, 0.74]; U4b1b1 ($p = 0.99$), $\Theta = 0.30$ [0.26, 0.34].

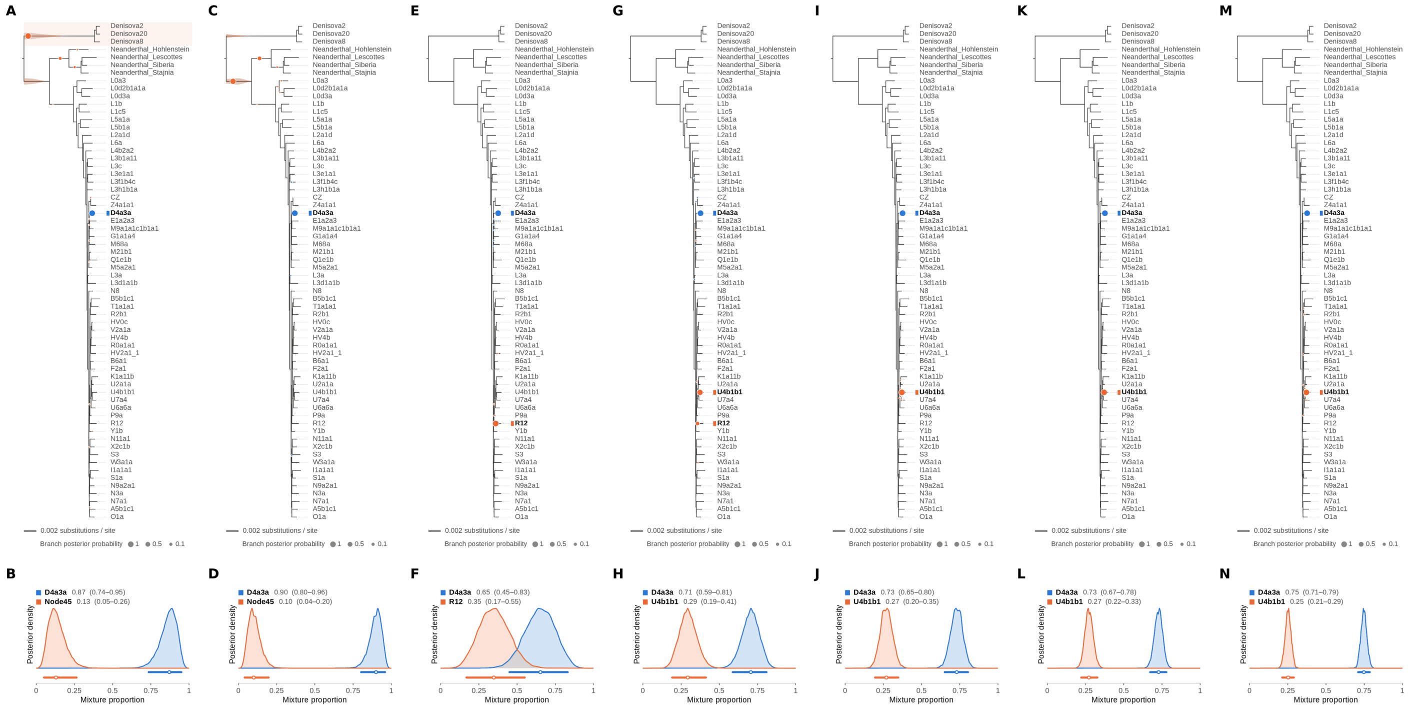

Figure S71: **Two-source simulated mixture of modern human D4a1d (FJ951517.1) (85% of reads) and modern human U5a2a (HM229344.1; 15%) at nominal mean coverage depths of 0.5X, 1X, 2X, 4X, 8X, 16X, 32X.** Both sources carry ancient DNA damage (U5a2a with the damage rates scaled by 0.75). For each coverage depth, the upper panel shows the phylogenetic placement posterior of both sources for the best of five chains ($k = 2$, 100,000 iterations each), colored by source (dot area: branch posterior probability; swelling: posterior density of the position along the branch; bold tips: highest-posterior branches), and the lower panel the posterior densities of the mixture proportions $\Theta$ with median and 95% credible interval. D4a1d is expected on the D4a3a branch (its closest relative in the reference); U5a2a, for which no U5 haplotype is present in the reference, is expected within the U clade. Modal branch of each source in the best chain ($p$: its posterior probability) and $\Theta$ median [95% CI]: (A,B) 0.5X: D4a3a ($p = 1.00$), $\Theta = 0.87$ [0.74, 0.95]; Node45 (stem of the Denisovan clade; $p = 0.90$), $\Theta = 0.13$ [0.05, 0.26]; (C,D) 1X: D4a3a ($p = 1.00$), $\Theta = 0.90$ [0.80, 0.96]; Node45 (stem of the Denisovan clade; $p = 0.79$), $\Theta = 0.10$ [0.04, 0.20]; (E,F) 2X: D4a3a ($p = 1.00$), $\Theta = 0.65$ [0.45, 0.83]; R12 ($p = 1.00$), $\Theta = 0.35$ [0.17, 0.55]; (G,H) 4X: D4a3a ($p = 1.00$), $\Theta = 0.71$ [0.59, 0.81]; U4b1b1 ($p = 0.83$), $\Theta = 0.29$ [0.19, 0.41]; (I,J) 8X: D4a3a ($p = 1.00$), $\Theta = 0.73$ [0.65, 0.80]; U4b1b1 ($p = 1.00$), $\Theta = 0.27$ [0.20, 0.35]; (K,L) 16X: D4a3a ($p = 1.00$), $\Theta = 0.73$ [0.67, 0.78]; U4b1b1 ($p = 1.00$), $\Theta = 0.27$ [0.22, 0.33]; (M,N) 32X: D4a3a ($p = 1.00$), $\Theta = 0.75$ [0.71, 0.79]; U4b1b1 ($p = 1.00$), $\Theta = 0.25$ [0.21, 0.29].

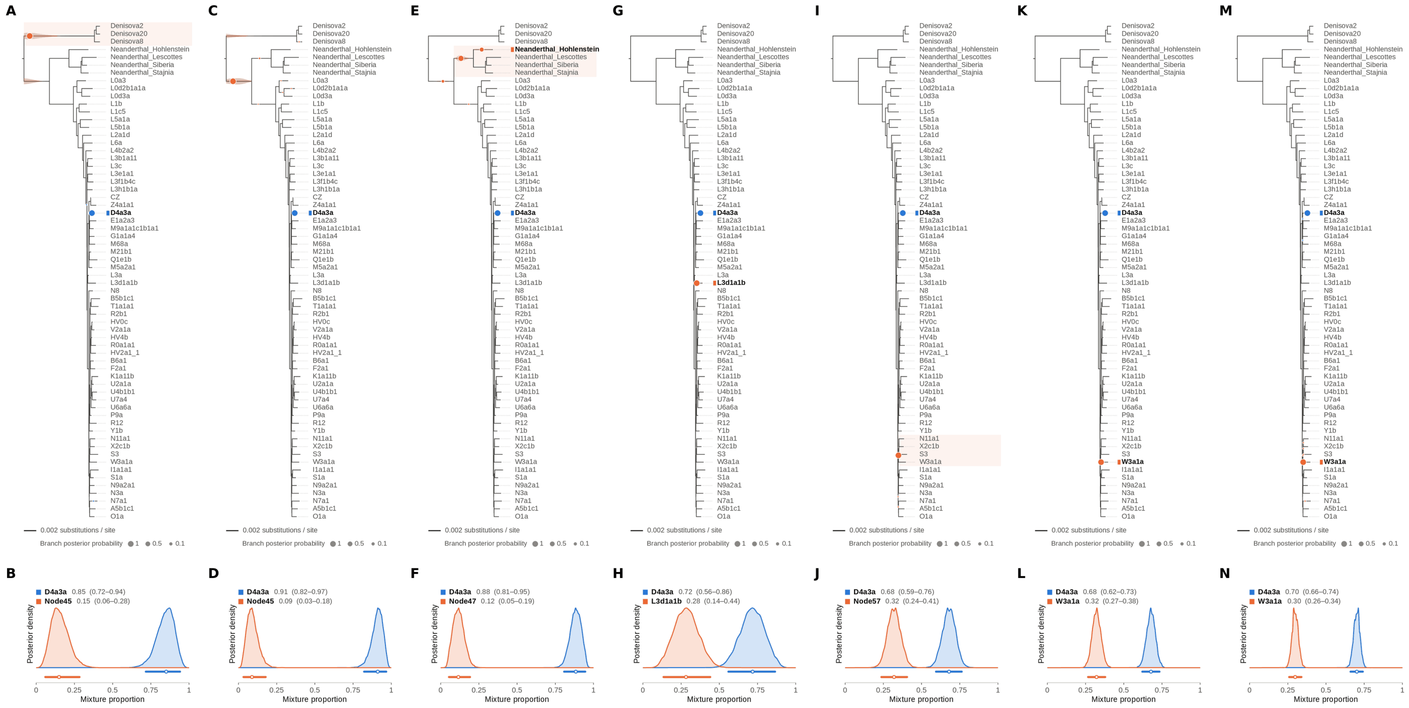


Figure S72: **Two-source simulated mixture of modern human D4a1d (FJ951517.1) (90% of reads) and modern human U5a2a (HM229344.1; 10%) at nominal mean coverage depths of 0.5X, 1X, 2X, 4X, 8X, 16X, 32X.** Both sources carry ancient DNA damage (U5a2a with the damage rates scaled by 0.75). For each coverage depth, the upper panel shows the phylogenetic placement posterior of both sources for the best of five chains ($k = 2$, 100,000 iterations each), colored by source (dot area: branch posterior probability; swelling: posterior density of the position along the branch; bold tips: highest-posterior branches), and the lower panel the posterior densities of the mixture proportions $\Theta$ with median and 95% credible interval. D4a1d is expected on the D4a3a branch (its closest relative in the reference); U5a2a, for which no U5 haplotype is present in the reference, is expected within the U clade. Modal branch of each source in the best chain ($p$: its posterior probability) and $\Theta$ median [95% CI]: (A,B) 0.5X: D4a3a ($p = 1.00$), $\Theta = 0.85$ [0.72, 0.94]; Node45 (stem of the Denisovan clade; $p = 1.00$), $\Theta = 0.15$ [0.06, 0.28]; (C,D) 1X: D4a3a ($p = 1.00$), $\Theta = 0.91$ [0.82, 0.97]; Node45 (stem of the Denisovan clade; $p = 1.00$), $\Theta = 0.09$ [0.03, 0.18]; (E,F) 2X: D4a3a ($p = 1.00$), $\Theta = 0.88$ [0.81, 0.95]; Node47 (stem of the Neanderthal clade; $p = 0.70$), $\Theta = 0.12$ [0.05, 0.19]; (G,H) 4X: D4a3a ($p = 1.00$), $\Theta = 0.72$ [0.56, 0.86]; L3d1a1b ($p = 1.00$), $\Theta = 0.28$ [0.14, 0.44]; (I,J) 8X: D4a3a ($p = 1.00$), $\Theta = 0.68$ [0.59, 0.76]; Node57 (deep internal branch; $p = 1.00$), $\Theta = 0.32$ [0.24, 0.41]; (K,L) 16X: D4a3a ($p = 1.00$), $\Theta = 0.68$ [0.62, 0.73]; W3a1a ($p = 1.00$), $\Theta = 0.32$ [0.27, 0.38]; (M,N) 32X: D4a3a ($p = 1.00$), $\Theta = 0.70$ [0.66, 0.74]; W3a1a ($p = 1.00$), $\Theta = 0.30$ [0.26, 0.34].

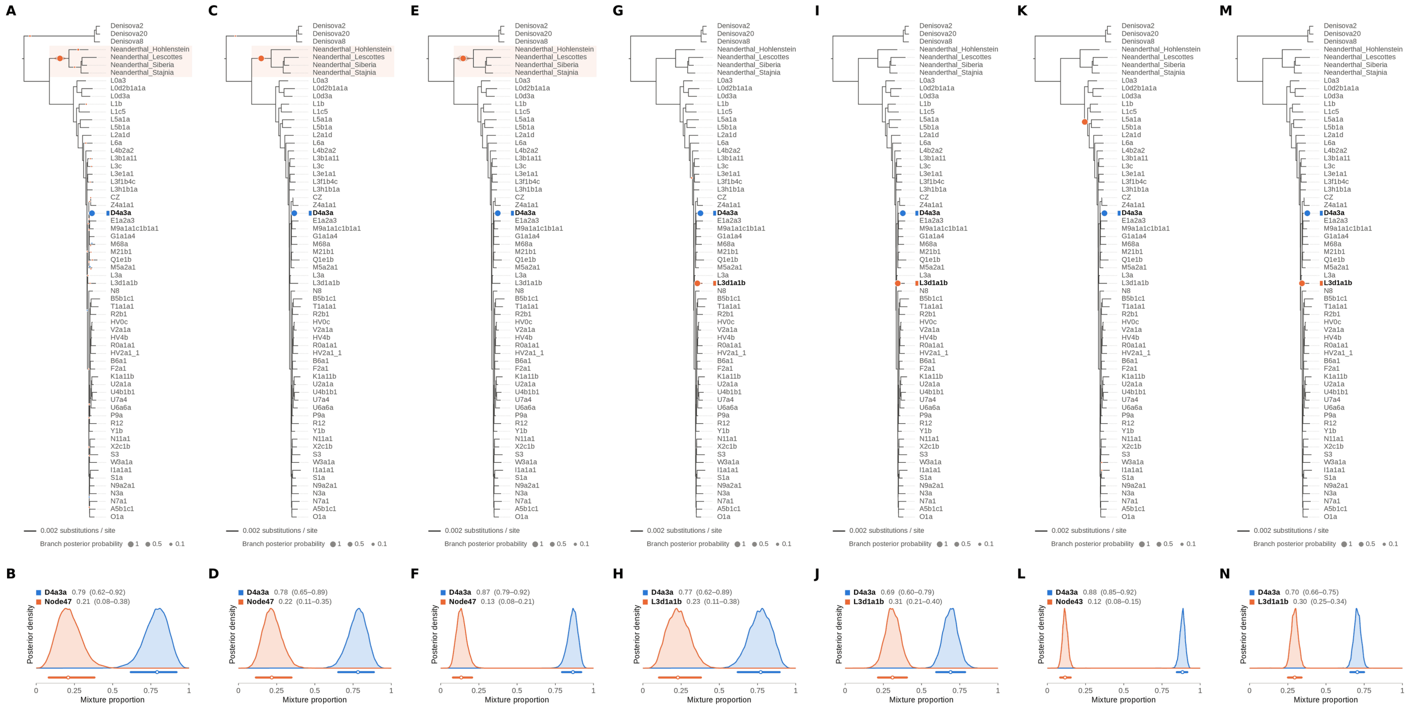

Figure S73: **Two-source simulated mixture of modern human D4a1d (FJ951517.1) (95% of reads) and modern human U5a2a (HM229344.1; 5%) at nominal mean coverage depths of 0.5X, 1X, 2X, 4X, 8X, 16X, 32X.** Both sources carry ancient DNA damage (U5a2a with the damage rates scaled by 0.75). For each coverage depth, the upper panel shows the phylogenetic placement posterior of both sources for the best of five chains ($k = 2$, 100,000 iterations each), colored by source (dot area: branch posterior probability; swelling: posterior density of the position along the branch; bold tips: highest-posterior branches), and the lower panel the posterior densities of the mixture proportions $\Theta$ with median and 95% credible interval. D4a1d is expected on the D4a3a branch (its closest relative in the reference); U5a2a, for which no U5 haplotype is present in the reference, is expected within the U clade. Modal branch of each source in the best chain ($p$: its posterior probability) and $\Theta$ median [95% CI]: (A,B) 0.5X: D4a3a ($p = 1.00$), $\Theta = 0.79$ [0.62, 0.92]; Node47 (stem of the Neanderthal clade; $p = 0.98$), $\Theta = 0.21$ [0.08, 0.38]; (C,D) 1X: D4a3a ($p = 1.00$), $\Theta = 0.78$ [0.65, 0.89]; Node47 (stem of the Neanderthal clade; $p = 1.00$), $\Theta = 0.22$ [0.11, 0.35]; (E,F) 2X: D4a3a ($p = 1.00$), $\Theta = 0.87$ [0.79, 0.92]; Node47 (stem of the Neanderthal clade; $p = 1.00$), $\Theta = 0.13$ [0.08, 0.21]; (G,H) 4X: D4a3a ($p = 1.00$), $\Theta = 0.77$ [0.62, 0.89]; L3d1a1b ($p = 1.00$), $\Theta = 0.23$ [0.11, 0.38]; (I,J) 8X: D4a3a ($p = 1.00$), $\Theta = 0.69$ [0.60, 0.79]; L3d1a1b ($p = 1.00$), $\Theta = 0.31$ [0.21, 0.40]; (K,L) 16X: D4a3a ($p = 1.00$), $\Theta = 0.88$ [0.85, 0.92]; Node43 (deep internal branch; $p = 1.00$), $\Theta = 0.12$ [0.08, 0.15]; (M,N) 32X: D4a3a ($p = 1.00$), $\Theta = 0.70$ [0.66, 0.75]; L3d1a1b ($p = 1.00$), $\Theta = 0.30$ [0.25, 0.34].

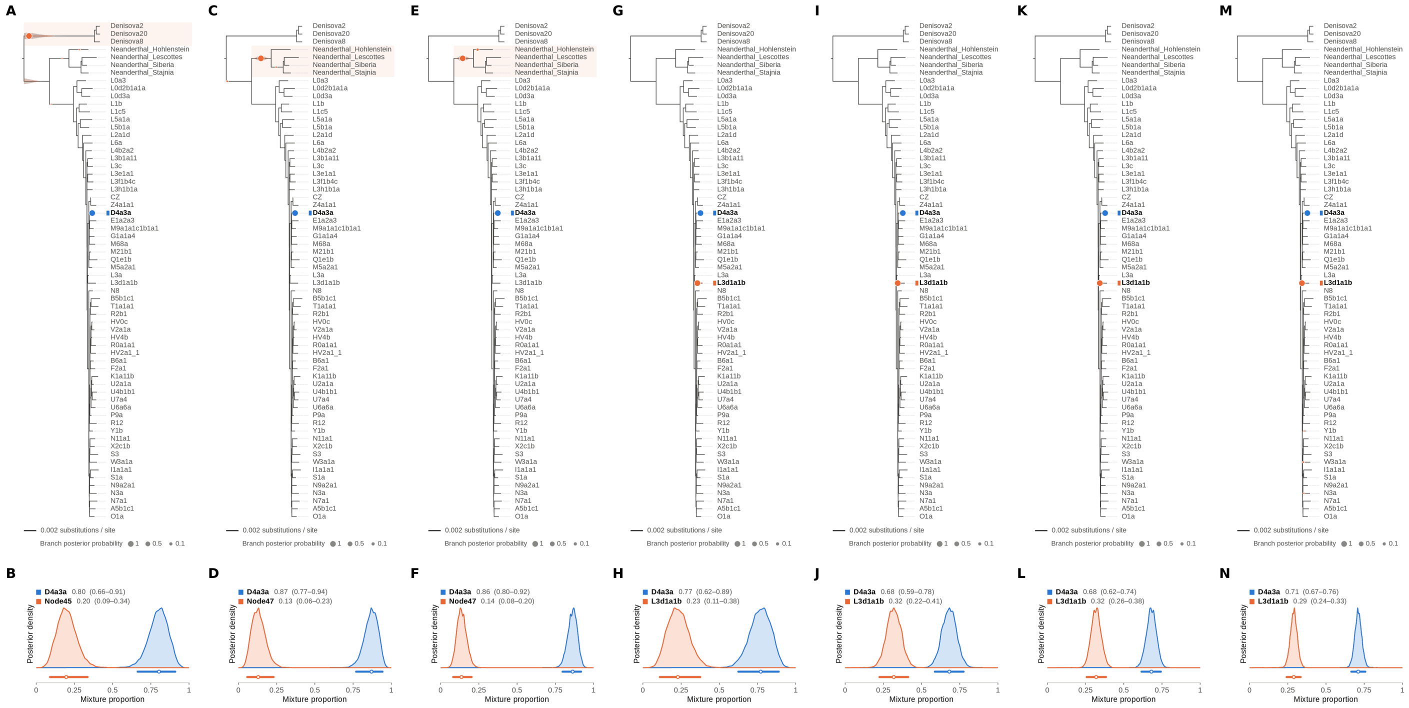


Figure S74: **Two-source simulated mixture of modern human D4a1d (FJ951517.1) (99% of reads) and modern human U5a2a (HM229344.1; 1%) at nominal mean coverage depths of 0.5X, 1X, 2X, 4X, 8X, 16X, 32X.** Both sources carry ancient DNA damage (U5a2a with the damage rates scaled by 0.75). For each coverage depth, the upper panel shows the phylogenetic placement posterior of both sources for the best of five chains ($k = 2$, 100,000 iterations each), colored by source (dot area: branch posterior probability; swelling: posterior density of the position along the branch; bold tips: highest-posterior branches), and the lower panel the posterior densities of the mixture proportions $\Theta$ with median and 95% credible interval. D4a1d is expected on the D4a3a branch (its closest relative in the reference); U5a2a, for which no U5 haplotype is present in the reference, is expected within the U clade. Modal branch of each source in the best chain ($p$: its posterior probability) and $\Theta$ median [95% CI]: (A,B) 0.5X: D4a3a ($p = 1.00$), $\Theta = 0.80$ [0.66, 0.91]; Node45 (stem of the Denisovan clade; $p = 1.00$), $\Theta = 0.20$ [0.09, 0.34]; (C,D) 1X: D4a3a ($p = 1.00$), $\Theta = 0.87$ [0.77, 0.94]; Node47 (stem of the Neanderthal clade; $p = 0.99$), $\Theta = 0.13$ [0.06, 0.23]; (E,F) 2X: D4a3a ($p = 1.00$), $\Theta = 0.86$ [0.80, 0.92]; Node47 (stem of the Neanderthal clade; $p = 0.97$), $\Theta = 0.14$ [0.08, 0.20]; (G,H) 4X: D4a3a ($p = 1.00$), $\Theta = 0.77$ [0.62, 0.89]; L3d1a1b ($p = 1.00$), $\Theta = 0.23$ [0.11, 0.38]; (I,J) 8X: D4a3a ($p = 1.00$), $\Theta = 0.68$ [0.59, 0.78]; L3d1a1b ($p = 1.00$), $\Theta = 0.32$ [0.22, 0.41]; (K,L) 16X: D4a3a ($p = 1.00$), $\Theta = 0.68$ [0.62, 0.74]; L3d1a1b ($p = 1.00$), $\Theta = 0.32$ [0.26, 0.38]; (M,N) 32X: D4a3a ($p = 1.00$), $\Theta = 0.71$ [0.67, 0.76]; L3d1a1b ($p = 0.99$), $\Theta = 0.29$ [0.24, 0.33].

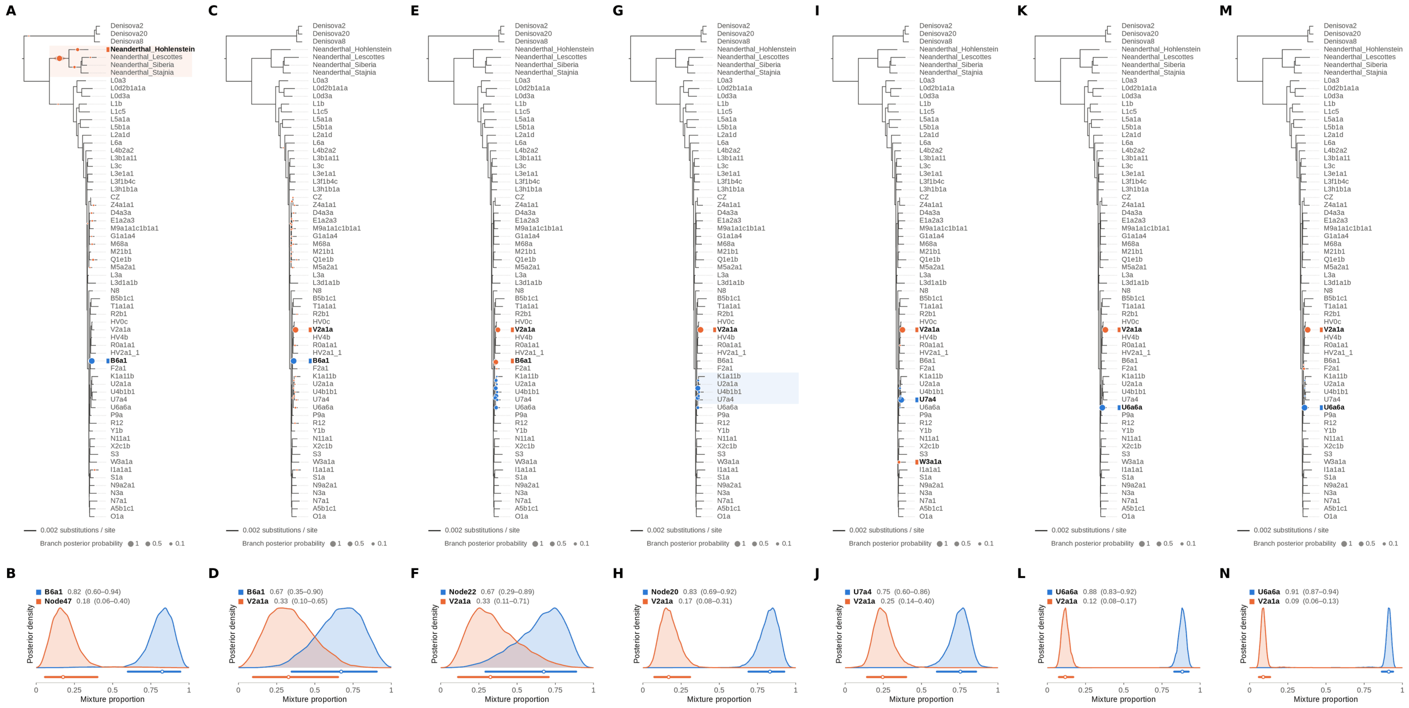


Figure S75: **Two-source simulated mixture of modern human L2 (AF381981.1) (1% of reads) and modern human U5a2a (HM229344.1; 99%) at nominal mean coverage depths of 0.5X, 1X, 2X, 4X, 8X, 16X, 32X.** Both sources carry ancient DNA damage (U5a2a with the damage rates scaled by 0.75). For each coverage depth, the upper panel shows the phylogenetic placement posterior of both sources for the best of five chains ($k = 2$, 100,000 iterations each), colored by source (dot area: branch posterior probability; swelling: posterior density of the position along the branch; bold tips: highest-posterior branches), and the lower panel the posterior densities of the mixture proportions $\Theta$ with median and 95% credible interval. L2 is expected on the L2a1d branch, the only L2 haplotype in the reference; U5a2a, for which no U5 haplotype is present in the reference, is expected within the U clade. Modal branch of each source in the best chain ($p$: its posterior probability) and $\Theta$ median [95% CI]: (A,B) 0.5X: B6a1 ($p = 1.00$), $\Theta = 0.82$ [0.60, 0.94]; Node47 (stem of the Neanderthal clade; $p = 0.83$), $\Theta = 0.18$ [0.06, 0.40]; (C,D) 1X: B6a1 ($p = 1.00$), $\Theta = 0.67$ [0.35, 0.90]; V2a1a ($p = 0.95$), $\Theta = 0.33$ [0.10, 0.65]; (E,F) 2X: Node22 (internal branch ancestral to U4b1b1 and U7a4; $p = 0.29$), $\Theta = 0.67$ [0.29, 0.89]; V2a1a ($p = 0.52$), $\Theta = 0.33$ [0.11, 0.71]; (G,H) 4X: Node20 (internal branch within the U clade; $p = 0.60$), $\Theta = 0.83$ [0.69, 0.92]; V2a1a ($p = 1.00$), $\Theta = 0.17$ [0.08, 0.31]; (I,J) 8X: U7a4 ($p = 0.98$), $\Theta = 0.75$ [0.60, 0.86]; V2a1a ($p = 0.94$), $\Theta = 0.25$ [0.14, 0.40]; (K,L) 16X: U6a6a ($p = 1.00$), $\Theta = 0.88$ [0.83, 0.92]; V2a1a ($p = 1.00$), $\Theta = 0.12$ [0.08, 0.17]; (M,N) 32X: U6a6a ($p = 1.00$), $\Theta = 0.91$ [0.87, 0.94]; V2a1a ($p = 0.98$), $\Theta = 0.09$ [0.06, 0.13].

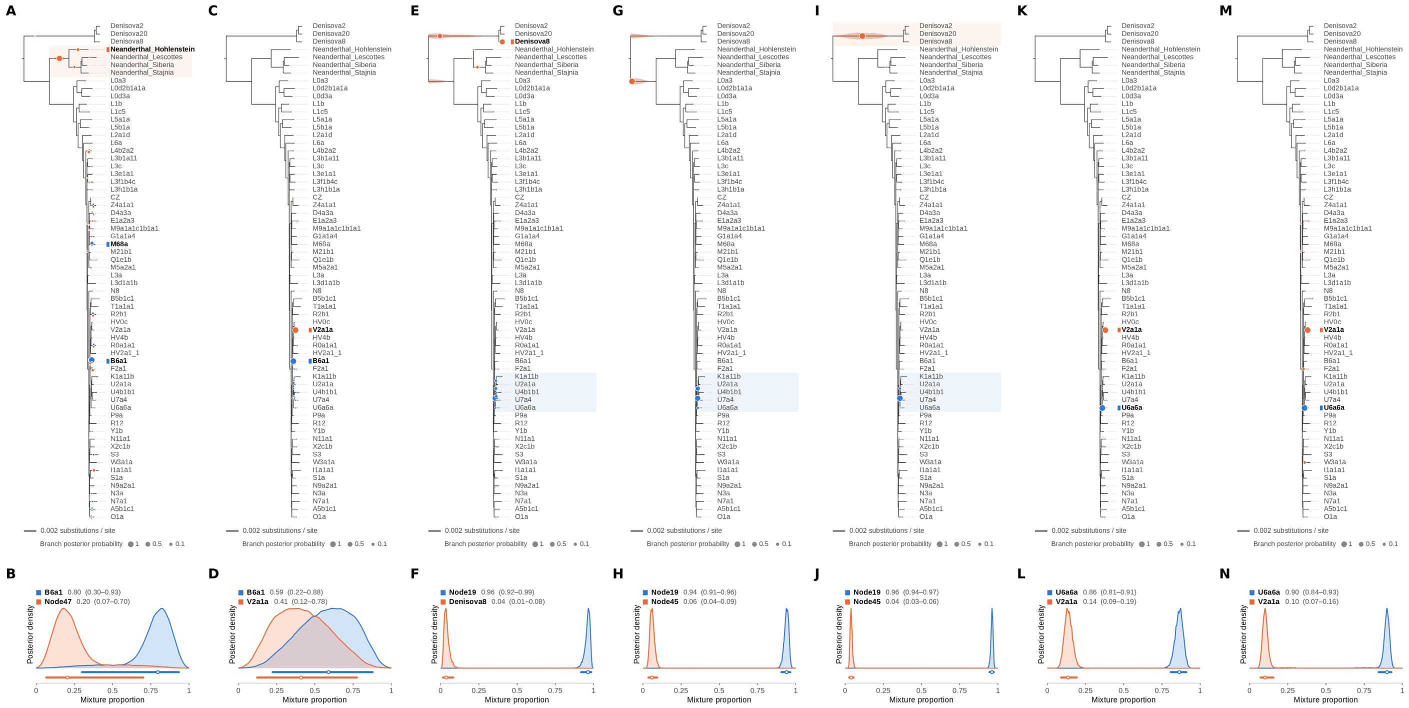


Figure S76: **Two-source simulated mixture of modern human L2 (AF381981.1) (5% of reads) and modern human U5a2a (HM229344.1; 95%) at nominal mean coverage depths of 0.5X, 1X, 2X, 4X, 8X, 16X, 32X.** Both sources carry ancient DNA damage (U5a2a with the damage rates scaled by 0.75). For each coverage depth, the upper panel shows the phylogenetic placement posterior of both sources for the best of five chains ($k = 2$, 100,000 iterations each), colored by source (dot area: branch posterior probability; swelling: posterior density of the position along the branch; bold tips: highest-posterior branches), and the lower panel the posterior densities of the mixture proportions $\Theta$ with median and 95% credible interval. L2 is expected on the L2a1d branch, the only L2 haplotype in the reference; U5a2a, for which no U5 haplotype is present in the reference, is expected within the U clade. Modal branch of each source in the best chain ($p$: its posterior probability) and $\Theta$ median [95% CI]: (A,B) 0.5X: B6a1 ($p = 0.88$), $\Theta = 0.80$ [0.30, 0.93]; Node47 (stem of the Neanderthal clade; $p = 0.76$), $\Theta = 0.20$ [0.07, 0.70]; (C,D) 1X: B6a1 ($p = 0.99$), $\Theta = 0.59$ [0.22, 0.88]; V2a1a ($p = 1.00$), $\Theta = 0.41$ [0.12, 0.78]; (E,F) 2X: Node19 (stem of the U clade; $p = 0.83$), $\Theta = 0.96$ [0.92, 0.99]; Denisova8 ($p = 0.60$), $\Theta = 0.04$ [0.01, 0.08]; (G,H) 4X: Node19 (stem of the U clade; $p = 0.70$), $\Theta = 0.94$ [0.91, 0.96]; Node45 (stem of the Denisovan clade; $p = 1.00$), $\Theta = 0.06$ [0.04, 0.09]; (I,J) 8X: Node19 (stem of the U clade; $p = 0.97$), $\Theta = 0.96$ [0.94, 0.97]; Node45 (stem of the Denisovan clade; $p = 1.00$), $\Theta = 0.04$ [0.03, 0.06]; (K,L) 16X: U6a6a ($p = 1.00$), $\Theta = 0.86$ [0.81, 0.91]; V2a1a ($p = 1.00$), $\Theta = 0.14$ [0.09, 0.19]; (M,N) 32X: U6a6a ($p = 1.00$), $\Theta = 0.90$ [0.84, 0.93]; V2a1a ($p = 0.94$), $\Theta = 0.10$ [0.07, 0.16].

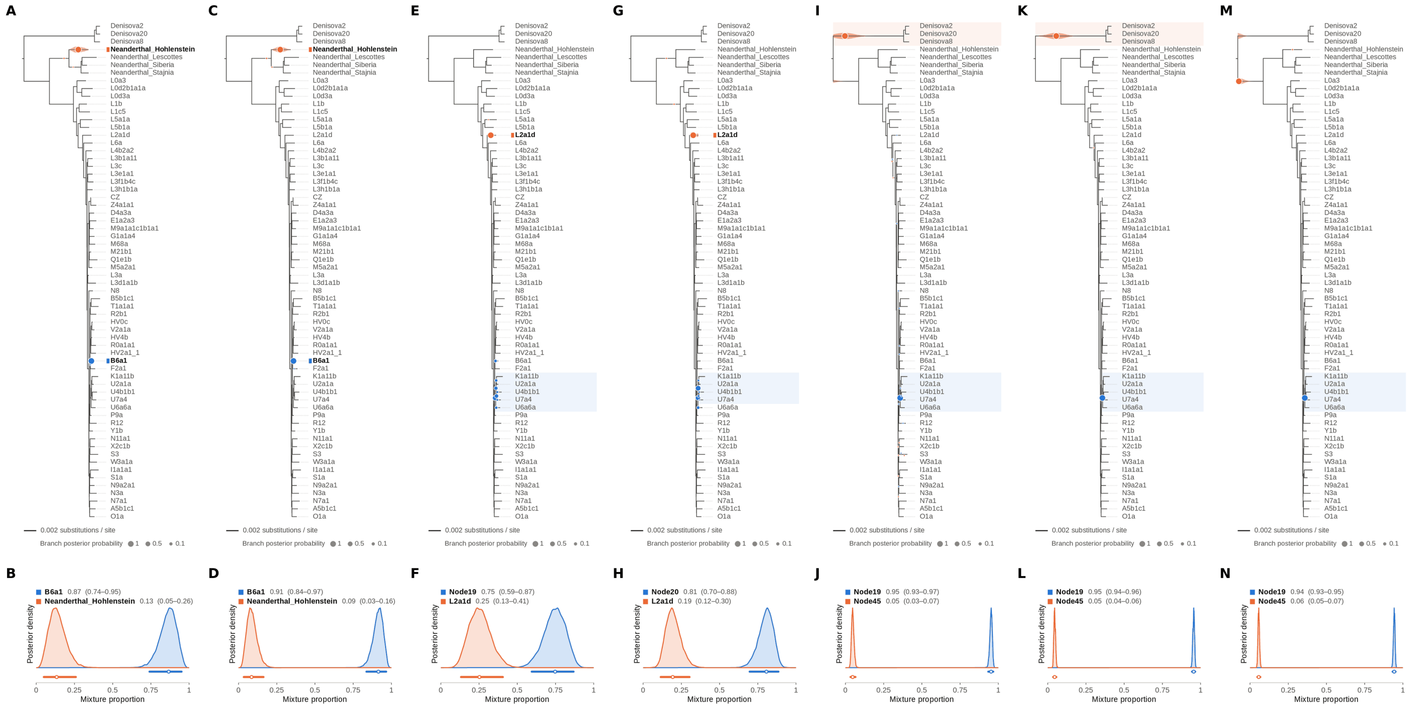


Figure S77: **Two-source simulated mixture of modern human L2 (AF381981.1) (10% of reads) and modern human U5a2a (HM229344.1; 90%) at nominal mean coverage depths of 0.5X, 1X, 2X, 4X, 8X, 16X, 32X.** Both sources carry ancient DNA damage (U5a2a with the damage rates scaled by 0.75). For each coverage depth, the upper panel shows the phylogenetic placement posterior of both sources for the best of five chains ($k = 2$, 100,000 iterations each), colored by source (dot area: branch posterior probability; swelling: posterior density of the position along the branch; bold tips: highest-posterior branches), and the lower panel the posterior densities of the mixture proportions $\Theta$ with median and 95% credible interval. L2 is expected on the L2a1d branch, the only L2 haplotype in the reference; U5a2a, for which no U5 haplotype is present in the reference, is expected within the U clade. Modal branch of each source in the best chain ($p$: its posterior probability) and $\Theta$ median [95% CI]: (A,B) 0.5X: B6a1 ($p = 1.00$), $\Theta = 0.87$ [0.74, 0.95]; Neanderthal_Hohlenstein ($p = 1.00$), $\Theta = 0.13$ [0.05, 0.26]; (C,D) 1X: B6a1 ($p = 1.00$), $\Theta = 0.91$ [0.84, 0.97]; Neanderthal_Hohlenstein ($p = 1.00$), $\Theta = 0.09$ [0.03, 0.16]; (E,F) 2X: Node19 (stem of the U clade; $p = 0.56$), $\Theta = 0.75$ [0.59, 0.87]; L2a1d ($p = 1.00$), $\Theta = 0.25$ [0.13, 0.41]; (G,H) 4X: Node20 (internal branch within the U clade; $p = 0.52$), $\Theta = 0.81$ [0.70, 0.88]; L2a1d ($p = 1.00$), $\Theta = 0.19$ [0.12, 0.30]; (I,J) 8X: Node19 (stem of the U clade; $p = 1.00$), $\Theta = 0.95$ [0.93, 0.97]; Node45 (stem of the Denisovan clade; $p = 1.00$), $\Theta = 0.05$ [0.03, 0.07]; (K,L) 16X: Node19 (stem of the U clade; $p = 1.00$), $\Theta = 0.95$ [0.94, 0.96]; Node45 (stem of the Denisovan clade; $p = 1.00$), $\Theta = 0.05$ [0.04, 0.06]; (M,N) 32X: Node19 (stem of the U clade; $p = 1.00$), $\Theta = 0.94$ [0.93, 0.95]; Node45 (stem of the Denisovan clade; $p = 1.00$), $\Theta = 0.06$ [0.05, 0.07].

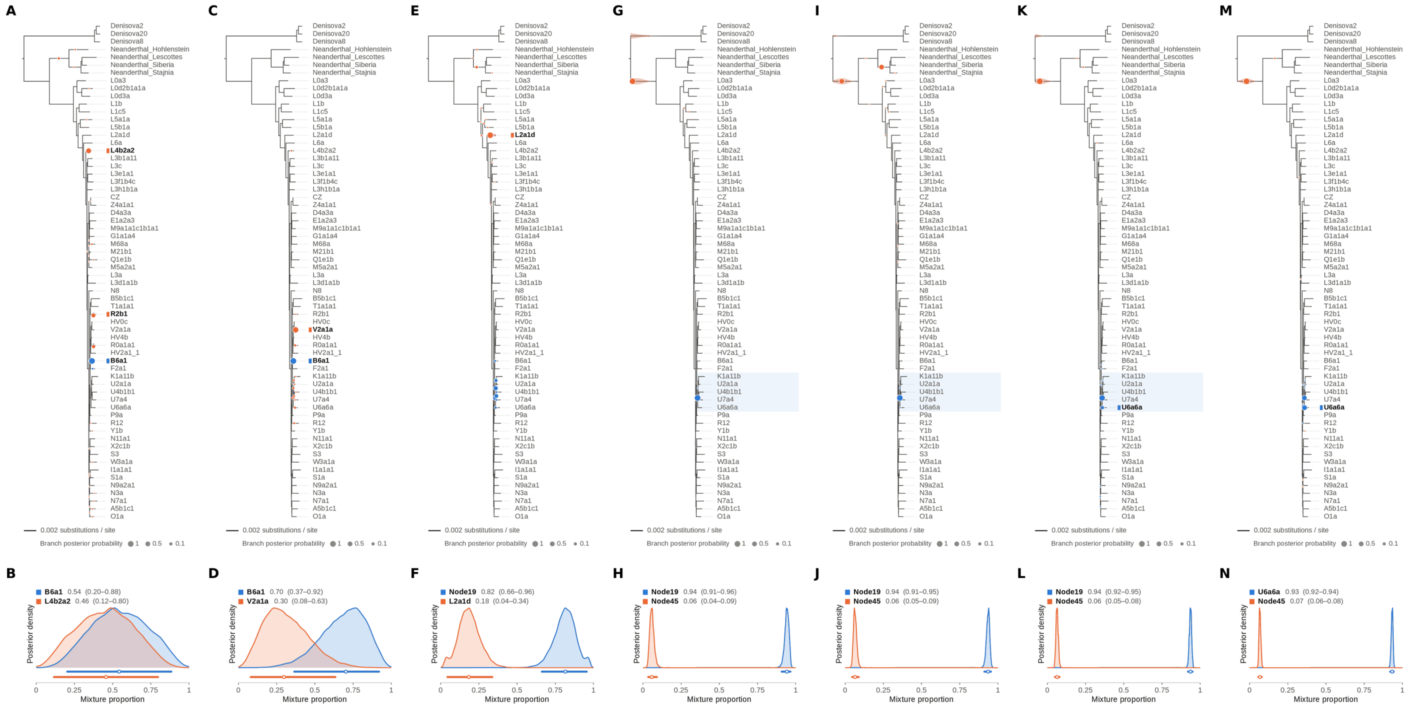

Figure S78: **Two-source simulated mixture of modern human L2 (AF381981.1) (15% of reads) and modern human U5a2a (HM229344.1; 85%) at nominal mean coverage depths of 0.5X, 1X, 2X, 4X, 8X, 16X, 32X.** Both sources carry ancient DNA damage (U5a2a with the damage rates scaled by 0.75). For each coverage depth, the upper panel shows the phylogenetic placement posterior of both sources for the best of five chains ($k = 2$, 100,000 iterations each), colored by source (dot area: branch posterior probability; swelling: posterior density of the position along the branch; bold tips: highest-posterior branches), and the lower panel the posterior densities of the mixture proportions $\Theta$ with median and 95% credible interval. L2 is expected on the L2a1d branch, the only L2 haplotype in the reference; U5a2a, for which no U5 haplotype is present in the reference, is expected within the U clade. Modal branch of each source in the best chain ($p$: its posterior probability) and $\Theta$ median [95% CI]: (A,B) 0.5X: B6a1 ($p = 0.99$), $\Theta = 0.54$ [0.20, 0.88]; L4b2a2 ($p = 0.57$), $\Theta = 0.46$ [0.12, 0.80]; (C,D) 1X: B6a1 ($p = 1.00$), $\Theta = 0.70$ [0.37, 0.92]; V2a1a ($p = 0.88$), $\Theta = 0.30$ [0.08, 0.63]; (E,F) 2X: Node19 (stem of the U clade; $p = 0.32$), $\Theta = 0.82$ [0.66, 0.96]; L2a1d ($p = 0.94$), $\Theta = 0.18$ [0.04, 0.34]; (G,H) 4X: Node19 (stem of the U clade; $p = 1.00$), $\Theta = 0.94$ [0.91, 0.96]; Node45 (stem of the Denisovan clade; $p = 1.00$), $\Theta = 0.06$ [0.04, 0.09]; (I,J) 8X: Node19 (stem of the U clade; $p = 1.00$), $\Theta = 0.94$ [0.91, 0.95]; Node45 (stem of the Denisovan clade; $p = 0.51$), $\Theta = 0.06$ [0.05, 0.09]; (K,L) 16X: Node19 (stem of the U clade; $p = 0.87$), $\Theta = 0.94$ [0.92, 0.95]; Node45 (stem of the Denisovan clade; $p = 1.00$), $\Theta = 0.06$ [0.05, 0.08]; (M,N) 32X: U6a6a ($p = 0.52$), $\Theta = 0.93$ [0.92, 0.94]; Node45 (stem of the Denisovan clade; $p = 1.00$), $\Theta = 0.07$ [0.06, 0.08].

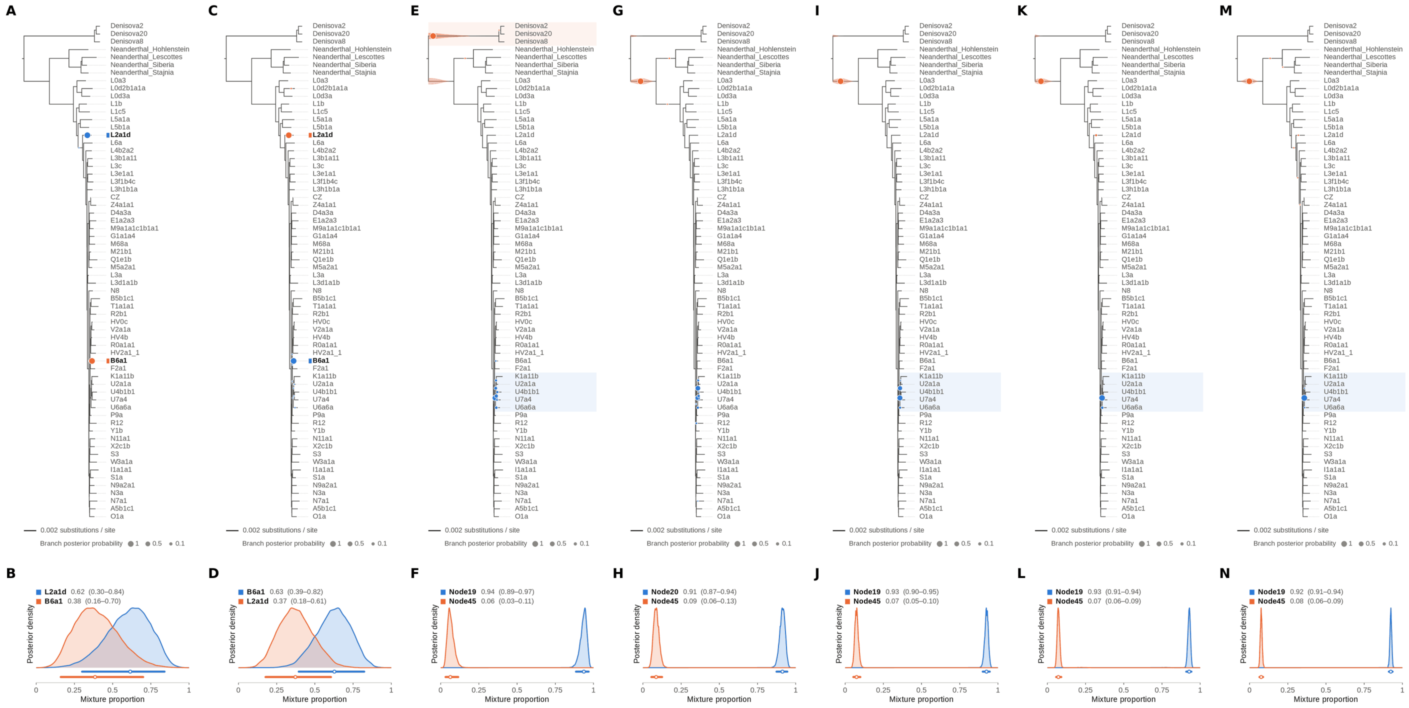


Figure S79: **Two-source simulated mixture of modern human L2 (AF381981.1) (20% of reads) and modern human U5a2a (HM229344.1; 80%) at nominal mean coverage depths of 0.5X, 1X, 2X, 4X, 8X, 16X, 32X.** Both sources carry ancient DNA damage (U5a2a with the damage rates scaled by 0.75). For each coverage depth, the upper panel shows the phylogenetic placement posterior of both sources for the best of five chains ($k$ = 2, 100,000 iterations each), colored by source (dot area: branch posterior probability; swelling: posterior density of the position along the branch; bold tips: highest-posterior branches), and the lower panel the posterior densities of the mixture proportions $\Theta$ with median and 95% credible interval. L2 is expected on the L2a1d branch, the only L2 haplotype in the reference; U5a2a, for which no U5 haplotype is present in the reference, is expected within the U clade. Modal branch of each source in the best chain ($p$: its posterior probability) and $\Theta$ median [95% CI]: (A,B) 0.5X: L2a1d ($p$ = 1.00), $\Theta$ = 0.62 [0.30, 0.84]; B6a1 ($p$ = 1.00), $\Theta$ = 0.38 [0.16, 0.70]; (C,D) 1X: B6a1 ($p$ = 1.00), $\Theta$ = 0.63 [0.39, 0.82]; L2a1d ($p$ = 1.00), $\Theta$ = 0.37 [0.18, 0.61]; (E,F) 2X: Node19 (stem of the U clade; $p$ = 0.69), $\Theta$ = 0.94 [0.89, 0.97]; Node45 (stem of the Denisovan clade; $p$ = 1.00), $\Theta$ = 0.06 [0.03, 0.11]; (G,H) 4X: Node20 (internal branch within the U clade; $p$ = 0.49), $\Theta$ = 0.91 [0.87, 0.94]; Node45 (stem of the Denisovan clade; $p$ = 1.00), $\Theta$ = 0.09 [0.06, 0.13]; (I,J) 8X: Node19 (stem of the U clade; $p$ = 0.64), $\Theta$ = 0.93 [0.90, 0.95]; Node45 (stem of the Denisovan clade; $p$ = 1.00), $\Theta$ = 0.07 [0.05, 0.10]; (K,L) 16X: Node19 (stem of the U clade; $p$ = 0.96), $\Theta$ = 0.93 [0.91, 0.94]; Node45 (stem of the Denisovan clade; $p$ = 0.99), $\Theta$ = 0.07 [0.06, 0.09]; (M,N) 32X: Node19 (stem of the U clade; $p$ = 1.00), $\Theta$ = 0.92 [0.91, 0.94]; Node45 (stem of the Denisovan clade; $p$ = 1.00), $\Theta$ = 0.08 [0.06, 0.09].

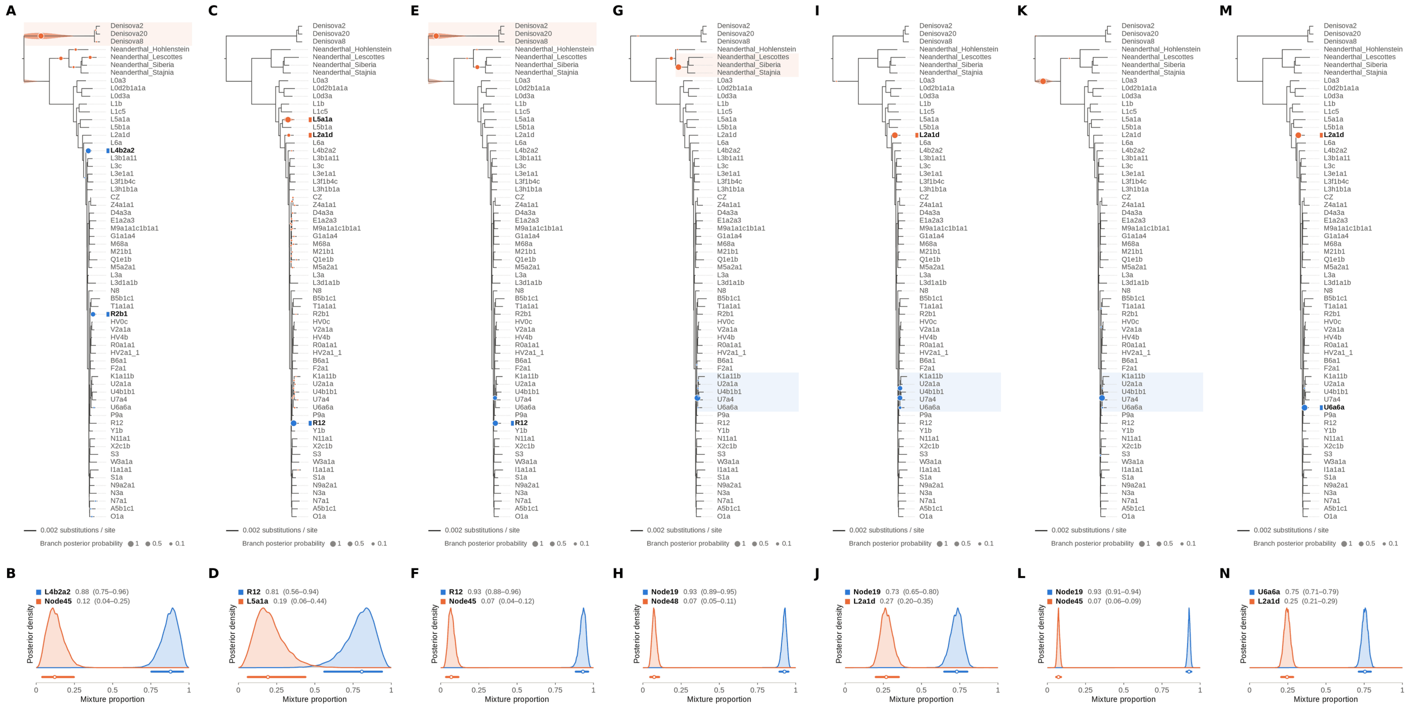


Figure S80: **Two-source simulated mixture of modern human L2 (AF381981.1) (25% of reads) and modern human U5a2a (HM229344.1; 75%) at nominal mean coverage depths of 0.5X, 1X, 2X, 4X, 8X, 16X, 32X.** Both sources carry ancient DNA damage (U5a2a with the damage rates scaled by 0.75). For each coverage depth, the upper panel shows the phylogenetic placement posterior of both sources for the best of five chains ($k = 2$, 100,000 iterations each), colored by source (dot area: branch posterior probability; swelling: posterior density of the position along the branch; bold tips: highest-posterior branches), and the lower panel the posterior densities of the mixture proportions $\Theta$ with median and 95% credible interval. L2 is expected on the L2a1d branch, the only L2 haplotype in the reference; U5a2a, for which no U5 haplotype is present in the reference, is expected within the U clade. Modal branch of each source in the best chain ($p$: its posterior probability) and $\Theta$ median [95% CI]: (A,B) 0.5X: L4b2a2 ($p = 0.61$), $\Theta = 0.88$ [0.75, 0.96]; Node45 (stem of the Denisovan clade; $p = 0.71$), $\Theta = 0.12$ [0.04, 0.25]; (C,D) 1X: R12 ($p = 1.00$), $\Theta = 0.81$ [0.56, 0.94]; L5a1a ($p = 0.88$), $\Theta = 0.19$ [0.06, 0.44]; (E,F) 2X: R12 ($p = 0.74$), $\Theta = 0.93$ [0.88, 0.96]; Node45 (stem of the Denisovan clade; $p = 0.74$), $\Theta = 0.07$ [0.04, 0.12]; (G,H) 4X: Node19 (stem of the U clade; $p = 1.00$), $\Theta = 0.93$ [0.89, 0.95]; Node48 (internal branch ancestral to Neanderthal_Lescottes, Neanderthal_Siberia and Neanderthal_Stajnia; $p = 0.90$), $\Theta = 0.07$ [0.05, 0.11]; (I,J) 8X: Node19 (stem of the U clade; $p = 0.53$), $\Theta = 0.73$ [0.65, 0.80]; L2a1d ($p = 1.00$), $\Theta = 0.27$ [0.20, 0.35]; (K,L) 16X: Node19 (stem of the U clade; $p = 1.00$), $\Theta = 0.93$ [0.91, 0.94]; Node45 (stem of the Denisovan clade; $p = 1.00$), $\Theta = 0.07$ [0.06, 0.09]; (M,N) 32X: U6a6a ($p = 1.00$), $\Theta = 0.75$ [0.71, 0.79]; L2a1d ($p = 1.00$), $\Theta = 0.25$ [0.21, 0.29].

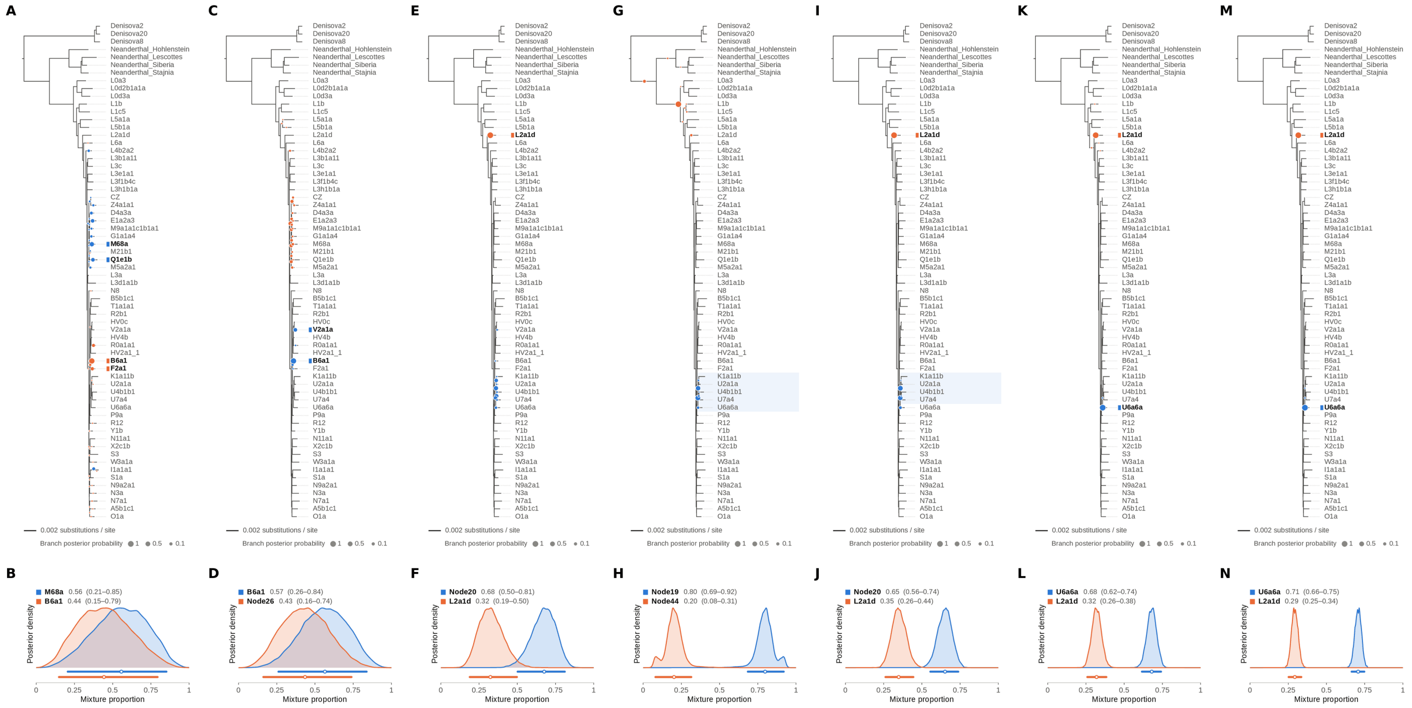


Figure S81: **Two-source simulated mixture of modern human L2 (AF381981.1) (30% of reads) and modern human U5a2a (HM229344.1; 70%) at nominal mean coverage depths of 0.5X, 1X, 2X, 4X, 8X, 16X, 32X.** Both sources carry ancient DNA damage (U5a2a with the damage rates scaled by 0.75). For each coverage depth, the upper panel shows the phylogenetic placement posterior of both sources for the best of five chains ($k = 2$, 100,000 iterations each), colored by source (dot area: branch posterior probability; swelling: posterior density of the position along the branch; bold tips: highest-posterior branches), and the lower panel the posterior densities of the mixture proportions $\Theta$ with median and 95% credible interval. L2 is expected on the L2a1d branch, the only L2 haplotype in the reference; U5a2a, for which no U5 haplotype is present in the reference, is expected within the U clade. Modal branch of each source in the best chain ($p$: its posterior probability) and $\Theta$ median [95% CI]: (A,B) 0.5X: M68a ($p = 0.35$), $\Theta = 0.56$ [0.21, 0.85]; B6a1 ($p = 0.75$), $\Theta = 0.44$ [0.15, 0.79]; (C,D) 1X: B6a1 ($p = 0.78$), $\Theta = 0.57$ [0.26, 0.84]; Node26 (stem of the M clade; $p = 0.28$), $\Theta = 0.43$ [0.16, 0.74]; (E,F) 2X: Node20 (internal branch within the U clade; $p = 0.33$), $\Theta = 0.68$ [0.50, 0.81]; L2a1d ($p = 1.00$), $\Theta = 0.32$ [0.19, 0.50]; (G,H) 4X: Node19 (stem of the U clade; $p = 0.54$), $\Theta = 0.80$ [0.69, 0.92]; Node44 (stem of the archaic hominin clade; $p = 0.89$), $\Theta = 0.20$ [0.08, 0.31]; (I,J) 8X: Node20 (internal branch within the U clade; $p = 0.51$), $\Theta = 0.65$ [0.56, 0.74]; L2a1d ($p = 1.00$), $\Theta = 0.35$ [0.26, 0.44]; (K,L) 16X: U6a6a ($p = 1.00$), $\Theta = 0.68$ [0.62, 0.74]; L2a1d ($p = 1.00$), $\Theta = 0.32$ [0.26, 0.38]; (M,N) 32X: U6a6a ($p = 1.00$), $\Theta = 0.71$ [0.66, 0.75]; L2a1d ($p = 1.00$), $\Theta = 0.29$ [0.25, 0.34].

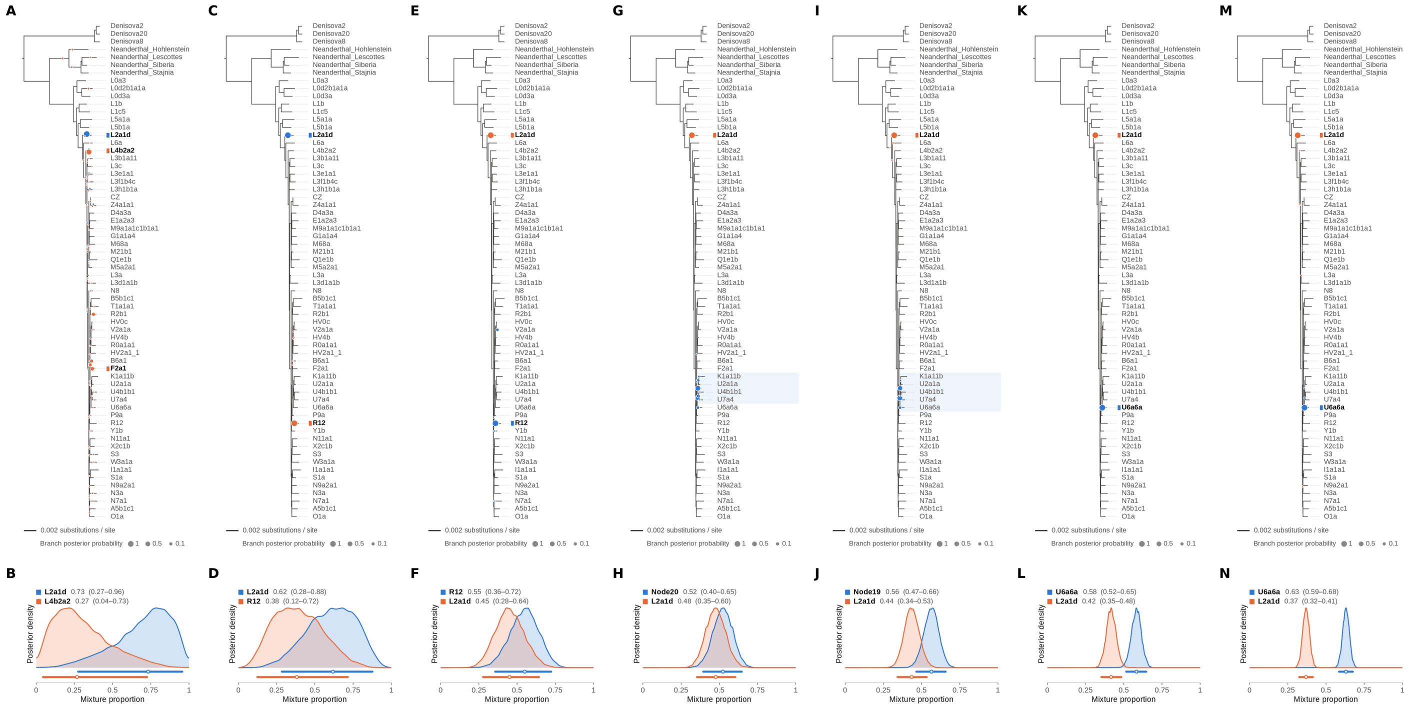

Figure S82: **Two-source simulated mixture of modern human L2 (AF381981.1) (35% of reads) and modern human U5a2a (HM229344.1; 65%) at nominal mean coverage depths of 0.5X, 1X, 2X, 4X, 8X, 16X, 32X.** Both sources carry ancient DNA damage (U5a2a with the damage rates scaled by 0.75). For each coverage depth, the upper panel shows the phylogenetic placement posterior of both sources for the best of five chains ($k = 2$, 100,000 iterations each), colored by source (dot area: branch posterior probability; swelling: posterior density of the position along the branch; bold tips: highest-posterior branches), and the lower panel the posterior densities of the mixture proportions $\Theta$ with median and 95% credible interval. L2 is expected on the L2a1d branch, the only L2 haplotype in the reference; U5a2a, for which no U5 haplotype is present in the reference, is expected within the U clade. Modal branch of each source in the best chain ($p$: its posterior probability) and $\Theta$ median [95% CI]: (A,B) 0.5X: L2a1d ($p = 0.98$), $\Theta = 0.73$ [0.27, 0.96]; L4b2a2 ($p = 0.56$), $\Theta = 0.27$ [0.04, 0.73]; (C,D) 1X: L2a1d ($p = 1.00$), $\Theta = 0.62$ [0.28, 0.88]; R12 ($p = 1.00$), $\Theta = 0.38$ [0.12, 0.72]; (E,F) 2X: R12 ($p = 0.95$), $\Theta = 0.55$ [0.36, 0.72]; L2a1d ($p = 1.00$), $\Theta = 0.45$ [0.28, 0.64]; (G,H) 4X: Node20 (internal branch within the U clade; $p = 0.51$), $\Theta = 0.52$ [0.40, 0.65]; L2a1d ($p = 1.00$), $\Theta = 0.48$ [0.35, 0.60]; (I,J) 8X: Node19 (stem of the U clade; $p = 0.57$), $\Theta = 0.56$ [0.47, 0.66]; L2a1d ($p = 1.00$), $\Theta = 0.44$ [0.34, 0.53]; (K,L) 16X: U6a6a ($p = 1.00$), $\Theta = 0.58$ [0.52, 0.65]; L2a1d ($p = 1.00$), $\Theta = 0.42$ [0.35, 0.48]; (M,N) 32X: U6a6a ($p = 1.00$), $\Theta = 0.63$ [0.59, 0.68]; L2a1d ($p = 1.00$), $\Theta = 0.37$ [0.32, 0.41].

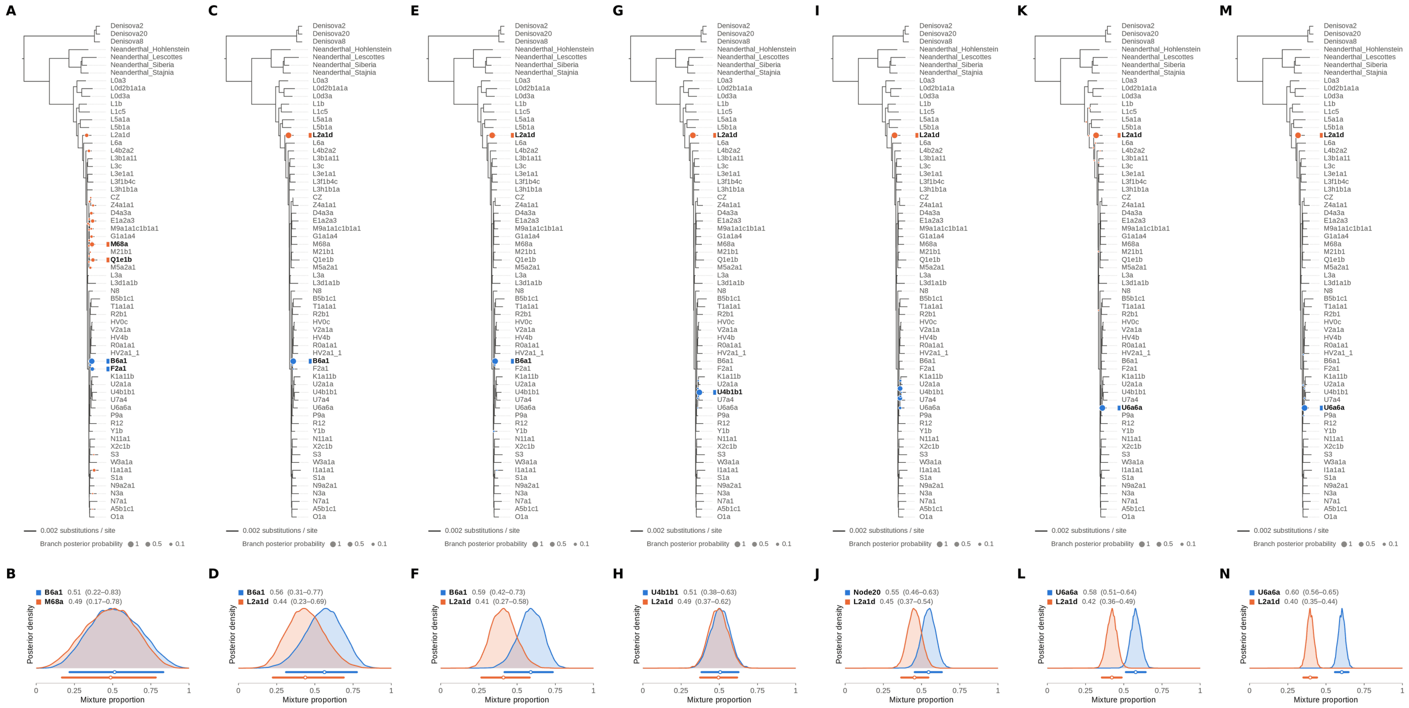

Figure S83: **Two-source simulated mixture of modern human L2 (AF381981.1) (40% of reads) and modern human U5a2a (HM229344.1; 60%) at nominal mean coverage depths of 0.5X, 1X, 2X, 4X, 8X, 16X, 32X.** Both sources carry ancient DNA damage (U5a2a with the damage rates scaled by 0.75). For each coverage depth, the upper panel shows the phylogenetic placement posterior of both sources for the best of five chains ($k = 2$, 100,000 iterations each), colored by source (dot area: branch posterior probability; swelling: posterior density of the position along the branch; bold tips: highest-posterior branches), and the lower panel the posterior densities of the mixture proportions $\Theta$ with median and 95% credible interval. L2 is expected on the L2a1d branch, the only L2 haplotype in the reference; U5a2a, for which no U5 haplotype is present in the reference, is expected within the U clade. Modal branch of each source in the best chain ($p$: its posterior probability) and $\Theta$ median [95% CI]: (A,B) 0.5X: B6a1 ($p = 0.81$), $\Theta = 0.51$ [0.22, 0.83]; M68a ($p = 0.29$), $\Theta = 0.49$ [0.17, 0.78]; (C,D) 1X: B6a1 ($p = 1.00$), $\Theta = 0.56$ [0.31, 0.77]; L2a1d ($p = 1.00$), $\Theta = 0.44$ [0.23, 0.69]; (E,F) 2X: B6a1 ($p = 0.99$), $\Theta = 0.59$ [0.42, 0.73]; L2a1d ($p = 1.00$), $\Theta = 0.41$ [0.27, 0.58]; (G,H) 4X: U4b1b1 ($p = 0.99$), $\Theta = 0.51$ [0.38, 0.63]; L2a1d ($p = 1.00$), $\Theta = 0.49$ [0.37, 0.62]; (I,J) 8X: Node20 (internal branch within the U clade; $p = 0.50$), $\Theta = 0.55$ [0.46, 0.63]; L2a1d ($p = 1.00$), $\Theta = 0.45$ [0.37, 0.54]; (K,L) 16X: U6a6a ($p = 1.00$), $\Theta = 0.58$ [0.51, 0.64]; L2a1d ($p = 1.00$), $\Theta = 0.42$ [0.36, 0.49]; (M,N) 32X: U6a6a ($p = 1.00$), $\Theta = 0.60$ [0.56, 0.65]; L2a1d ($p = 1.00$), $\Theta = 0.40$ [0.35, 0.44].

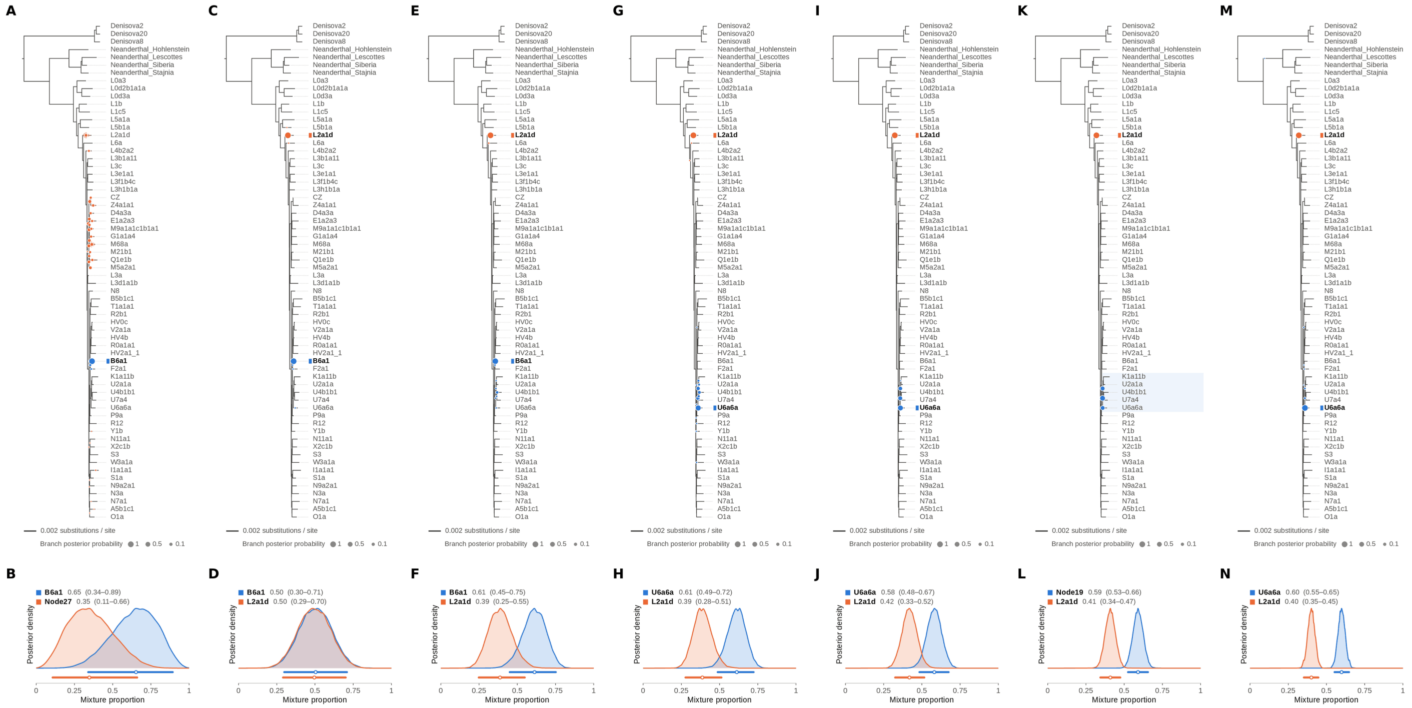


Figure S84: **Two-source simulated mixture of modern human L2 (AF381981.1) (45% of reads) and modern human U5a2a (HM229344.1; 55%) at nominal mean coverage depths of 0.5X, 1X, 2X, 4X, 8X, 16X, 32X.** Both sources carry ancient DNA damage (U5a2a with the damage rates scaled by 0.75). For each coverage depth, the upper panel shows the phylogenetic placement posterior of both sources for the best of five chains ($k = 2$, 100,000 iterations each), colored by source (dot area: branch posterior probability; swelling: posterior density of the position along the branch; bold tips: highest-posterior branches), and the lower panel the posterior densities of the mixture proportions $\Theta$ with median and 95% credible interval. L2 is expected on the L2a1d branch, the only L2 haplotype in the reference; U5a2a, for which no U5 haplotype is present in the reference, is expected within the U clade. Modal branch of each source in the best chain ($p$: its posterior probability) and $\Theta$ median [95% CI]: (A,B) 0.5X: B6a1 ($p = 0.98$), $\Theta = 0.65$ [0.34, 0.89]; Node27 (internal branch ancestral to CZ and Z4a1a1; $p = 0.09$), $\Theta = 0.35$ [0.11, 0.66]; (C,D) 1X: B6a1 ($p = 1.00$), $\Theta = 0.50$ [0.30, 0.71]; L2a1d ($p = 1.00$), $\Theta = 0.50$ [0.29, 0.70]; (E,F) 2X: B6a1 ($p = 0.94$), $\Theta = 0.61$ [0.45, 0.75]; L2a1d ($p = 1.00$), $\Theta = 0.39$ [0.25, 0.55]; (G,H) 4X: U6a6a ($p = 0.63$), $\Theta = 0.61$ [0.49, 0.72]; L2a1d ($p = 1.00$), $\Theta = 0.39$ [0.28, 0.51]; (I,J) 8X: U6a6a ($p = 0.58$), $\Theta = 0.58$ [0.48, 0.67]; L2a1d ($p = 1.00$), $\Theta = 0.42$ [0.33, 0.52]; (K,L) 16X: Node19 (stem of the U clade; $p = 0.52$), $\Theta = 0.59$ [0.53, 0.66]; L2a1d ($p = 1.00$), $\Theta = 0.41$ [0.34, 0.47]; (M,N) 32X: U6a6a ($p = 0.98$), $\Theta = 0.60$ [0.55, 0.65]; L2a1d ($p = 1.00$), $\Theta = 0.40$ [0.35, 0.45].

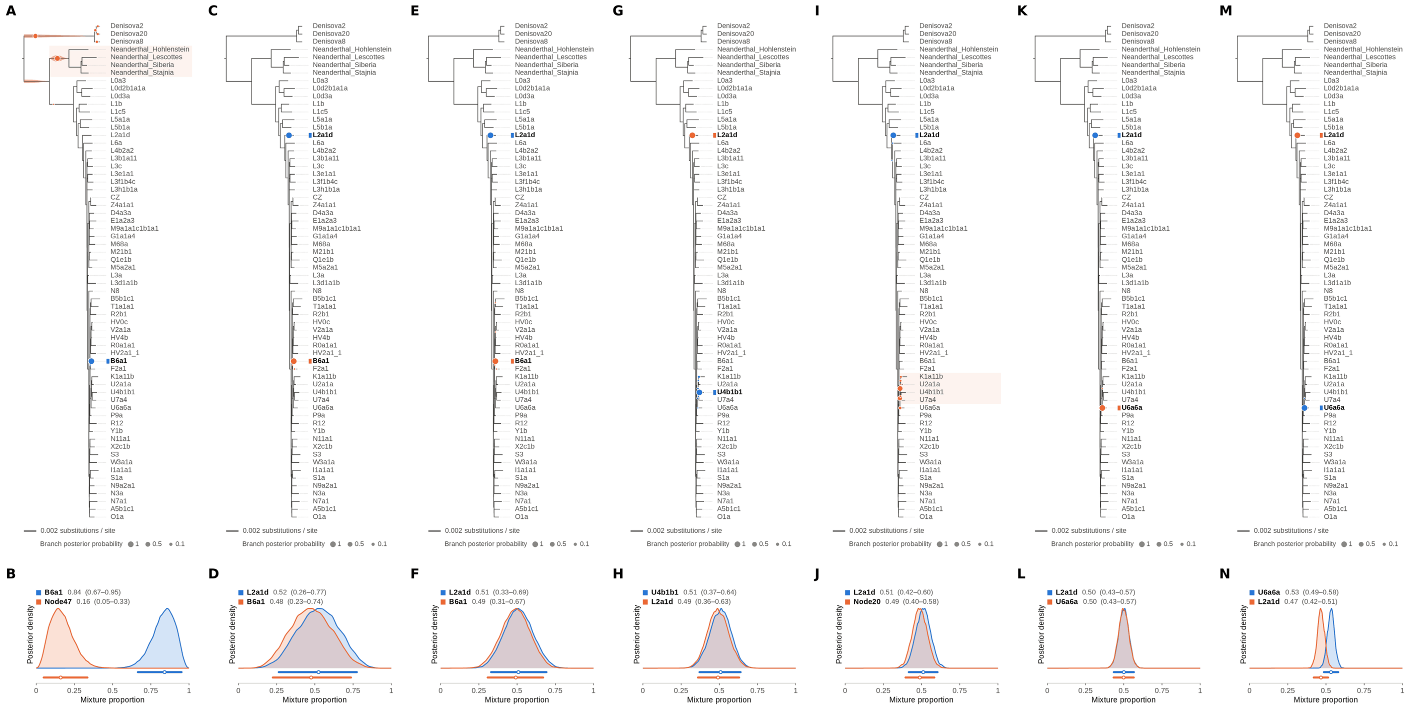


Figure S85: **Two-source simulated mixture of modern human L2 (AF381981.1) (50% of reads) and modern human U5a2a (HM229344.1; 50%) at nominal mean coverage depths of 0.5X, 1X, 2X, 4X, 8X, 16X, 32X.** Both sources carry ancient DNA damage (U5a2a with the damage rates scaled by 0.75). For each coverage depth, the upper panel shows the phylogenetic placement posterior of both sources for the best of five chains ($k = 2$, 100,000 iterations each), colored by source (dot area: branch posterior probability; swelling: posterior density of the position along the branch; bold tips: highest-posterior branches), and the lower panel the posterior densities of the mixture proportions $\Theta$ with median and 95% credible interval. L2 is expected on the L2a1d branch, the only L2 haplotype in the reference; U5a2a, for which no U5 haplotype is present in the reference, is expected within the U clade. Modal branch of each source in the best chain ($p$: its posterior probability) and $\Theta$ median [95% CI]: (A,B) 0.5X: B6a1 ($p = 1.00$), $\Theta = 0.84$ [0.67, 0.95]; Node47 (stem of the Neanderthal clade; $p = 0.57$), $\Theta = 0.16$ [0.05, 0.33]; (C,D) 1X: L2a1d ($p = 1.00$), $\Theta = 0.52$ [0.26, 0.77]; B6a1 ($p = 1.00$), $\Theta = 0.48$ [0.23, 0.74]; (E,F) 2X: L2a1d ($p = 1.00$), $\Theta = 0.51$ [0.33, 0.69]; B6a1 ($p = 1.00$), $\Theta = 0.49$ [0.31, 0.67]; (G,H) 4X: U4b1b1 ($p = 0.97$), $\Theta = 0.51$ [0.37, 0.64]; L2a1d ($p = 1.00$), $\Theta = 0.49$ [0.36, 0.63]; (I,J) 8X: L2a1d ($p = 1.00$), $\Theta = 0.51$ [0.42, 0.60]; Node20 (internal branch within the U clade; $p = 0.55$), $\Theta = 0.49$ [0.40, 0.58]; (K,L) 16X: L2a1d ($p = 1.00$), $\Theta = 0.50$ [0.43, 0.57]; U6a6a ($p = 1.00$), $\Theta = 0.50$ [0.43, 0.57]; (M,N) 32X: U6a6a ($p = 1.00$), $\Theta = 0.53$ [0.49, 0.58]; L2a1d ($p = 1.00$), $\Theta = 0.47$ [0.42, 0.51].

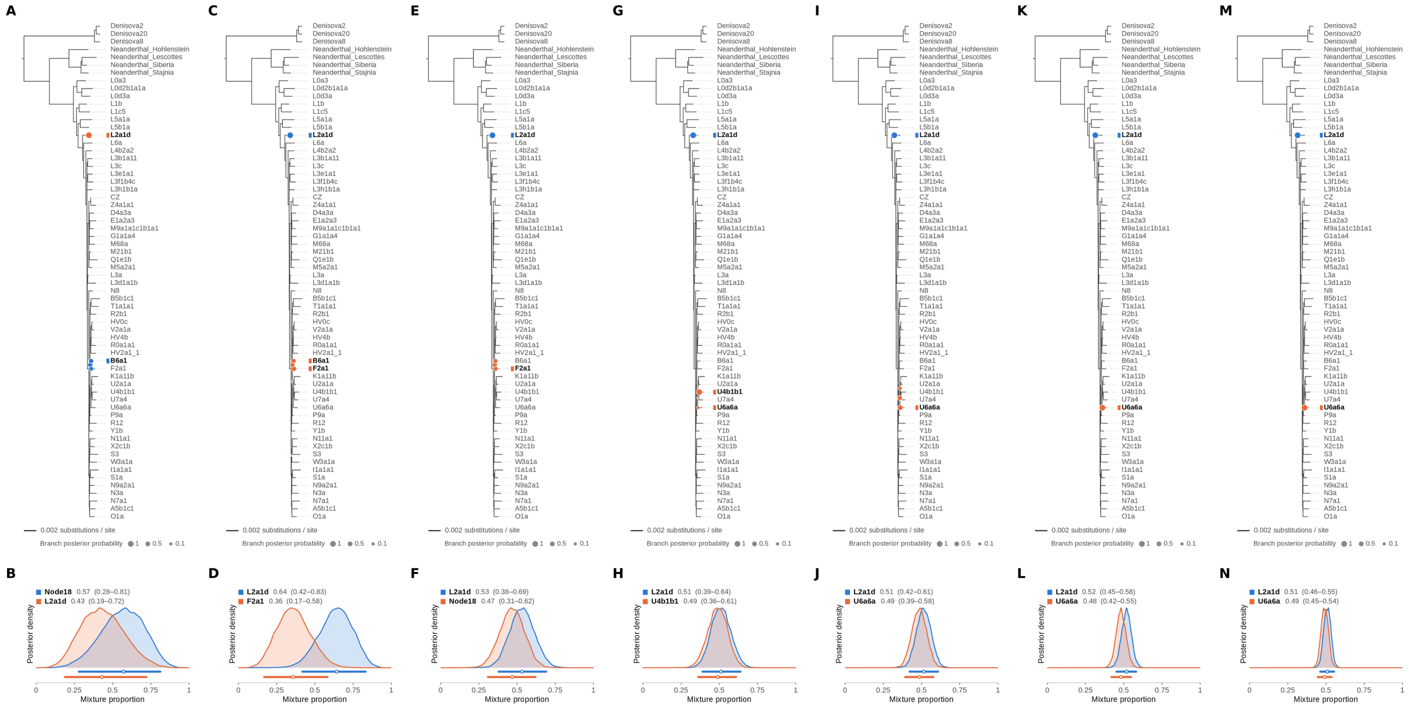


Figure S86: **Two-source simulated mixture of modern human L2 (AF381981.1) (55% of reads) and modern human U5a2a (HM229344.1; 45%) at nominal mean coverage depths of 0.5X, 1X, 2X, 4X, 8X, 16X, 32X.** Both sources carry ancient DNA damage (U5a2a with the damage rates scaled by 0.75). For each coverage depth, the upper panel shows the phylogenetic placement posterior of both sources for the best of five chains ($k = 2$, 100,000 iterations each), colored by source (dot area: branch posterior probability; swelling: posterior density of the position along the branch; bold tips: highest-posterior branches), and the lower panel the posterior densities of the mixture proportions $\Theta$ with median and 95% credible interval. L2 is expected on the L2a1d branch, the only L2 haplotype in the reference; U5a2a, for which no U5 haplotype is present in the reference, is expected within the U clade. Modal branch of each source in the best chain ($p$: its posterior probability) and $\Theta$ median [95% CI]: (A,B) 0.5X: Node18 (internal branch ancestral to B6a1 and F2a1; $p = 0.39$), $\Theta = 0.57$ [0.28, 0.81]; L2a1d ($p = 1.00$), $\Theta = 0.43$ [0.19, 0.72]; (C,D) 1X: L2a1d ($p = 1.00$), $\Theta = 0.64$ [0.42, 0.83]; F2a1 ($p = 0.42$), $\Theta = 0.36$ [0.17, 0.58]; (E,F) 2X: L2a1d ($p = 1.00$), $\Theta = 0.53$ [0.38, 0.69]; Node18 (internal branch ancestral to B6a1 and F2a1; $p = 0.39$), $\Theta = 0.47$ [0.31, 0.62]; (G,H) 4X: L2a1d ($p = 1.00$), $\Theta = 0.51$ [0.39, 0.64]; U4b1b1 ($p = 0.93$), $\Theta = 0.49$ [0.36, 0.61]; (I,J) 8X: L2a1d ($p = 1.00$), $\Theta = 0.51$ [0.42, 0.61]; U6a6a ($p = 0.49$), $\Theta = 0.49$ [0.39, 0.58]; (K,L) 16X: L2a1d ($p = 1.00$), $\Theta = 0.52$ [0.45, 0.58]; U6a6a ($p = 1.00$), $\Theta = 0.48$ [0.42, 0.55]; (M,N) 32X: L2a1d ($p = 1.00$), $\Theta = 0.51$ [0.46, 0.55]; U6a6a ($p = 1.00$), $\Theta = 0.49$ [0.45, 0.54].

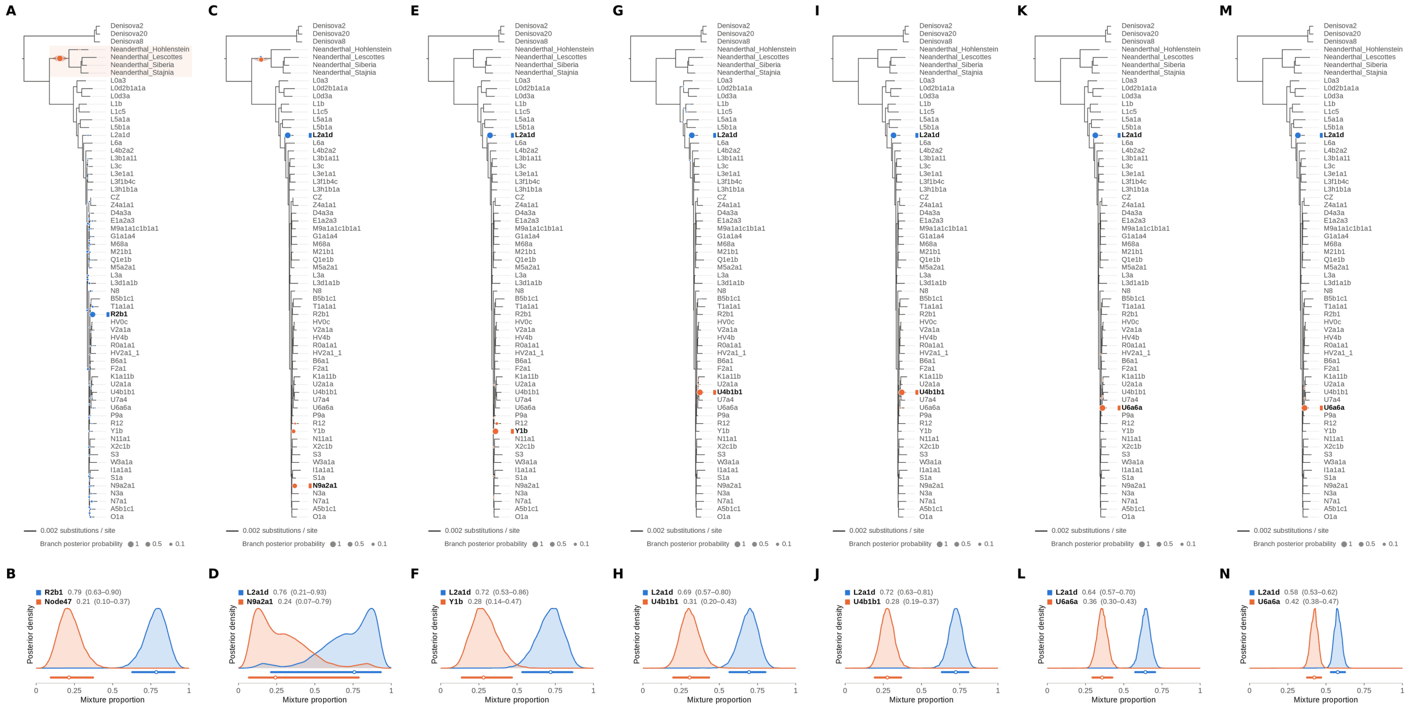

Figure S87: **Two-source simulated mixture of modern human L2 (AF381981.1) (60% of reads) and modern human U5a2a (HM229344.1; 40%) at nominal mean coverage depths of 0.5X, 1X, 2X, 4X, 8X, 16X, 32X.** Both sources carry ancient DNA damage (U5a2a with the damage rates scaled by 0.75). For each coverage depth, the upper panel shows the phylogenetic placement posterior of both sources for the best of five chains ($k$ = 2, 100,000 iterations each), colored by source (dot area: branch posterior probability; swelling: posterior density of the position along the branch; bold tips: highest-posterior branches), and the lower panel the posterior densities of the mixture proportions Θ with median and 95% credible interval. L2 is expected on the L2a1d branch, the only L2 haplotype in the reference; U5a2a, for which no U5 haplotype is present in the reference, is expected within the U clade. Modal branch of each source in the best chain ($p$: its posterior probability) and Θ median [95% CI]: (A,B) 0.5X: R2b1 ($p$ = 0.79), Θ = 0.79 [0.63, 0.90]; Node47 (stem of the Neanderthal clade; $p$ = 1.00), Θ = 0.21 [0.10, 0.37]; (C,D) 1X: L2a1d ($p$ = 0.97), Θ = 0.76 [0.21, 0.93]; N9a2a1 ($p$ = 0.42), Θ = 0.24 [0.07, 0.79]; (E,F) 2X: L2a1d ($p$ = 1.00), Θ = 0.72 [0.53, 0.86]; Y1b ($p$ = 0.95), Θ = 0.28 [0.14, 0.47]; (G,H) 4X: L2a1d ($p$ = 1.00), Θ = 0.69 [0.57, 0.80]; U4b1b1 ($p$ = 1.00), Θ = 0.31 [0.20, 0.43]; (I,J) 8X: L2a1d ($p$ = 1.00), Θ = 0.72 [0.63, 0.81]; U4b1b1 ($p$ = 1.00), Θ = 0.28 [0.19, 0.37]; (K,L) 16X: L2a1d ($p$ = 1.00), Θ = 0.64 [0.57, 0.70]; U6a6a ($p$ = 1.00), Θ = 0.36 [0.30, 0.43]; (M,N) 32X: L2a1d ($p$ = 1.00), Θ = 0.58 [0.53, 0.62]; U6a6a ($p$ = 1.00), Θ = 0.42 [0.38, 0.47].

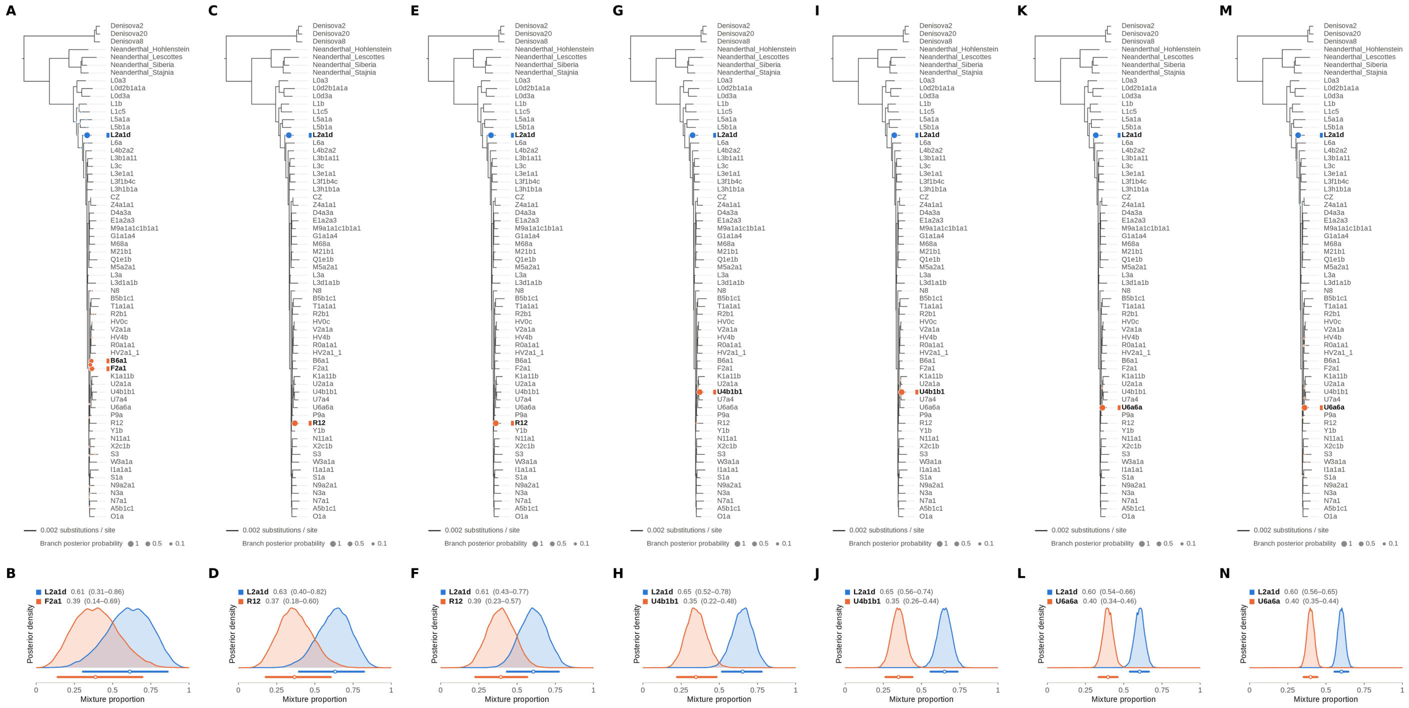


Figure S88: **Two-source simulated mixture of modern human L2 (AF381981.1) (65% of reads) and modern human U5a2a (HM229344.1; 35%) at nominal mean coverage depths of 0.5X, 1X, 2X, 4X, 8X, 16X, 32X.** Both sources carry ancient DNA damage (U5a2a with the damage rates scaled by 0.75). For each coverage depth, the upper panel shows the phylogenetic placement posterior of both sources for the best of five chains ($k = 2$, 100,000 iterations each), colored by source (dot area: branch posterior probability; swelling: posterior density of the position along the branch; bold tips: highest-posterior branches), and the lower panel the posterior densities of the mixture proportions $\Theta$ with median and 95% credible interval. L2 is expected on the L2a1d branch, the only L2 haplotype in the reference; U5a2a, for which no U5 haplotype is present in the reference, is expected within the U clade. Modal branch of each source in the best chain ($p$: its posterior probability) and $\Theta$ median [95% CI]: (A,B) 0.5X: L2a1d ($p = 1.00$), $\Theta = 0.61$ [0.31, 0.86]; F2a1 ($p = 0.46$), $\Theta = 0.39$ [0.14, 0.69]; (C,D) 1X: L2a1d ($p = 1.00$), $\Theta = 0.63$ [0.40, 0.82]; R12 ($p = 1.00$), $\Theta = 0.37$ [0.18, 0.60]; (E,F) 2X: L2a1d ($p = 1.00$), $\Theta = 0.61$ [0.43, 0.77]; R12 ($p = 1.00$), $\Theta = 0.39$ [0.23, 0.57]; (G,H) 4X: L2a1d ($p = 1.00$), $\Theta = 0.65$ [0.52, 0.78]; U4b1b1 ($p = 1.00$), $\Theta = 0.35$ [0.22, 0.48]; (I,J) 8X: L2a1d ($p = 1.00$), $\Theta = 0.65$ [0.56, 0.74]; U4b1b1 ($p = 1.00$), $\Theta = 0.35$ [0.26, 0.44]; (K,L) 16X: L2a1d ($p = 1.00$), $\Theta = 0.60$ [0.54, 0.66]; U6a6a ($p = 1.00$), $\Theta = 0.40$ [0.34, 0.46]; (M,N) 32X: L2a1d ($p = 1.00$), $\Theta = 0.60$ [0.56, 0.65]; U6a6a ($p = 1.00$), $\Theta = 0.40$ [0.35, 0.44].

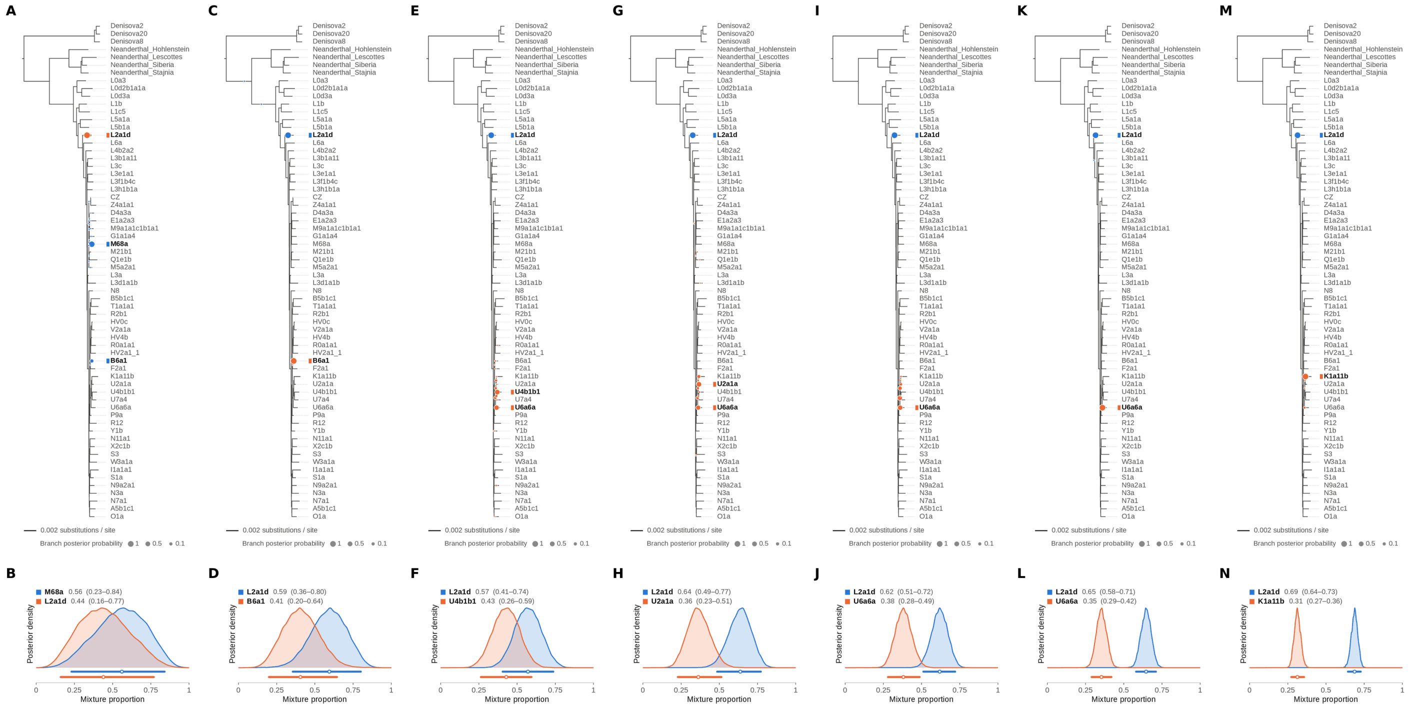

Figure S89: **Two-source simulated mixture of modern human L2 (AF381981.1) (70% of reads) and modern human U5a2a (HM229344.1; 30%) at nominal mean coverage depths of 0.5X, 1X, 2X, 4X, 8X, 16X, 32X.** Both sources carry ancient DNA damage (U5a2a with the damage rates scaled by 0.75). For each coverage depth, the upper panel shows the phylogenetic placement posterior of both sources for the best of five chains ($k = 2$, 100,000 iterations each), colored by source (dot area: branch posterior probability; swelling: posterior density of the position along the branch; bold tips: highest-posterior branches), and the lower panel the posterior densities of the mixture proportions $\Theta$ with median and 95% credible interval. L2 is expected on the L2a1d branch, the only L2 haplotype in the reference; U5a2a, for which no U5 haplotype is present in the reference, is expected within the U clade. Modal branch of each source in the best chain ($p$: its posterior probability) and $\Theta$ median [95% CI]: (A,B) 0.5X: M68a ($p = 0.88$), $\Theta = 0.56$ [0.23, 0.84]; L2a1d ($p = 1.00$), $\Theta = 0.44$ [0.16, 0.77]; (C,D) 1X: L2a1d ($p = 1.00$), $\Theta = 0.59$ [0.36, 0.80]; B6a1 ($p = 1.00$), $\Theta = 0.41$ [0.20, 0.64]; (E,F) 2X: L2a1d ($p = 1.00$), $\Theta = 0.57$ [0.41, 0.74]; U4b1b1 ($p = 0.47$), $\Theta = 0.43$ [0.26, 0.59]; (G,H) 4X: L2a1d ($p = 1.00$), $\Theta = 0.64$ [0.49, 0.77]; U2a1a ($p = 0.49$), $\Theta = 0.36$ [0.23, 0.51]; (I,J) 8X: L2a1d ($p = 1.00$), $\Theta = 0.62$ [0.51, 0.72]; U6a6a ($p = 0.41$), $\Theta = 0.38$ [0.28, 0.49]; (K,L) 16X: L2a1d ($p = 1.00$), $\Theta = 0.65$ [0.58, 0.71]; U6a6a ($p = 1.00$), $\Theta = 0.35$ [0.29, 0.42]; (M,N) 32X: L2a1d ($p = 1.00$), $\Theta = 0.69$ [0.64, 0.73]; K1a11b ($p = 0.98$), $\Theta = 0.31$ [0.27, 0.36].

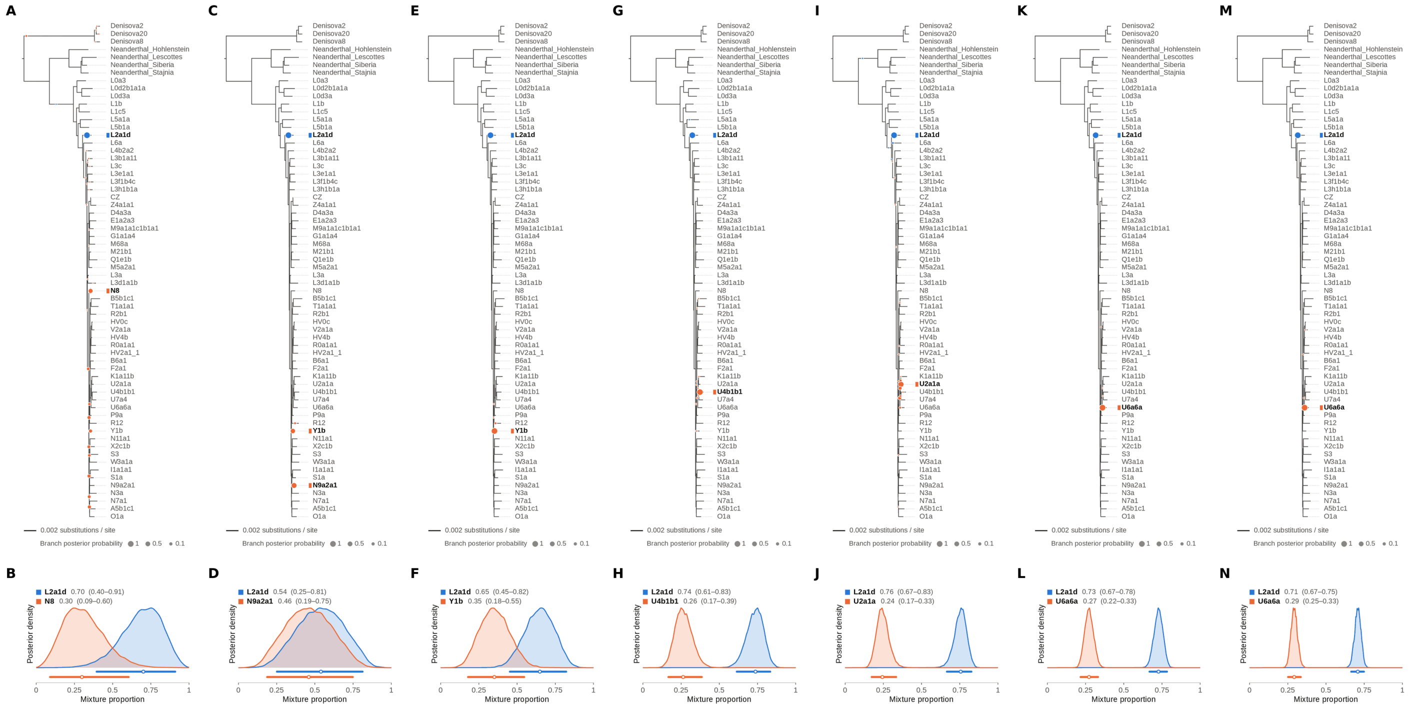


Figure S90: **Two-source simulated mixture of modern human L2 (AF381981.1) (75% of reads) and modern human U5a2a (HM229344.1; 25%) at nominal mean coverage depths of 0.5X, 1X, 2X, 4X, 8X, 16X, 32X.** Both sources carry ancient DNA damage (U5a2a with the damage rates scaled by 0.75). For each coverage depth, the upper panel shows the phylogenetic placement posterior of both sources for the best of five chains ($k = 2$, 100,000 iterations each), colored by source (dot area: branch posterior probability; swelling: posterior density of the position along the branch; bold tips: highest-posterior branches), and the lower panel the posterior densities of the mixture proportions $\Theta$ with median and 95% credible interval. L2 is expected on the L2a1d branch, the only L2 haplotype in the reference; U5a2a, for which no U5 haplotype is present in the reference, is expected within the U clade. Modal branch of each source in the best chain ($p$: its posterior probability) and $\Theta$ median [95% CI]: (A,B) 0.5X: L2a1d ($p = 1.00$), $\Theta = 0.70$ [0.40, 0.91]; N8 ($p = 0.29$), $\Theta = 0.30$ [0.09, 0.60]; (C,D) 1X: L2a1d ($p = 1.00$), $\Theta = 0.54$ [0.25, 0.81]; N9a2a1 ($p = 0.60$), $\Theta = 0.46$ [0.19, 0.75]; (E,F) 2X: L2a1d ($p = 1.00$), $\Theta = 0.65$ [0.45, 0.82]; Y1b ($p = 0.99$), $\Theta = 0.35$ [0.18, 0.55]; (G,H) 4X: L2a1d ($p = 1.00$), $\Theta = 0.74$ [0.61, 0.83]; U4b1b1 ($p = 0.99$), $\Theta = 0.26$ [0.17, 0.39]; (I,J) 8X: L2a1d ($p = 0.99$), $\Theta = 0.76$ [0.67, 0.83]; U2a1a ($p = 0.73$), $\Theta = 0.24$ [0.17, 0.33]; (K,L) 16X: L2a1d ($p = 1.00$), $\Theta = 0.73$ [0.67, 0.78]; U6a6a ($p = 1.00$), $\Theta = 0.27$ [0.22, 0.33]; (M,N) 32X: L2a1d ($p = 1.00$), $\Theta = 0.71$ [0.67, 0.75]; U6a6a ($p = 1.00$), $\Theta = 0.29$ [0.25, 0.33].

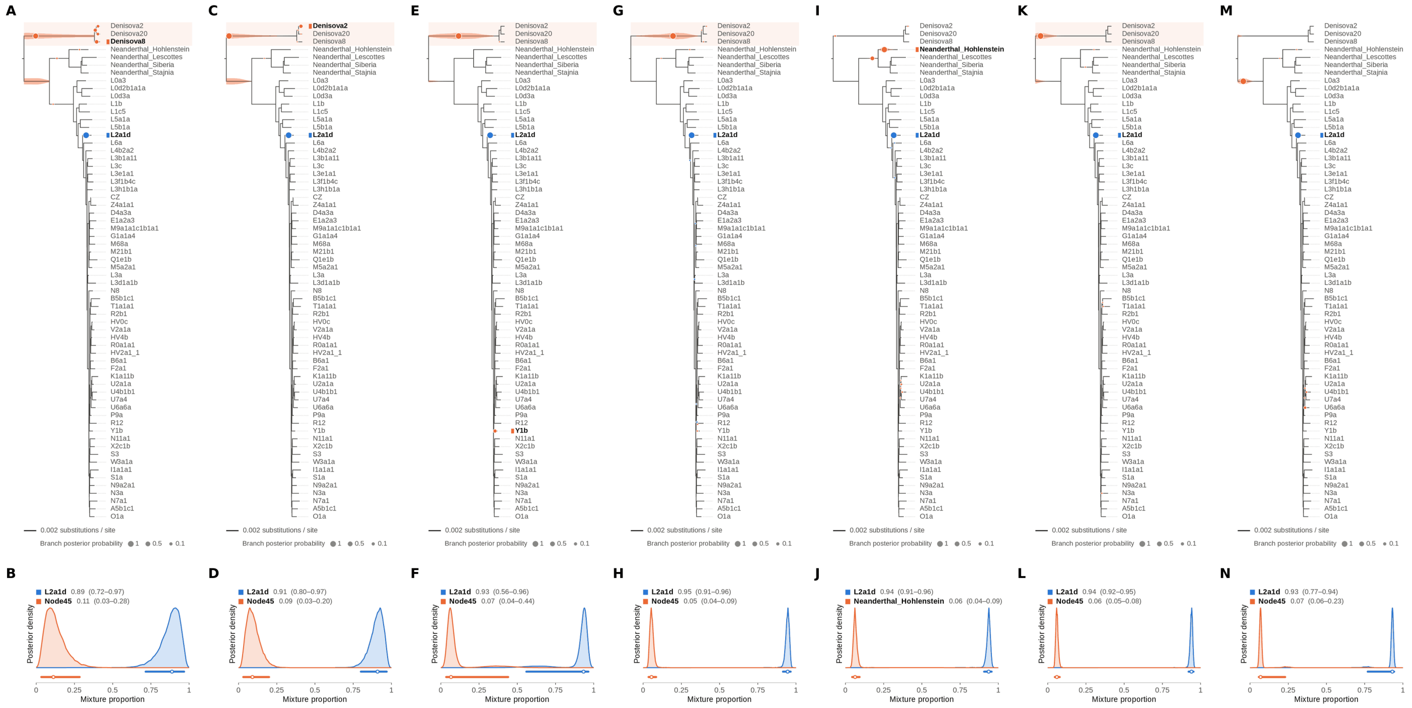


Figure S91: **Two-source simulated mixture of modern human L2 (AF381981.1) (80% of reads) and modern human U5a2a (HM229344.1; 20%) at nominal mean coverage depths of 0.5X, 1X, 2X, 4X, 8X, 16X, 32X.** Both sources carry ancient DNA damage (U5a2a with the damage rates scaled by 0.75). For each coverage depth, the upper panel shows the phylogenetic placement posterior of both sources for the best of five chains ($k = 2$, 100,000 iterations each), colored by source (dot area: branch posterior probability; swelling: posterior density of the position along the branch; bold tips: highest-posterior branches), and the lower panel the posterior densities of the mixture proportions $\Theta$ with median and 95% credible interval. L2 is expected on the L2a1d branch, the only L2 haplotype in the reference; U5a2a, for which no U5 haplotype is present in the reference, is expected within the U clade. Modal branch of each source in the best chain ($p$: its posterior probability) and $\Theta$ median [95% CI]: (A,B) 0.5X: L2a1d ($p = 1.00$), $\Theta = 0.89$ [0.72, 0.97]; Node45 (stem of the Denisovan clade; $p = 0.62$), $\Theta = 0.11$ [0.03, 0.28]; (C,D) 1X: L2a1d ($p = 1.00$), $\Theta = 0.91$ [0.80, 0.97]; Node45 (stem of the Denisovan clade; $p = 0.88$), $\Theta = 0.09$ [0.03, 0.20]; (E,F) 2X: L2a1d ($p = 1.00$), $\Theta = 0.93$ [0.56, 0.96]; Node45 (stem of the Denisovan clade; $p = 0.90$), $\Theta = 0.07$ [0.04, 0.44]; (G,H) 4X: L2a1d ($p = 0.99$), $\Theta = 0.95$ [0.91, 0.96]; Node45 (stem of the Denisovan clade; $p = 1.00$), $\Theta = 0.05$ [0.04, 0.09]; (I,J) 8X: L2a1d ($p = 1.00$), $\Theta = 0.94$ [0.91, 0.96]; Neanderthal_Hohlenstein ($p = 0.79$), $\Theta = 0.06$ [0.04, 0.09]; (K,L) 16X: L2a1d ($p = 1.00$), $\Theta = 0.94$ [0.92, 0.95]; Node45 (stem of the Denisovan clade; $p = 1.00$), $\Theta = 0.06$ [0.05, 0.08]; (M,N) 32X: L2a1d ($p = 1.00$), $\Theta = 0.93$ [0.77, 0.94]; Node45 (stem of the Denisovan clade; $p = 0.96$), $\Theta = 0.07$ [0.06, 0.23].

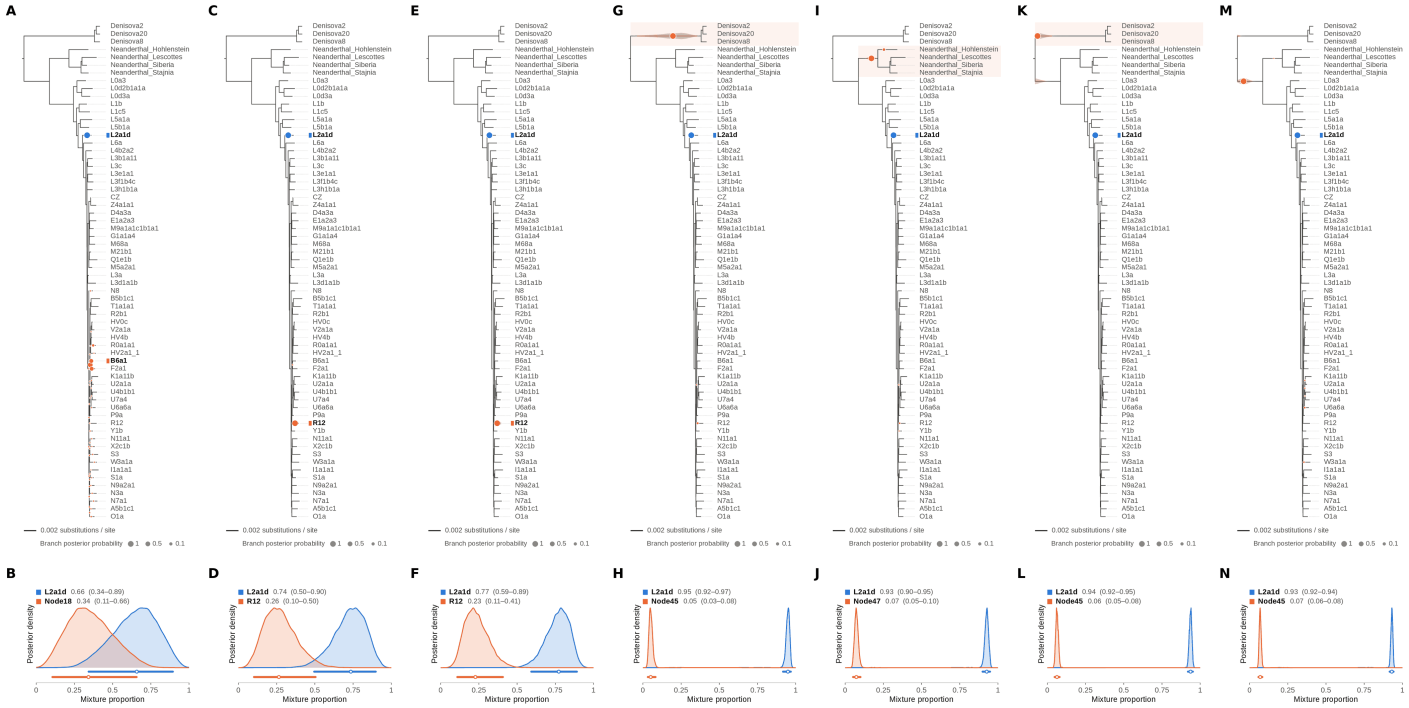


Figure S92: **Two-source simulated mixture of modern human L2 (AF381981.1) (85% of reads) and modern human U5a2a (HM229344.1; 15%) at nominal mean coverage depths of 0.5X, 1X, 2X, 4X, 8X, 16X, 32X.** Both sources carry ancient DNA damage (U5a2a with the damage rates scaled by 0.75). For each coverage depth, the upper panel shows the phylogenetic placement posterior of both sources for the best of five chains ($k = 2$, 100,000 iterations each), colored by source (dot area: branch posterior probability; swelling: posterior density of the position along the branch; bold tips: highest-posterior branches), and the lower panel the posterior densities of the mixture proportions $\Theta$ with median and 95% credible interval. L2 is expected on the L2a1d branch, the only L2 haplotype in the reference; U5a2a, for which no U5 haplotype is present in the reference, is expected within the U clade. Modal branch of each source in the best chain ($p$: its posterior probability) and $\Theta$ median [95% CI]: (A,B) 0.5X: L2a1d ($p = 1.00$), $\Theta = 0.66$ [0.34, 0.89]; Node18 (internal branch ancestral to B6a1 and F2a1; $p = 0.49$), $\Theta = 0.34$ [0.11, 0.66]; (C,D) 1X: L2a1d ($p = 1.00$), $\Theta = 0.74$ [0.50, 0.90]; R12 ($p = 1.00$), $\Theta = 0.26$ [0.10, 0.50]; (E,F) 2X: L2a1d ($p = 1.00$), $\Theta = 0.77$ [0.59, 0.89]; R12 ($p = 1.00$), $\Theta = 0.23$ [0.11, 0.41]; (G,H) 4X: L2a1d ($p = 1.00$), $\Theta = 0.95$ [0.92, 0.97]; Node45 (stem of the Denisovan clade; $p = 0.99$), $\Theta = 0.05$ [0.03, 0.08]; (I,J) 8X: L2a1d ($p = 1.00$), $\Theta = 0.93$ [0.90, 0.95]; Node47 (stem of the Neanderthal clade; $p = 0.96$), $\Theta = 0.07$ [0.05, 0.10]; (K,L) 16X: L2a1d ($p = 1.00$), $\Theta = 0.94$ [0.92, 0.95]; Node45 (stem of the Denisovan clade; $p = 1.00$), $\Theta = 0.06$ [0.05, 0.08]; (M,N) 32X: L2a1d ($p = 1.00$), $\Theta = 0.93$ [0.92, 0.94]; Node45 (stem of the Denisovan clade; $p = 0.99$), $\Theta = 0.07$ [0.06, 0.08].

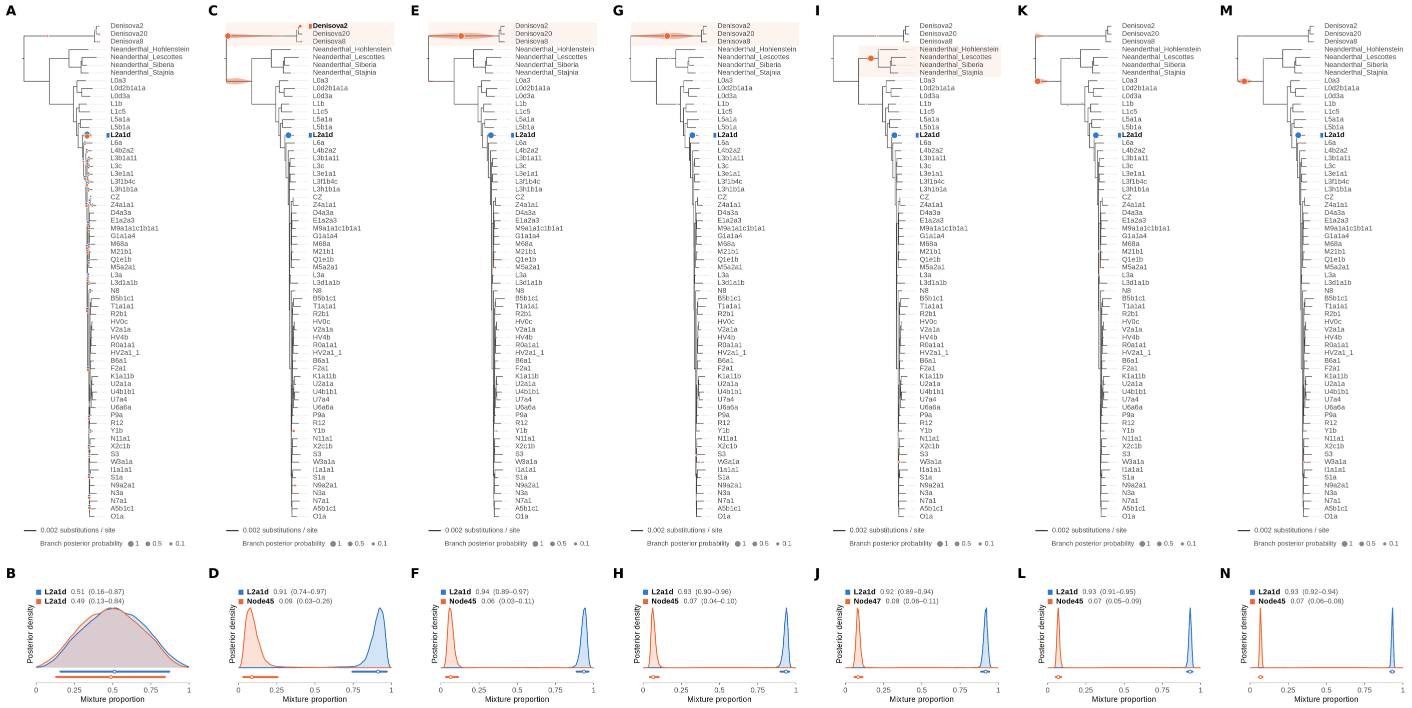


Figure S93: **Two-source simulated mixture of modern human L2 (AF381981.1) (90% of reads) and modern human U5a2a (HM229344.1; 10%) at nominal mean coverage depths of 0.5X, 1X, 2X, 4X, 8X, 16X, 32X.** Both sources carry ancient DNA damage (U5a2a with the damage rates scaled by 0.75). For each coverage depth, the upper panel shows the phylogenetic placement posterior of both sources for the best of five chains ($k = 2$, 100,000 iterations each), colored by source (dot area: branch posterior probability; swelling: posterior density of the position along the branch; bold tips: highest-posterior branches), and the lower panel the posterior densities of the mixture proportions $\Theta$ with median and 95% credible interval. L2 is expected on the L2a1d branch, the only L2 haplotype in the reference; U5a2a, for which no U5 haplotype is present in the reference, is expected within the U clade. Modal branch of each source in the best chain ($p$: its posterior probability) and $\Theta$ median [95% CI]: (A,B) 0.5X: L2a1d ($p = 0.60$), $\Theta = 0.51$ [0.16, 0.87]; L2a1d ($p = 0.39$), $\Theta = 0.49$ [0.13, 0.84]; (C,D) 1X: L2a1d ($p = 1.00$), $\Theta = 0.91$ [0.74, 0.97]; Node45 (stem of the Denisovan clade; $p = 0.90$), $\Theta = 0.09$ [0.03, 0.26]; (E,F) 2X: L2a1d ($p = 1.00$), $\Theta = 0.94$ [0.89, 0.97]; Node45 (stem of the Denisovan clade; $p = 0.99$), $\Theta = 0.06$ [0.03, 0.11]; (G,H) 4X: L2a1d ($p = 1.00$), $\Theta = 0.93$ [0.90, 0.96]; Node45 (stem of the Denisovan clade; $p = 1.00$), $\Theta = 0.07$ [0.04, 0.10]; (I,J) 8X: L2a1d ($p = 1.00$), $\Theta = 0.92$ [0.89, 0.94]; Node47 (stem of the Neanderthal clade; $p = 0.99$), $\Theta = 0.08$ [0.06, 0.11]; (K,L) 16X: L2a1d ($p = 1.00$), $\Theta = 0.93$ [0.91, 0.95]; Node45 (stem of the Denisovan clade; $p = 0.99$), $\Theta = 0.07$ [0.05, 0.09]; (M,N) 32X: L2a1d ($p = 1.00$), $\Theta = 0.93$ [0.92, 0.94]; Node45 (stem of the Denisovan clade; $p = 1.00$), $\Theta = 0.07$ [0.06, 0.08].

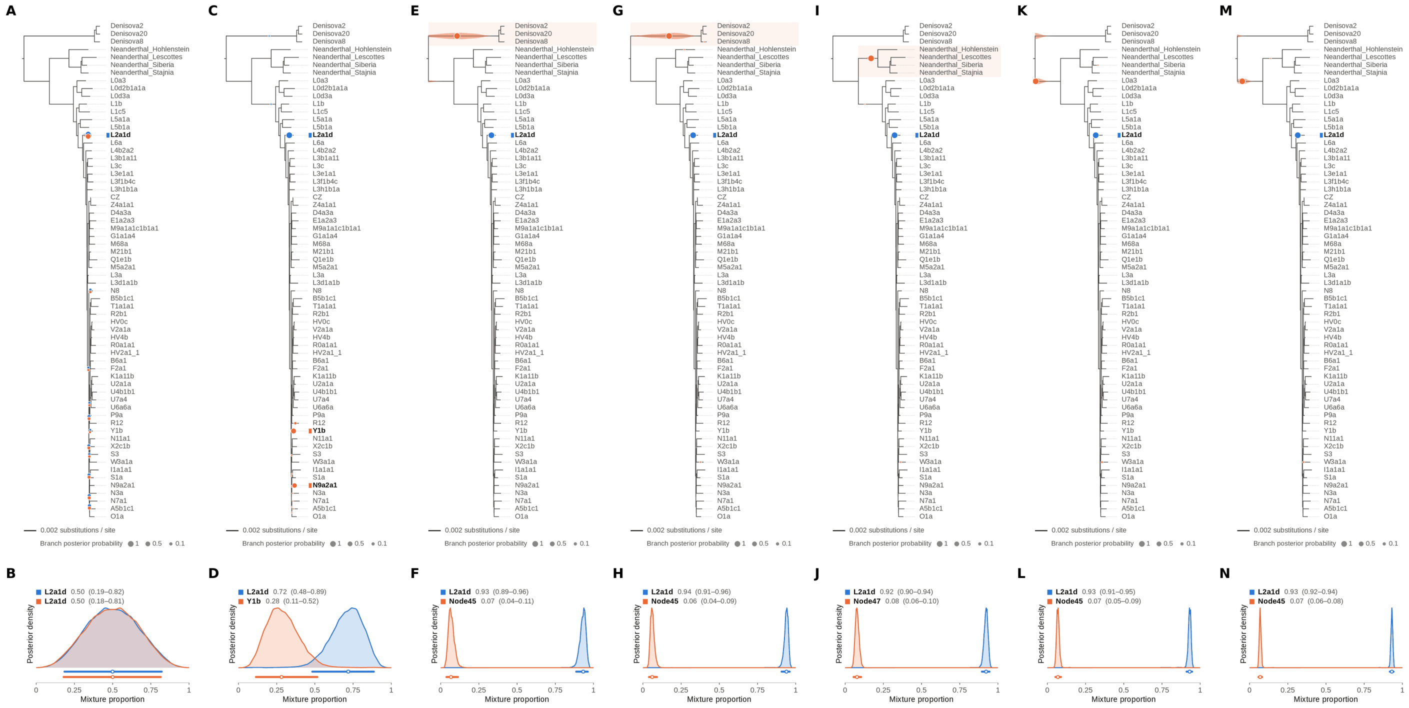


Figure S94: **Two-source simulated mixture of modern human L2 (AF381981.1) (95% of reads) and modern human U5a2a (HM229344.1; 5%) at nominal mean coverage depths of 0.5X, 1X, 2X, 4X, 8X, 16X, 32X.** Both sources carry ancient DNA damage (U5a2a with the damage rates scaled by 0.75). For each coverage depth, the upper panel shows the phylogenetic placement posterior of both sources for the best of five chains ($k = 2$, 100,000 iterations each), colored by source (dot area: branch posterior probability; swelling: posterior density of the position along the branch; bold tips: highest-posterior branches), and the lower panel the posterior densities of the mixture proportions $\Theta$ with median and 95% credible interval. L2 is expected on the L2a1d branch, the only L2 haplotype in the reference; U5a2a, for which no U5 haplotype is present in the reference, is expected within the U clade. Modal branch of each source in the best chain ($p$: its posterior probability) and $\Theta$ median [95% CI]: (A,B) 0.5X: L2a1d ($p = 0.50$), $\Theta = 0.50$ [0.18, 0.81]; L2a1d ($p = 0.50$), $\Theta = 0.50$ [0.19, 0.82]; (C,D) 1X: L2a1d ($p = 1.00$), $\Theta = 0.72$ [0.48, 0.89]; Y1b ($p = 0.57$), $\Theta = 0.28$ [0.11, 0.52]; (E,F) 2X: L2a1d ($p = 1.00$), $\Theta = 0.93$ [0.89, 0.96]; Node45 (stem of the Denisovan clade; $p = 1.00$), $\Theta = 0.07$ [0.04, 0.11]; (G,H) 4X: L2a1d ($p = 1.00$), $\Theta = 0.94$ [0.91, 0.96]; Node45 (stem of the Denisovan clade; $p = 1.00$), $\Theta = 0.06$ [0.04, 0.09]; (I,J) 8X: L2a1d ($p = 1.00$), $\Theta = 0.92$ [0.90, 0.94]; Node47 (stem of the Neanderthal clade; $p = 1.00$), $\Theta = 0.08$ [0.06, 0.10]; (K,L) 16X: L2a1d ($p = 1.00$), $\Theta = 0.93$ [0.91, 0.95]; Node45 (stem of the Denisovan clade; $p = 0.99$), $\Theta = 0.07$ [0.05, 0.09]; (M,N) 32X: L2a1d ($p = 1.00$), $\Theta = 0.93$ [0.92, 0.94]; Node45 (stem of the Denisovan clade; $p = 1.00$), $\Theta = 0.07$ [0.06, 0.08].

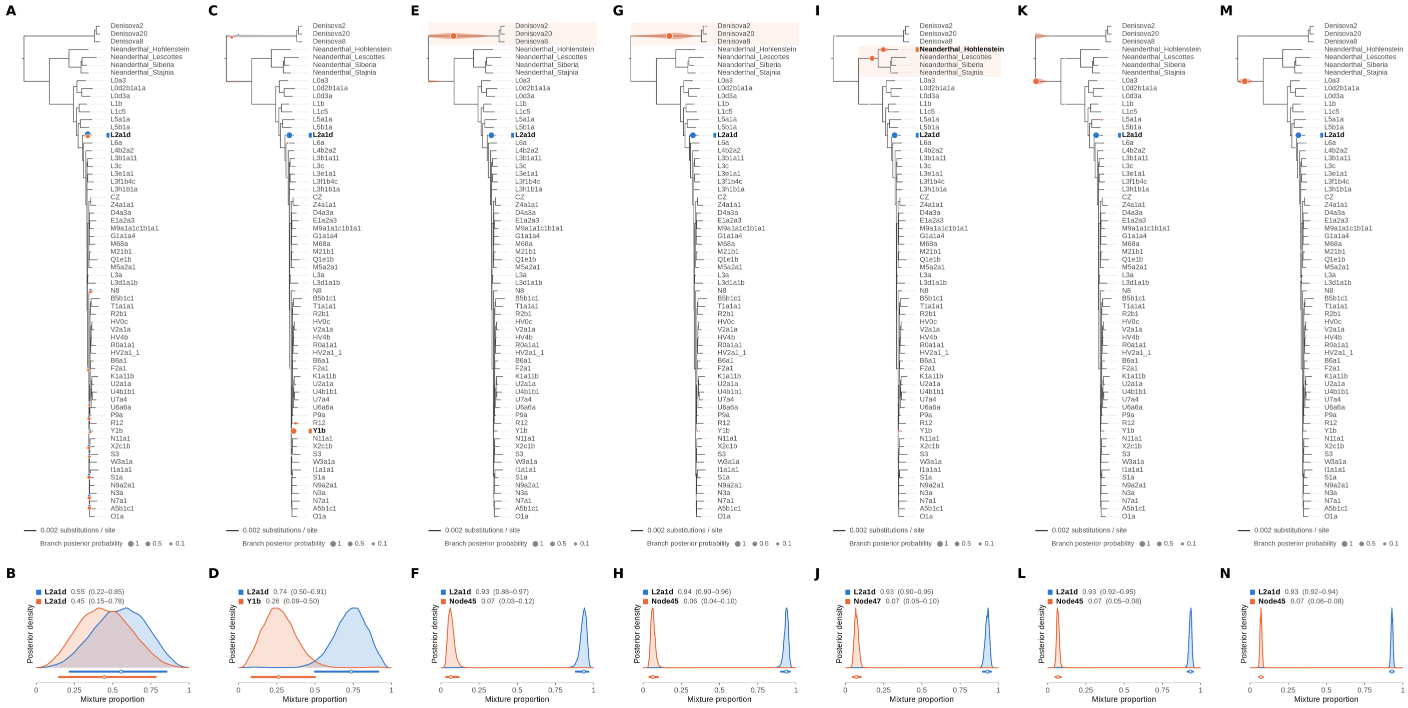


Figure S95: **Two-source simulated mixture of modern human L2 (AF381981.1) (99% of reads) and modern human U5a2a (HM229344.1; 1%) at nominal mean coverage depths of 0.5X, 1X, 2X, 4X, 8X, 16X, 32X.** Both sources carry ancient DNA damage (U5a2a with the damage rates scaled by 0.75). For each coverage depth, the upper panel shows the phylogenetic placement posterior of both sources for the best of five chains ($k = 2$, 100,000 iterations each), colored by source (dot area: branch posterior probability; swelling: posterior density of the position along the branch; bold tips: highest-posterior branches), and the lower panel the posterior densities of the mixture proportions $\Theta$ with median and 95% credible interval. L2 is expected on the L2a1d branch, the only L2 haplotype in the reference; U5a2a, for which no U5 haplotype is present in the reference, is expected within the U clade. Modal branch of each source in the best chain ($p$: its posterior probability) and $\Theta$ median [95% CI]: (A,B) 0.5X: L2a1d ($p = 0.81$), $\Theta = 0.55$ [0.22, 0.85]; L2a1d ($p = 0.19$), $\Theta = 0.45$ [0.15, 0.78]; (C,D) 1X: L2a1d ($p = 1.00$), $\Theta = 0.74$ [0.50, 0.91]; Y1b ($p = 0.89$), $\Theta = 0.26$ [0.09, 0.50]; (E,F) 2X: L2a1d ($p = 1.00$), $\Theta = 0.93$ [0.88, 0.97]; Node45 (stem of the Denisovan clade; $p = 1.00$), $\Theta = 0.07$ [0.03, 0.12]; (G,H) 4X: L2a1d ($p = 1.00$), $\Theta = 0.94$ [0.90, 0.96]; Node45 (stem of the Denisovan clade; $p = 1.00$), $\Theta = 0.06$ [0.04, 0.10]; (I,J) 8X: L2a1d ($p = 1.00$), $\Theta = 0.93$ [0.90, 0.95]; Node47 (stem of the Neanderthal clade; $p = 0.50$), $\Theta = 0.07$ [0.05, 0.10]; (K,L) 16X: L2a1d ($p = 1.00$), $\Theta = 0.93$ [0.92, 0.95]; Node45 (stem of the Denisovan clade; $p = 0.99$), $\Theta = 0.07$ [0.05, 0.08]; (M,N) 32X: L2a1d ($p = 1.00$), $\Theta = 0.93$ [0.92, 0.94]; Node45 (stem of the Denisovan clade; $p = 1.00$), $\Theta = 0.07$ [0.06, 0.08].

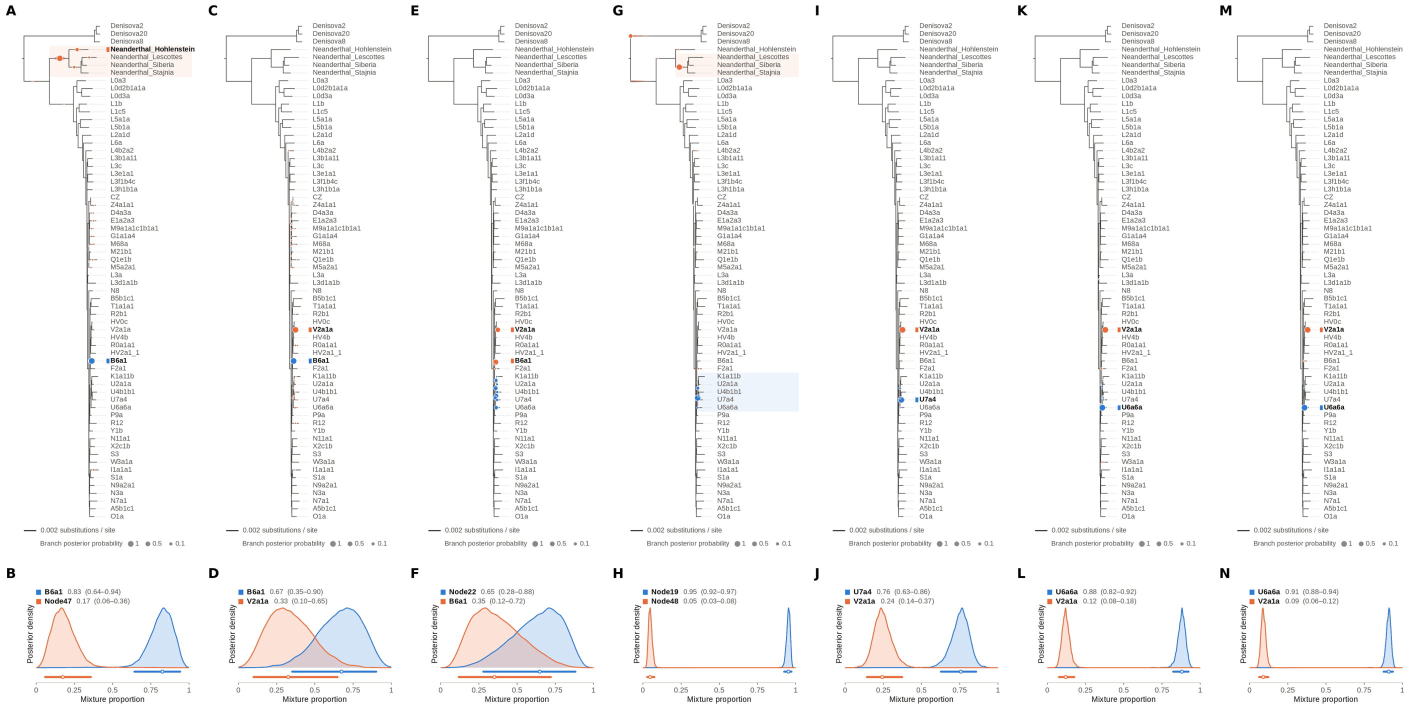


Figure S96: **Two-source simulated mixture of modern human T2a1 (KY671094.1) (1% of reads) and modern human U5a2a (HM229344.1; 99%) at nominal mean coverage depths of 0.5X, 1X, 2X, 4X, 8X, 16X, 32X.** Both sources carry ancient DNA damage (U5a2a with the damage rates scaled by 0.75). For each coverage depth, the upper panel shows the phylogenetic placement posterior of both sources for the best of five chains ($k = 2$, 100,000 iterations each), colored by source (dot area: branch posterior probability; swelling: posterior density of the position along the branch; bold tips: highest-posterior branches), and the lower panel the posterior densities of the mixture proportions $\Theta$ with median and 95% credible interval. T2a1 is expected on the T1a1a1 branch, the only haplogroup T haplotype in the reference; U5a2a, for which no U5 haplotype is present in the reference, is expected within the U clade. Modal branch of each source in the best chain ($p$: its posterior probability) and $\Theta$ median [95% CI]: (A,B) 0.5X: B6a1 ($p = 1.00$), $\Theta = 0.83$ [0.64, 0.94]; Node47 (stem of the Neanderthal clade; $p = 0.83$), $\Theta = 0.17$ [0.06, 0.36]; (C,D) 1X: B6a1 ($p = 1.00$), $\Theta = 0.67$ [0.35, 0.90]; V2a1a ($p = 0.97$), $\Theta = 0.33$ [0.10, 0.65]; (E,F) 2X: Node22 (internal branch ancestral to U4b1b1 and U7a4; $p = 0.29$), $\Theta = 0.65$ [0.28, 0.88]; B6a1 ($p = 0.59$), $\Theta = 0.35$ [0.12, 0.72]; (G,H) 4X: Node19 (stem of the U clade; $p = 0.91$), $\Theta = 0.95$ [0.92, 0.97]; Node48 (internal branch ancestral to Neanderthal_Lescottes, Neanderthal_Siberia and Neanderthal_Stajnia; $p = 0.82$), $\Theta = 0.05$ [0.03, 0.08]; (I,J) 8X: U7a4 ($p = 0.98$), $\Theta = 0.76$ [0.63, 0.86]; V2a1a ($p = 1.00$), $\Theta = 0.24$ [0.14, 0.37]; (K,L) 16X: U6a6a ($p = 1.00$), $\Theta = 0.88$ [0.82, 0.92]; V2a1a ($p = 1.00$), $\Theta = 0.12$ [0.08, 0.18]; (M,N) 32X: U6a6a ($p = 1.00$), $\Theta = 0.91$ [0.88, 0.94]; V2a1a ($p = 1.00$), $\Theta = 0.09$ [0.06, 0.12].

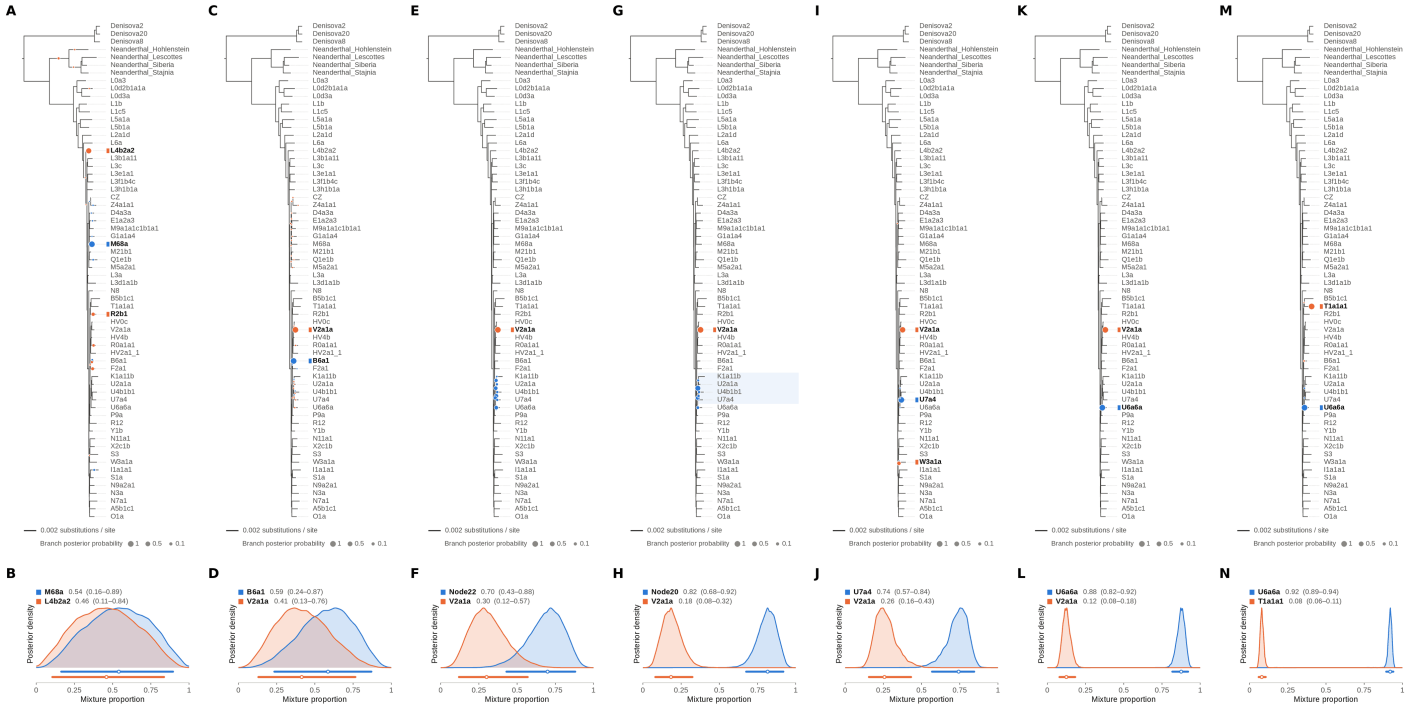

Figure S97: **Two-source simulated mixture of modern human T2a1 (KY671094.1) (5% of reads) and modern human U5a2a (HM229344.1; 95%) at nominal mean coverage depths of 0.5X, 1X, 2X, 4X, 8X, 16X, 32X.** Both sources carry ancient DNA damage (U5a2a with the damage rates scaled by 0.75). For each coverage depth, the upper panel shows the phylogenetic placement posterior of both sources for the best of five chains ($k = 2$, 100,000 iterations each), colored by source (dot area: branch posterior probability; swelling: posterior density of the position along the branch; bold tips: highest-posterior branches), and the lower panel the posterior densities of the mixture proportions $\Theta$ with median and 95% credible interval. T2a1 is expected on the T1a1a1 branch, the only haplogroup T haplotype in the reference; U5a2a, for which no U5 haplotype is present in the reference, is expected within the U clade. Modal branch of each source in the best chain ($p$: its posterior probability) and $\Theta$ median [95% CI]: (A,B) 0.5X: M68a ($p = 0.97$), $\Theta = 0.54$ [0.16, 0.89]; L4b2a2 ($p = 0.70$), $\Theta = 0.46$ [0.11, 0.84]; (C,D) 1X: B6a1 ($p = 1.00$), $\Theta = 0.59$ [0.24, 0.87]; V2a1a ($p = 0.99$), $\Theta = 0.41$ [0.13, 0.76]; (E,F) 2X: Node22 (internal branch ancestral to U4b1b1 and U7a4; $p = 0.27$), $\Theta = 0.70$ [0.43, 0.88]; V2a1a ($p = 1.00$), $\Theta = 0.30$ [0.12, 0.57]; (G,H) 4X: Node20 (internal branch within the U clade; $p = 0.62$), $\Theta = 0.82$ [0.68, 0.92]; V2a1a ($p = 1.00$), $\Theta = 0.18$ [0.08, 0.32]; (I,J) 8X: U7a4 ($p = 1.00$), $\Theta = 0.74$ [0.57, 0.84]; V2a1a ($p = 0.84$), $\Theta = 0.26$ [0.16, 0.43]; (K,L) 16X: U6a6a ($p = 1.00$), $\Theta = 0.88$ [0.82, 0.92]; V2a1a ($p = 1.00$), $\Theta = 0.12$ [0.08, 0.18]; (M,N) 32X: U6a6a ($p = 1.00$), $\Theta = 0.92$ [0.89, 0.94]; T1a1a1 ($p = 1.00$), $\Theta = 0.08$ [0.06, 0.11].

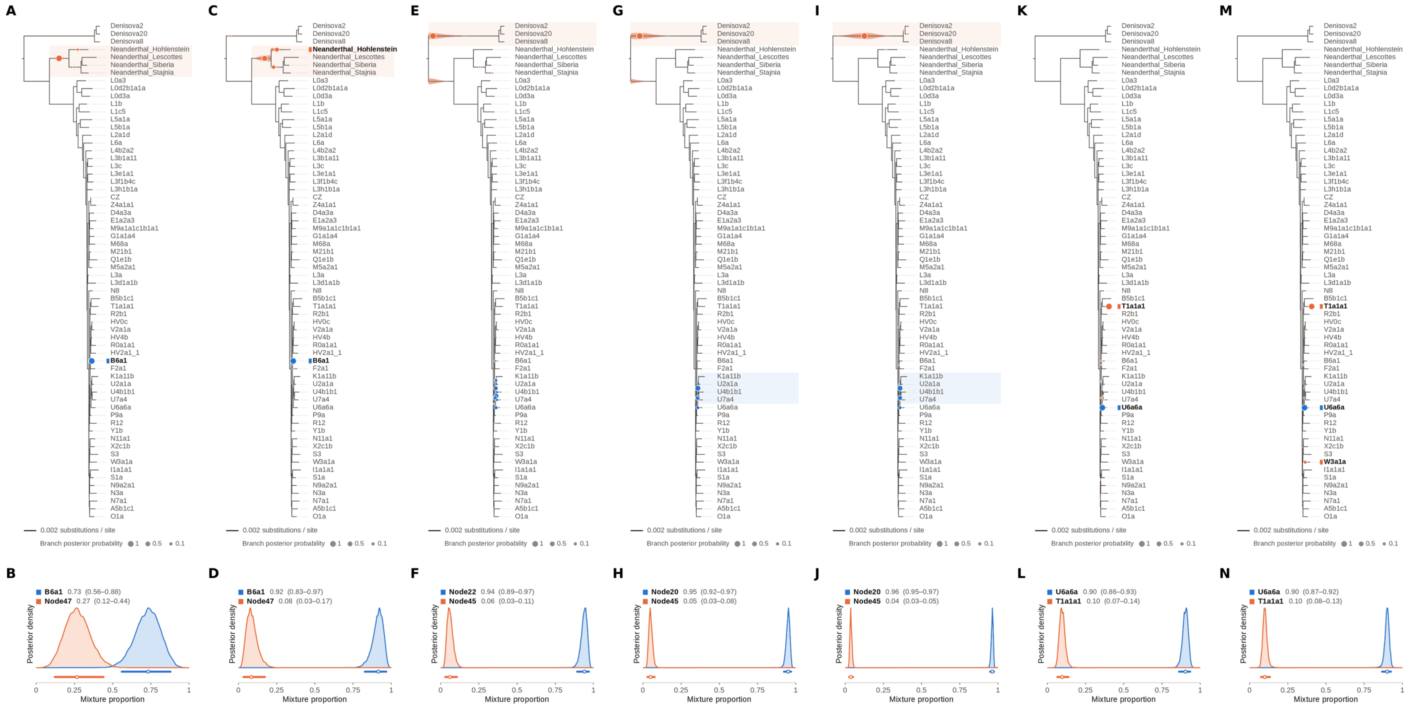

Figure S98: **Two-source simulated mixture of modern human T2a1 (KY671094.1) (10% of reads) and modern human U5a2a (HM229344.1; 90%) at nominal mean coverage depths of 0.5X, 1X, 2X, 4X, 8X, 16X, 32X.** Both sources carry ancient DNA damage (U5a2a with the damage rates scaled by 0.75). For each coverage depth, the upper panel shows the phylogenetic placement posterior of both sources for the best of five chains ($k = 2$, 100,000 iterations each), colored by source (dot area: branch posterior probability; swelling: posterior density of the position along the branch; bold tips: highest-posterior branches), and the lower panel the posterior densities of the mixture proportions Θ with median and 95% credible interval. T2a1 is expected on the T1a1a1 branch, the only haplogroup T haplotype in the reference; U5a2a, for which no U5 haplotype is present in the reference, is expected within the U clade. Modal branch of each source in the best chain ($p$: its posterior probability) and Θ median [95% CI]: (A,B) 0.5X: B6a1 ($p = 1.00$), $\Theta = 0.73$ [0.56, 0.88]; Node47 (stem of the Neanderthal clade; $p = 0.97$), $\Theta = 0.27$ [0.12, 0.44]; (C,D) 1X: B6a1 ($p = 1.00$), $\Theta = 0.92$ [0.83, 0.97]; Node47 (stem of the Neanderthal clade; $p = 0.59$), $\Theta = 0.08$ [0.03, 0.17]; (E,F) 2X: Node22 (internal branch ancestral to U4b1b1 and U7a4; $p = 0.36$), $\Theta = 0.94$ [0.89, 0.97]; Node45 (stem of the Denisovan clade; $p = 1.00$), $\Theta = 0.06$ [0.03, 0.11]; (G,H) 4X: Node20 (internal branch within the U clade; $p = 0.65$), $\Theta = 0.95$ [0.92, 0.97]; Node45 (stem of the Denisovan clade; $p = 1.00$), $\Theta = 0.05$ [0.03, 0.08]; (I,J) 8X: Node20 (internal branch within the U clade; $p = 0.57$), $\Theta = 0.96$ [0.95, 0.97]; Node45 (stem of the Denisovan clade; $p = 1.00$), $\Theta = 0.04$ [0.03, 0.05]; (K,L) 16X: U6a6a ($p = 1.00$), $\Theta = 0.90$ [0.86, 0.93]; T1a1a1 ($p = 1.00$), $\Theta = 0.10$ [0.07, 0.14]; (M,N) 32X: U6a6a ($p = 1.00$), $\Theta = 0.90$ [0.87, 0.92]; T1a1a1 ($p = 0.93$), $\Theta = 0.10$ [0.08, 0.13].

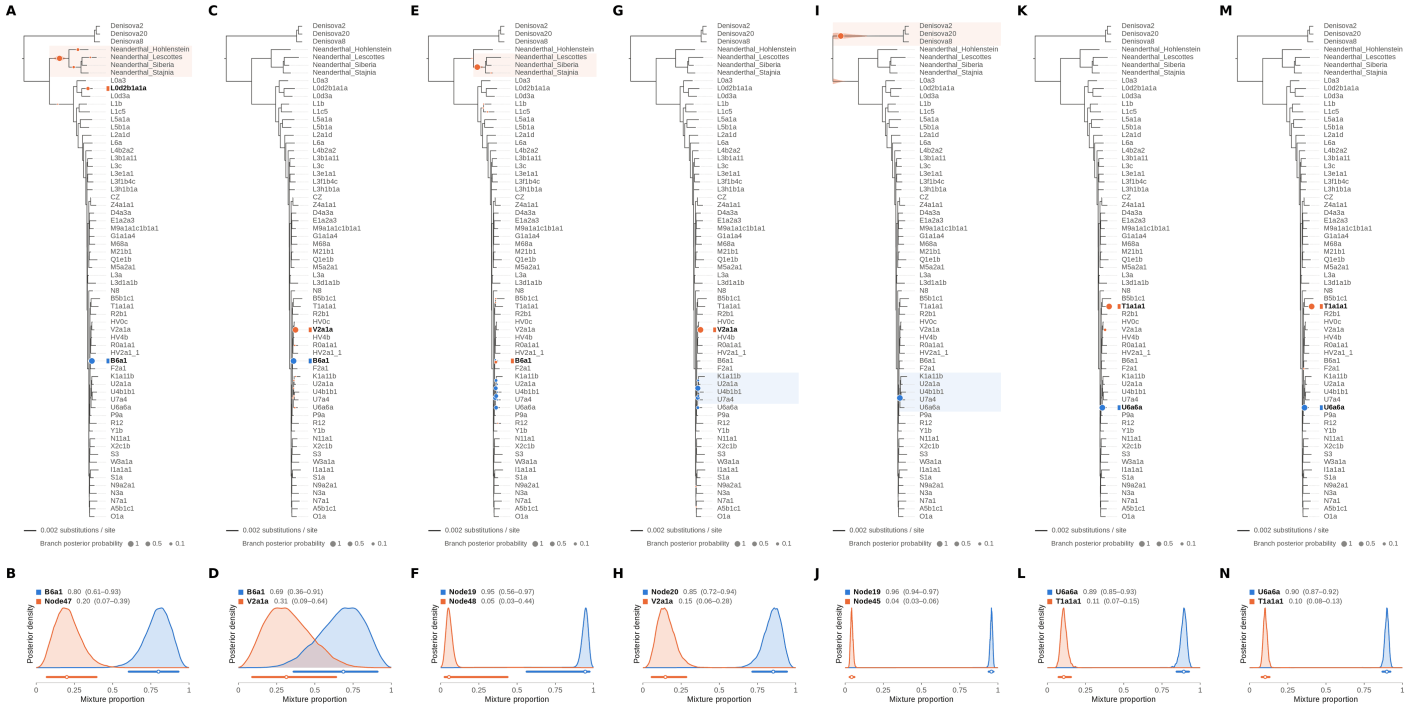


Figure S99: **Two-source simulated mixture of modern human T2a1 (KY671094.1) (15% of reads) and modern human U5a2a (HM229344.1; 85%) at nominal mean coverage depths of 0.5X, 1X, 2X, 4X, 8X, 16X, 32X.** Both sources carry ancient DNA damage (U5a2a with the damage rates scaled by 0.75). For each coverage depth, the upper panel shows the phylogenetic placement posterior of both sources for the best of five chains ($k = 2$, 100,000 iterations each), colored by source (dot area: branch posterior probability; swelling: posterior density of the position along the branch; bold tips: highest-posterior branches), and the lower panel the posterior densities of the mixture proportions $\Theta$ with median and 95% credible interval. T2a1 is expected on the T1a1a1 branch, the only haplogroup T haplotype in the reference; U5a2a, for which no U5 haplotype is present in the reference, is expected within the U clade. Modal branch of each source in the best chain ($p$: its posterior probability) and $\Theta$ median [95% CI]: (A,B) 0.5X: B6a1 ($p = 1.00$), $\Theta = 0.80$ [0.61, 0.93]; Node47 (stem of the Neanderthal clade; $p = 0.78$), $\Theta = 0.20$ [0.07, 0.39]; (C,D) 1X: B6a1 ($p = 1.00$), $\Theta = 0.69$ [0.36, 0.91]; V2a1a ($p = 0.99$), $\Theta = 0.31$ [0.09, 0.64]; (E,F) 2X: Node19 (stem of the U clade; $p = 0.28$), $\Theta = 0.95$ [0.56, 0.97]; Node48 (internal branch ancestral to Neanderthal_Lescottes, Neanderthal_Siberia and Neanderthal_Stajnia; $p = 0.94$), $\Theta = 0.05$ [0.03, 0.44]; (G,H) 4X: Node20 (internal branch within the U clade; $p = 0.66$), $\Theta = 0.85$ [0.72, 0.94]; V2a1a ($p = 1.00$), $\Theta = 0.15$ [0.06, 0.28]; (I,J) 8X: Node19 (stem of the U clade; $p = 1.00$), $\Theta = 0.96$ [0.94, 0.97]; Node45 (stem of the Denisovan clade; $p = 1.00$), $\Theta = 0.04$ [0.03, 0.06]; (K,L) 16X: U6a6a ($p = 1.00$), $\Theta = 0.89$ [0.85, 0.93]; T1a1a1 ($p = 0.96$), $\Theta = 0.11$ [0.07, 0.15]; (M,N) 32X: U6a6a ($p = 1.00$), $\Theta = 0.90$ [0.87, 0.92]; T1a1a1 ($p = 1.00$), $\Theta = 0.10$ [0.08, 0.13].

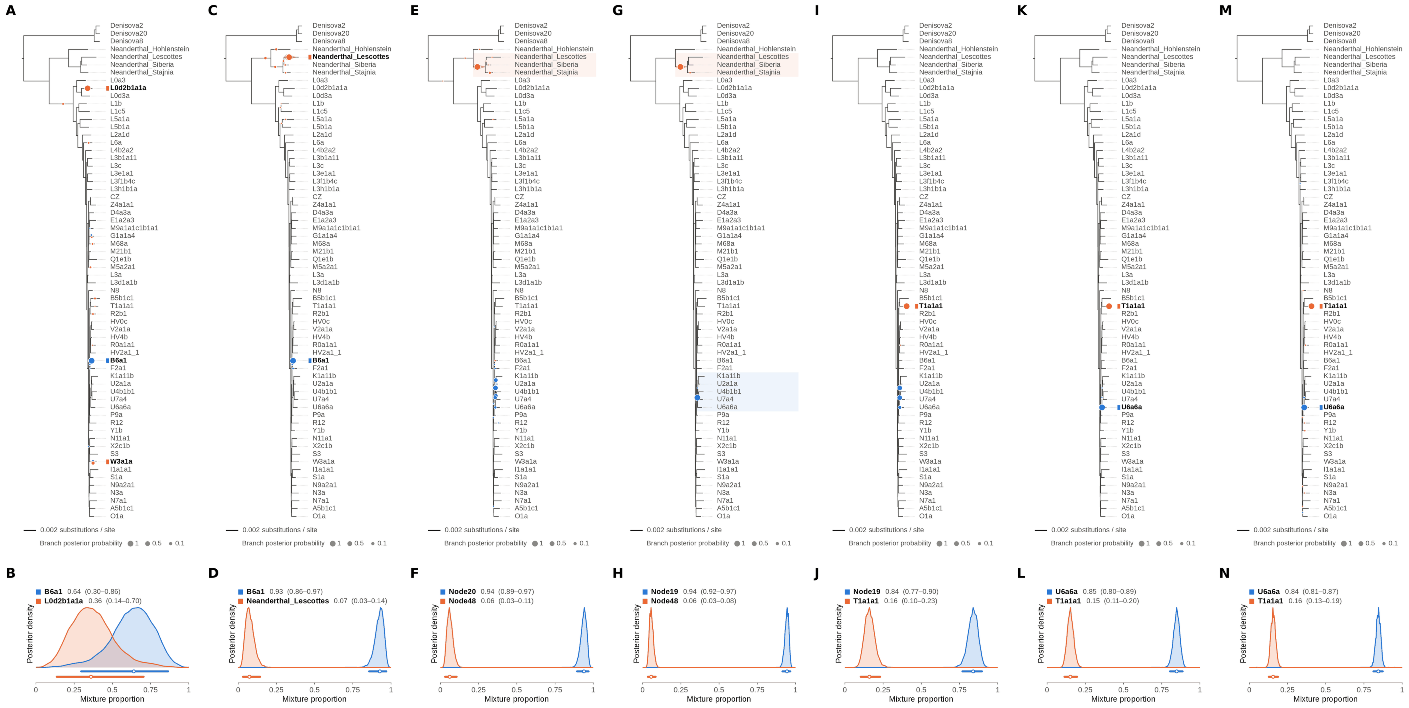

Figure S100: **Two-source simulated mixture of modern human T2a1 (KY671094.1) (20% of reads) and modern human U5a2a (HM229344.1; 80%) at nominal mean coverage depths of 0.5X, 1X, 2X, 4X, 8X, 16X, 32X.** Both sources carry ancient DNA damage (U5a2a with the damage rates scaled by 0.75). For each coverage depth, the upper panel shows the phylogenetic placement posterior of both sources for the best of five chains ($k = 2$, 100,000 iterations each), colored by source (dot area: branch posterior probability; swelling: posterior density of the position along the branch; bold tips: highest-posterior branches), and the lower panel the posterior densities of the mixture proportions $\Theta$ with median and 95% credible interval. T2a1 is expected on the T1a1a1 branch, the only haplogroup T haplotype in the reference; U5a2a, for which no U5 haplotype is present in the reference, is expected within the U clade. Modal branch of each source in the best chain ($p$: its posterior probability) and $\Theta$ median [95% CI]: (A,B) 0.5X: B6a1 ($p = 0.99$), $\Theta = 0.64$ [0.30, 0.86]; L0d2b1a1a ($p = 0.86$), $\Theta = 0.36$ [0.14, 0.70]; (C,D) 1X: B6a1 ($p = 0.99$), $\Theta = 0.93$ [0.86, 0.97]; Neanderthal_Lescottes ($p = 0.91$), $\Theta = 0.07$ [0.03, 0.14]; (E,F) 2X: Node20 (internal branch within the U clade; $p = 0.47$), $\Theta = 0.94$ [0.89, 0.97]; Node48 (internal branch ancestral to Neanderthal_Lescottes, Neanderthal_Siberia and Neanderthal_Stajnia; $p = 0.97$), $\Theta = 0.06$ [0.03, 0.11]; (G,H) 4X: Node19 (stem of the U clade; $p = 1.00$), $\Theta = 0.94$ [0.92, 0.97]; Node48 (internal branch ancestral to Neanderthal_Lescottes, Neanderthal_Siberia and Neanderthal_Stajnia; $p = 1.00$), $\Theta = 0.06$ [0.03, 0.08]; (I,J) 8X: Node19 (stem of the U clade; $p = 0.46$), $\Theta = 0.84$ [0.77, 0.90]; T1a1a1 ($p = 1.00$), $\Theta = 0.16$ [0.10, 0.23]; (K,L) 16X: U6a6a ($p = 1.00$), $\Theta = 0.85$ [0.80, 0.89]; T1a1a1 ($p = 1.00$), $\Theta = 0.15$ [0.11, 0.20]; (M,N) 32X: U6a6a ($p = 1.00$), $\Theta = 0.84$ [0.81, 0.87]; T1a1a1 ($p = 1.00$), $\Theta = 0.16$ [0.13, 0.19].

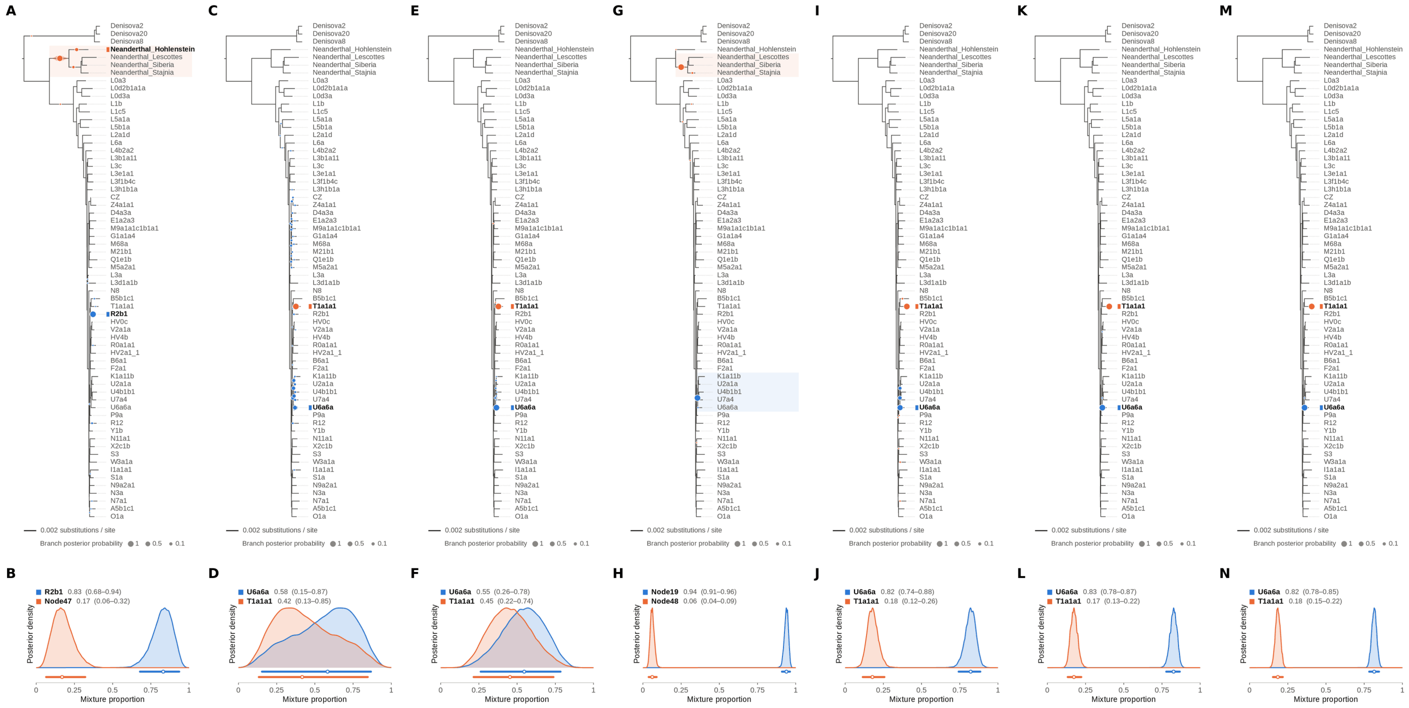


Figure S101: **Two-source simulated mixture of modern human T2a1 (KY671094.1) (25% of reads) and modern human U5a2a (HM229344.1; 75%) at nominal mean coverage depths of 0.5X, 1X, 2X, 4X, 8X, 16X, 32X.** Both sources carry ancient DNA damage (U5a2a with the damage rates scaled by 0.75). For each coverage depth, the upper panel shows the phylogenetic placement posterior of both sources for the best of five chains ($k = 2$, 100,000 iterations each), colored by source (dot area: branch posterior probability; swelling: posterior density of the position along the branch; bold tips: highest-posterior branches), and the lower panel the posterior densities of the mixture proportions $\Theta$ with median and 95% credible interval. T2a1 is expected on the T1a1a1 branch, the only haplogroup T haplotype in the reference; U5a2a, for which no U5 haplotype is present in the reference, is expected within the U clade. Modal branch of each source in the best chain ($p$: its posterior probability) and $\Theta$ median [95% CI]: (A,B) 0.5X: R2b1 ($p = 0.99$), $\Theta = 0.83$ [0.68, 0.94]; Node47 (stem of the Neanderthal clade; $p = 0.86$), $\Theta = 0.17$ [0.06, 0.32]; (C,D) 1X: U6a6a ($p = 0.24$), $\Theta = 0.58$ [0.15, 0.87]; T1a1a1 ($p = 1.00$), $\Theta = 0.42$ [0.13, 0.85]; (E,F) 2X: U6a6a ($p = 0.98$), $\Theta = 0.55$ [0.26, 0.78]; T1a1a1 ($p = 1.00$), $\Theta = 0.45$ [0.22, 0.74]; (G,H) 4X: Node19 (stem of the U clade; $p = 1.00$), $\Theta = 0.94$ [0.91, 0.96]; Node48 (internal branch ancestral to Neanderthal_Lescottes, Neanderthal_Siberia and Neanderthal_Stajnia; $p = 0.99$), $\Theta = 0.06$ [0.04, 0.09]; (I,J) 8X: U6a6a ($p = 0.74$), $\Theta = 0.82$ [0.74, 0.88]; T1a1a1 ($p = 0.99$), $\Theta = 0.18$ [0.12, 0.26]; (K,L) 16X: U6a6a ($p = 1.00$), $\Theta = 0.83$ [0.78, 0.87]; T1a1a1 ($p = 1.00$), $\Theta = 0.17$ [0.13, 0.22]; (M,N) 32X: U6a6a ($p = 1.00$), $\Theta = 0.82$ [0.78, 0.85]; T1a1a1 ($p = 1.00$), $\Theta = 0.18$ [0.15, 0.22].

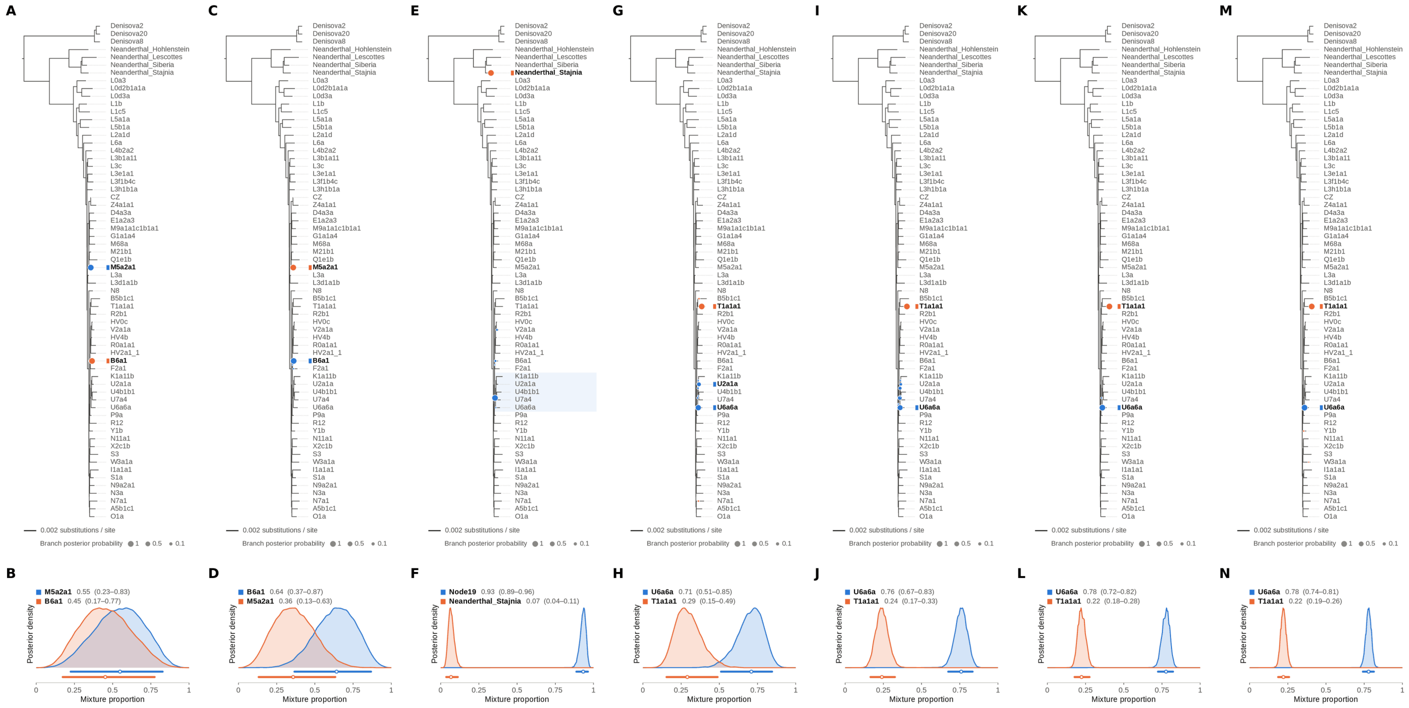

Figure S102: **Two-source simulated mixture of modern human T2a1 (KY671094.1) (30% of reads) and modern human U5a2a (HM229344.1; 70%) at nominal mean coverage depths of 0.5X, 1X, 2X, 4X, 8X, 16X, 32X.** Both sources carry ancient DNA damage (U5a2a with the damage rates scaled by 0.75). For each coverage depth, the upper panel shows the phylogenetic placement posterior of both sources for the best of five chains ($k = 2$, 100,000 iterations each), colored by source (dot area: branch posterior probability; swelling: posterior density of the position along the branch; bold tips: highest-posterior branches), and the lower panel the posterior densities of the mixture proportions $\Theta$ with median and 95% credible interval. T2a1 is expected on the T1a1a1 branch, the only haplogroup T haplotype in the reference; U5a2a, for which no U5 haplotype is present in the reference, is expected within the U clade. Modal branch of each source in the best chain ($p$: its posterior probability) and $\Theta$ median [95% CI]: (A,B) 0.5X: M5a2a1 ($p = 1.00$), $\Theta = 0.55$ [0.23, 0.83]; B6a1 ($p = 1.00$), $\Theta = 0.45$ [0.17, 0.77]; (C,D) 1X: B6a1 ($p = 1.00$), $\Theta = 0.64$ [0.37, 0.87]; M5a2a1 ($p = 1.00$), $\Theta = 0.36$ [0.13, 0.63]; (E,F) 2X: Node19 (stem of the U clade; $p = 0.97$), $\Theta = 0.93$ [0.89, 0.96]; Neanderthal_Stajnia ($p = 1.00$), $\Theta = 0.07$ [0.04, 0.11]; (G,H) 4X: U6a6a ($p = 0.77$), $\Theta = 0.71$ [0.51, 0.85]; T1a1a1 ($p = 1.00$), $\Theta = 0.29$ [0.15, 0.49]; (I,J) 8X: U6a6a ($p = 0.64$), $\Theta = 0.76$ [0.67, 0.83]; T1a1a1 ($p = 1.00$), $\Theta = 0.24$ [0.17, 0.33]; (K,L) 16X: U6a6a ($p = 1.00$), $\Theta = 0.78$ [0.72, 0.82]; T1a1a1 ($p = 1.00$), $\Theta = 0.22$ [0.18, 0.28]; (M,N) 32X: U6a6a ($p = 1.00$), $\Theta = 0.78$ [0.74, 0.81]; T1a1a1 ($p = 1.00$), $\Theta = 0.22$ [0.19, 0.26].

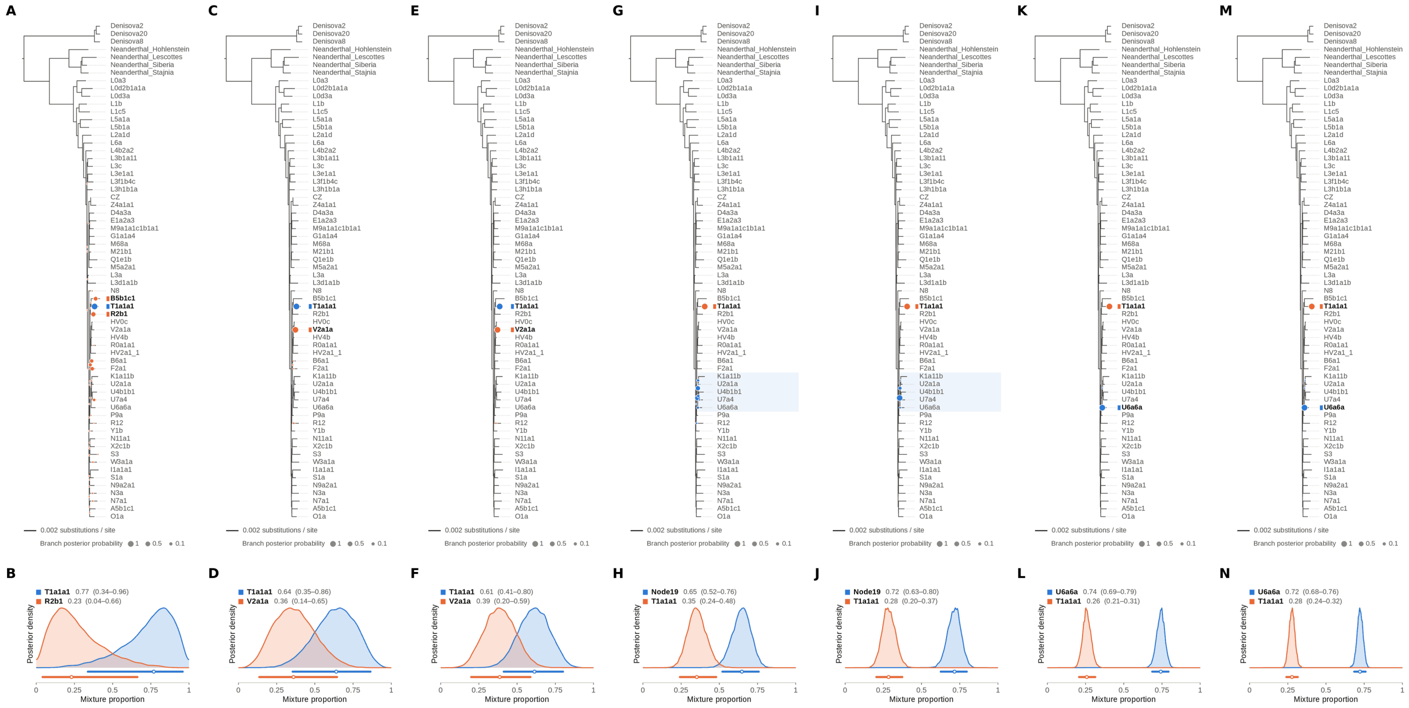


Figure S103: **Two-source simulated mixture of modern human T2a1 (KY671094.1) (35% of reads) and modern human U5a2a (HM229344.1; 65%) at nominal mean coverage depths of 0.5X, 1X, 2X, 4X, 8X, 16X, 32X.** Both sources carry ancient DNA damage (U5a2a with the damage rates scaled by 0.75). For each coverage depth, the upper panel shows the phylogenetic placement posterior of both sources for the best of five chains ($k = 2$, 100,000 iterations each), colored by source (dot area: branch posterior probability; swelling: posterior density of the position along the branch; bold tips: highest-posterior branches), and the lower panel the posterior densities of the mixture proportions $\Theta$ with median and 95% credible interval. T2a1 is expected on the T1a1a1 branch, the only haplogroup T haplotype in the reference; U5a2a, for which no U5 haplotype is present in the reference, is expected within the U clade. Modal branch of each source in the best chain ($p$: its posterior probability) and $\Theta$ median [95% CI]: (A,B) 0.5X: T1a1a1 ($p = 1.00$), $\Theta = 0.77$ [0.34, 0.96]; R2b1 ($p = 0.33$), $\Theta = 0.23$ [0.04, 0.66]; (C,D) 1X: T1a1a1 ($p = 1.00$), $\Theta = 0.64$ [0.35, 0.86]; V2a1a ($p = 0.99$), $\Theta = 0.36$ [0.14, 0.65]; (E,F) 2X: T1a1a1 ($p = 1.00$), $\Theta = 0.61$ [0.41, 0.80]; V2a1a ($p = 1.00$), $\Theta = 0.39$ [0.20, 0.59]; (G,H) 4X: Node19 (stem of the U clade; $p = 0.53$), $\Theta = 0.65$ [0.52, 0.76]; T1a1a1 ($p = 1.00$), $\Theta = 0.35$ [0.24, 0.48]; (I,J) 8X: Node19 (stem of the U clade; $p = 0.90$), $\Theta = 0.72$ [0.63, 0.80]; T1a1a1 ($p = 1.00$), $\Theta = 0.28$ [0.20, 0.37]; (K,L) 16X: U6a6a ($p = 1.00$), $\Theta = 0.74$ [0.69, 0.79]; T1a1a1 ($p = 1.00$), $\Theta = 0.26$ [0.21, 0.31]; (M,N) 32X: U6a6a ($p = 1.00$), $\Theta = 0.72$ [0.68, 0.76]; T1a1a1 ($p = 1.00$), $\Theta = 0.28$ [0.24, 0.32].

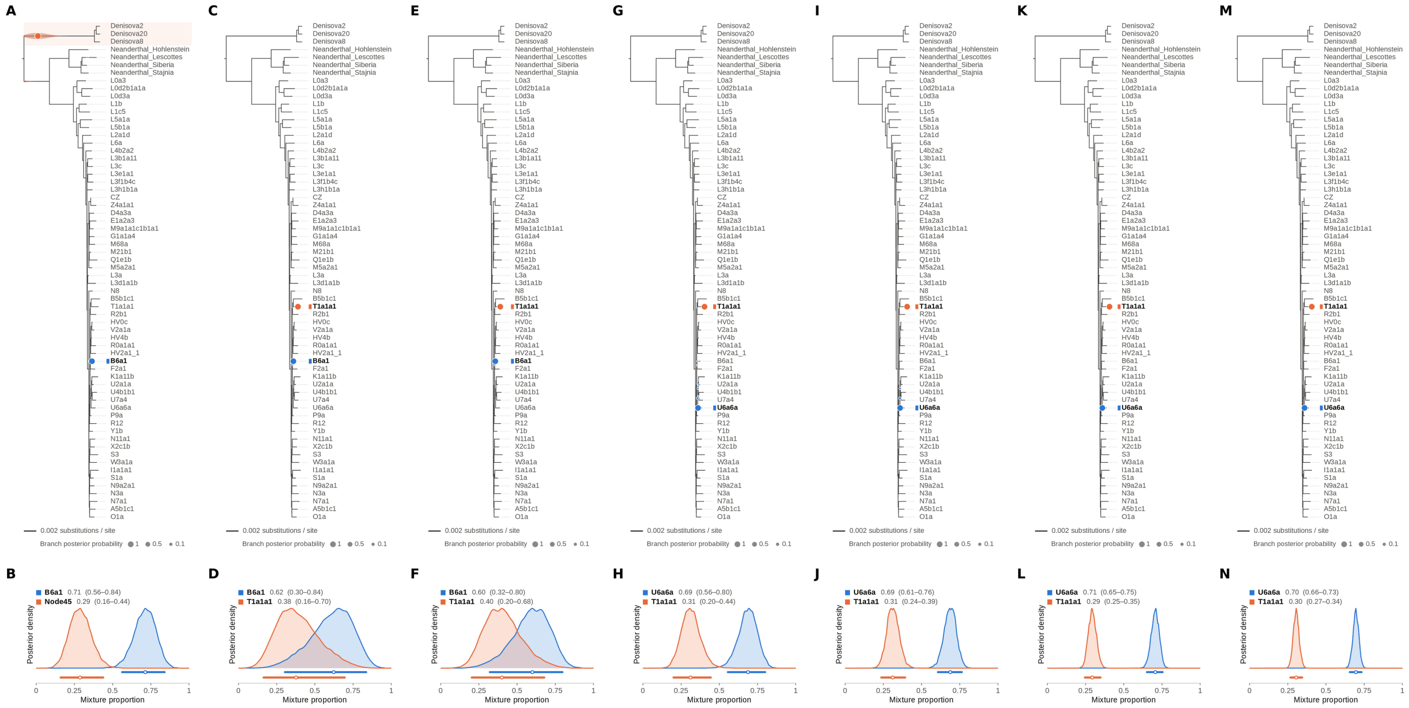

Figure S104: **Two-source simulated mixture of modern human T2a1 (KY671094.1) (40% of reads) and modern human U5a2a (HM229344.1; 60%) at nominal mean coverage depths of 0.5X, 1X, 2X, 4X, 8X, 16X, 32X.** Both sources carry ancient DNA damage (U5a2a with the damage rates scaled by 0.75). For each coverage depth, the upper panel shows the phylogenetic placement posterior of both sources for the best of five chains ($k = 2$, 100,000 iterations each), colored by source (dot area: branch posterior probability; swelling: posterior density of the position along the branch; bold tips: highest-posterior branches), and the lower panel the posterior densities of the mixture proportions $\Theta$ with median and 95% credible interval. T2a1 is expected on the T1a1a1 branch, the only haplogroup T haplotype in the reference; U5a2a, for which no U5 haplotype is present in the reference, is expected within the U clade. Modal branch of each source in the best chain ($p$: its posterior probability) and $\Theta$ median [95% CI]: (A,B) 0.5X: B6a1 ($p = 1.00$), $\Theta = 0.71$ [0.56, 0.84]; Node45 (stem of the Denisovan clade; $p = 1.00$), $\Theta = 0.29$ [0.16, 0.44]; (C,D) 1X: B6a1 ($p = 1.00$), $\Theta = 0.62$ [0.30, 0.84]; T1a1a1 ($p = 1.00$), $\Theta = 0.38$ [0.16, 0.70]; (E,F) 2X: B6a1 ($p = 1.00$), $\Theta = 0.60$ [0.32, 0.80]; T1a1a1 ($p = 1.00$), $\Theta = 0.40$ [0.20, 0.68]; (G,H) 4X: U6a6a ($p = 0.99$), $\Theta = 0.69$ [0.56, 0.80]; T1a1a1 ($p = 1.00$), $\Theta = 0.31$ [0.20, 0.44]; (I,J) 8X: U6a6a ($p = 0.98$), $\Theta = 0.69$ [0.61, 0.76]; T1a1a1 ($p = 1.00$), $\Theta = 0.31$ [0.24, 0.39]; (K,L) 16X: U6a6a ($p = 1.00$), $\Theta = 0.71$ [0.65, 0.75]; T1a1a1 ($p = 1.00$), $\Theta = 0.29$ [0.25, 0.35]; (M,N) 32X: U6a6a ($p = 1.00$), $\Theta = 0.70$ [0.66, 0.73]; T1a1a1 ($p = 1.00$), $\Theta = 0.30$ [0.27, 0.34].

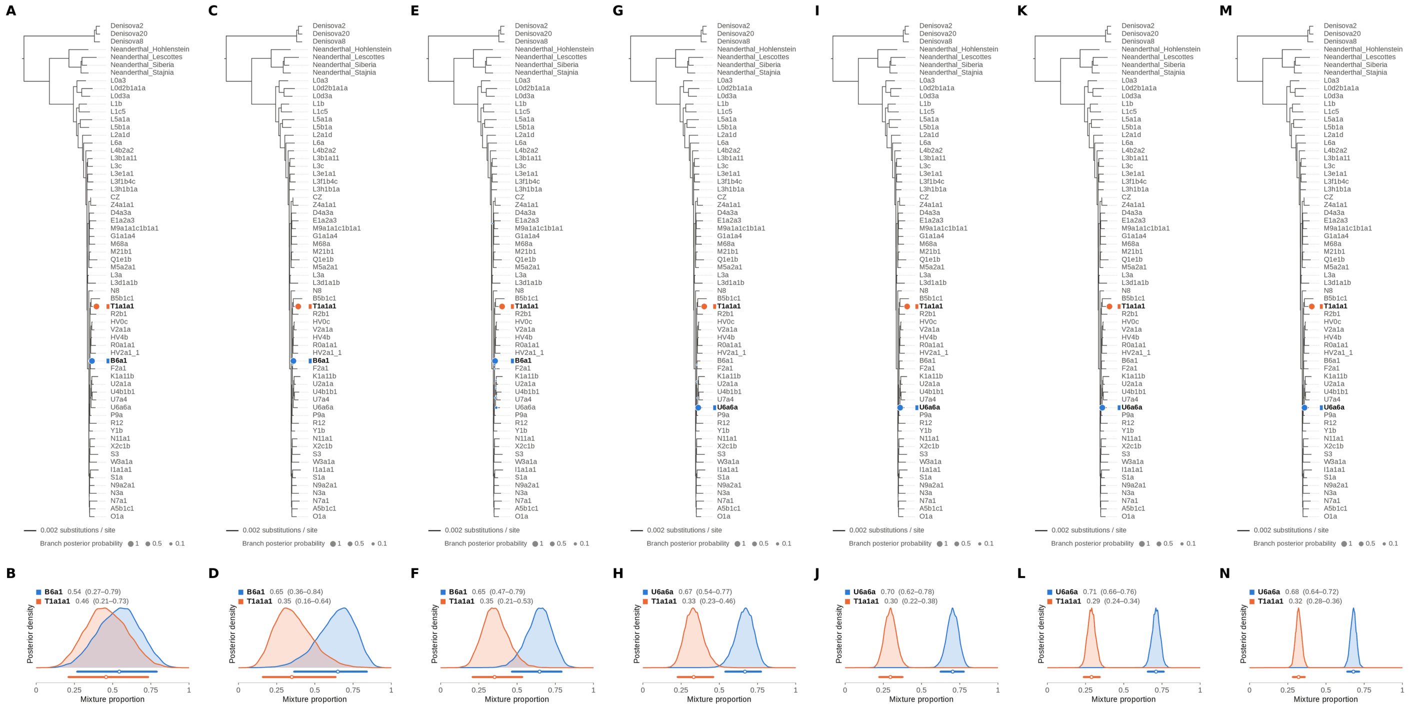

Figure S105: **Two-source simulated mixture of modern human T2a1 (KY671094.1) (45% of reads) and modern human U5a2a (HM229344.1; 55%) at nominal mean coverage depths of 0.5X, 1X, 2X, 4X, 8X, 16X, 32X.** Both sources carry ancient DNA damage (U5a2a with the damage rates scaled by 0.75). For each coverage depth, the upper panel shows the phylogenetic placement posterior of both sources for the best of five chains ($k = 2$, 100,000 iterations each), colored by source (dot area: branch posterior probability; swelling: posterior density of the position along the branch; bold tips: highest-posterior branches), and the lower panel the posterior densities of the mixture proportions $\Theta$ with median and 95% credible interval. T2a1 is expected on the T1a1a1 branch, the only haplogroup T haplotype in the reference; U5a2a, for which no U5 haplotype is present in the reference, is expected within the U clade. Modal branch of each source in the best chain ($p$: its posterior probability) and $\Theta$ median [95% CI]: (A,B) 0.5X: B6a1 ($p = 1.00$), $\Theta = 0.54$ [0.27, 0.79]; T1a1a1 ($p = 1.00$), $\Theta = 0.46$ [0.21, 0.73]; (C,D) 1X: B6a1 ($p = 1.00$), $\Theta = 0.65$ [0.36, 0.84]; T1a1a1 ($p = 1.00$), $\Theta = 0.35$ [0.16, 0.64]; (E,F) 2X: B6a1 ($p = 0.96$), $\Theta = 0.65$ [0.47, 0.79]; T1a1a1 ($p = 1.00$), $\Theta = 0.35$ [0.21, 0.53]; (G,H) 4X: U6a6a ($p = 1.00$), $\Theta = 0.67$ [0.54, 0.77]; T1a1a1 ($p = 1.00$), $\Theta = 0.33$ [0.23, 0.46]; (I,J) 8X: U6a6a ($p = 1.00$), $\Theta = 0.70$ [0.62, 0.78]; T1a1a1 ($p = 1.00$), $\Theta = 0.30$ [0.22, 0.38]; (K,L) 16X: U6a6a ($p = 1.00$), $\Theta = 0.71$ [0.66, 0.76]; T1a1a1 ($p = 1.00$), $\Theta = 0.29$ [0.24, 0.34]; (M,N) 32X: U6a6a ($p = 1.00$), $\Theta = 0.68$ [0.64, 0.72]; T1a1a1 ($p = 1.00$), $\Theta = 0.32$ [0.28, 0.36].

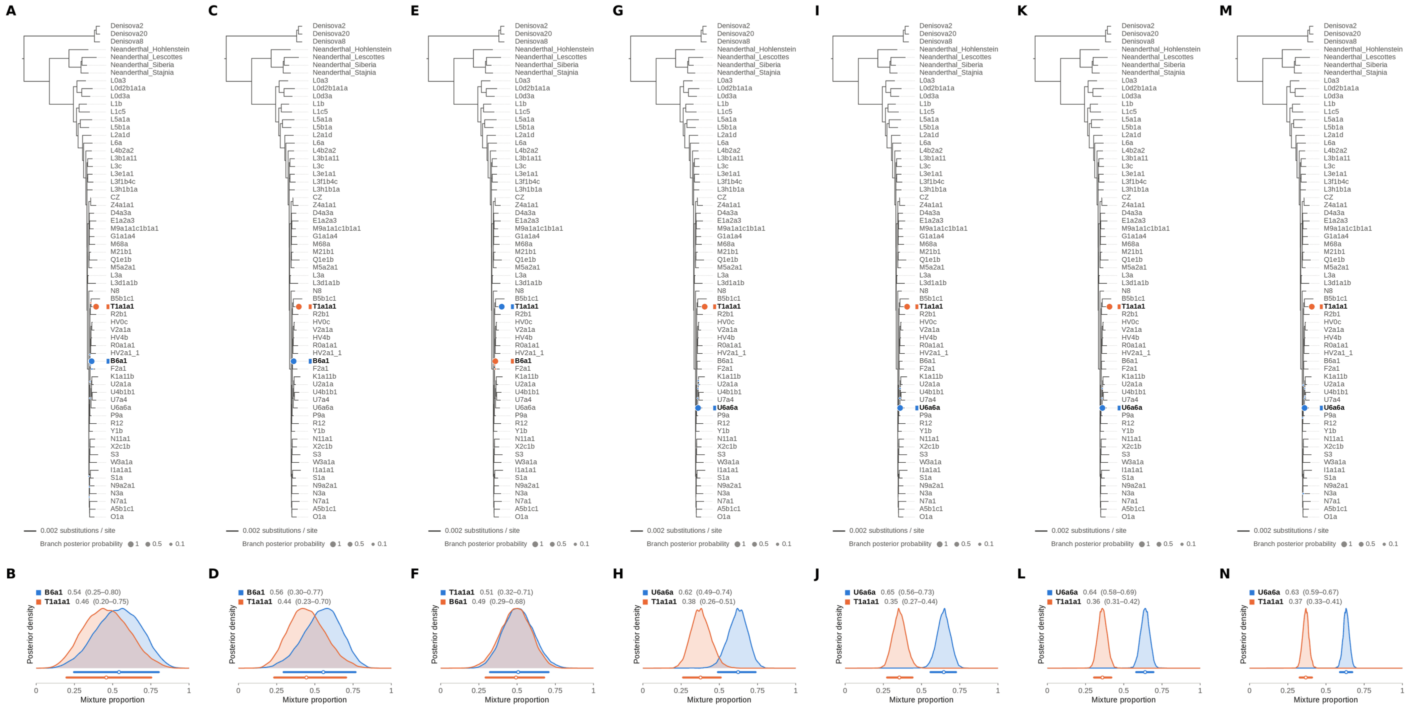


Figure S106: **Two-source simulated mixture of modern human T2a1 (KY671094.1) (50% of reads) and modern human U5a2a (HM229344.1; 50%) at nominal mean coverage depths of 0.5X, 1X, 2X, 4X, 8X, 16X, 32X.** Both sources carry ancient DNA damage (U5a2a with the damage rates scaled by 0.75). For each coverage depth, the upper panel shows the phylogenetic placement posterior of both sources for the best of five chains ($k = 2$, 100,000 iterations each), colored by source (dot area: branch posterior probability; swelling: posterior density of the position along the branch; bold tips: highest-posterior branches), and the lower panel the posterior densities of the mixture proportions $\Theta$ with median and 95% credible interval. T2a1 is expected on the T1a1a1 branch, the only haplogroup T haplotype in the reference; U5a2a, for which no U5 haplotype is present in the reference, is expected within the U clade. Modal branch of each source in the best chain ($p$: its posterior probability) and $\Theta$ median [95% CI]: (A,B) 0.5X: B6a1 ($p = 1.00$), $\Theta = 0.54$ [0.25, 0.80]; T1a1a1 ($p = 1.00$), $\Theta = 0.46$ [0.20, 0.75]; (C,D) 1X: B6a1 ($p = 1.00$), $\Theta = 0.56$ [0.30, 0.77]; T1a1a1 ($p = 1.00$), $\Theta = 0.44$ [0.23, 0.70]; (E,F) 2X: T1a1a1 ($p = 1.00$), $\Theta = 0.51$ [0.32, 0.71]; B6a1 ($p = 1.00$), $\Theta = 0.49$ [0.29, 0.68]; (G,H) 4X: U6a6a ($p = 1.00$), $\Theta = 0.62$ [0.49, 0.74]; T1a1a1 ($p = 1.00$), $\Theta = 0.38$ [0.26, 0.51]; (I,J) 8X: U6a6a ($p = 1.00$), $\Theta = 0.65$ [0.56, 0.73]; T1a1a1 ($p = 1.00$), $\Theta = 0.35$ [0.27, 0.44]; (K,L) 16X: U6a6a ($p = 1.00$), $\Theta = 0.64$ [0.58, 0.69]; T1a1a1 ($p = 1.00$), $\Theta = 0.36$ [0.31, 0.42]; (M,N) 32X: U6a6a ($p = 1.00$), $\Theta = 0.63$ [0.59, 0.67]; T1a1a1 ($p = 1.00$), $\Theta = 0.37$ [0.33, 0.41].

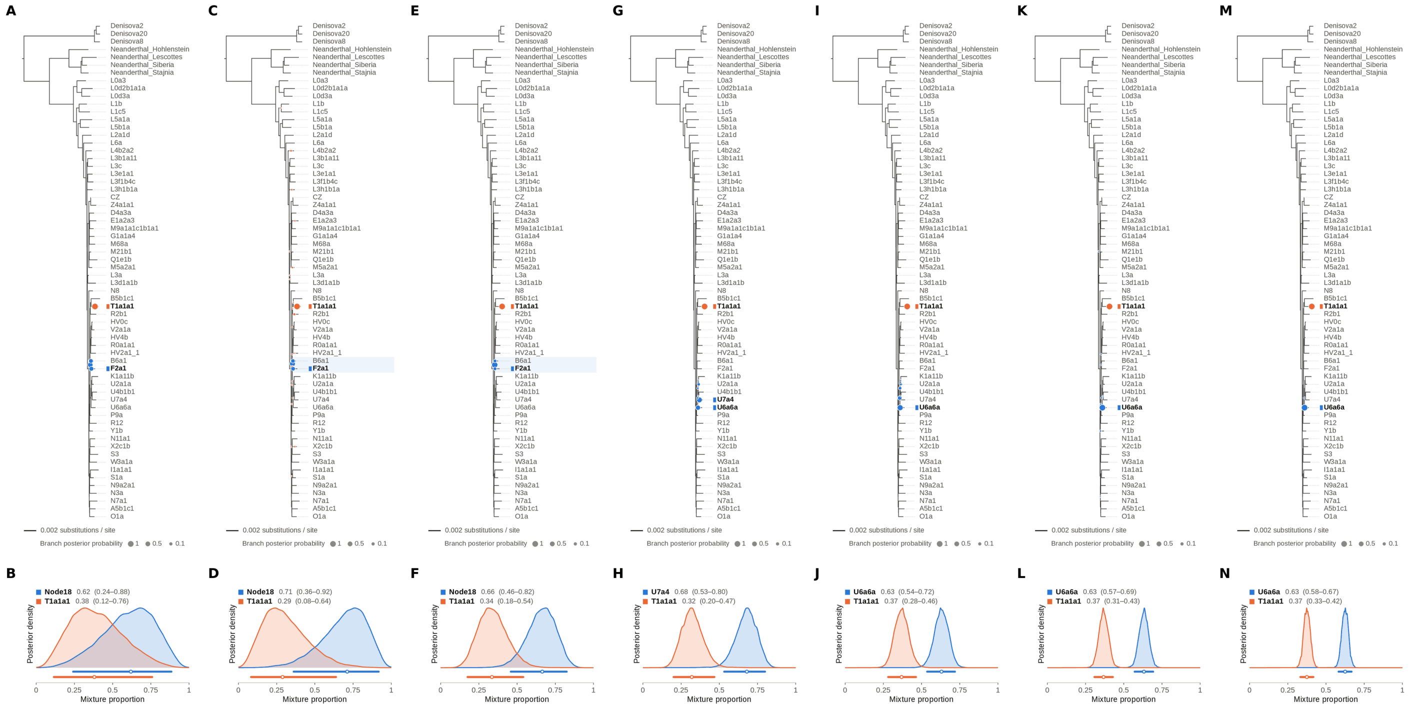

Figure S107: **Two-source simulated mixture of modern human T2a1 (KY671094.1) (55% of reads) and modern human U5a2a (HM229344.1; 45%) at nominal mean coverage depths of 0.5X, 1X, 2X, 4X, 8X, 16X, 32X.** Both sources carry ancient DNA damage (U5a2a with the damage rates scaled by 0.75). For each coverage depth, the upper panel shows the phylogenetic placement posterior of both sources for the best of five chains ($k = 2$, 100,000 iterations each), colored by source (dot area: branch posterior probability; swelling: posterior density of the position along the branch; bold tips: highest-posterior branches), and the lower panel the posterior densities of the mixture proportions $\Theta$ with median and 95% credible interval. T2a1 is expected on the T1a1a1 branch, the only haplogroup T haplotype in the reference; U5a2a, for which no U5 haplotype is present in the reference, is expected within the U clade. Modal branch of each source in the best chain ($p$: its posterior probability) and $\Theta$ median [95% CI]: (A,B) 0.5X: Node18 (internal branch ancestral to B6a1 and F2a1; $p = 0.49$), $\Theta = 0.62$ [0.24, 0.88]; T1a1a1 ($p = 1.00$), $\Theta = 0.38$ [0.12, 0.76]; (C,D) 1X: Node18 (internal branch ancestral to B6a1 and F2a1; $p = 0.63$), $\Theta = 0.71$ [0.36, 0.92]; T1a1a1 ($p = 0.97$), $\Theta = 0.29$ [0.08, 0.64]; (E,F) 2X: Node18 (internal branch ancestral to B6a1 and F2a1; $p = 0.85$), $\Theta = 0.66$ [0.46, 0.82]; T1a1a1 ($p = 1.00$), $\Theta = 0.34$ [0.18, 0.54]; (G,H) 4X: U7a4 ($p = 0.60$), $\Theta = 0.68$ [0.53, 0.80]; T1a1a1 ($p = 1.00$), $\Theta = 0.32$ [0.20, 0.47]; (I,J) 8X: U6a6a ($p = 0.73$), $\Theta = 0.63$ [0.54, 0.72]; T1a1a1 ($p = 1.00$), $\Theta = 0.37$ [0.28, 0.46]; (K,L) 16X: U6a6a ($p = 0.99$), $\Theta = 0.63$ [0.57, 0.69]; T1a1a1 ($p = 1.00$), $\Theta = 0.37$ [0.31, 0.43]; (M,N) 32X: U6a6a ($p = 1.00$), $\Theta = 0.63$ [0.58, 0.67]; T1a1a1 ($p = 1.00$), $\Theta = 0.37$ [0.33, 0.42].

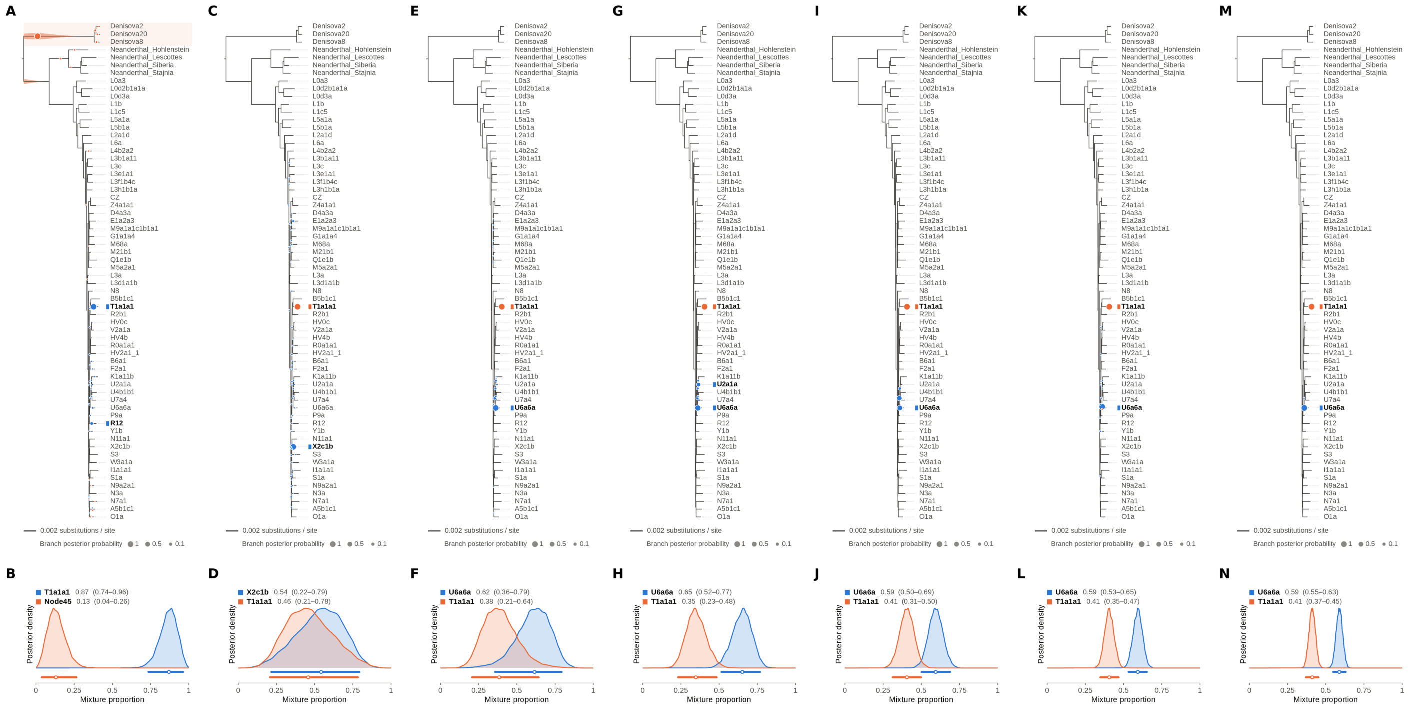

Figure S108: **Two-source simulated mixture of modern human T2a1 (KY671094.1) (60% of reads) and modern human U5a2a (HM229344.1; 40%) at nominal mean coverage depths of 0.5X, 1X, 2X, 4X, 8X, 16X, 32X.** Both sources carry ancient DNA damage (U5a2a with the damage rates scaled by 0.75). For each coverage depth, the upper panel shows the phylogenetic placement posterior of both sources for the best of five chains ($k = 2$, 100,000 iterations each), colored by source (dot area: branch posterior probability; swelling: posterior density of the position along the branch; bold tips: highest-posterior branches), and the lower panel the posterior densities of the mixture proportions $\Theta$ with median and 95% credible interval. T2a1 is expected on the T1a1a1 branch, the only haplogroup T haplotype in the reference; U5a2a, for which no U5 haplotype is present in the reference, is expected within the U clade. Modal branch of each source in the best chain ($p$: its posterior probability) and $\Theta$ median [95% CI]: (A,B) 0.5X: T1a1a1 ($p = 0.93$), $\Theta = 0.87$ [0.74, 0.96]; Node45 (stem of the Denisovan clade; $p = 0.96$), $\Theta = 0.13$ [0.04, 0.26]; (C,D) 1X: X2c1b ($p = 0.96$), $\Theta = 0.54$ [0.22, 0.79]; T1a1a1 ($p = 1.00$), $\Theta = 0.46$ [0.21, 0.78]; (E,F) 2X: U6a6a ($p = 0.90$), $\Theta = 0.62$ [0.36, 0.79]; T1a1a1 ($p = 1.00$), $\Theta = 0.38$ [0.21, 0.64]; (G,H) 4X: U6a6a ($p = 0.76$), $\Theta = 0.65$ [0.52, 0.77]; T1a1a1 ($p = 1.00$), $\Theta = 0.35$ [0.23, 0.48]; (I,J) 8X: U6a6a ($p = 0.52$), $\Theta = 0.59$ [0.50, 0.69]; T1a1a1 ($p = 1.00$), $\Theta = 0.41$ [0.31, 0.50]; (K,L) 16X: U6a6a ($p = 0.99$), $\Theta = 0.59$ [0.53, 0.65]; T1a1a1 ($p = 1.00$), $\Theta = 0.41$ [0.35, 0.47]; (M,N) 32X: U6a6a ($p = 1.00$), $\Theta = 0.59$ [0.55, 0.63]; T1a1a1 ($p = 1.00$), $\Theta = 0.41$ [0.37, 0.45].

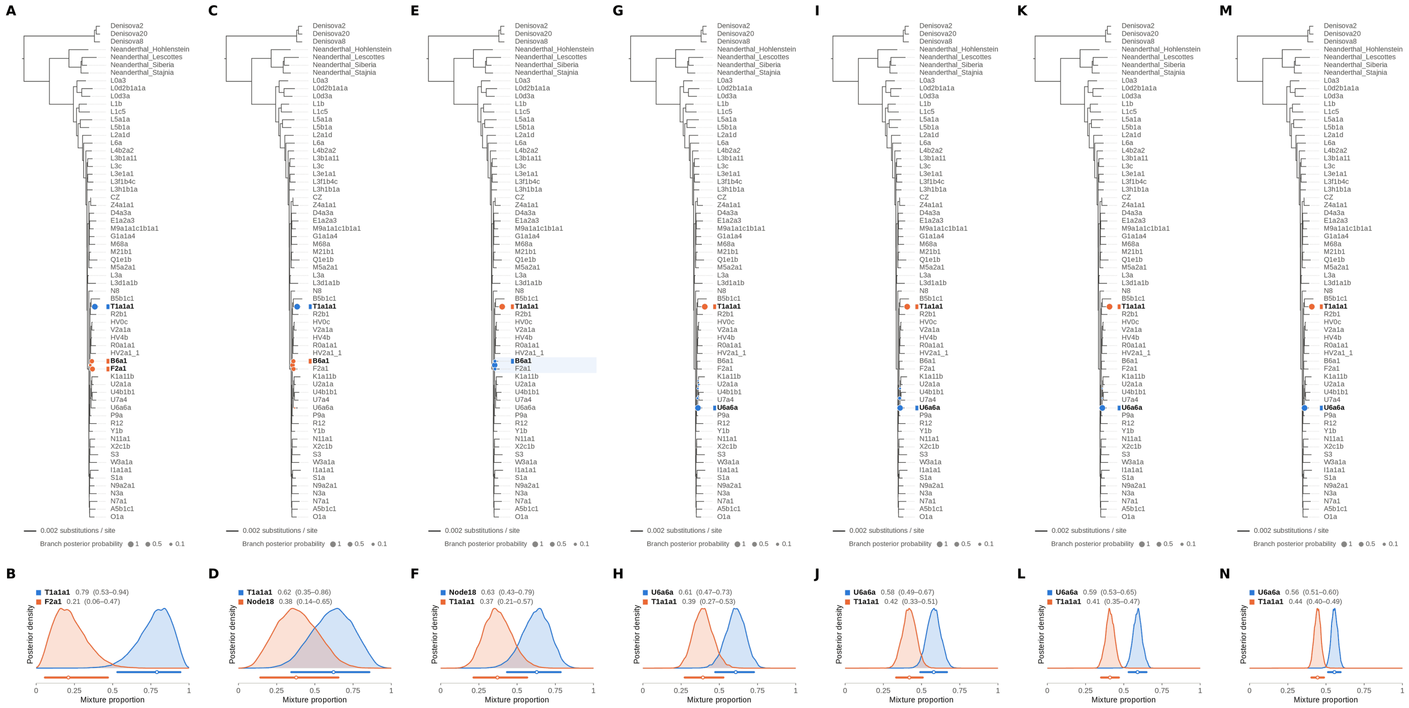

Figure S109: **Two-source simulated mixture of modern human T2a1 (KY671094.1) (65% of reads) and modern human U5a2a (HM229344.1; 35%) at nominal mean coverage depths of 0.5X, 1X, 2X, 4X, 8X, 16X, 32X.** Both sources carry ancient DNA damage (U5a2a with the damage rates scaled by 0.75). For each coverage depth, the upper panel shows the phylogenetic placement posterior of both sources for the best of five chains ($k = 2$, 100,000 iterations each), colored by source (dot area: branch posterior probability; swelling: posterior density of the position along the branch; bold tips: highest-posterior branches), and the lower panel the posterior densities of the mixture proportions $\Theta$ with median and 95% credible interval. T2a1 is expected on the T1a1a1 branch, the only haplogroup T haplotype in the reference; U5a2a, for which no U5 haplotype is present in the reference, is expected within the U clade. Modal branch of each source in the best chain ($p$: its posterior probability) and $\Theta$ median [95% CI]: (A,B) 0.5X: T1a1a1 ($p = 1.00$), $\Theta = 0.79$ [0.53, 0.94]; F2a1 ($p = 0.60$), $\Theta = 0.21$ [0.06, 0.47]; (C,D) 1X: T1a1a1 ($p = 1.00$), $\Theta = 0.62$ [0.35, 0.86]; Node18 (internal branch ancestral to B6a1 and F2a1; $p = 0.41$), $\Theta = 0.38$ [0.14, 0.65]; (E,F) 2X: Node18 (internal branch ancestral to B6a1 and F2a1; $p = 0.85$), $\Theta = 0.63$ [0.43, 0.79]; T1a1a1 ($p = 1.00$), $\Theta = 0.37$ [0.21, 0.57]; (G,H) 4X: U6a6a ($p = 0.97$), $\Theta = 0.61$ [0.47, 0.73]; T1a1a1 ($p = 1.00$), $\Theta = 0.39$ [0.27, 0.53]; (I,J) 8X: U6a6a ($p = 0.91$), $\Theta = 0.58$ [0.49, 0.67]; T1a1a1 ($p = 1.00$), $\Theta = 0.42$ [0.33, 0.51]; (K,L) 16X: U6a6a ($p = 1.00$), $\Theta = 0.59$ [0.53, 0.65]; T1a1a1 ($p = 1.00$), $\Theta = 0.41$ [0.35, 0.47]; (M,N) 32X: U6a6a ($p = 1.00$), $\Theta = 0.56$ [0.51, 0.60]; T1a1a1 ($p = 1.00$), $\Theta = 0.44$ [0.40, 0.49].

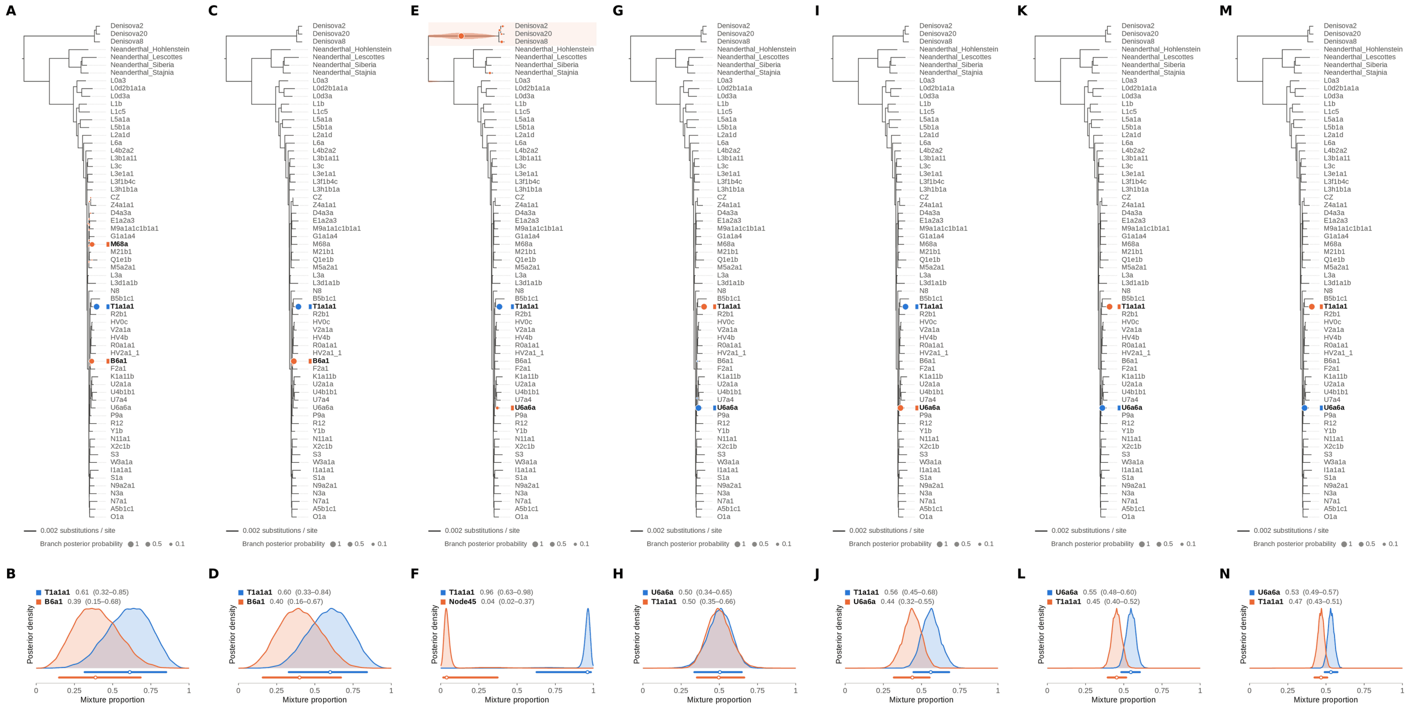

Figure S110: **Two-source simulated mixture of modern human T2a1 (KY671094.1) (70% of reads) and modern human U5a2a (HM229344.1; 30%) at nominal mean coverage depths of 0.5X, 1X, 2X, 4X, 8X, 16X, 32X.** Both sources carry ancient DNA damage (U5a2a with the damage rates scaled by 0.75). For each coverage depth, the upper panel shows the phylogenetic placement posterior of both sources for the best of five chains ($k = 2$, 100,000 iterations each), colored by source (dot area: branch posterior probability; swelling: posterior density of the position along the branch; bold tips: highest-posterior branches), and the lower panel the posterior densities of the mixture proportions $\Theta$ with median and 95% credible interval. T2a1 is expected on the T1a1a1 branch, the only haplogroup T haplotype in the reference; U5a2a, for which no U5 haplotype is present in the reference, is expected within the U clade. Modal branch of each source in the best chain ($p$: its posterior probability) and $\Theta$ median [95% CI]: (A,B) 0.5X: T1a1a1 ($p = 1.00$), $\Theta = 0.61$ [0.32, 0.85]; B6a1 ($p = 0.53$), $\Theta = 0.39$ [0.15, 0.68]; (C,D) 1X: T1a1a1 ($p = 1.00$), $\Theta = 0.60$ [0.33, 0.84]; B6a1 ($p = 1.00$), $\Theta = 0.40$ [0.16, 0.67]; (E,F) 2X: T1a1a1 ($p = 1.00$), $\Theta = 0.96$ [0.63, 0.98]; Node45 (stem of the Denisovan clade; $p = 0.84$), $\Theta = 0.04$ [0.02, 0.37]; (G,H) 4X: U6a6a ($p = 1.00$), $\Theta = 0.50$ [0.34, 0.65]; T1a1a1 ($p = 1.00$), $\Theta = 0.50$ [0.35, 0.66]; (I,J) 8X: T1a1a1 ($p = 1.00$), $\Theta = 0.56$ [0.45, 0.68]; U6a6a ($p = 1.00$), $\Theta = 0.44$ [0.32, 0.55]; (K,L) 16X: U6a6a ($p = 1.00$), $\Theta = 0.55$ [0.48, 0.60]; T1a1a1 ($p = 1.00$), $\Theta = 0.45$ [0.40, 0.52]; (M,N) 32X: U6a6a ($p = 1.00$), $\Theta = 0.53$ [0.49, 0.57]; T1a1a1 ($p = 1.00$), $\Theta = 0.47$ [0.43, 0.51].

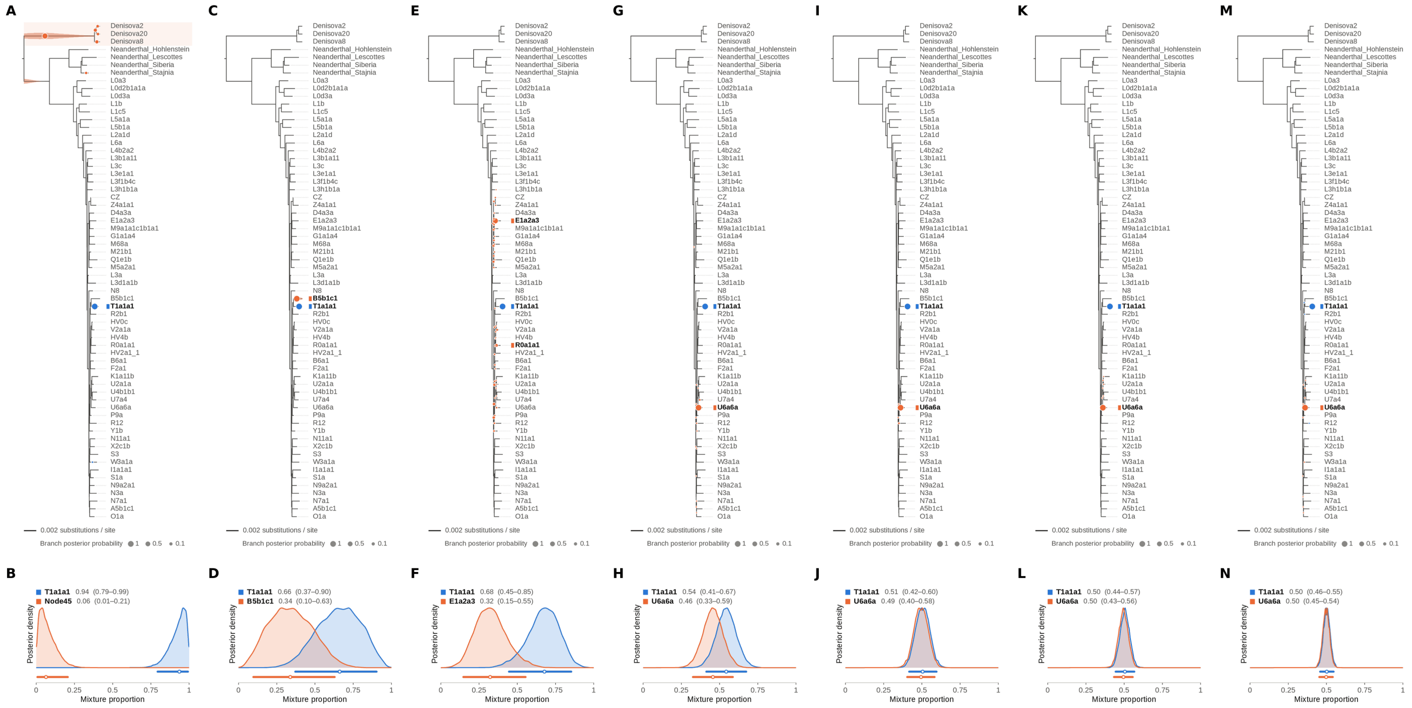


Figure S111: **Two-source simulated mixture of modern human T2a1 (KY671094.1) (75% of reads) and modern human U5a2a (HM229344.1; 25%) at nominal mean coverage depths of 0.5X, 1X, 2X, 4X, 8X, 16X, 32X.** Both sources carry ancient DNA damage (U5a2a with the damage rates scaled by 0.75). For each coverage depth, the upper panel shows the phylogenetic placement posterior of both sources for the best of five chains ($k = 2$, 100,000 iterations each), colored by source (dot area: branch posterior probability; swelling: posterior density of the position along the branch; bold tips: highest-posterior branches), and the lower panel the posterior densities of the mixture proportions $\Theta$ with median and 95% credible interval. T2a1 is expected on the T1a1a1 branch, the only haplogroup T haplotype in the reference; U5a2a, for which no U5 haplotype is present in the reference, is expected within the U clade. Modal branch of each source in the best chain ($p$: its posterior probability) and $\Theta$ median [95% CI]: (A,B) 0.5X: T1a1a1 ($p = 1.00$), $\Theta = 0.94$ [0.79, 0.99]; Node45 (stem of the Denisovan clade; $p = 0.66$), $\Theta = 0.06$ [0.01, 0.21]; (C,D) 1X: T1a1a1 ($p = 1.00$), $\Theta = 0.66$ [0.37, 0.90]; B5b1c1 ($p = 1.00$), $\Theta = 0.34$ [0.10, 0.63]; (E,F) 2X: T1a1a1 ($p = 1.00$), $\Theta = 0.68$ [0.45, 0.85]; E1a2a3 ($p = 0.65$), $\Theta = 0.32$ [0.15, 0.55]; (G,H) 4X: T1a1a1 ($p = 1.00$), $\Theta = 0.54$ [0.41, 0.67]; U6a6a ($p = 0.99$), $\Theta = 0.46$ [0.33, 0.59]; (I,J) 8X: T1a1a1 ($p = 1.00$), $\Theta = 0.51$ [0.42, 0.60]; U6a6a ($p = 0.99$), $\Theta = 0.49$ [0.40, 0.58]; (K,L) 16X: T1a1a1 ($p = 1.00$), $\Theta = 0.50$ [0.44, 0.57]; U6a6a ($p = 1.00$), $\Theta = 0.50$ [0.43, 0.56]; (M,N) 32X: T1a1a1 ($p = 1.00$), $\Theta = 0.50$ [0.46, 0.55]; U6a6a ($p = 1.00$), $\Theta = 0.50$ [0.45, 0.54].

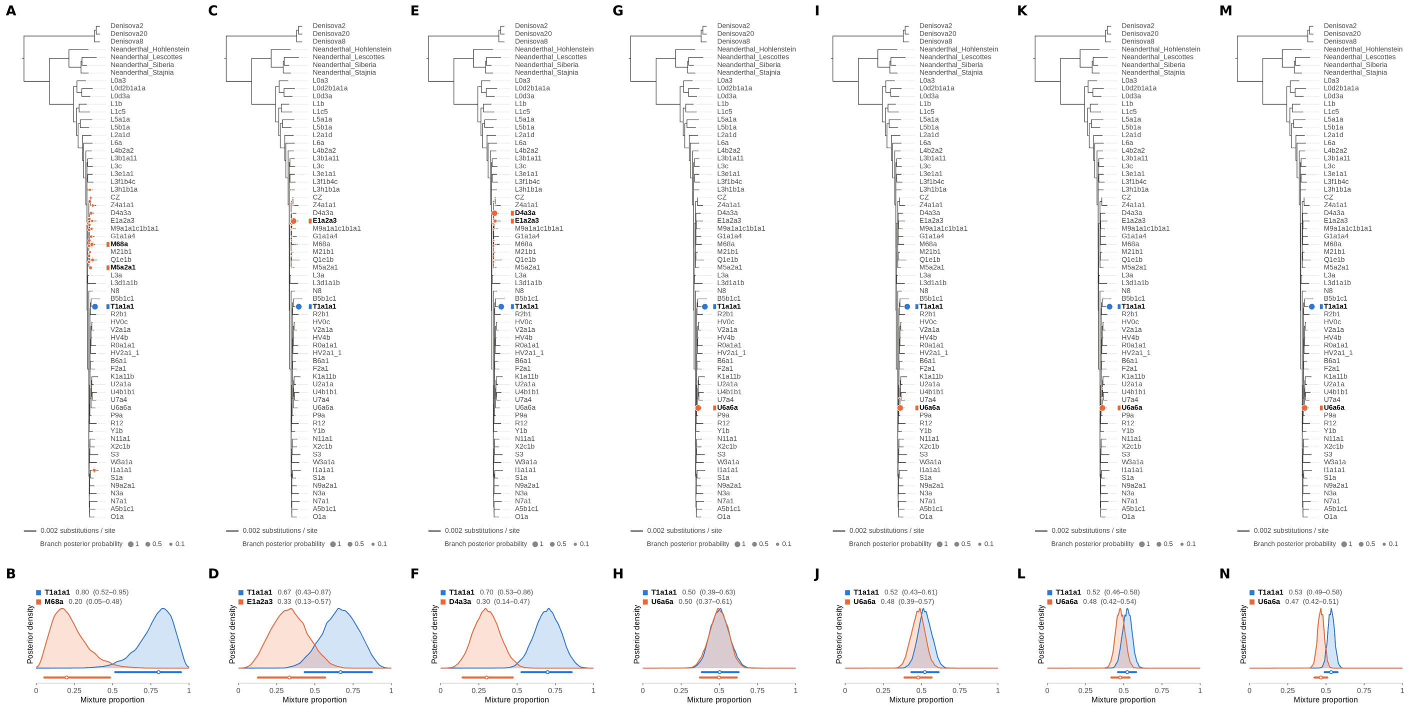


Figure S112: **Two-source simulated mixture of modern human T2a1 (KY671094.1) (80% of reads) and modern human U5a2a (HM229344.1; 20%) at nominal mean coverage depths of 0.5X, 1X, 2X, 4X, 8X, 16X, 32X.** Both sources carry ancient DNA damage (U5a2a with the damage rates scaled by 0.75). For each coverage depth, the upper panel shows the phylogenetic placement posterior of both sources for the best of five chains ($k = 2$, 100,000 iterations each), colored by source (dot area: branch posterior probability; swelling: posterior density of the position along the branch; bold tips: highest-posterior branches), and the lower panel the posterior densities of the mixture proportions $\Theta$ with median and 95% credible interval. T2a1 is expected on the T1a1a1 branch, the only haplogroup T haplotype in the reference; U5a2a, for which no U5 haplotype is present in the reference, is expected within the U clade. Modal branch of each source in the best chain ($p$: its posterior probability) and $\Theta$ median [95% CI]: (A,B) 0.5X: T1a1a1 ($p = 1.00$), $\Theta = 0.80$ [0.52, 0.95]; M68a ($p = 0.09$), $\Theta = 0.20$ [0.05, 0.48]; (C,D) 1X: T1a1a1 ($p = 1.00$), $\Theta = 0.67$ [0.43, 0.87]; E1a2a3 ($p = 0.99$), $\Theta = 0.33$ [0.13, 0.57]; (E,F) 2X: T1a1a1 ($p = 1.00$), $\Theta = 0.70$ [0.53, 0.86]; D4a3a ($p = 0.85$), $\Theta = 0.30$ [0.14, 0.47]; (G,H) 4X: T1a1a1 ($p = 1.00$), $\Theta = 0.50$ [0.39, 0.63]; U6a6a ($p = 1.00$), $\Theta = 0.50$ [0.37, 0.61]; (I,J) 8X: T1a1a1 ($p = 1.00$), $\Theta = 0.52$ [0.43, 0.61]; U6a6a ($p = 1.00$), $\Theta = 0.48$ [0.39, 0.57]; (K,L) 16X: T1a1a1 ($p = 1.00$), $\Theta = 0.52$ [0.46, 0.58]; U6a6a ($p = 1.00$), $\Theta = 0.48$ [0.42, 0.54]; (M,N) 32X: T1a1a1 ($p = 1.00$), $\Theta = 0.53$ [0.49, 0.58]; U6a6a ($p = 1.00$), $\Theta = 0.47$ [0.42, 0.51].

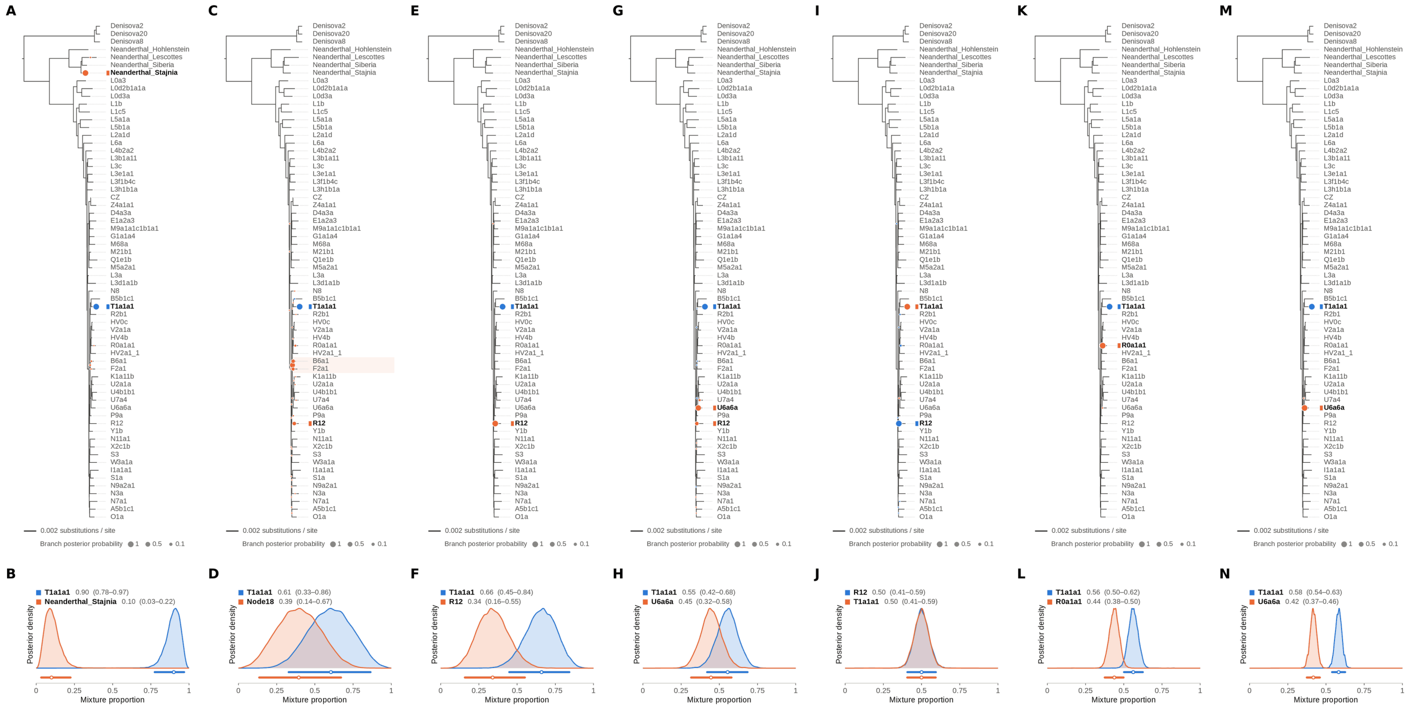


Figure S113: **Two-source simulated mixture of modern human T2a1 (KY671094.1) (85% of reads) and modern human U5a2a (HM229344.1; 15%) at nominal mean coverage depths of 0.5X, 1X, 2X, 4X, 8X, 16X, 32X.** Both sources carry ancient DNA damage (U5a2a with the damage rates scaled by 0.75). For each coverage depth, the upper panel shows the phylogenetic placement posterior of both sources for the best of five chains ($k = 2$, 100,000 iterations each), colored by source (dot area: branch posterior probability; swelling: posterior density of the position along the branch; bold tips: highest-posterior branches), and the lower panel the posterior densities of the mixture proportions $\Theta$ with median and 95% credible interval. T2a1 is expected on the T1a1a1 branch, the only haplogroup T haplotype in the reference; U5a2a, for which no U5 haplotype is present in the reference, is expected within the U clade. Modal branch of each source in the best chain ($p$: its posterior probability) and $\Theta$ median [95% CI]: (A,B) 0.5X: T1a1a1 ($p = 1.00$), $\Theta = 0.90$ [0.78, 0.97]; Neanderthal_Stajnia ($p = 0.98$), $\Theta = 0.10$ [0.03, 0.22]; (C,D) 1X: T1a1a1 ($p = 1.00$), $\Theta = 0.61$ [0.33, 0.86]; Node18 (internal branch ancestral to B6a1 and F2a1; $p = 0.57$), $\Theta = 0.39$ [0.14, 0.67]; (E,F) 2X: T1a1a1 ($p = 1.00$), $\Theta = 0.66$ [0.45, 0.84]; R12 ($p = 1.00$), $\Theta = 0.34$ [0.16, 0.55]; (G,H) 4X: T1a1a1 ($p = 1.00$), $\Theta = 0.55$ [0.42, 0.68]; U6a6a ($p = 0.83$), $\Theta = 0.45$ [0.32, 0.58]; (I,J) 8X: R12 ($p = 0.98$), $\Theta = 0.50$ [0.41, 0.59]; T1a1a1 ($p = 1.00$), $\Theta = 0.50$ [0.41, 0.59]; (K,L) 16X: T1a1a1 ($p = 1.00$), $\Theta = 0.56$ [0.50, 0.62]; R0a1a1 ($p = 1.00$), $\Theta = 0.44$ [0.38, 0.50]; (M,N) 32X: T1a1a1 ($p = 1.00$), $\Theta = 0.58$ [0.54, 0.63]; U6a6a ($p = 1.00$), $\Theta = 0.42$ [0.37, 0.46].

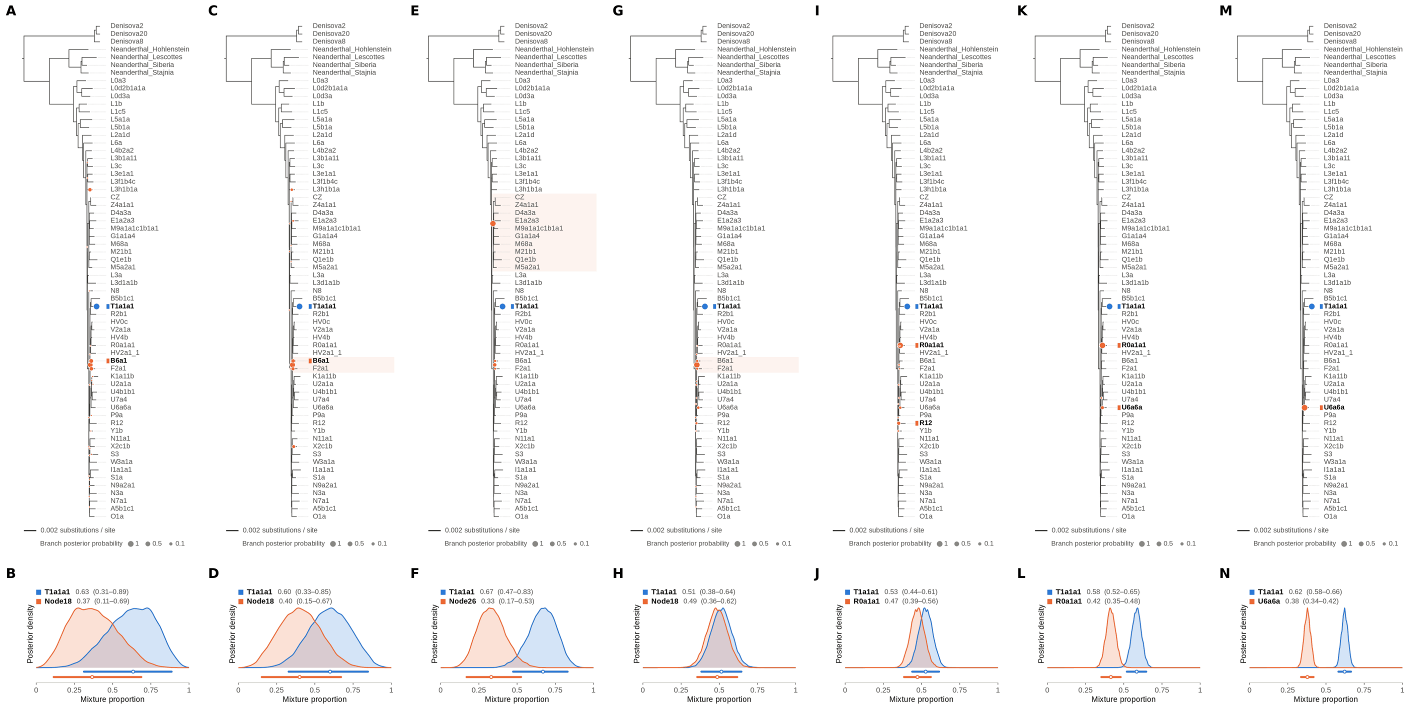


Figure S114: **Two-source simulated mixture of modern human T2a1 (KY671094.1) (90% of reads) and modern human U5a2a (HM229344.1; 10%) at nominal mean coverage depths of 0.5X, 1X, 2X, 4X, 8X, 16X, 32X.** Both sources carry ancient DNA damage (U5a2a with the damage rates scaled by 0.75). For each coverage depth, the upper panel shows the phylogenetic placement posterior of both sources for the best of five chains ($k = 2$, 100,000 iterations each), colored by source (dot area: branch posterior probability; swelling: posterior density of the position along the branch; bold tips: highest-posterior branches), and the lower panel the posterior densities of the mixture proportions $\Theta$ with median and 95% credible interval. T2a1 is expected on the T1a1a1 branch, the only haplogroup T haplotype in the reference; U5a2a, for which no U5 haplotype is present in the reference, is expected within the U clade. Modal branch of each source in the best chain ($p$: its posterior probability) and $\Theta$ median [95% CI]: (A,B) 0.5X: T1a1a1 ($p = 1.00$), $\Theta = 0.63$ [0.31, 0.89]; Node18 (internal branch ancestral to B6a1 and F2a1; $p = 0.48$), $\Theta = 0.37$ [0.11, 0.69]; (C,D) 1X: T1a1a1 ($p = 1.00$), $\Theta = 0.60$ [0.33, 0.85]; Node18 (internal branch ancestral to B6a1 and F2a1; $p = 0.63$), $\Theta = 0.40$ [0.15, 0.67]; (E,F) 2X: T1a1a1 ($p = 1.00$), $\Theta = 0.67$ [0.47, 0.83]; Node26 (stem of the M clade; $p = 0.83$), $\Theta = 0.33$ [0.17, 0.53]; (G,H) 4X: T1a1a1 ($p = 1.00$), $\Theta = 0.51$ [0.38, 0.64]; Node18 (internal branch ancestral to B6a1 and F2a1; $p = 0.89$), $\Theta = 0.49$ [0.36, 0.62]; (I,J) 8X: T1a1a1 ($p = 1.00$), $\Theta = 0.53$ [0.44, 0.61]; R0a1a1 ($p = 0.84$), $\Theta = 0.47$ [0.39, 0.56]; (K,L) 16X: T1a1a1 ($p = 1.00$), $\Theta = 0.58$ [0.52, 0.65]; R0a1a1 ($p = 0.91$), $\Theta = 0.42$ [0.35, 0.48]; (M,N) 32X: T1a1a1 ($p = 1.00$), $\Theta = 0.62$ [0.58, 0.66]; U6a6a ($p = 1.00$), $\Theta = 0.38$ [0.34, 0.42].

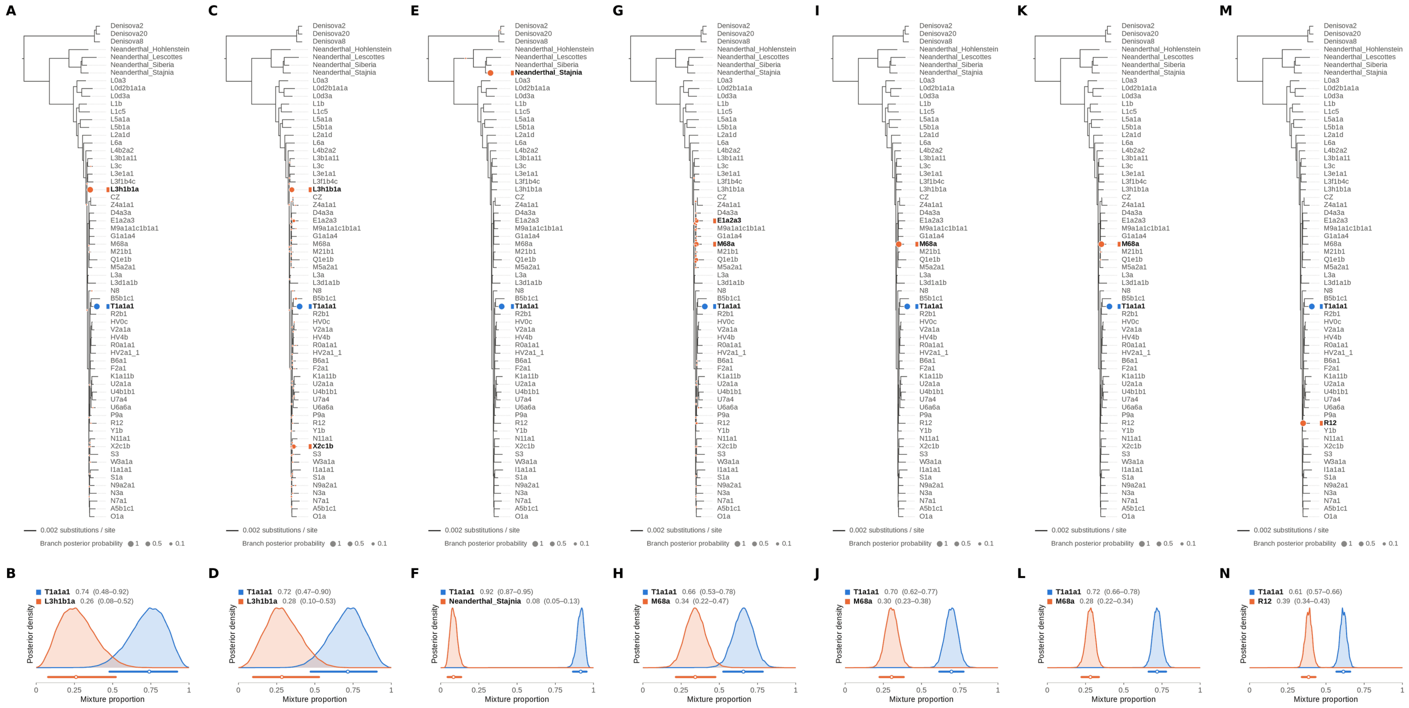

Figure S115: **Two-source simulated mixture of modern human T2a1 (KY671094.1) (95% of reads) and modern human U5a2a (HM229344.1; 5%) at nominal mean coverage depths of 0.5X, 1X, 2X, 4X, 8X, 16X, 32X.** Both sources carry ancient DNA damage (U5a2a with the damage rates scaled by 0.75). For each coverage depth, the upper panel shows the phylogenetic placement posterior of both sources for the best of five chains ($k = 2$, 100,000 iterations each), colored by source (dot area: branch posterior probability; swelling: posterior density of the position along the branch; bold tips: highest-posterior branches), and the lower panel the posterior densities of the mixture proportions $\Theta$ with median and 95% credible interval. T2a1 is expected on the T1a1a1 branch, the only haplogroup T haplotype in the reference; U5a2a, for which no U5 haplotype is present in the reference, is expected within the U clade. Modal branch of each source in the best chain ($p$: its posterior probability) and $\Theta$ median [95% CI]: (A,B) 0.5X: T1a1a1 ($p = 1.00$), $\Theta = 0.74$ [0.48, 0.92]; L3h1b1a ($p = 1.00$), $\Theta = 0.26$ [0.08, 0.52]; (C,D) 1X: T1a1a1 ($p = 1.00$), $\Theta = 0.72$ [0.47, 0.90]; L3h1b1a ($p = 0.57$), $\Theta = 0.28$ [0.10, 0.53]; (E,F) 2X: T1a1a1 ($p = 1.00$), $\Theta = 0.92$ [0.87, 0.95]; Neanderthal_Stajnia ($p = 1.00$), $\Theta = 0.08$ [0.05, 0.13]; (G,H) 4X: T1a1a1 ($p = 1.00$), $\Theta = 0.66$ [0.53, 0.78]; M68a ($p = 0.46$), $\Theta = 0.34$ [0.22, 0.47]; (I,J) 8X: T1a1a1 ($p = 1.00$), $\Theta = 0.70$ [0.62, 0.77]; M68a ($p = 1.00$), $\Theta = 0.30$ [0.23, 0.38]; (K,L) 16X: T1a1a1 ($p = 1.00$), $\Theta = 0.72$ [0.66, 0.78]; M68a ($p = 1.00$), $\Theta = 0.28$ [0.22, 0.34]; (M,N) 32X: T1a1a1 ($p = 1.00$), $\Theta = 0.61$ [0.57, 0.66]; R12 ($p = 1.00$), $\Theta = 0.39$ [0.34, 0.43].

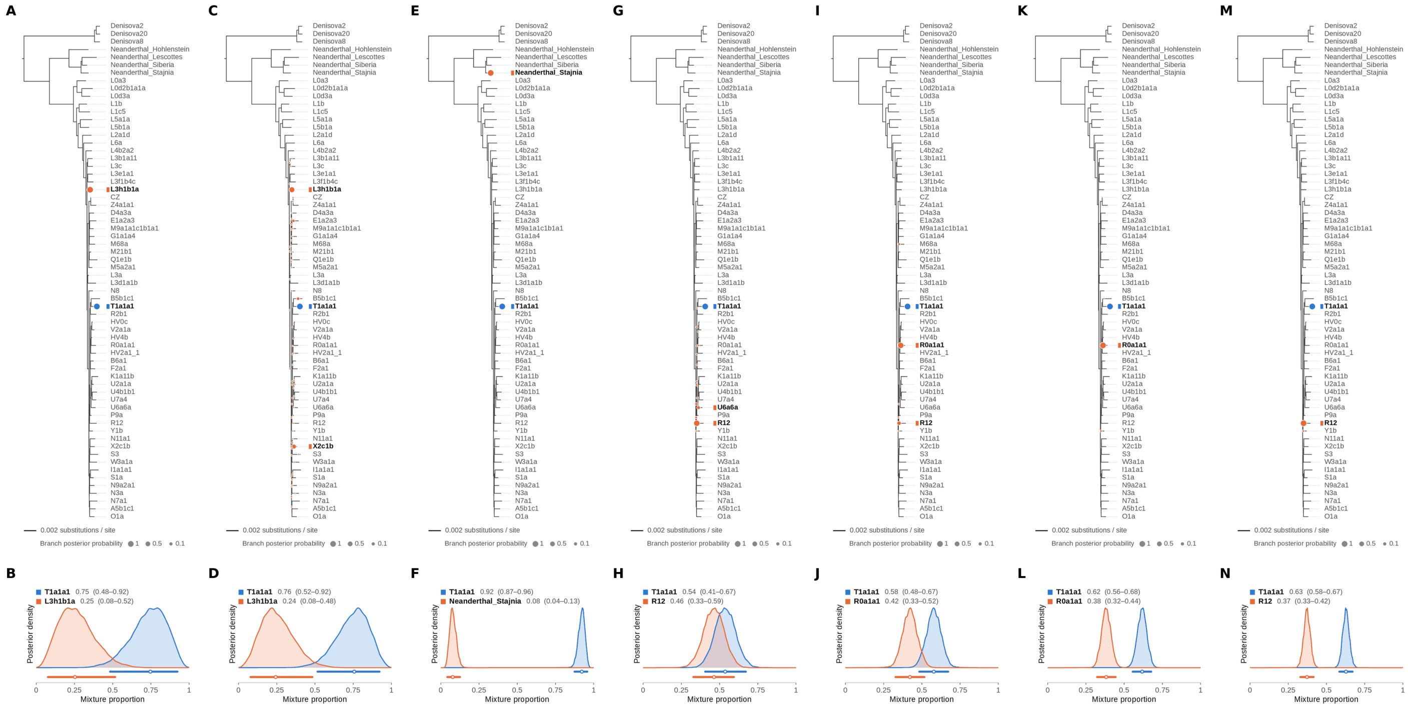

Figure S116: **Two-source simulated mixture of modern human T2a1 (KY671094.1) (99% of reads) and modern human U5a2a (HM229344.1; 1%) at nominal mean coverage depths of 0.5X, 1X, 2X, 4X, 8X, 16X, 32X.** Both sources carry ancient DNA damage (U5a2a with the damage rates scaled by 0.75). For each coverage depth, the upper panel shows the phylogenetic placement posterior of both sources for the best of five chains ($k = 2$, 100,000 iterations each), colored by source (dot area: branch posterior probability; swelling: posterior density of the position along the branch; bold tips: highest-posterior branches), and the lower panel the posterior densities of the mixture proportions $\Theta$ with median and 95% credible interval. T2a1 is expected on the T1a1a1 branch, the only haplogroup T haplotype in the reference; U5a2a, for which no U5 haplotype is present in the reference, is expected within the U clade. Modal branch of each source in the best chain ($p$: its posterior probability) and $\Theta$ median [95% CI]: (A,B) 0.5X: T1a1a1 ($p = 1.00$), $\Theta = 0.75$ [0.48, 0.92]; L3h1b1a ($p = 1.00$), $\Theta = 0.25$ [0.08, 0.52]; (C,D) 1X: T1a1a1 ($p = 1.00$), $\Theta = 0.76$ [0.52, 0.92]; L3h1b1a ($p = 0.64$), $\Theta = 0.24$ [0.08, 0.48]; (E,F) 2X: T1a1a1 ($p = 1.00$), $\Theta = 0.92$ [0.87, 0.96]; Neanderthal_Stajnia ($p = 1.00$), $\Theta = 0.08$ [0.04, 0.13]; (G,H) 4X: T1a1a1 ($p = 1.00$), $\Theta = 0.54$ [0.41, 0.67]; R12 ($p = 0.86$), $\Theta = 0.46$ [0.33, 0.59]; (I,J) 8X: T1a1a1 ($p = 1.00$), $\Theta = 0.58$ [0.48, 0.67]; R0a1a1 ($p = 0.83$), $\Theta = 0.42$ [0.33, 0.52]; (K,L) 16X: T1a1a1 ($p = 1.00$), $\Theta = 0.62$ [0.56, 0.68]; R0a1a1 ($p = 0.99$), $\Theta = 0.38$ [0.32, 0.44]; (M,N) 32X: T1a1a1 ($p = 1.00$), $\Theta = 0.63$ [0.58, 0.67]; R12 ($p = 0.99$), $\Theta = 0.37$ [0.33, 0.42].

#### S7 Estimated versus Simulated Mixture Proportions

For each of the 735 two-source mixtures we compared the minor (lower) mixture proportion estimated by `TrailMix` with the simulated one, and with the estimates of two other methods (main Figure 5E and Supplementary Figures S117–S126). For `TrailMix` we took the posterior mean of the smaller of the two proportions in the best chain (selected as for the figures), with its 5th and 95th posterior percentiles. For the `RPVG` initialization, we took the read counts assigned by the `haplotype-transcripts` mode to the two haplotypes of the most probable haplotype pair, and computed the smaller of the two as a fraction of their sum. For `kallisto` [1] (version 0.51.1), we built an index ($k = 19$) of the 126 sequences of the `TrailMix` reference graph and quantified the reads of each mixture with `kallisto quant --single -l 50 -s 15`; the proportion is computed from the two reference sequences with the highest estimated read counts, in the same way as for `RPVG`. Neither `RPVG` nor `kallisto` provides an interval for these estimates. For the mixtures of an archaic and a modern source, we also summed the `kallisto` estimated read counts over the 12 archaic sequences of the index (the 7 archaic tips and the 5 ancestral sequences of the Neanderthal and Denisovan lineages) and compared this archaic fraction with the simulated share of Chagyrskaya 8 or Denisova 21: its RMSE over the 21 mixtures was 0.04–0.05 for Chagyrskaya 8 and 0.03–0.04 for Denisova 21 at every coverage depth, compared with 0.03–0.09 and 0.02–0.09, respectively, for `TrailMix`. The comparison is label-free: the smaller estimated proportion is compared with the smaller simulated proportion regardless of which source it was assigned to. The x-axis of each panel is the nominal simulated proportion; the realized proportion after downsampling differs from it by 0.03 on average at 0.5X and by less than 0.01 from 8X.

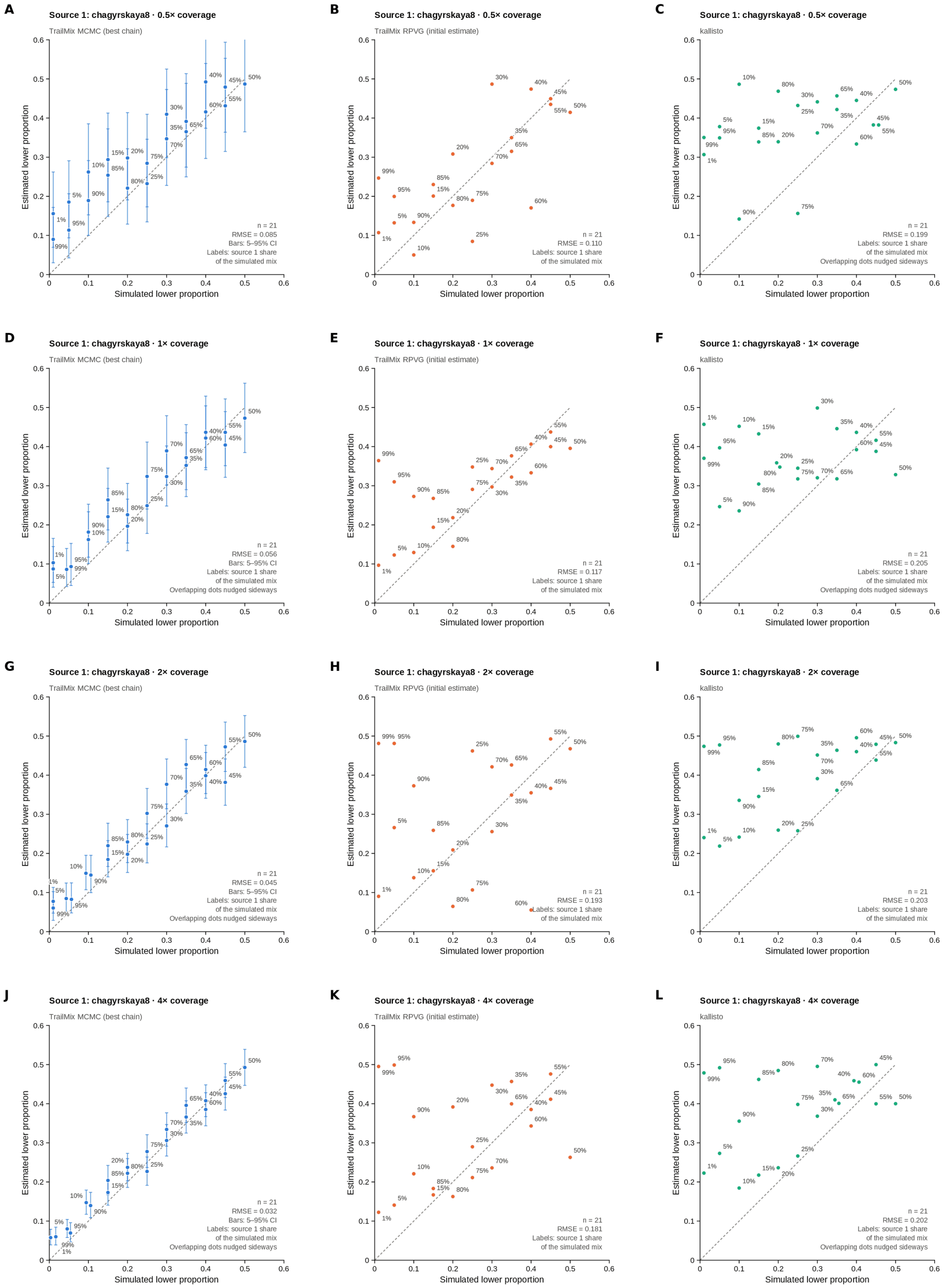


Figure S117: **Estimated versus simulated minor mixture proportion for Chagyrskaya 8 Neanderthal (MK388903.1) mixed with U5a2a at 21 proportions (1–99%), at nominal mean coverage depths of 0.5X, 1X, 2X, 4X (rows).** Columns: `TrailMix` MCMC (best chain; posterior mean with 5–95% interval), the `RPVG` estimate used to initialize `TrailMix`, and `kallisto`. The x-axis is the simulated (nominal) proportion of the less abundant source and the y-axis the corresponding estimate; each point is labeled with the share of Chagyrskaya 8 in the simulated mixture, the dashed line is the identity and the RMSE over the 21 mixtures is given in each panel. The `TrailMix` MCMC panels at 1X and 4X are also shown in main Figure 5.

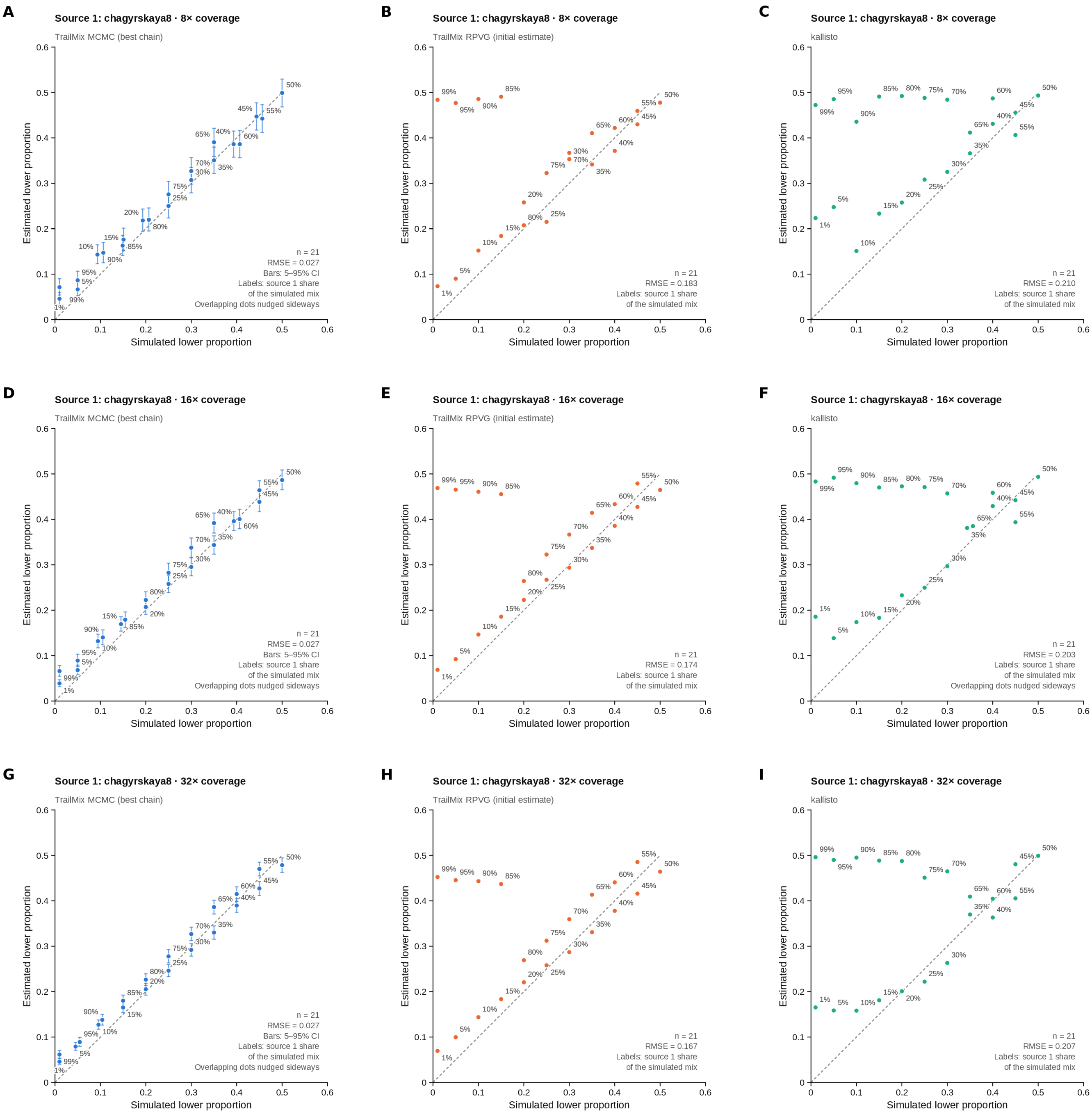


Figure S118: **Estimated versus simulated minor mixture proportion for Chagyrskaya 8 Neanderthal (MK388903.1) mixed with U5a2a at 21 proportions (1–99%), at nominal mean coverage depths of 8X, 16X, 32X (rows).** Columns: `TrailMix` MCMC (best chain; posterior mean with 5–95% interval), the `RPVG` estimate used to initialize `TrailMix`, and `kallisto`. The x-axis is the simulated (nominal) proportion of the less abundant source and the y-axis the corresponding estimate; each point is labeled with the share of Chagyrskaya 8 in the simulated mixture, the dashed line is the identity and the RMSE over the 21 mixtures is given in each panel. The `TrailMix` MCMC panels at 8X and 16X are also shown in main Figure 5.

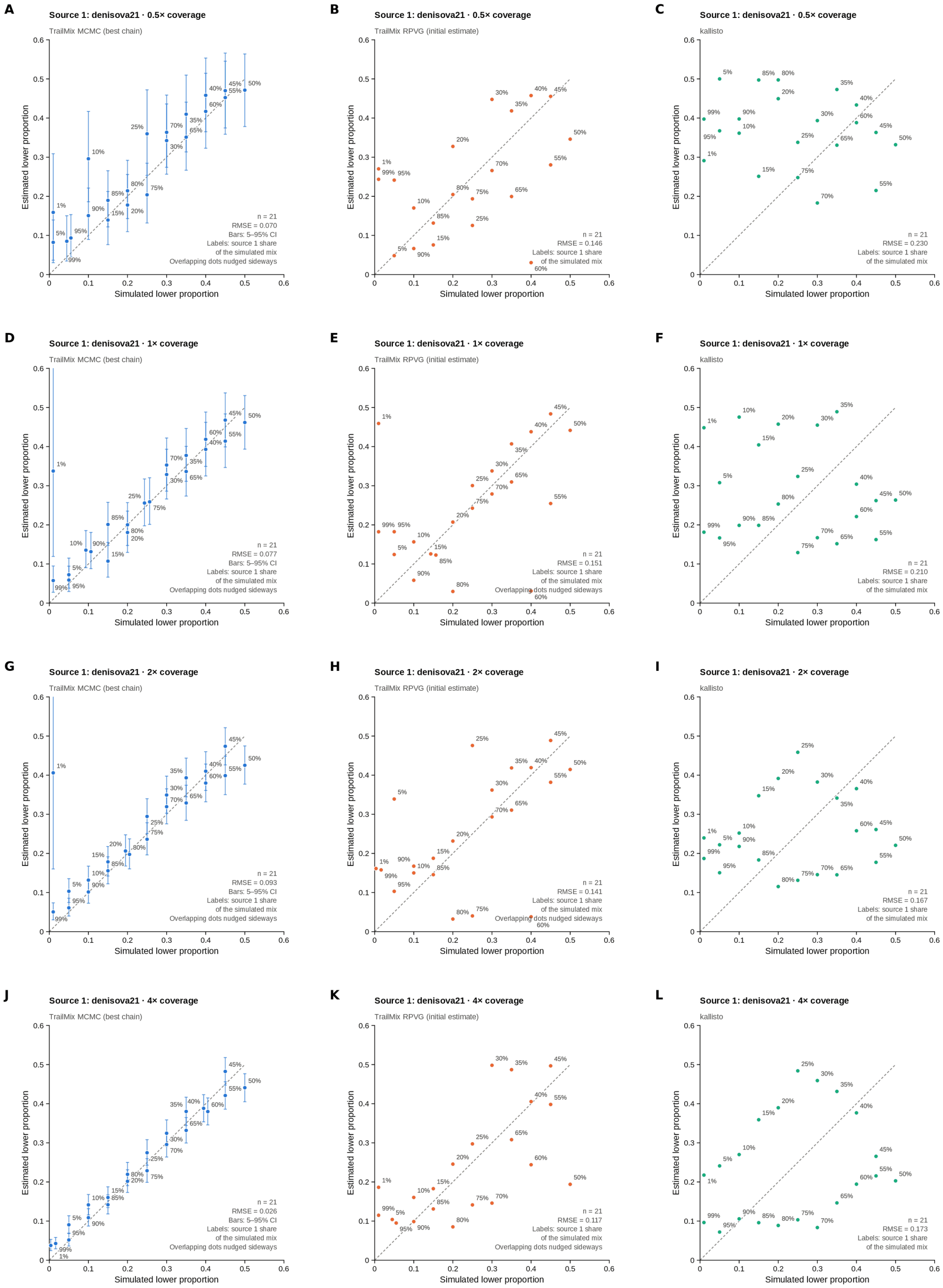


Figure S119: **Estimated versus simulated minor mixture proportion for Denisova 21 (MT576653.1) mixed with U5a2a at 21 proportions (1–99%), at nominal mean coverage depths of 0.5X, 1X, 2X, 4X (rows).** Columns: `TrailMix` MCMC (best chain; posterior mean with 5–95% interval), the `RPVG` estimate used to initialize `TrailMix`, and `kallisto`. The x-axis is the simulated (nominal) proportion of the less abundant source and the y-axis the corresponding estimate; each point is labeled with the share of Denisova 21 in the simulated mixture, the dashed line is the identity and the RMSE over the 21 mixtures is given in each panel.

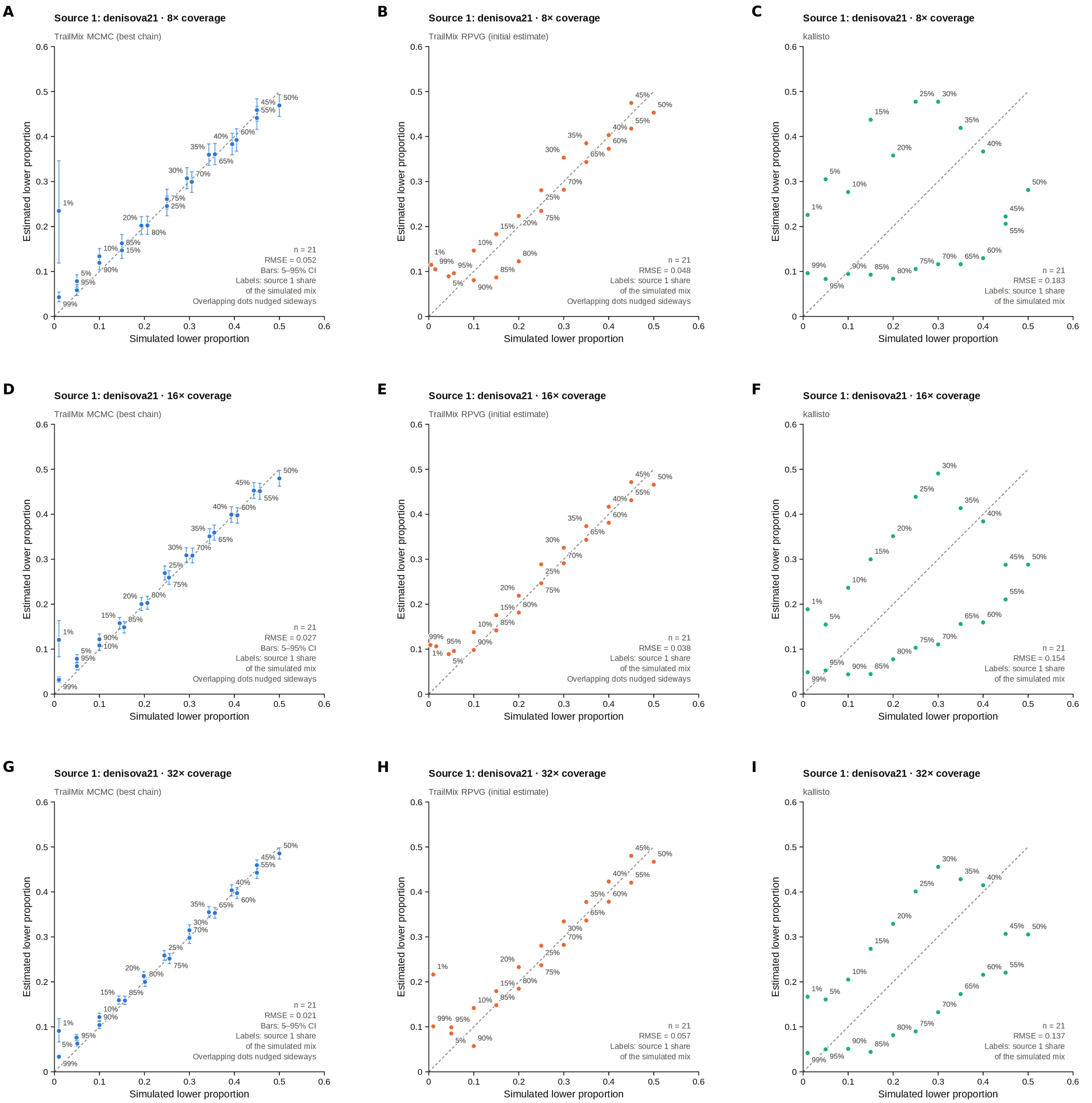


Figure S120: **Estimated versus simulated minor mixture proportion for Denisova 21 (MT576653.1) mixed with U5a2a at 21 proportions (1–99%), at nominal mean coverage depths of 8X, 16X, 32X (rows).** Columns: `TrailMix` MCMC (best chain; posterior mean with 5–95% interval), the `RPVG` estimate used to initialize `TrailMix`, and `kallisto`. The x-axis is the simulated (nominal) proportion of the less abundant source and the y-axis the corresponding estimate; each point is labeled with the share of Denisova 21 in the simulated mixture, the dashed line is the identity and the RMSE over the 21 mixtures is given in each panel.

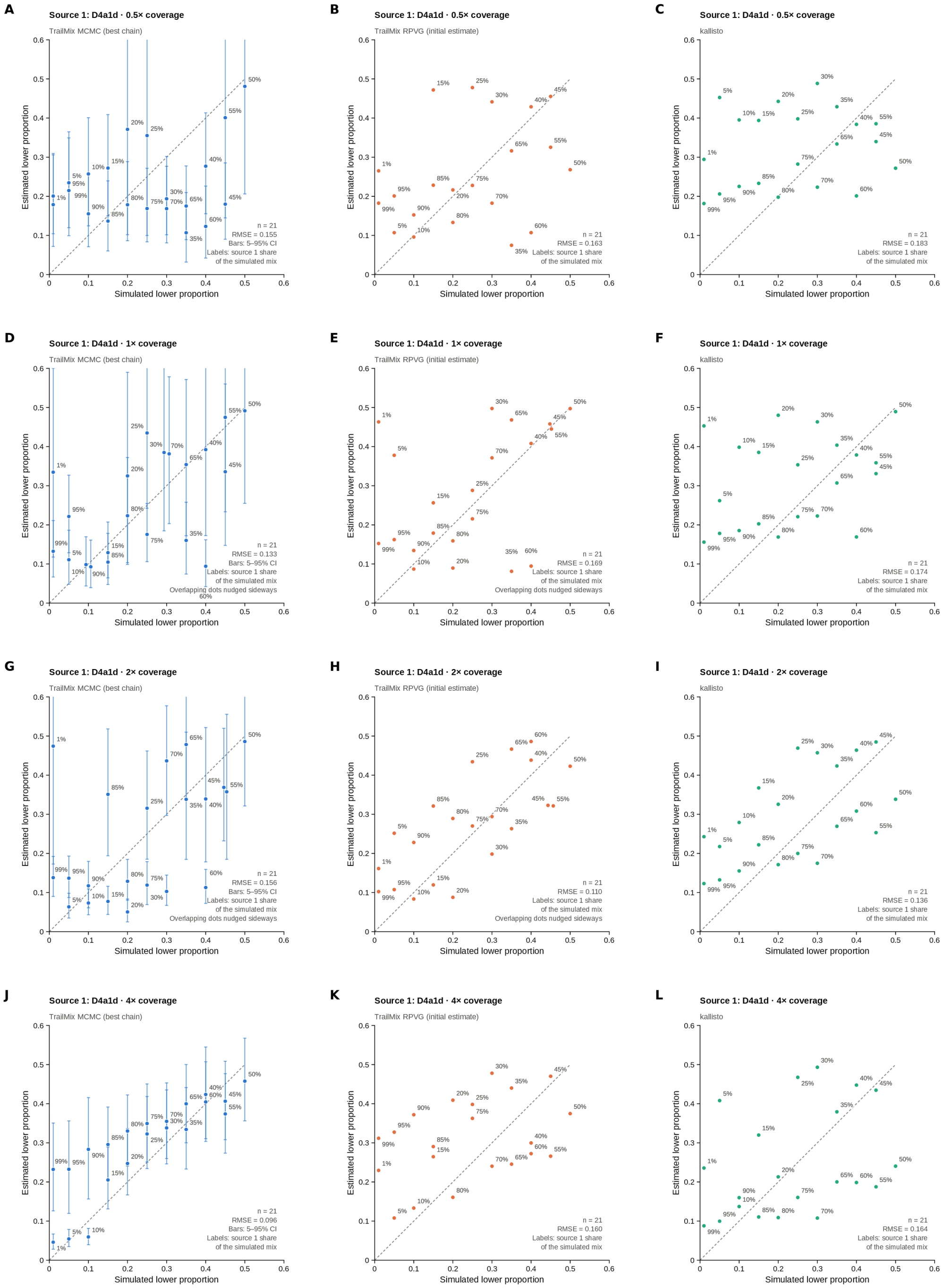

Figure S121: **Estimated versus simulated minor mixture proportion for modern human D4a1d (FJ951517.1) mixed with U5a2a at 21 proportions (1–99%), at nominal mean coverage depths of 0.5X, 1X, 2X, 4X (rows).** Columns: `TrailMix` MCMC (best chain; posterior mean with 5–95% interval), the `RPVG` estimate used to initialize `TrailMix`, and `kallisto`. The x-axis is the simulated (nominal) proportion of the less abundant source and the y-axis the corresponding estimate; each point is labeled with the share of D4a1d in the simulated mixture, the dashed line is the identity and the RMSE over the 21 mixtures is given in each panel.

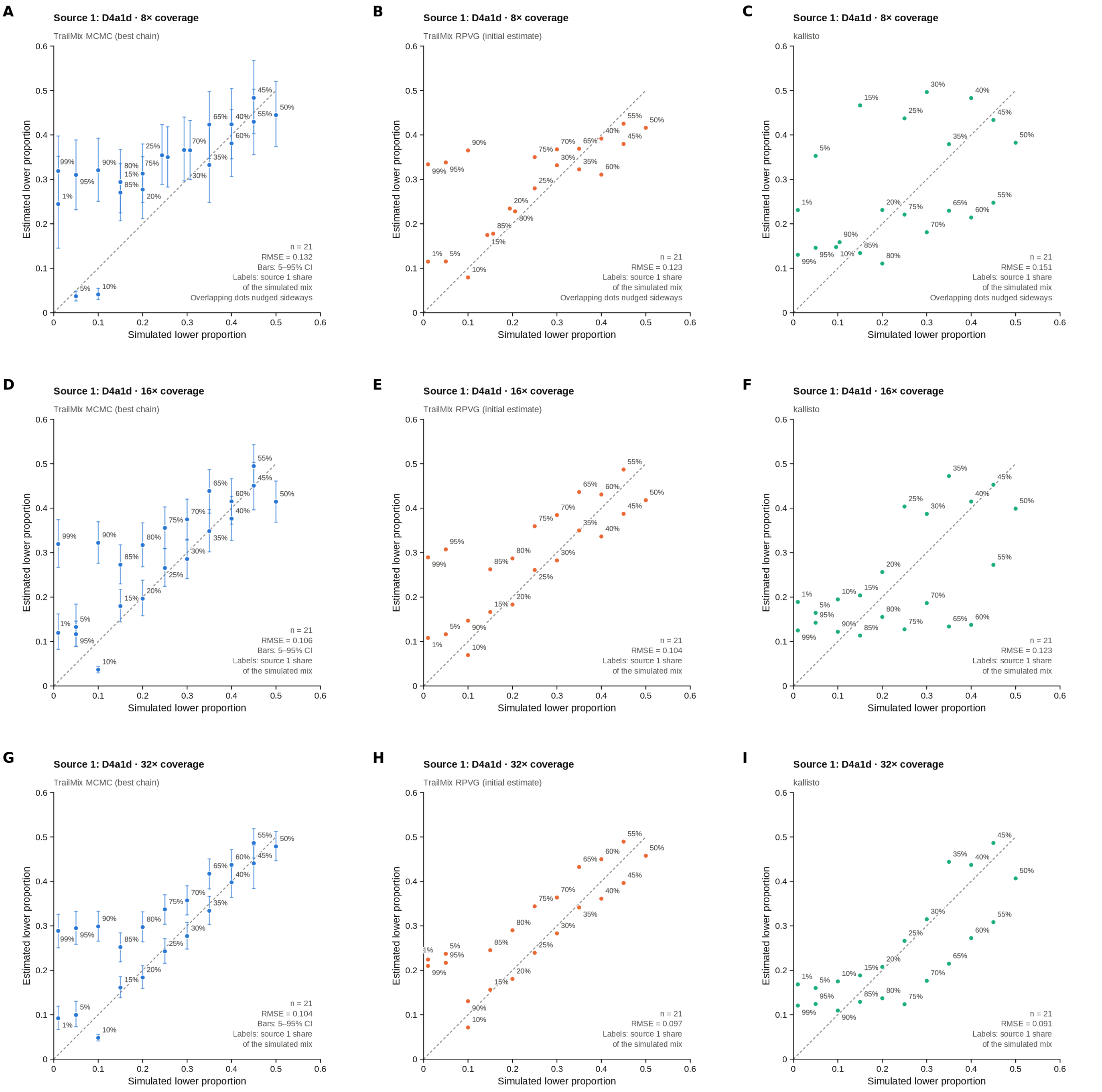


Figure S122: **Estimated versus simulated minor mixture proportion for modern human D4a1d (FJ951517.1) mixed with U5a2a at 21 proportions (1–99%), at nominal mean coverage depths of 8X, 16X, 32X (rows).** Columns: `TrailMix` MCMC (best chain; posterior mean with 5–95% interval), the `RPVG` estimate used to initialize `TrailMix`, and `kallisto`. The x-axis is the simulated (nominal) proportion of the less abundant source and the y-axis the corresponding estimate; each point is labeled with the share of D4a1d in the simulated mixture, the dashed line is the identity and the RMSE over the 21 mixtures is given in each panel.

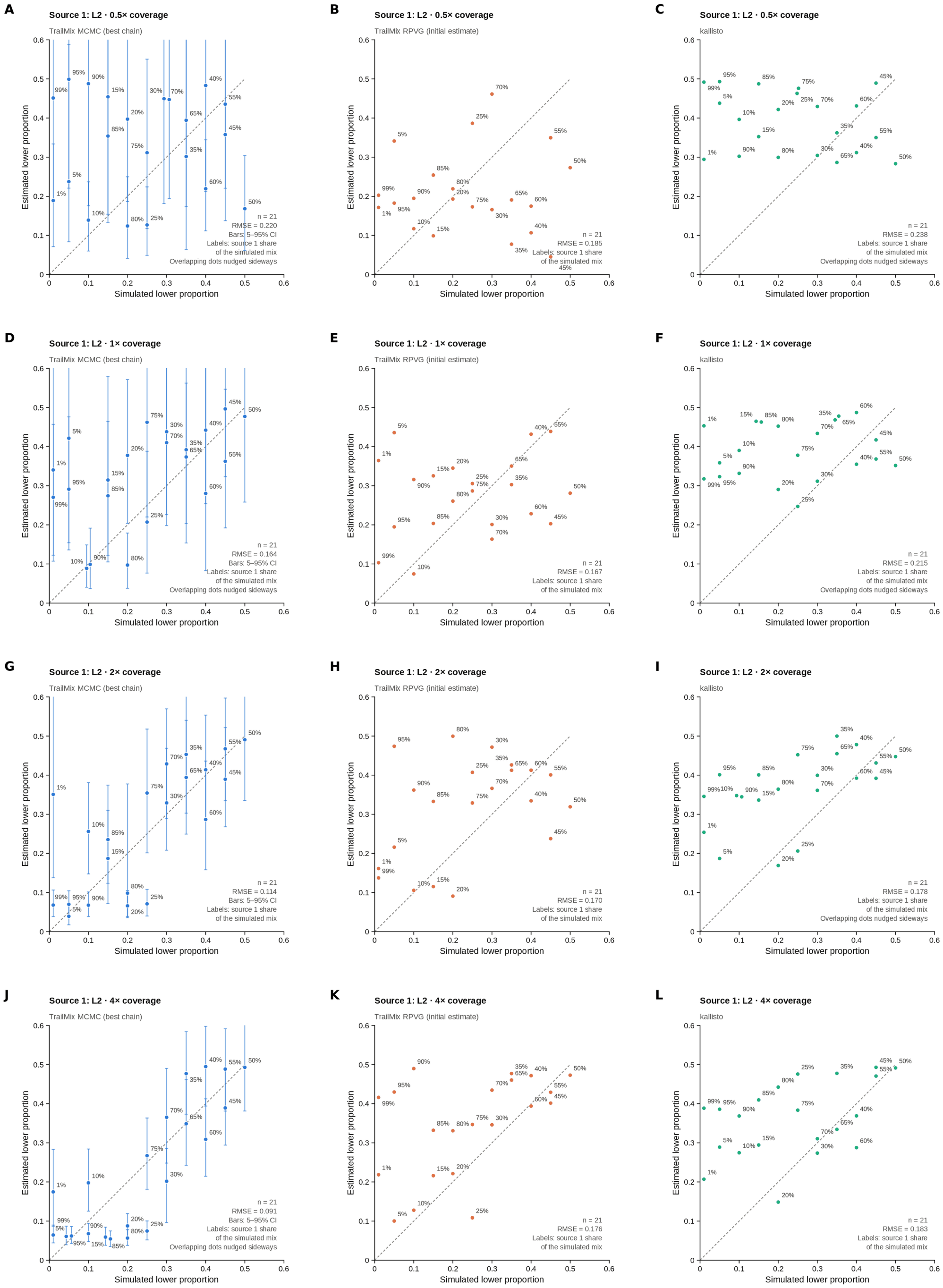


Figure S123: **Estimated versus simulated minor mixture proportion for modern human L2 (AF381981.1) mixed with U5a2a at 21 proportions (1–99%), at nominal mean coverage depths of 0.5X, 1X, 2X, 4X (rows).** Columns: `TrailMix` MCMC (best chain; posterior mean with 5–95% interval), the `RPVG` estimate used to initialize `TrailMix`, and `kallisto`. The x-axis is the simulated (nominal) proportion of the less abundant source and the y-axis the corresponding estimate; each point is labeled with the share of L2 in the simulated mixture, the dashed line is the identity and the RMSE over the 21 mixtures is given in each panel.

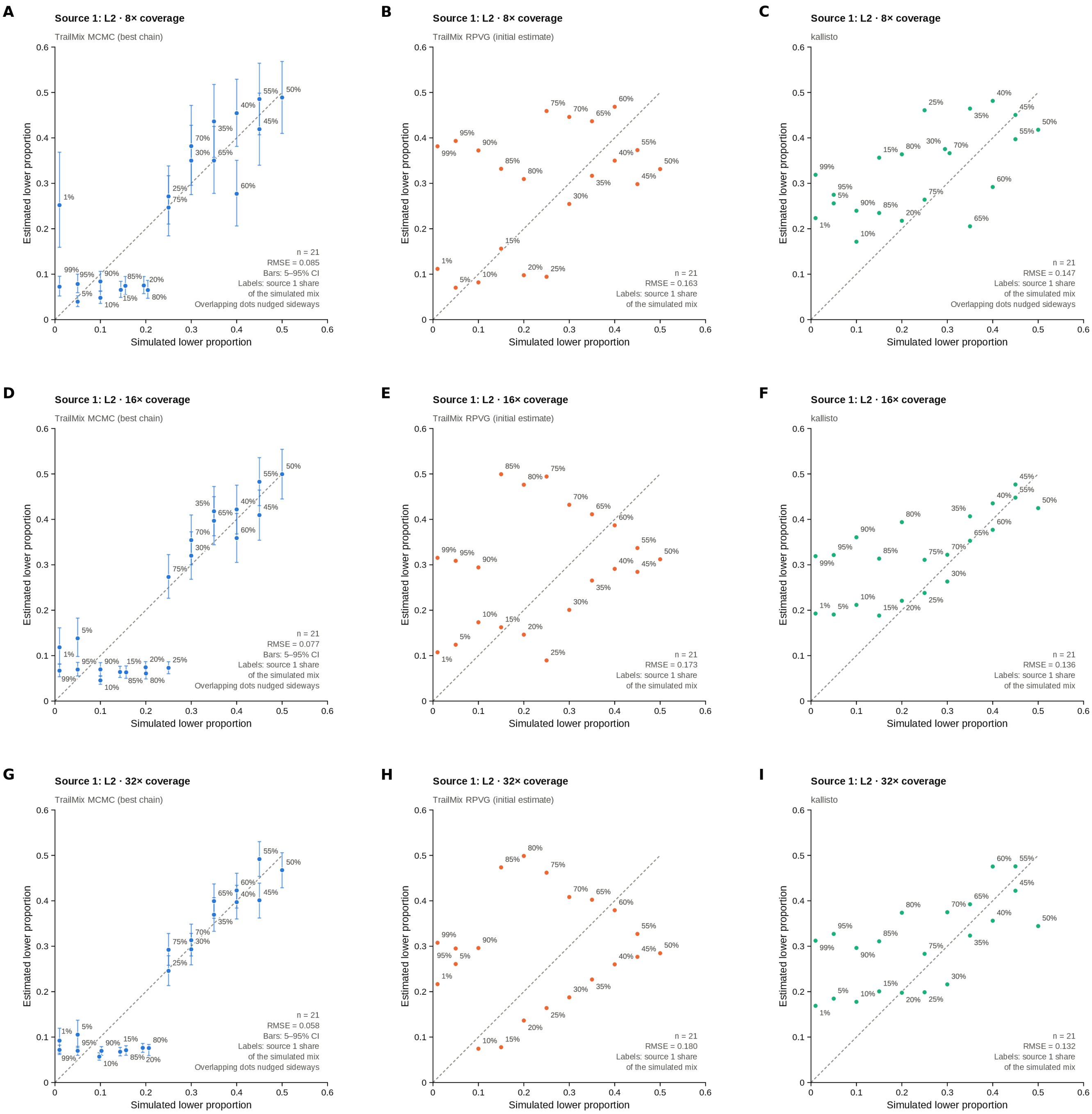


Figure S124: **Estimated versus simulated minor mixture proportion for modern human L2 (AF381981.1) mixed with U5a2a at 21 proportions (1–99%), at nominal mean coverage depths of 8X, 16X, 32X (rows).** Columns: `TrailMix` MCMC (best chain; posterior mean with 5–95% interval), the `RPVG` estimate used to initialize `TrailMix`, and `kallisto`. The x-axis is the simulated (nominal) proportion of the less abundant source and the y-axis the corresponding estimate; each point is labeled with the share of L2 in the simulated mixture, the dashed line is the identity and the RMSE over the 21 mixtures is given in each panel.

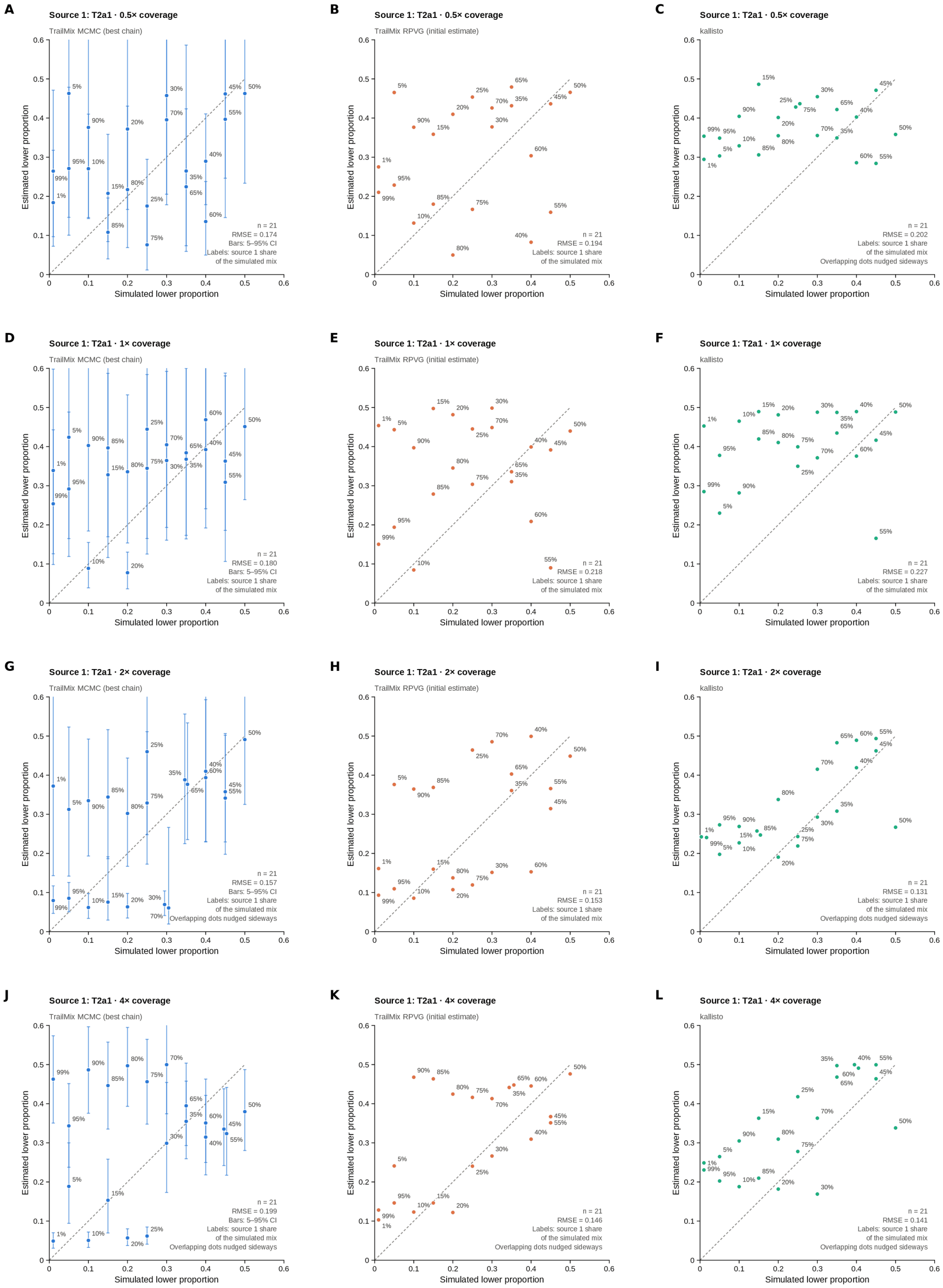


Figure S125: **Estimated versus simulated minor mixture proportion for modern human T2a1 (KY671094.1) mixed with U5a2a at 21 proportions (1–99%), at nominal mean coverage depths of 0.5X, 1X, 2X, 4X (rows).** Columns: `TrailMix` MCMC (best chain; posterior mean with 5–95% interval), the `RPVG` estimate used to initialize `TrailMix`, and `kallisto`. The x-axis is the simulated (nominal) proportion of the less abundant source and the y-axis the corresponding estimate; each point is labeled with the share of T2a1 in the simulated mixture, the dashed line is the identity and the RMSE over the 21 mixtures is given in each panel.

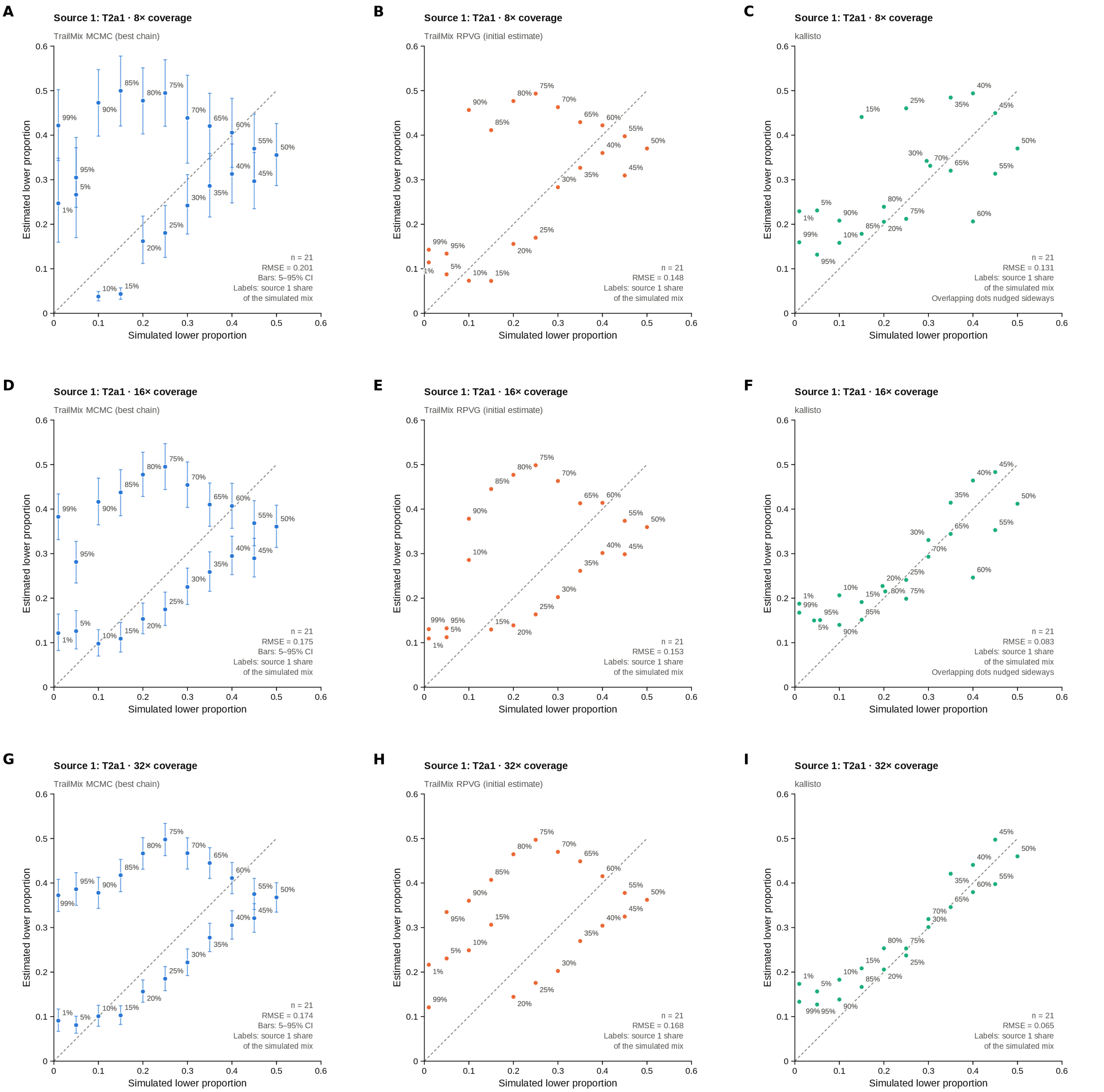


Figure S126: **Estimated versus simulated minor mixture proportion for modern human T2a1 (KY671094.1) mixed with U5a2a at 21 proportions (1–99%), at nominal mean coverage depths of 8X, 16X, 32X (rows).** Columns: `TrailMix` MCMC (best chain; posterior mean with 5–95% interval), the `RPVG` estimate used to initialize `TrailMix`, and `kallisto`. The x-axis is the simulated (nominal) proportion of the less abundant source and the y-axis the corresponding estimate; each point is labeled with the share of T2a1 in the simulated mixture, the dashed line is the identity and the RMSE over the 21 mixtures is given in each panel.

#### S8 Empirical Samples

Table S4 lists the nine sediment libraries analyzed in the main text and here. Their reads were mapped to the rCRS, PCR duplicates were removed and reads shorter than 35 bp were discarded; damage rates were estimated for each library with `bam2prof` [12]. `TrailMix` was run with $k = 1$ and $k = 2$ (100,000 iterations, 5 chains). The $k = 1$ runs of all nine libraries are shown in Supplementary Figure S127 and their $k = 2$ runs in Supplementary Figures S128–S130, with the full tree; the $k = 2$ runs of M76, E251, E256, Ch-3007, GE-I-B09 and GE-I-A4 are also shown, with collapsed trees, in main Figures 6 and 7.

| Sample | Library | Site | Publication | Reads | Mean length (bp) | Coverage |
|---|---|---|---|---|---|---|
| M76 | A20702 | Denisova Cave | [24] | 4,785 | 49.1 | 14.2X |
| E221 | M5300 | Denisova Cave | [24] | 4,154 | 70.2 | 17.6X |
| E227 | M5410 | Denisova Cave | [24] | 5,047 | 75.6 | 23.0X |
| E249 | M5438 | Denisova Cave | [24] | 4,571 | 77.4 | 21.4X |
| E251 | M5440 | Denisova Cave | [24] | 4,756 | 78.0 | 22.4X |
| E256 | M5450 | Denisova Cave | [24] | 4,757 | 76.7 | 22.0X |
| Ch-3007 | A15850 | Chagyrskaya Cave | [23] | 11,644 | 64.4 | 45.3X |
| GE-I-B09 | A16044 | Galería de las Estatuas | [23] | 15,932 | 62.2 | 59.8X |
| GE-I-A4 | A11423 | Galería de las Estatuas | [23] | 11,241 | 59.1 | 40.1X |

Table S4: **Empirical sediment libraries.** Reads: number of reads of at least 35 bp after duplicate removal; coverage: total number of bases divided by the length of the rCRS.

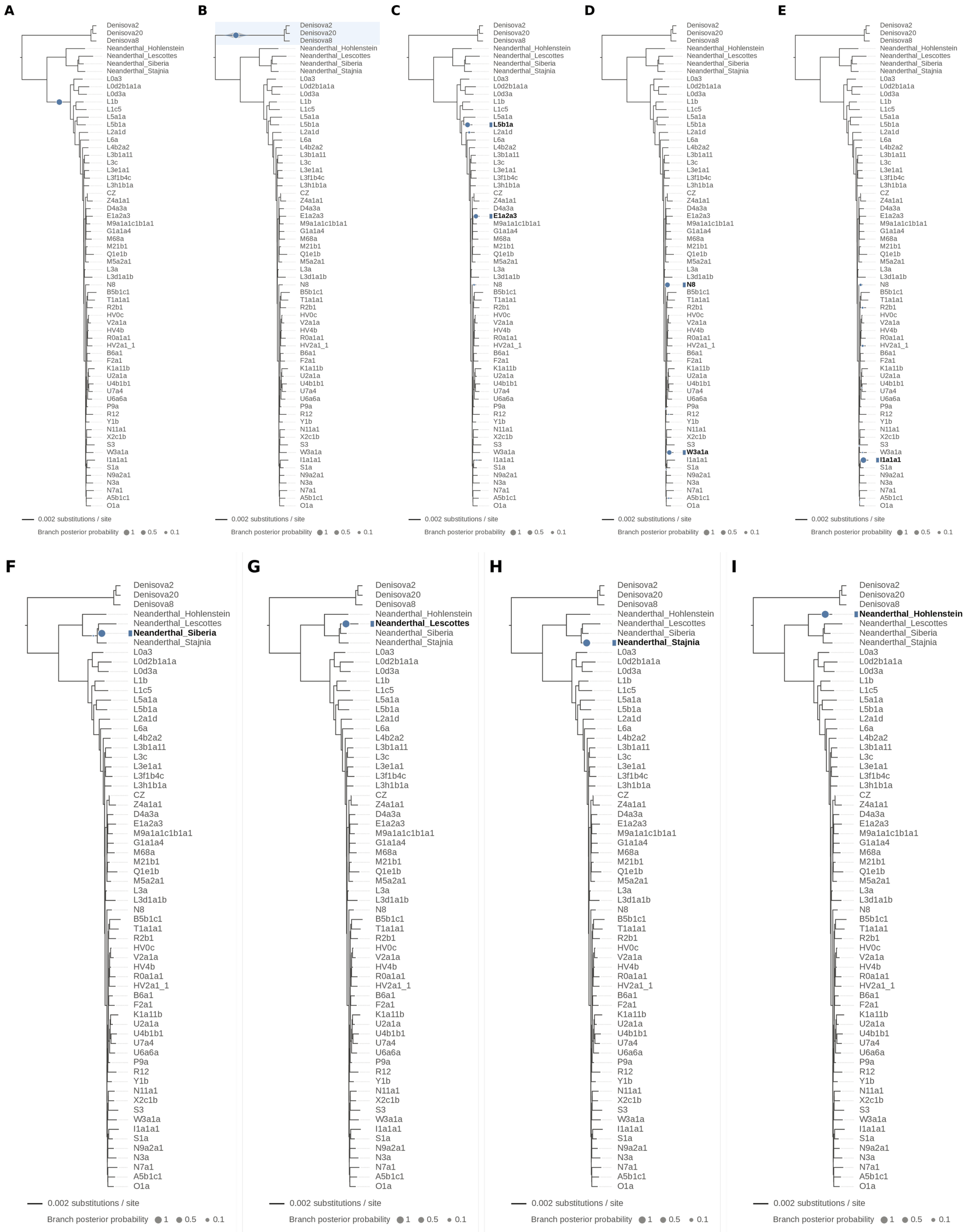


Figure S127: **Single-source ($k = 1$) `TrailMix` runs on the nine empirical sediment libraries.** (A) M76 (library A20702), (B) E221 (M5300), (C) E227 (M5410), (D) E249 (M5438), (E) E251 (M5440) and (F) E256 (M5450) from Denisova Cave [24]; (G) Ch-3007 (A15850) from Chagyrskaya Cave, and (H) GE-I-B09 (A16044) and (I) GE-I-A4 (A11423) from the Galería de las Estatuas [23]. Each panel shows the phylogenetic placement posterior of the best of five chains (100,000 iterations each); the five chains agree on the modal branch in every run except E227 and E249, in which one chain differs. When the sample is a mixture, a single source is forced onto a compromise branch: M76 is placed on the branch separating the archaic hominins from the modern humans (Node44), and E227, a mixture of a Denisovan and a Neanderthal with $k = 2$, on the branch leading to L5b1a (haplogroup L5); the lineages resolved with $k = 2$ are shown in main Figure 6 and Supplementary Figures S128–S130. E221 is placed on the stem of the Denisovan clade (Node45), E249 on the branch leading to N8 and E251 on the branch leading to I1a1a1 (both macrohaplogroup N), E256 on the branch of the Denisova 17 Neanderthal (labeled Neanderthal_Siberia), Ch-3007 on the Les Cottés branch, GE-I-B09 on the Stajnia branch and GE-I-A4 on the Hohlenstein-Stadel branch.

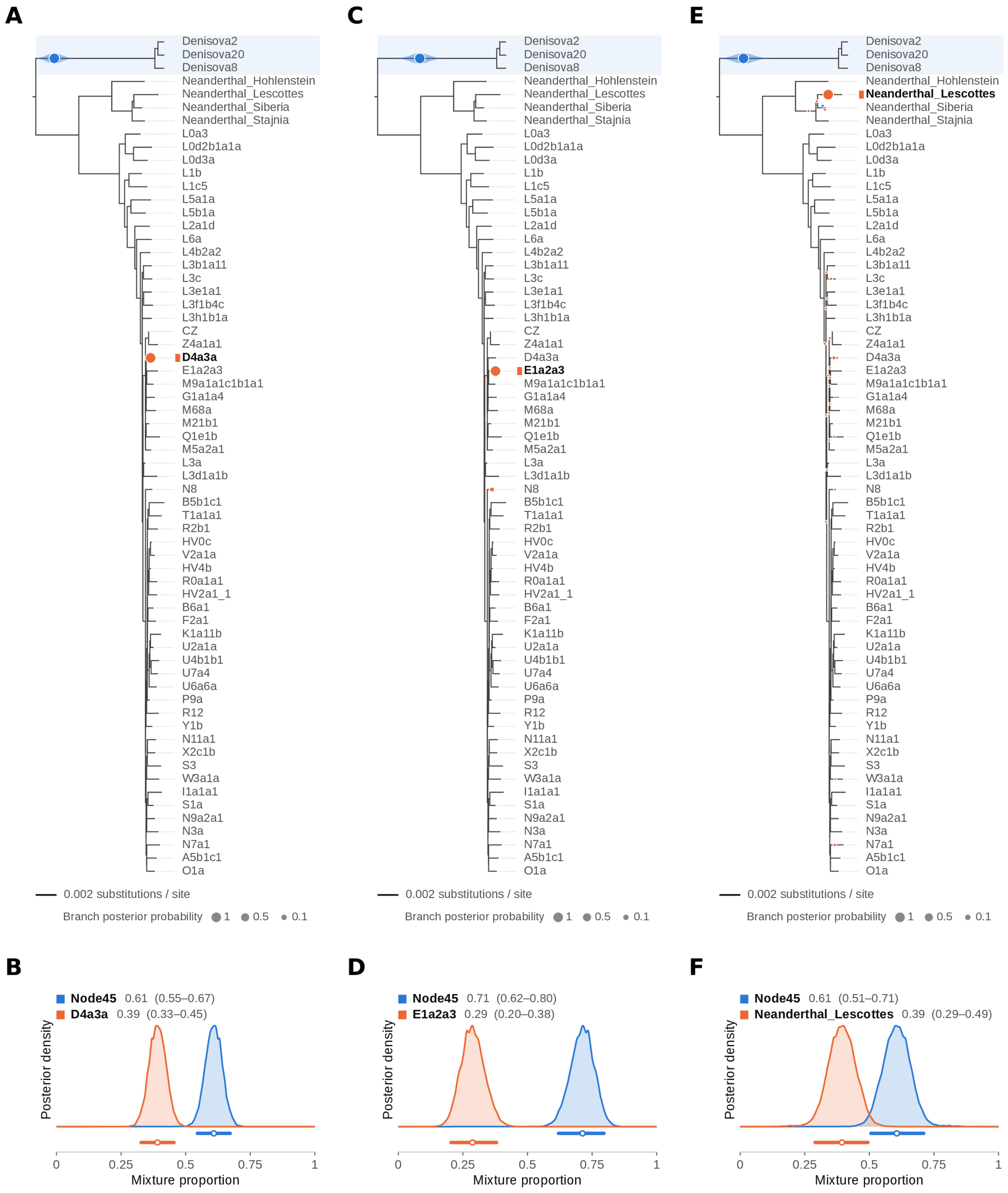


Figure S128: **Two-source ($k = 2$) `TrailMix` runs on the Denisova Cave sediment libraries M76, E221 and E227 [24].** (A,B) M76 (library A20702); (C,D) E221 (M5300); (E,F) E227 (M5410). Top: phylogenetic placement posterior of both sources on the full tree for the best of five chains (100,000 iterations each); bottom: posterior densities of the mixture proportions $\Theta$ with median and 95% credible interval. M76 is deconvolved into a deep lineage on the stem of the Denisovan clade (Node45; $\Theta = 0.61$ [0.55, 0.67]) and a modern human on the D4a3a branch (macrohaplogroup M; 0.39 [0.33, 0.46]); this run is also shown, with a collapsed tree, in main Figure 6A–B. E221 is deconvolved into a lineage on the stem of the Denisovan clade (0.71 [0.62, 0.80]) and a modern human on the E1a2a3 branch (macrohaplogroup M; 0.29 [0.20, 0.38]), and E227 into a lineage on the stem of the Denisovan clade (0.61 [0.51, 0.71]) and a Neanderthal on the Les Cottés branch (0.39 [0.29, 0.49]). For all three libraries, all five chains converge to the same pair of lineages.

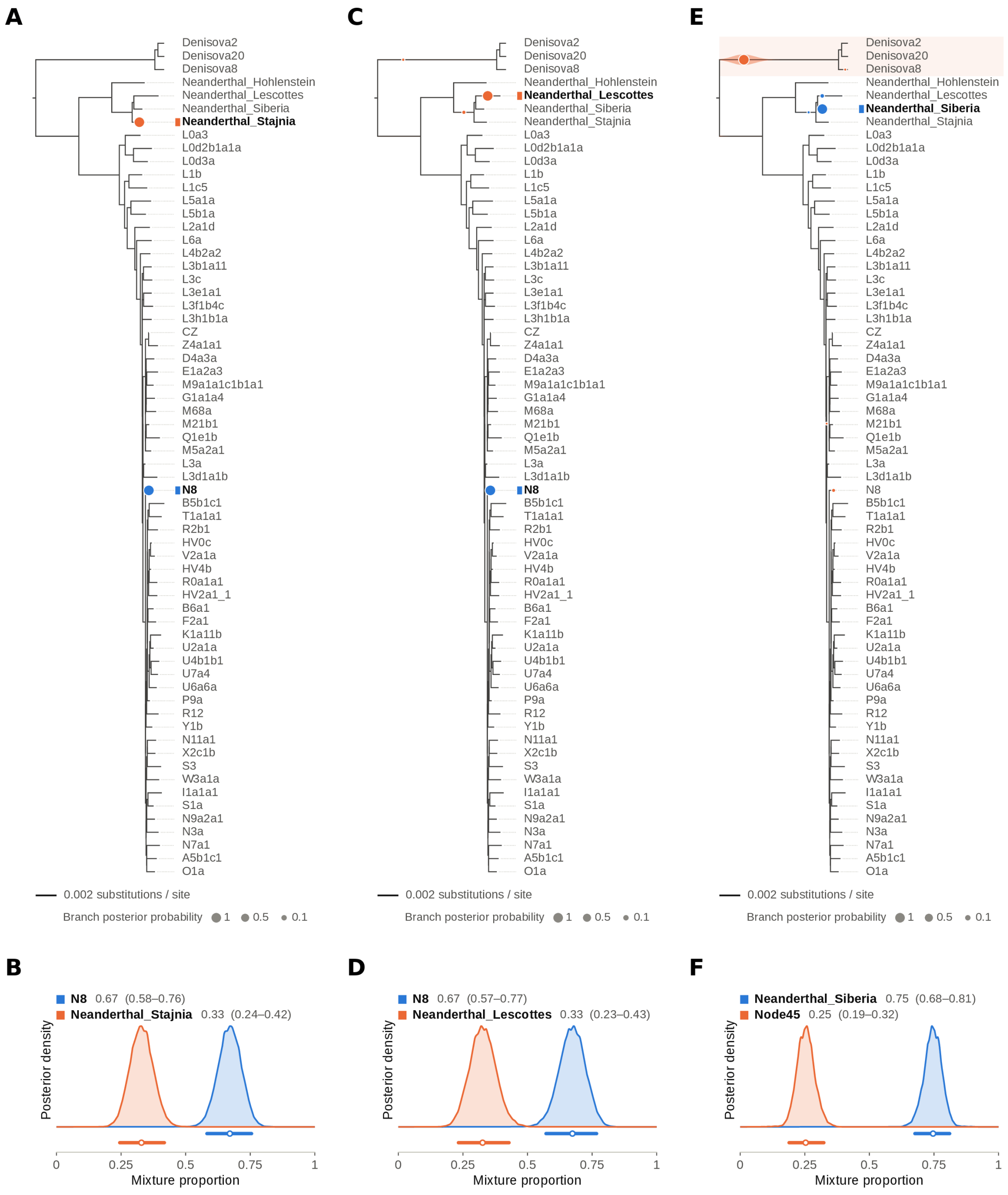


Figure S129: **Two-source ($k = 2$) `TrailMix` runs on the Denisova Cave sediment libraries E249, E251 and E256 [24].** (A,B) E249 (library M5438); (C,D) E251 (M5440); (E,F) E256 (M5450). Top: phylogenetic placement posterior of both sources on the full tree for the best of five chains (100,000 iterations each); bottom: posterior densities of the mixture proportions $\Theta$ with median and 95% credible interval. E249 is deconvolved into a modern human on the N8 branch ($\Theta$ = 0.67 [0.58, 0.76]) and a Neanderthal on the Stajnia branch (0.33 [0.25, 0.42]); E251 into a modern human on the N8 branch (0.67 [0.57, 0.77]) and a Neanderthal on the Les Cottés branch (0.33 [0.23, 0.43]); and E256 into a Neanderthal on the branch of the Denisova 17 Neanderthal (labeled Neanderthal_Siberia; 0.75 [0.68, 0.81]) and a lineage on the stem of the Denisovan clade (Node45; 0.25 [0.19, 0.32]). The runs of E251 and E256 are also shown, with collapsed trees, in main Figure 6C–F. For all three libraries, all five chains converge to the same pair of lineages.

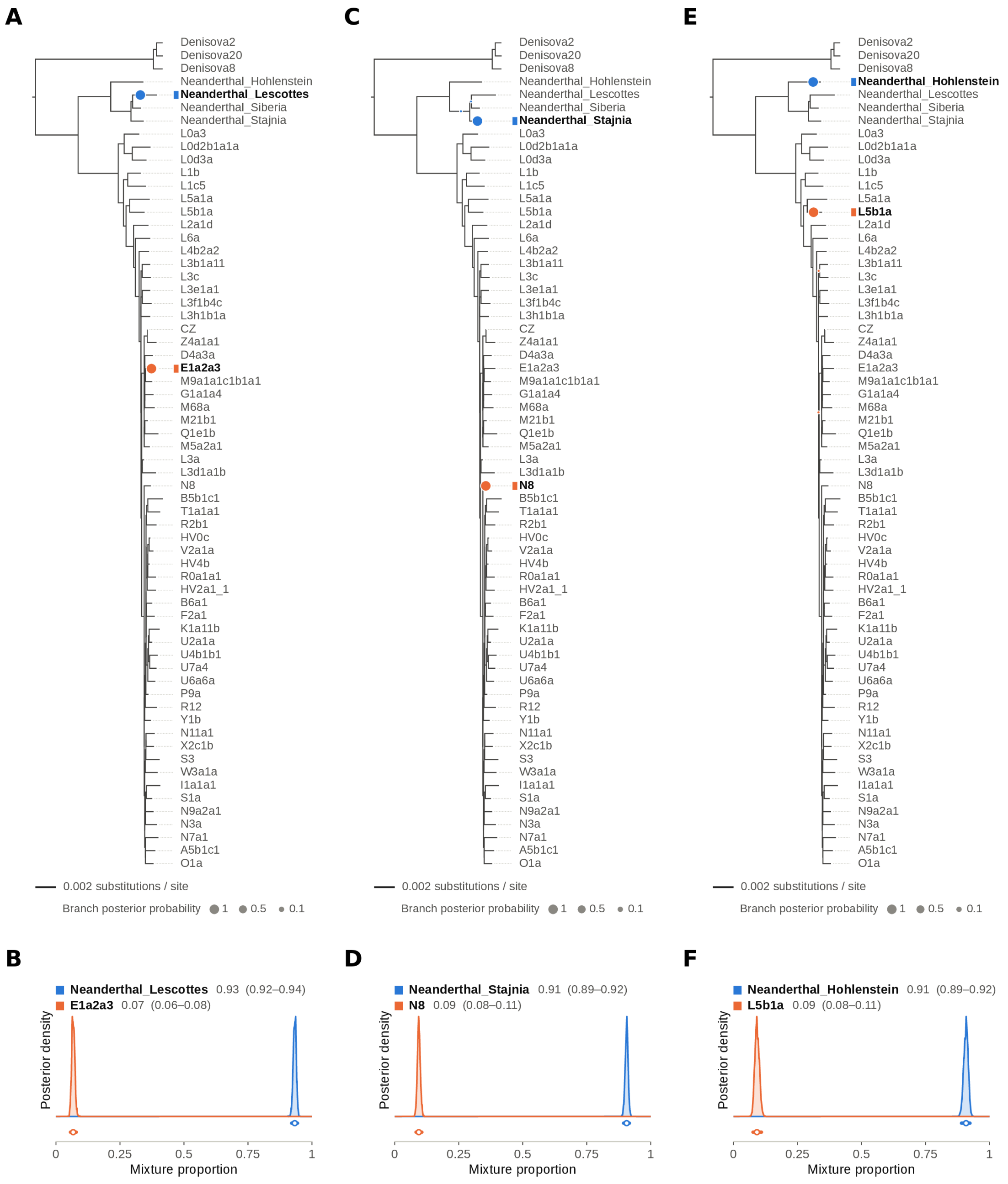


Figure S130: **Two-source ($k = 2$) `TrailMix` runs on the sediment libraries Ch-3007 from Chagyrskaya Cave and GE-I-B09 and GE-I-A4 from the Galería de las Estatuas [23].** (A,B) Ch-3007 (library A15850); (C,D) GE-I-B09 (A16044); (E,F) GE-I-A4 (A11423). Top: phylogenetic placement posterior of both sources on the full tree for the best of five chains (100,000 iterations each); bottom: posterior densities of the mixture proportions $\Theta$ with median and 95% credible interval. Ch-3007 is deconvolved into a Neanderthal on the Les Cottés branch ($\Theta = 0.93$ [0.92, 0.94]) and a modern human on the E1a2a3 branch (macrohaplogroup M; 0.07 [0.06, 0.08]); GE-I-B09 into a Neanderthal on the Stajnia branch (0.91 [0.89, 0.92]) and a modern human on the N8 branch (0.09 [0.08, 0.11]); and GE-I-A4 into a Neanderthal on the Hohlenstein-Stadel branch (0.91 [0.89, 0.92]) and a modern human on the L5b1a branch (haplogroup L5; 0.09 [0.08, 0.11]). All five chains converge to the same pair of lineages, except for one chain of Ch-3007. These runs are also shown, with collapsed trees, in main Figure 7.

### References


[1] Nicolas L Bray et al. “Near-optimal probabilistic RNA-seq quantification”. In: *Nature Biotechnology* 34.5 (2016), pp. 525–527. DOI: 10.1038/nbt.3519.

[2] Samantha Brown et al. “The earliest Denisovans and their cultural adaptation”. In: *Nature Ecology & Evolution* 6.1 (2022), pp. 28–35.

[3] Matei David et al. “SHRiMP2: sensitive yet practical short read mapping”. In: *Bioinformatics* 27.7 (2011), pp. 1011–1012.

[4] Miroslava Derenko et al. “Origin and post-glacial dispersal of mitochondrial DNA haplogroups C and D in northern Asia”. In: *PloS ONE* 5.12 (2010), e15214.

[5] Jordan M Eizenga. *Graph Methods for Computational Pangenomics*. University of California, Santa Cruz, 2021.

[6] Erik Garrison et al. “Building Pangenome Graphs”. In: *Nature Methods* (2024), pp. 1–5.

[7] Erik Garrison et al. “Variation Graph Toolkit Improves Read Mapping by Representing Genetic Variation in the Reference”. In: *Nature Biotechnology* 36.9 (2018), pp. 875–879.

[8] Torsten Günther et al. “Ancient Genomes Link Early Farmers from Atapuerca in Spain to Modern-day Basques”. In: *Proceedings of the National Academy of Sciences* 112.38 (2015), pp. 11917–11922.

[9] Weichun Huang et al. “ART: a Next-generation Sequencing Read Simulator”. In: *Bioinformatics* 28.4 (2012), pp. 593–594.

[10] Kazutaka Katoh et al. “MAFFT: a Novel Method for Rapid Multiple Sequence Alignment Based on Fast Fourier Transform”. In: *Nucleic Acids Research* 30.14 (2002), pp. 3059–3066.

[11] Johannes Köster and Sven Rahmann. “Snakemake—a Scalable Bioinformatics Workflow Engine”. In: *Bioinformatics* 28.19 (2012), pp. 2520–2522.

[12] Louis Kraft et al. “AdDeam: a fast and scalable tool for estimating and clustering reference-level damage profiles”. In: *Bioinformatics* 41.8 (2025), btaf407.

[13] Heng Li et al. “The Sequence Alignment/Map Format and SAMtools”. In: *Bioinformatics* 25.16 (2009), pp. 2078–2079.

[14] Nicole Maca-Meyer et al. “Major genomic mitochondrial lineages delineate early human expansions”. In: *BMC Genetics* 2.1 (2001), p. 13.

[15] Fabrizio Mafessoni et al. “A high-coverage Neandertal genome from Chagyrskaya Cave”. In: *Proceedings of the National Academy of Sciences* 117.26 (2020), pp. 15132–15136.

[16] Boris Malyarchuk et al. “Mitogenomic diversity in Russians and Poles”. In: *Forensic Science International: Genetics* 30 (2017), pp. 51–56.

[17] Lam-Tung Nguyen et al. “IQ-TREE: a fast and effective stochastic algorithm for estimating maximum-likelihood phylogenies”. In: *Molecular Biology and Evolution* 32.1 (2015), pp. 268–274.

[18] Gabriel Renaud, Udo Stenzel, and Janet Kelso. “leeHom: Adaptor Trimming and Merging for Illumina Sequencing Reads”. In: *Nucleic Acids Research* 42.18 (2014), e141–e141.

[19] Gabriel Renaud et al. “gargammel: a Sequence Simulator for Ancient DNA”. In: *Bioinformatics* 33.4 (2017), pp. 577–579.

[20] Joshua Rubin et al. “SAFARI: Pangenome Alignment of Ancient DNA Using Purine/Pyrimidine Encodings”. In: *NAR Genomics and Bioinformatics* 7.4 (2025), lqaf170.

[21] Jonas A Sibbesen et al. “Haplotype-aware pantranscriptome analyses using spliced pangenome graphs”. In: *Nature Methods* 20.2 (2023), pp. 239–247.

[22] Jouni Sirén et al. “Pangenomics Enables Genotyping of Known Structural Variants in 5202 Diverse Genomes”. In: *Science* 374.6574 (2021), abg8871.

[23] Benjamin Vernot et al. “Unearthing Neanderthal population history using nuclear and mitochondrial DNA from cave sediments”. In: *Science* 372.6542 (2021), eabf1667.

[24] Elena I Zavala et al. “Pleistocene sediment DNA reveals hominin and faunal turnovers at Denisova Cave”. In: *Nature* 595.7867 (2021), pp. 399–403.
